# FACET: Criterion-level face-validity of 32,535 mouse genotypes across five DSM-5-TR psychiatric disorders

**DOI:** 10.64898/2026.11.21.768352

**Authors:** An Le, Donovan Nguyen, Andrew R. Craig, Jonathan Hess, Chunyu Liu

## Abstract

A mouse is typically declared a model of a psychiatric disorder from a few behavioral tests, while a diagnosis requires several criteria together, so a model’s scope of inference is seldom stated. FACET makes that scope explicit by auditing the face validity of every Mouse Genome Informatics (MGI) genotype against the criteria of DSM-5-TR. Each criterion is resolved into human phenotype terms (anchors), and a large language model (LLM) blind to genes judges mouse terms against each anchor. We report two main measures: Diagnostic Coverage, the number of criteria a genotype’s annotated phenotype profile models, and a support score, how rarely a human-mouse phenotype match is seen for a random profile of equal size. Across 32,535 genotypes and five disorders (autism, schizophrenia, intellectual disability, major neurocognitive disorder and major depressive disorder), most curated models recapitulate part of their definition, and no profile can meet full diagnosis for autism or schizophrenia. Ten criteria are modeled by no curated model. The support score ranks curated models above the background (support AUROC 0.96, 0.68, 0.75 and 0.67 for autism, schizophrenia, intellectual disability and neurocognitive disorder). Outside autism, behavioral annotation count alone separates the classes, and at equal depth of behavioral annotation only autism shows a clear separation (0.90). Nearly every curated model’s profile comes from the paper that generated it, so label recovery is not independent validation. The LLM agrees with a single expert about as often as two experts agree with each other (77 against 79%). In an interim expert study, eight human raters confirm 88% of the pairs it refuses and 66% of the pairs it admits. FACET therefore bounds each model’s inference by the criteria its profile models.

**Author summary:** Thousands of genetically altered mice are used to study psychiatric disorders, on the premise that the mouse reproduces part of the human condition. A mouse can show a measurable analog of some symptoms, such as anhedonia, but not of others, such as hallucinations. A mouse is usually called a model after a few behavioral tests, while a diagnosis requires several symptoms together. What a model is meant to recapitulate and what it has been shown to recapitulate can therefore differ, and the difference is rarely measured. We built FACET to ask, for every mouse genotype in the Mouse Genome Informatics catalogue, which diagnostic criteria of five psychiatric disorders its recorded phenotypes model. FACET resolves each criterion into human phenotype terms and uses a large language model under strict constraints to decide whether a recorded mouse phenotype models the feature each term names. Most curated models recapitulate only part of their disorder’s definition. No genotype’s recorded phenotypes can meet the full diagnostic rule for schizophrenia or autism. Five criteria have no admitted mouse analog, one has no mouse term and four are modeled only by mice that nobody has called a model of the disorder. In an interim rating study, FACET agreed with an expert about as often as two experts agreed with each other. The experts agreed with most of its refusals and with two in three of its admissions. The released catalogue shows which criteria a candidate model does and does not recapitulate. It also points to existing mice that carry criteria of a disorder they were never proposed for.

## Introduction

Mice carrying mutations in psychiatric risk genes are the main experimental system for studying what those genes do to the brain and behavior, yet findings from animal models of psychiatric disorders translate poorly to patients [1,2]. One explanation is the mismatch between how a psychiatric disorder is diagnosed in a person and how it is modeled in a mouse. For most disorders in the Diagnostic and Statistical Manual of Mental Disorders, fifth edition, text revision (DSM-5-TR), the etiology is unknown and no biomarker or objective test exists, thus, diagnosis rests on symptoms and their course [3,6]. A patient receives diagnosis only when several distinct criteria hold together, often in specified combinations and with clinically significant impairment [3]. A mouse genotype, by contrast, is often declared a model based on a few behavioral tests, without meeting any combination of the criteria required for a human diagnosis [2]. The resemblance between the symptoms a model displays and those of the disorder is its face validity [4,7], and a model of a syndromic disorder should be evaluated on it. Predictive validity (response to the drugs that work in patients) is the usual alternative standard. It does not replace face validity, since the screens that find psychiatric drugs detect compounds resembling existing drugs and are blind to the disorder itself [6]. Some criteria, such as disorganized speech or delusions, have no mouse analog, and most of the remaining criteria are not recorded for a given genotype. Much of the preclinical record in psychiatry therefore rests on mice that capture only part of the definition they are named for. A systematic criterion-level evaluation is needed to measure how much, one that reads every criterion of the definition and covers every genotype in the inventory.

Such an evaluation does not require any single genotype to reproduce the full syndrome, in agreement with the view that no model is expected to reproduce a full syndrome [5–7]. The field has turned to specific behaviors and endophenotypes (heritable traits intermediate between genotype and syndrome) [8,9] and to multiple-hit models that combine risk factors to reflect the multifactorial etiology of a disorder [10]. A narrower question remains open across the thousands of genetically altered mice already created: which criteria of a disorder’s diagnosis does a given mouse recapitulate? Criteria are the natural benchmark because model claims are made in reference to them and the manual itself is organized around them. Polythetic criteria are heterogeneous by construction, so a criterion-level record provides a floor for regrouping under symptom dimensions or Research Domain Criteria constructs, without re-judging any pair. Answering this would allow existing models to be compared on an equal footing by the extent and composition of the diagnostic definition each recapitulates. It would also inform the choice of complementary genotypes in multiple-hit designs. Evaluation against the diagnostic criteria has been proposed [11,12] and contested [13], and manual mappings of criteria to rodent phenotypes have been drawn up for depression [14,15], autism [16,17] and schizophrenia [18]. The closest precedent aligned fourteen behavioral measures to the DSM-5 autism criteria and validated the alignment by factor analysis on 65 mice [19]. Each of these was built for a single disorder and a few models.

The records needed for such an evaluation are already public, though not utilized in this manner. Mouse Genome Informatics (MGI) holds curated phenotype annotations for more than 70,000 genotypes, stored as Mammalian Phenotype (MP) terms [20], and curates genotypes as disease models from the papers that describe them. These curated models serve as the reference labels in this paper, and an uncurated genotype carries no label of either type. The five disorders studied are autism spectrum disorder (ASD), schizophrenia (SCZ), intellectual developmental disorder (intellectual disability, IDD), major neurocognitive disorder (NCD), and major depressive disorder (MDD), the last as an exploratory disorder. On the human side, symptoms are recorded in the Human Phenotype Ontology (HPO). Face validity at scale therefore becomes a problem of cross-species phenotype correspondence, decided for every symptom and every recorded mouse phenotype, in uncurated genotypes as well as curated ones. Existing tools for cross-species correspondence were not built for this question. PhenoDigm [21] and the exome-prioritization tools built on it [22] were designed for Mendelian disease, and they compare terms by shared ontology ancestry, logical definitions and annotation frequency without reading the terms’ free-text definitions. Two psychiatric terms describing the same behavior can therefore score as unrelated, and the increased and decreased forms of a behavior score alike. In the HPO annotation release [23], almost none of the psychiatric symptom terms are attached to the diagnoses themselves (S1 Text), so no human profile of schizophrenia exists for a mouse phenotype to be compared with. Deciding correspondence at this scale therefore needs a tool that reads phenotype descriptions as a human evaluator would and whose readings can be checked against human evaluators.

This paper presents FACET (Face-validity Assessment via Criterion-Anchored-LLM Extraction and Translation), a pipeline that credits each genotype in a phenotype catalogue with the diagnostic criteria its recorded phenotypes model. It breaks each diagnostic criterion into observable human features, asks which mouse phenotypes correspond to those features, and records which criteria each genotype’s annotated phenotypes model. Each feature is represented by an HPO term, its anchor. We used FACET to re-evaluate MGI’s psychiatric model inventory. The anchor sets, the mouse taxonomy and the per-criterion catalogue it produces are released. Its unit of output is the admitted pair: a human anchor term and a mouse phenotype term that the LLM accepts as a direct or close analog of it. Admission means the analog is strong enough to count as modeling the feature; every other judged pair is refused, and no whole-profile similarity method returns such pairs. A genotype is credited with a criterion when at least one of its recorded MP phenotypes forms an admitted pair with an anchor of that criterion. Large language models (LLMs) supply the reading the pipeline needs [24,25], having outperformed crowd workers on text annotation [26], populated biomedical knowledge bases [27] and drafted ontology terms and definitions [28]. Deciding whether a human term and a mouse term describe the same observable is a reading task with a narrow answer, which an LLM can perform under constraints. We restricted what the LLM could read and what it could return to increase transparency. We also validated FACET’s verdicts against alternate LLMs and a panel of human experts. FACET quantifies the face validity of psychiatric animal models at scale, applied here to the mouse. FACET reports two measures per genotype, Diagnostic Coverage (the criteria its recorded phenotypes model) and the FACET support score (how unusually well its recorded phenotypes match a disorder’s criteria compared with random phenotype profiles of the same size), and we evaluate how well each separates curated models from other genotypes and what that separation rests on. We applied FACET to ASD, SCZ, IDD, NCD, and MDD across the 32,535 MGI genotypes carrying four or more annotated phenotype terms. These disorders account for most of the curated psychiatric models in MGI and span definitions whose cardinal symptoms range from directly observable in a mouse to having no mouse analog. To our knowledge, this is the first criterion-level measurement of face validity at the scale of a mouse phenotype catalogue. The three released products are the HPO disorder anchor sets, the criterion-mapped taxonomy of the mouse vocabulary and a searchable per-criterion catalogue of the MGI genotypes carrying a phenotype that models any DSM-5-TR criterion (https://dsm-5-map-mouse.vercel.app).

## Results

### FACET records which criteria every genotype’s phenotypes model

FACET breaks each diagnostic criterion into observable human features, asks which mouse phenotypes correspond to those features, and records which criteria each genotype’s annotated phenotypes model. It reads the diagnostic criteria of a disorder and the phenotype annotations of a catalogue in three steps: anchoring, judgment, and aggregation (Figure 1, Methods). Anchoring turns each criterion into a set of HPO terms (its anchors), each representing one observable feature of the criterion, and a separate LLM step blinded to the anchors assigns each MP term to the criteria it could serve (mouse criterion mapping). Judgment retrieves candidate MP terms for each anchor by text and ontology similarity and has the LLM grade how well each candidate models the anchor. Each candidate pair carries a similarity score between 0 and 1, the combined retrieval score, built from text-embedding, lexical and ontology similarity, and the LLM’s verdict adjusts it into a final pair score (Methods). A judged pair is admitted when the LLM finds the mouse term strictly eligible, grades it a direct or close analog, and an automatic direction check passes. It is refused otherwise. Aggregation credits each admitted term, and every more specific term beneath it in MP, to the genotypes annotated with them. A criterion unit is one countable symptom or functional component of the diagnostic definition (the phenotypic content of one numbered criterion that counts toward the diagnosis). A genotype is credited with a unit, and recapitulates it, when at least one of its annotated phenotypes is an admitted term for one of the unit’s anchors. The bottom row of Figure 1 follows one criterion through the three steps.

**Figure 1.**
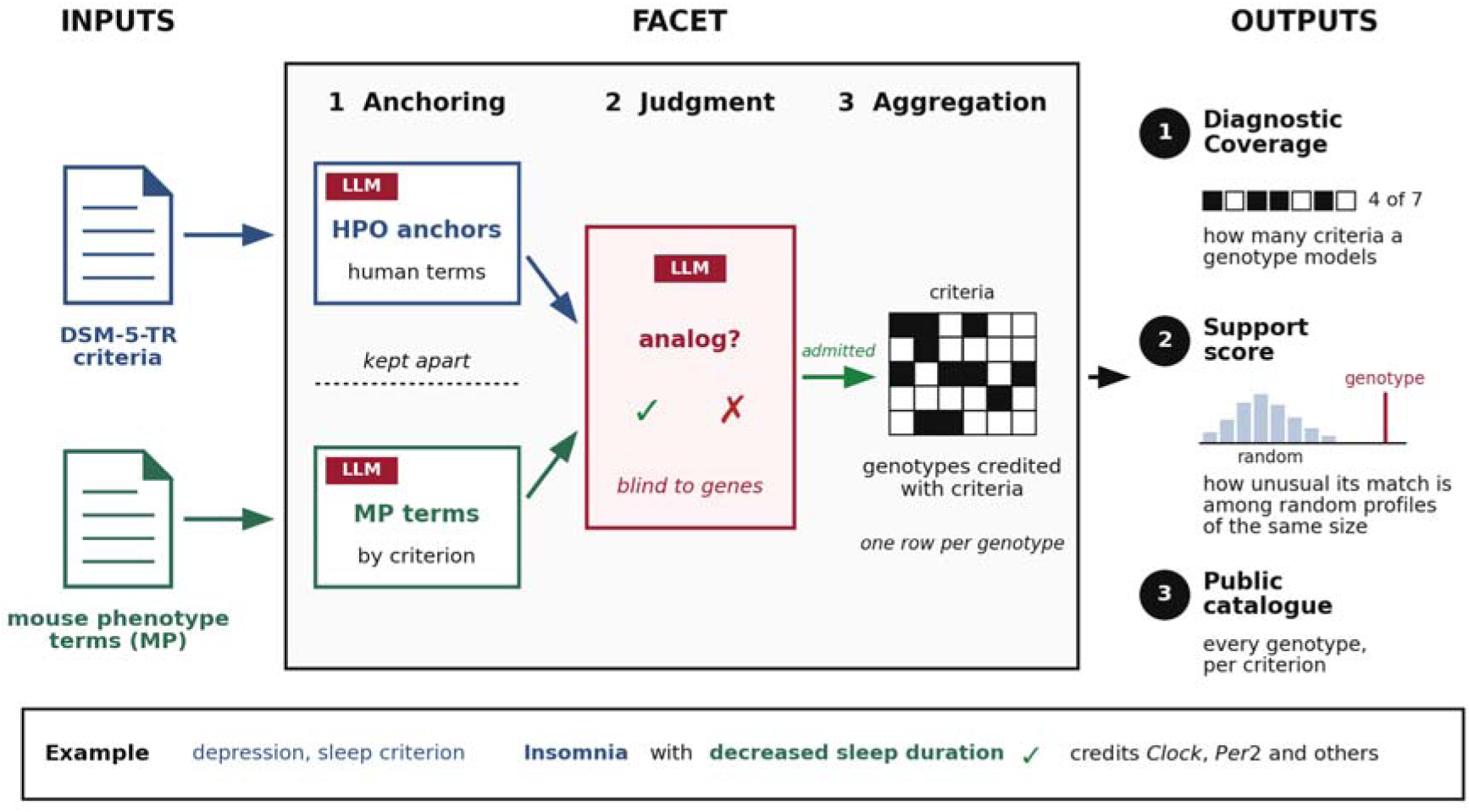
FACET from inputs to outputs. Crimson marks the steps decided by the LLM. Diagnostic criteria are resolved into HPO anchors and mouse phenotype terms are assigned to criteria separately, and the two meet only at judgment. There the LLM admits or refuses each candidate pair without seeing genes or genotypes. Aggregation credits genotypes with the criteria their admitted terms model. The grid stands for the table of credits, one row per genotype and one column per criterion. The support score places a genotype’s match among those of random profiles of the same size. The bottom row follows one example. MDD criterion A4 (sleeping too little or too much on most days, paraphrased) resolves to six HPO anchors. For the Insomnia anchor, 37 candidate MP terms are retrieved and the five highest scoring are judged in one call. The admitted term decreased sleep duration credits the genotypes annotated with it or with a more specific descendant. Values are from the primary analysis run.

Two measures summarize the credited phenotypes of a genotype. Diagnostic Coverage is the number of a disorder’s scored criterion units that the genotype recapitulates; that is, the units for which it has at least one admitted mouse phenotype. The FACET support score measures how unusually well the genotype’s recorded phenotypes match a disorder’s criteria compared with random phenotype profiles containing the same number of annotations. A genotype’s match to the criteria is computed from these similarity scores: the final pair scores of its admitted matches are combined within each anchor and averaged over the disorder’s anchors (Methods). The support score asks whether this match is higher than that of 10,000 random sets of MP terms as large as the genotype’s annotation. It is the negative base-10 logarithm of the share of random sets scoring as high, capped at 4. Each measure is evaluated against the curated models (the genotypes MGI curates as models of the disorder) by an AUROC (area under the receiver operating characteristic curve). An AUROC is the probability that a randomly selected curated model outranks a random background genotype. The coverage AUROC ranks all 32,535 genotypes, and the support AUROC ranks the genotypes carrying a candidate term for the disorder (its evaluation set). The LLM sees no gene, genotype, MGI disease label or curated list at any step. The primary analysis run is the frozen set of LLM verdicts from which every number in this paper is computed, and the primary LLM (gpt-5.4) produced them.

HPO contains psychiatric symptom terms but does not group them under the diagnoses (17 subtype or severity terms beneath the five disorders, five annotated phenotypes for idiopathic schizophrenia and none for MDD), so FACET builds the anchor sets itself. Decomposition of the five DSM-5-TR chapters yielded 174 anchors. Of these, 138 are attributable to a criterion unit and 133 to a scored unit (the five others serve SCZ criterion B, which is not scored). The remaining 41 come from onset, exclusion, duration and associated-feature text (features not typically studied in mice) and are judged but never credited. The 138 attributable anchors are a machine-readable decomposition of the criteria of five disorders into HPO terms and are released (S11 Table). On the mouse side, 11,485 of the 14,750 MP terms are annotated to at least one MGI genotype and are candidates for judgment.

Over the 174 anchors, the pair judgment produced 21,138 verdicts, of which 14,209 lie on anchors of scored criteria and can reach a score (Table 1). The LLM refuses far more candidate pairs than it admits in every disorder, and 420 of the 616 admitted pairs credit genotypes.

**Table 1.** What FACET built and read, per disorder. Curated models are counted among the 32,535 genotypes under the label rules of Methods. Behavioral gives the percent of curated models with at least one annotation at or below abnormal behavior (MP:0004924), and negatives are MGI records stating that a genotype is not a model. Anchors are the HPO terms resolved from the chapter, with the number attributable to a criterion unit in parentheses. Judged and admitted pairs are counted on anchors of scored criteria, and admitted pairs also among all judged pairs.

| Disorder | Curated | Behavioral, % | Negatives | Anchors (attributable) | Scored units | Judged pairs, scored | Admitted, all | Admitted, scored |
|---|---|---|---|---|---|---|---|---|
| ASD | 85 | 97.6 | 1 | 43 (37) | 7 | 4,925 | 145 | 125 |
| SCZ | 42 | 100 | 0 | 41 (15) | 5 | 3,213 | 196 | 50 |
| IDD | 121 | 83.5 | 0 | 25 (21) | 2 | 3,201 | 138 | 108 |
| NCD | 71 | 87.3 | 12 | 28 (28) | 2 | 2,607 | 95 | 95 |
| MDD | 8 | 100 | 1 | 37 (37) | 9 | 263 | 42 | 42 |
| Total | 327 | 26.2 of all 32,535 genotypes | 14 | 174 (138) | 25 | 14,209 | 616 | 420 |

### Diagnostic Coverage: which criteria mice model and which curated models recapitulate

#### Ten criteria are credited to no curated model for three different reasons

Six criteria have no admitted phenotype in any of the 32,535 genotypes (S28 Table). Five are assessed through speech or self-report: delusions, hallucinations and disorganized speech (SCZ A1 to A3) and worthlessness and thoughts of death (MDD A7 and A9). Candidates were retrieved and judged for them and the primary LLM admitted none. The alternate LLMs admit 5 and 41 pairs on SCZ A1 to A3 (Validation), so the zero for those three criteria is a verdict of the primary LLM. The 5 pairs join the delusion and hallucination anchors to latent inhibition and startle terms, and the disordered-thought anchor to cognition, vocalization and latent inhibition terms. The 41 add learning, conditioning, habituation and stereotypy terms and, for disordered thought, eye-movement and brain-morphology terms. None of these pairs is an analog of the symptom itself, and the primary LLM refused all of them (S33 Table). Those admissions were not expert-rated, while the raters agreed with every rated SCZ refusal. Four further criteria are credited to background genotypes but to no curated model. They are psychomotor change, fatigue, and impaired concentration for MDD (A5, A6, and A8) and interference with independence for NCD (B). The ten fall into three kinds (Figure 2B). Five have no admitted analog since the primary LLM judged every candidate and admitted none. One has no mouse term: MP records none of ASD A2’s content. Four are phenotyping gaps, where the mouse vocabulary expresses them and background genotypes carry admitted terms for them, but no curated model carries a recorded abnormal annotation for them. MGI records abnormal results only, so a normal result leaves no trace.

**Figure 2.**
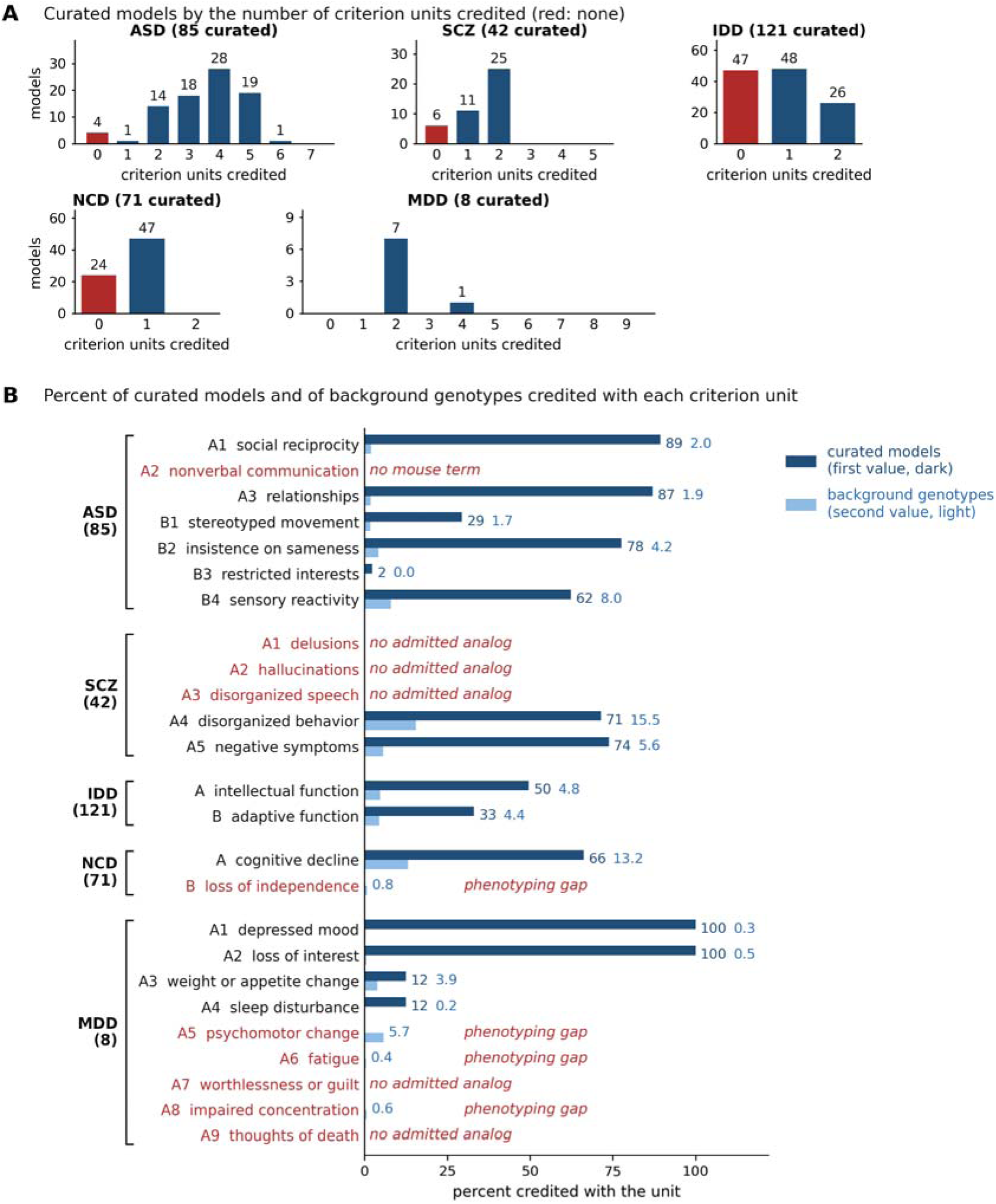
The criterion units credited to curated models, per disorder. (A) Curated models by the number of criterion units credited. Red bars give the models with no admitted phenotype. (B) Percent of curated models and of background genotypes credited with each criterion unit, grouped by disorder (number of curated models in parentheses). Red unit labels mark units credited to no curated model, and red italic text names the kind of gap. No admitted analog: SCZ A1 to A3, MDD A7 and A9, credited to no genotype. No mouse term: ASD A2, excluded by rule. Phenotyping gap: MDD A5, A6 and A8 and NCD B, credited to background genotypes only. The SCZ A5 value is an upper bound (Results). Value labels are percentages, the darker for curated models and the lighter for background genotypes.

SCZ A5 (negative symptoms) is credited through abnormal emotion or affect behavior and bradykinesia, which together reach 1,865 genotypes, and its avolition component has no MP vocabulary at all. The emotion term is a broad class, so curated SCZ models reach A5 through thigmotaxis, exploration, and anxiety-related annotations (S19 Table), the loosest credit in the results. Without it, A5 is credited to no curated model and to 72 background genotypes, and six curated models lose their only credit, so the 31 of 42 below is an upper bound. NCD B is credited through decreased grooming behavior and aphagia under the construct rule that a motor term does not model interference with independence (Methods). Without that rule, impaired coordination alone would credit criterion B to 1,292 genotypes and move the NCD coverage and support AUROCs by 0.014 or less (S1 Text).

#### The annotations of curated models follow the type of tests each field runs

About 7% of all annotations in the genotype set describe behavior, and three in four genotypes carry no behavioral annotation. At least 7.5% of those were assayed by the International Mouse Phenotyping Consortium (IMPC) and found normal, so absence of annotation is not absence of testing (S1 Text, S7 Table). Curated models are the exception: 83.5 to 100% are behaviorally annotated against 26.2% of the set (Table 1), and behavioral terms make up 30 to 61% of their annotations against 7% in the set (Figure 3A). Figure 3 reads the annotations by nine phenotype classes that are MP’s own branches (Methods, S16 Table). Which behavioral classes a curated set carries tracks each field’s assays rather than the manual’s criteria (Figure 3B). Nine in ten curated ASD models carry a social term. Curated SCZ models carry cognitive and motor terms more often than social terms, and cognition is an associated feature of SCZ rather than a criterion A symptom. Curated NCD models carry motor and nervous-system morphology terms far more often than the cognitive terms criterion A names, and almost never social or emotion terms. All eight MDD models carry an emotion term, and two carry a motor term, which is the phenotyping gap behind MDD A5. In every disorder, the admitted terms fall almost entirely in the behavioral classes (red dots in Figure 3B). The classes the field assays and the classes the criteria need therefore coincide for ASD and IDD and diverge for SCZ and NCD.

**Figure 3.**
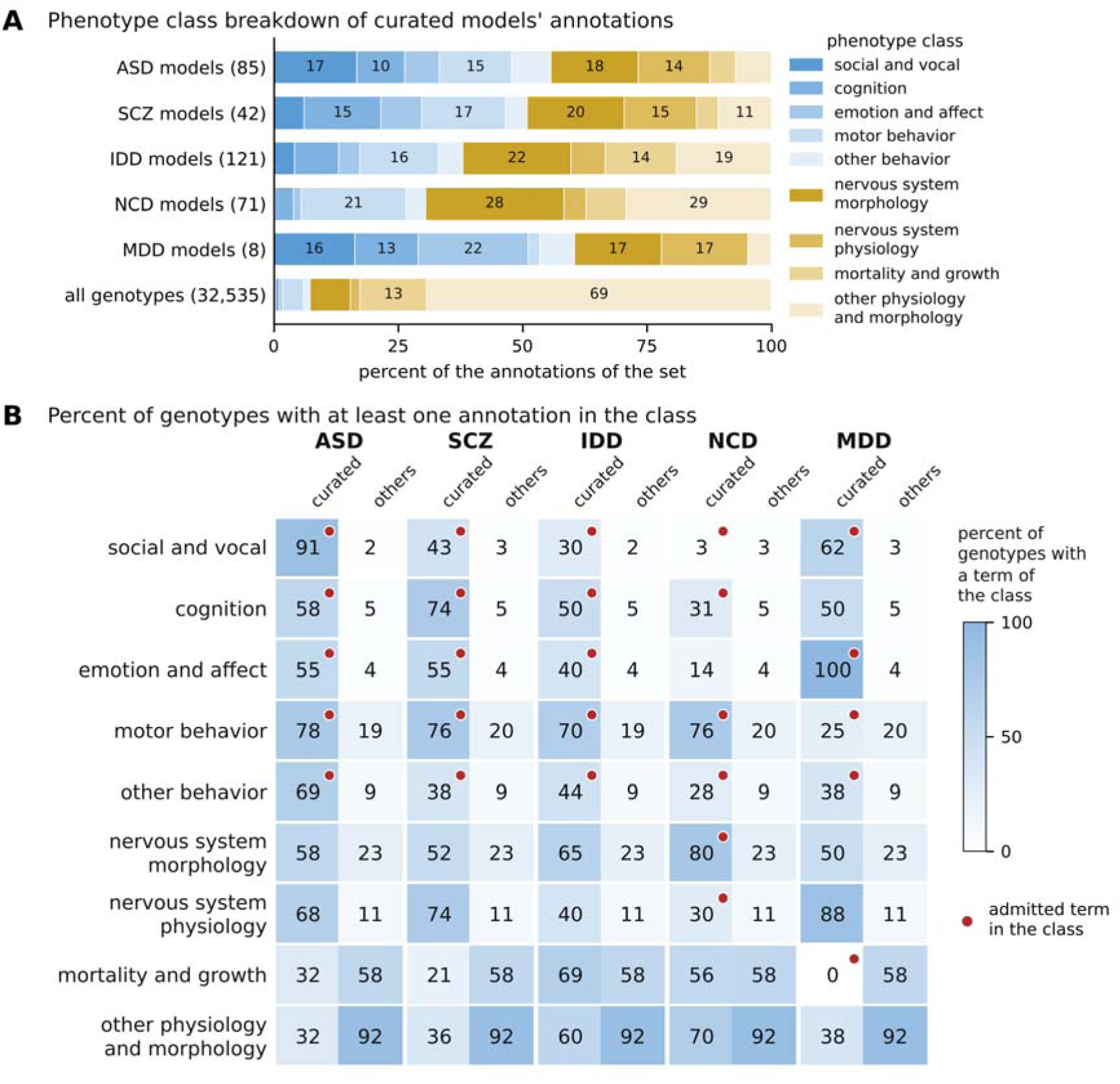
What the annotations of curated models record. (A) Share of each curated set’s annotations by phenotype class, against all 32,535 genotypes. Classes are MP’s own branches (Methods). (B) Percent of curated models and of all other genotypes carrying at least one annotation of the class. A red dot marks a class in which an admitted term of the disorder falls (match export of the primary analysis run). MDD rests on eight models.

#### Most curated models recapitulate part of their definition

Most curated models carry an admitted phenotype for at least one criterion of their disorder, and each credited criterion is far more common among curated models than among background genotypes (Figures 2 and 4, Table 2). The criteria recapitulated differ by disorder. Of the 85 curated ASD models, 81 recapitulate at least one unit and the median is four of seven, led by the social criteria A1 and A3, insistence on sameness and sensory reactivity, while restricted interests reach 2. For SCZ, 36 of 42 models recapitulate a criterion, an upper bound (30 of 42 without the broad emotion term). The two recapitulated are the only two a mouse can reach, disorganized behavior (A4) and negative symptoms (A5). For IDD, 74 of 121 models recapitulate intellectual function or adaptive function, and 26 both. For NCD, 47 of 71 recapitulate cognitive decline (A), and none does for interference with independence (B), which 251 background genotypes do. Of the 47, 18 are credited only through criterion A’s perceptual-motor domain, whose apraxia anchor admits impaired coordination and ataxia, so their cognitive credit rests on motor phenotypes. All eight MDD models recapitulate depressed mood and anhedonia through behavioral despair, anhedonia and decreased coping response, the forced-swim, tail-suspension and sucrose-preference class of readouts whose standing as mood measures is contested (S19 Table). Only Adcy3 adds weight or appetite change and sleep disturbance, reaching four of nine criteria.

**Figure 4.**
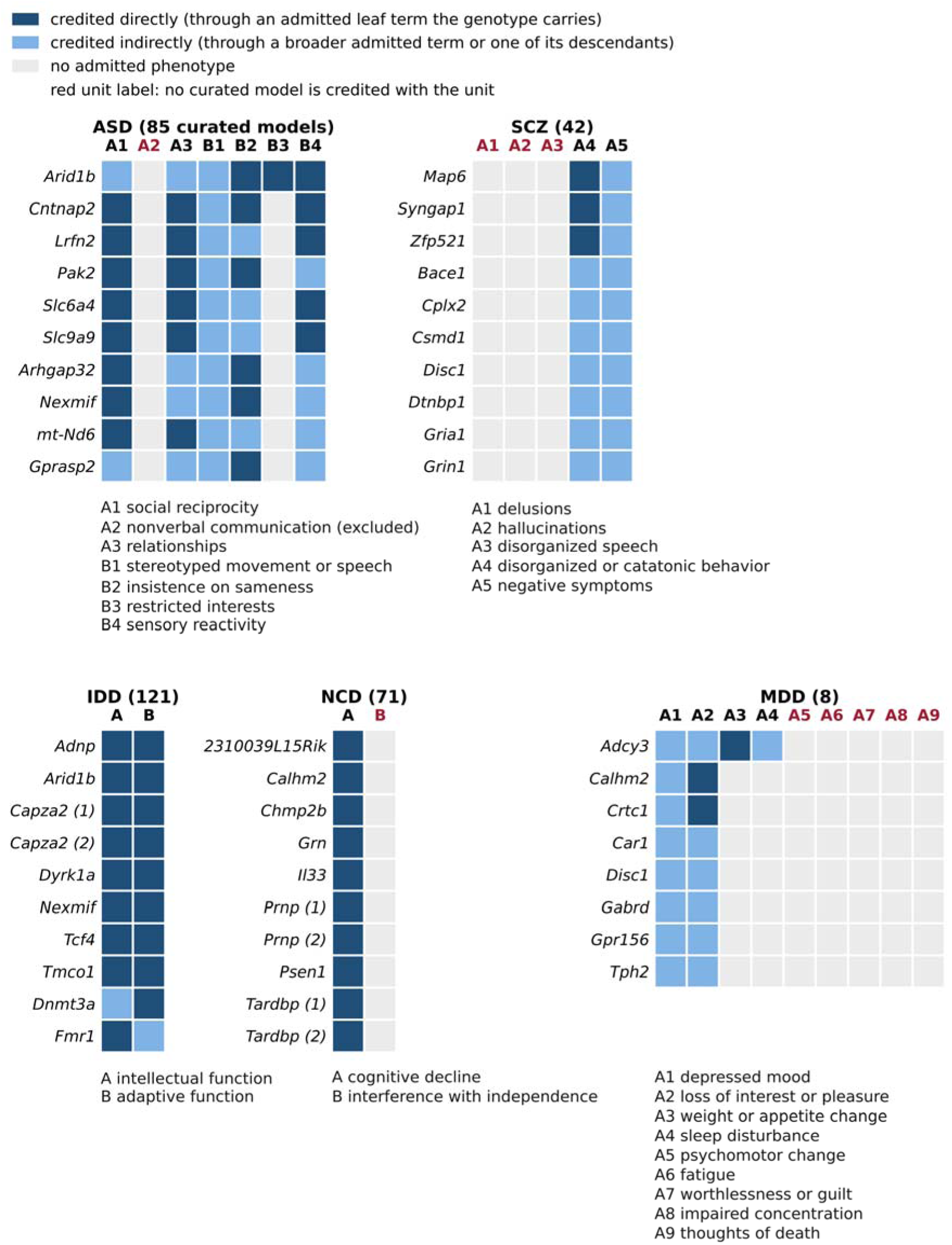
Criterion units credited to the ten highest-coverage curated models per disorder (all eight for MDD), sorted by Diagnostic Coverage, then by direct Diagnostic Coverage and gene symbol (Shank3 and Shank2 rank below the ten ASD models shown). Dark cells are units credited directly (through an admitted leaf term the genotype carries). Light cells are units credited indirectly (through a broader admitted term or one of its descendants), and empty cells carry no admitted phenotype. Red unit labels mark units no curated model is credited with. The profiles of all 327 curated models are released with the catalogue.

**Table 2.** Criterion-level credit per disorder, over the 32,535 genotypes. Highest coverage is the largest number of scored units any genotype is credited with. Genotypes meeting the rule satisfy the symptom-count part of the DSM-5-TR rule (the only part scored here), and the number of curated models among them is in parentheses. Coverage AUROC ranks the genotype set by Diagnostic Coverage and is given with its 95% gene-clustered interval. The restricted column counts only behaviorally annotated genotypes in both classes.

| Disorder | Curated models credited with a unit | Background genotypes credited with a unit | Highest coverage | Genotypes meeting the rule (curated) | Coverage AUROC (interval) | Coverage AUROC, both classes behaviorally annotated |
|---|---|---|---|---|---|---|
| ASD (85 curated) | 81 | 3,319 | 6 of 7 | 0 | 0.964 (0.928 to 0.990) | 0.944 |
| SCZ (42) | 36 | 5,638 | 2 of 5 | 0 | 0.877 (0.807 to 0.933) | 0.732 |
| IDD (121) | 74 | 2,676 | 2 of 2 | 333 (26) | 0.771 (0.729 to 0.836) | 0.735 |
| NCD (71) | 47 | 4,475 | 2 of 2 | 66 (0) | 0.761 (0.690 to 0.824) | 0.604 |
| MDD (8) | 8 | 3,429 | 4 of 9 | 0 | 0.995 (descriptive) | 0.982 |

Ranking the genotype set by Diagnostic Coverage recovers the curated lists well above chance in every disorder, from coverage AUROC 0.964 for ASD to 0.761 for NCD among the four primary disorders (Table 2, S2 Fig.). Intervals come from a bootstrap that resamples genes in place of genotypes; the four primary disorders (those with 30 or more curated models, Methods) carry 42 to 121 curated models, and all four values survive Holm correction (a correction for multiple testing) over the 16 primary claims (S4 Table). The MDD value rests on eight models and is descriptive. With both classes restricted to behaviorally annotated genotypes, the point estimates stay above 0.5. The drop is small for ASD and IDD and about 0.15 for SCZ and NCD, the part of the agreement due only to curated models carrying some behavioral annotation.

#### One profile read against its definition

The homozygous Cntnap2 knockout (Cntnap2 tm1Pele, 18 annotations from one paper) is the one curated ASD model that meets the strict requirement. It is credited with A1 and A3 through social withdrawal and decreased vocalization, with B1 and B2 through increased stereotypic and grooming behavior, and with B4 through a decreased thermal nociceptive threshold. That is five of seven units, and it reaches the support ceiling. It is not credited with A2, which no MP term can express, or with B3 (restricted interests), for which it carries no annotation. Its seizures, motor coordination, and olfaction annotations credit nothing. The inference the results support is therefore a social, repetitive, and sensory phenotype, with no evidence either way on restricted interests, and the sensory credit rests on one nociception term.

In every disorder, some uncurated genotypes are credited with as many criterion units as the best curated models, and for NCD with more (S15 Table). ASD’s highest coverage of six of seven is reached by Arid1b (a curated ASD model) and a CHMP2B transgene at the Atp2b1 locus (a curated frontotemporal dementia model). In all, 44 uncurated genotypes reach five or six, 8 of them deaf or vestibular mutants credited through circling, startle and sensory terms. Four genotypes (Cntnap2, Efna2, Syngap1 and Tcf4) meet the strictest ASD requirement available without A2 (direct credit for A1, A3 and two of B1 to B4). The symptom-count rules of IDD and NCD are met by 333 and 66 genotypes. No genotype can meet the ASD or SCZ rule, because the ASD rule requires A2 and the SCZ rule one of A1 to A3, so Diagnostic Coverage is read within each disorder only.

### Support AUROC: the support score separates curated models from the background, most clearly for ASD

The support score ranks genotypes by the admitted phenotypes behind their credited criteria. It places curated models above the background in all four primary disorders, strongly for ASD (support AUROC 0.956) and moderately for SCZ, IDD and NCD (Table 3, S2 Fig). Every interval lies above chance and all four survive Holm correction (S4 Table). Outside ASD, the separation is mostly an admission rate. Since 84 to 92% of the background carries no admitted term, using a genotype’s ownership of one as a binary indicator already gives 0.896, 0.684, 0.721 and 0.680 for ASD, SCZ, IDD and NCD. Among genotypes that carry one, the score ranks curated models above the background for ASD (0.89) and IDD (0.79) and below it for SCZ (0.44) and NCD (0.40). The comparison with random sets (the permutation null) is kept because it is what makes the ASD and IDD rankings graded, and because it is the only part of the score that reads criterion content rather than admission alone. Average precision (the share of curated models among the genotypes ranked at or above each curated model, averaged) is 25 times prevalence for ASD, 7 times for IDD, and two to three times for SCZ and NCD. The separation does not depend on any one gene or on the minimum of four annotations, and it survives the restriction to behaviorally annotated genotypes (Table 3, S8 Table). Genotypes that no random annotation set matches as well share the maximum score as a tie block (62 curated ASD and 43 IDD models).

**Table 3.** Support AUROC per disorder. Values are support AUROCs with 95% gene-clustered intervals, in the evaluation set (the genotypes carrying a candidate term for the disorder: ASD 11,961, SCZ 12,486, IDD 12,382, NCD 11,603, holding 85, 42, 111 and 67 curated models). The retrieval score is the best embedding and lexical match of each anchor over the genotype’s terms with no LLM verdict (S1 Text). The behavioral count is the number of a genotype’s annotations at or below abnormal behavior (MP:0004924) and carries no criterion content. Depth matching reweights the background to the curated models’ distribution of behavioral annotation depth (0, 1 to 2, 3 to 5, 6 to 10, 11 or more), and the depth-matched count row is a check that the matching worked. PhenoDigm is recomputed over the full phenotype profile, and its difference from FACET carries a two-sided bootstrap p. The retrieval baseline and the other PhenoDigm configurations are in S8 and S10 Tables.

| Comparison | ASD (85 curated) | SCZ (42) | IDD (111) | NCD (67) |
|---|---|---|---|---|
| Support score | 0.956 (0.920 to 0.983) | 0.682 (0.603 to 0.761) | 0.751 (0.705 to 0.820) | 0.668 (0.595 to 0.737) |
| Average precision (prevalence) | 0.175 (0.0071) | 0.0095 (0.0034) | 0.062 (0.0090) | 0.0118 (0.0058) |
| Support score, both classes behaviorally annotated | 0.961 (0.936 to 0.979) | 0.662 (0.584 to 0.739) | 0.761 (0.710 to 0.847) | 0.636 (0.557 to 0.707) |
| Number of behavioral annotations | 0.895 | 0.849 | 0.787 | 0.714 |
| Number of behavioral annotations, depth matched | 0.524 (0.499 to 0.545) | 0.527 (0.493 to 0.565) | 0.511 (0.499 to 0.531) | 0.508 (0.484 to 0.533) |
| Support score, depth matched | 0.895 (0.852 to 0.931) | 0.596 (0.517 to 0.674) | 0.667 (0.618 to 0.736) | 0.582 (0.508 to 0.653) |
| Retrieval score without LLM admission | 0.938 | 0.827 | 0.784 | 0.781 |
| Retrieval score without LLM admission, depth matched | 0.768 (0.718 to 0.819) | 0.540 (0.487 to 0.592) | 0.603 (0.563 to 0.652) | 0.654 (0.611 to 0.707) |
| PhenoDigm recomputed | 0.772 | 0.772 | 0.706 | 0.593 |
| FACET minus PhenoDigm (interval, p) | +0.184 (0.126 to 0.248, p below 0.001) | -0.090 (-0.191 to 0.001, p 0.054) | +0.045 (-0.028 to 0.158, p 0.28) | +0.075 (0.007 to 0.150, p 0.031) |

#### Much of the separation outside ASD reflects how much behavioral phenotyping a genotype received

The number of a genotype’s behavioral annotations is a baseline without criterion content, and it separates curated models from the background above the support score for SCZ, IDD, and NCD (Table 3). The random sets behind the support score match a genotype’s total annotation count but not its behavioral depth, so depth matching reweights the background to the curated models’ distribution of behavioral depth. The count itself then falls to near chance. The support score falls to 0.895 for ASD, 0.596 for SCZ, 0.667 for IDD and 0.582 for NCD: the separation that survives equal phenotyping depth is large for ASD, moderate for IDD and small for SCZ and NCD. A retrieval score without any LLM verdict (the best embedding and lexical match of each anchor over the genotype’s terms) behaves the same way. Before depth matching, it outranks the support score for SCZ, IDD, and NCD. At equal depth, the LLM admission adds separation for ASD and IDD, is within noise for SCZ, and subtracts for NCD (Table 3, S8 Table). The LLM step therefore earns its place by deciding which pairs count, not by raising the ranking. For SCZ, the admitted terms alone rank curated models below equally deep background genotypes (0.397), so the SCZ separation comes from the comparison with random annotation sets of the same size. Building the depth correction into the null instead changes nothing, because whether a genotype has any admitted phenotype explains most of the ranking, and the exact value of the support score adds little (S18 Table). ASD is also the disorder on which the fixed parameters and three of the eight mapping rules were set (Methods). The one separation that survives equal depth is therefore the one that is not out of sample for those settings.

#### FACET exceeds PhenoDigm for ASD and does not differ elsewhere after correction

No automated method evaluates a genotype against individual diagnostic criteria, so the comparison with PhenoDigm (a widely used cross-species phenotype similarity method [21]) concerns the support AUROC only. PhenoDigm is recomputed over the full phenotype profile, because the published table omits term pairs below a cutoff. FACET leads PhenoDigm for ASD by 0.184 and does not differ significantly for IDD (Table 3). It leads for NCD and trails for SCZ before correction, and only the ASD lead survives Holm correction (S4 Table). On FACET’s own admitted pairs, the PhenoDigm scorer comes within 0.062 of FACET in every disorder (S10 Table). PhenoDigm rises with annotation depth, which the four-annotation gate limits (Methods). Referring PhenoDigm to the same nulls as the support score (post hoc, S18 Table) leaves the pre-specified comparison unchanged, except that under the behavior-matched null, FACET leads for ASD, IDD, and NCD and ties SCZ. The paired contrasts detect differences of about 0.09 to 0.14 at 80% power, so smaller differences are not resolved here.

### Validation: human experts, alternate LLMs, and the source of the labels

#### The LLM agrees with an expert as often as two experts agree with each other

The expert study is the decisive test of the pair judgment, so it is reported first. The LLM agrees with a single rater on 77% of ratings (268 ratings on 152 pairs, 95% interval 0.71 to 0.83), and two raters rating the same pair agree 79% of the time (107 pairs, 0.71 to 0.87). On the pairs rated twice or more, the difference is -0.02 (95% interval -0.09 to 0.05). The raters also agree with each other less often on the pairs the LLM admitted (0.70) than on the pairs it refused (0.90). Human experts agree with most of the pipeline’s refusals and with two-thirds of its admissions (Figure 5, S13 Table). Ratings from eight raters cover all 168 pairs with one to three ratings per pair (336 ratings, against the three per pair of the 12-block design). The study is interim: eight of the 12 blocks have been returned, and these results will be updated as the remaining blocks arrive. The raters are seven of the invited experts and the first author, who developed the pipeline and rated one block that no invited rater returned. One of the seven invited raters (ARC) later joined as an author. He rated blind, under the same conditions as every invited rater, before joining. The instrument pairs were drawn by script from the sealed key, so the first author did not choose or know them, and removing those ratings gives confirmation 0.654 and rejection 0.880 with the same verdict (S13 Table). The pre-specified rule sets two levels. The judgment is expert-supported when the confirmation rate of admitted pairs and the rejection rate of refused pairs both reach 0.85 with one-sided lower bounds of 0.70. It is not expert-confirmed when either rate falls below the 0.70 floor and measured otherwise. The rejection rate passes the bar (0.884, lower bound 0.824), although most sampled refusals were clear ones (54 of the 75 rated refused pairs lie in the stratum farthest from admission). The confirmation rate (0.662, lower bound 0.517) falls below the floor, so the pair judgment is not expert-confirmed at this sample size. These rates exclude “cannot tell” (S13 Table). The LLM agrees with the rater majority on 79% of pairs (kappa 0.64). The 0.85 bar exceeds the agreement between raters themselves and was met by the rejection rate alone. Much of the gap lies in how strong an analog must be to count: raters grade the pairs the LLM called direct 0.83 steps lower on the four-step analog scale. Confirmation of admitted pairs is 77% for IDD, 75 for MDD, 67 for ASD, 62 for NCD and 52 for SCZ, and the raters agree with every SCZ refusal (Figure 5B).

**Figure 5.**
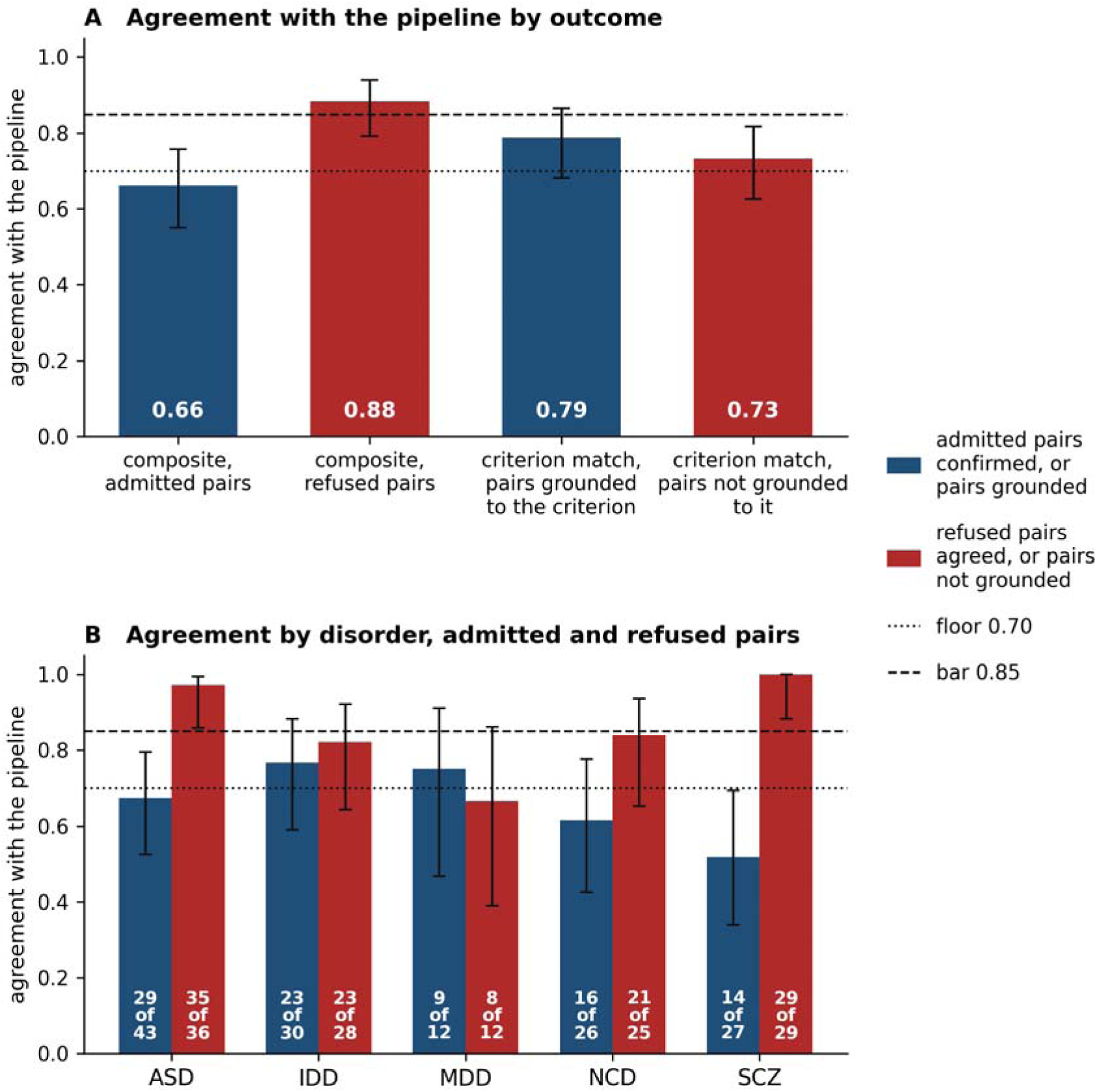
Interim expert ratings from eight raters, seven invited and the first author (77 admitted and 75 refused pairs with a definite rating). (A) Agreement with the pipeline by outcome, on the composite verdict and on the criterion-match question alone. Error bars are Wilson 95% intervals (suited to small samples). The dotted line marks the pre-specified floor of 0.70 and the dashed line the bar of 0.85. On the composite verdict, blue bars are admitted and red bars refused pairs. On the criterion-match question, blue bars are pairs whose mouse term the mapping step assigned to the criterion and red bars those it did not. (B) Agreement by disorder on the composite verdict: confirmation of admitted pairs (blue) and rejection of refused pairs (red), over definite ratings (“cannot tell” excluded). Labels give agreeing ratings of all ratings. The floor and bar apply to the pooled rates and are shown for reference.

#### Alternate LLMs reproduce the ASD result and score SCZ higher

Re-judging every pair with two alternate LLMs (claude-opus-4-8 and gemini-2.5-pro) leaves the ASD support AUROC within the pre-specified margin of 0.05 (0.956 against 0.950 and 0.942) and raises SCZ by 0.182 and 0.159 (0.682 against 0.864 and 0.842, S23 Table). A difference is equivalent when its 90% interval lies inside -0.05 to 0.05 (a conventional margin), not equivalent when it lies wholly outside and inconclusive otherwise. IDD is equivalent under claude-opus-4-8 and inconclusive under gemini-2.5-pro, and NCD is inconclusive under both, which with 67 curated models reflects power. The SCZ result therefore depends on which LLM performs the judgment step. The three LLMs differ mainly in how readily they admit a pair. The alternates keep 87 to 99% of the primary run’s admissions and admit 2.2 to 4.2 times as many, with their extra SCZ admissions falling mainly on A4 and A5 (S1 Text). The primary LLM is the strictest of the three. Against the rater majority on the 140 rated pairs, its admissions have precision 0.66 (the confirmation rate above, measured against the majority) and recall 0.88, against 0.68 and 0.85 for claude-opus-4-8 and 0.68 and 0.84 for gemini-2.5-pro (S13 Table). The instrument was drawn from the primary run’s own verdicts, so recall against the raters is inflated for the primary run and comparable only across runs. Retrieval alone, with no LLM verdict, can admit the same 71 of the 140 rated pairs. By retrieval score it then agrees with the rater majority on 73% of pairs and by embedding similarity alone on 74%, against 79% for the LLM (S29 Table). The LLM’s lead is +0.071 over the combined retrieval score within each disorder (95% interval 0.000 to +0.139) and +0.050 over embedding similarity (-0.02 to +0.12), so the present sample does not resolve either. Retrieval and the LLM form one selection: retrieval ranks candidates by a language-model embedding and word overlap, and the LLM prunes them, admitting 420 of the 14,205 judged pairs. Within the pairs the LLM admitted, the retrieval score barely separates the pairs the experts confirm from those they reject (0.56), so similarity alone cannot make that cut. Because the instrument was stratified on the LLM’s verdicts, the comparison is not neutral between the two rules. Under a disorder-specific variant of its developer prompt, the same LLM admits a similar number of pairs and moves the support AUROC by 0.045 or less, although 19 to 32% of the admitted set turns over (Methods, S1 Text). In the admission control, the PhenoDigm scorer separates curated models from the background better on the admitted pairs than on 100 matched draws of refused pairs. It does so for ASD (p 0.010) and IDD (p 0.020) but not for SCZ (p 0.109) or NCD (p 0.554). The admission control and the expert ratings therefore single out the same two disorders. The upstream LLM steps are more stable. Re-attribution of the anchors by the alternate LLMs agrees with the primary run on all but at most two anchors, except for SCZ, where 10 and 11 of 41 differ (S1 Text, S12 Table).

A majority vote of the three LLMs, examined post hoc as a panel, agrees with the rater majority slightly more often than the primary run (81 against 79%). It confirms 0.710 of its admissions, which clears the 0.70 floor and stays below the 0.85 bar (S1 Text, S13 Table). The primary run is kept as the main result because every validation sample was drawn from it.

#### Label agreement traces to the describing paper, and MGI holds little independent annotation

For most curated models every phenotype annotation comes from the claim paper, the paper MGI cites for the disease claim (ASD 78 of 85, SCZ 34 of 42, IDD 105 of 121, NCD 52 of 71). Removing the claim paper’s annotations, with the same removal applied to comparable background genotypes, leaves 4 of 85 ASD, 6 of 42 SCZ, 8 of 121 IDD and 6 of 71 NCD models with any credited unit. Both AUROC families then sit at 0.49 to 0.54 (S24 Table). The fall measures how little annotation outside the claim paper exists, not whether the score reads it: label recovery is agreement with the paper that made the claim. Genotypes of the same genes recorded by other papers (relatives) score modestly above background within behavioral depth (0.567 to 0.614) and no higher than genotypes of human-genetics genes without a curated model (S14 Table). Uncurated genotypes of human-genetics genes reach the top coverage tier more often than other uncurated genotypes for ASD and IDD, partly through phenotyping attention (S1 Text, S14 Table).

Where phenotyping is uniform and tied to no claim, in the 5,126 genotypes carrying an IMPC allele, neither FACET, PhenoDigm, nor the behavioral annotation count ranks the genotypes of human-genetics genes above the rest (support AUROC 0.48 to 0.52). That test is weak by construction, since the IMPC battery carries almost no social or cognitive assays (S1 Text, S17 and S24 Tables).

## Discussion

### Bounding translational inference through criterion-level auditing

FACET augments mouse model disorder designations with an explicit mapping of which DSM-5-TR diagnostic criteria a genotype’s recorded phenotypes recapitulate. Applied across 32,535 Mouse Genome Informatics (MGI) genotypes and five psychiatric disorders, our framework demonstrates that while most curated mouse lines reproduce subsets of their target condition, none recapitulates a disorder in its entirety. A model organism need not satisfy a complete human clinical diagnosis to yield translational value; rather, its utility depends on bounding experimental claims to the specific symptom domains it empirically reproduces. FACET establishes those inferential boundaries systematically for every catalogued genotype.

FACET’s two measures provide complementary perspectives on face validity. Diagnostic Coverage defines the scope of clinical inference by enumerating both met and unmet criteria, revealing that no annotated mouse profile satisfies the full diagnostic rule for autism spectrum disorder (ASD) or schizophrenia (SCZ). In contrast, intellectual developmental disorder (IDD) and major neurocognitive disorder (NCD) require two-component conjunctions that are structurally less stringent than the polythetic symptom thresholds of ASD, SCZ, and major depressive disorder (MDD). Consequently, diagnostic rule attainment should be interpreted within individual disorders. Whereas Diagnostic Coverage weights all diagnostic units equally (mirroring clinical nosology), the support score penalizes non-specific phenotypes frequently observed across mouse lines (such as sleep disturbances or negative-like symptoms). This weighting explains why curated SCZ models separate from the background more sharply under Diagnostic Coverage (coverage AUROC 0.877) than under the support score. Together, the released catalogue enables investigators to audit the exact criterion-level coverage of mouse genotypes of interest, with retrieval utility calibrated to the average precision benchmarks established in the Results.

### Sources of incomplete diagnostic coverage

The ten criteria credited to no curated model fall into three distinct categories, each requiring a different remedy; distinguishing among them is a central contribution of this study. The fundamental species boundary is the first kind. For MDD A7 and A9, the LLM’s universal refusal of candidate human-mouse phenotype pairs aligns with prior manual evaluations finding no valid rodent assays for guilt or suicidality. Similarly, for SCZ A1 through A3, the pair-judgment module evaluated 1,137 candidates and admitted none, whereas the two alternative LLMs admitted 5 and 41 pairs, respectively. None of those pairs is an analog of a delusion, a hallucination or disorganized speech: they join the three anchors to endophenotype, cognitive and brain-morphology terms (S33 Table). The zero-coverage observed here thus reflects the verdict of the most conservative LLM on clinical features inherently assessed through human speech.

The second is a vocabulary gap that ontology curation could address. ASD A2 cannot be assessed by FACET because no MP term records facial expression, although facial expressions of emotional state have been described in mice [30,31].

The third, an absence of empirical assays, is one researchers can act on. Psychomotor change, fatigue, and impaired concentration are expressible in the mouse vocabulary, credited to hundreds to thousands of background genotypes, yet recorded for no curated MDD model. Figure 3B illustrates this discrepancy directly: only two of the eight curated MDD models carry any motor-related annotation. These gaps motivate a minimum phenotyping standard wherein any model curated for a psychiatric condition is systematically assayed across all criteria expressible in the mouse vocabulary, with normal (negative) results deposited alongside positive findings. NCD B is credited through two terms only, suggesting its absence among curated models reflects a vocabulary bottleneck as much as an assay gap. More broadly, any ontology-based audit is bounded by existing curation conventions. MGI records phenotypes primarily from the initial describing publication, captures almost no negative results, and leaves conditional deletions and obligate intellectual disability syndromes unlabeled. On the other hand, the Human Phenotype Ontology (HPO) lacks structured symptom profiles beneath psychiatric diagnoses, a gap our released anchor sets address across five disorders. Meanwhile, standard ontology similarity metrics are blind to phenotypic directionality, which FACET resolves through an explicit direction check.

### Annotation depth and label circularity

Performance across the four primary disorders is not uniform, and our claims should be interpreted in descending order of evidential strength: ASD, then IDD, then SCZ and NCD. For ASD, the support score separates curated models from the background at 0.90 after depth matching, so the separation rests on criterion content and not on how much phenotyping the models received. For IDD, a moderate separation remains (0.67). For SCZ and NCD, however, total behavioral annotation count separates curated models from background genotypes as effectively as the support score, leaving minimal signal once behavioral annotation depth is equalized (support AUROCs of 0.60 and 0.58, respectively). This equalization should be read with care. It matches genotypes on their number of behavioral annotations, and MGI records abnormal results only, so a genotype’s behavioral depth reflects the abnormalities it showed as well as how much it was tested. Matching on depth therefore removes some true signal together with the phenotyping bias, and the depth-matched values are best read as a lower bound on the separation that criterion content carries. Outside ASD, the support AUROC functions largely as a binary admission indicator: carrying any admitted term separates the classes as well as the score does, and among admitted genotypes, curated SCZ and NCD models rank below the background. For SCZ and NCD, therefore, FACET’s primary utility lies in cataloging precisely which genotypes carry an admitted phenotype for a given criterion, rather than establishing graded whole-syndrome resemblance. Notably, the SCZ benchmark is also the most sensitive to LLM choice, indicating that its lower bound is partly governed by the strictness of our primary LLM. NCD serves as an instructive boundary case as its two criteria are satisfied primarily through common cognitive and grooming terms shared widely across background genotypes.

Recovery of the curated labels is the expected result for a valid method, and the source of the annotation sets its scope. Nearly every curated model’s phenotype profile derives from the same publication that originally claimed the disease association. So, label recovery is in part agreement with that paper, and removing it leaves too little annotation to rank. Independent phenotypic replication of curated lines remains scarce, and the primary uniform phenotyping effort, the International Mouse Phenotyping Consortium (IMPC), does not assay the behavioral domains specified by psychiatric diagnostic criteria. This explains why neither FACET nor PhenoDigm detects a signal in IMPC data. This circularity constrains all methods benchmarked against MGI disease assertions, including PhenoDigm and downstream exome-prioritization tools. Overcoming this limit will require independent phenotypic replication and testing of genotypes assayed after disease candidacy is proposed.

### Field-specific phenotyping conventions across disorders

Figure 3 locates the cause of the order of the four disorders in the phenotyping conventions of each field. The autism field phenotyped its candidate genes (catalogued in resources such as SFARI Gene) [29] to a shared standard of social, communication and repetitive-behavior batteries designed around the diagnostic criteria [16,17]. Consequently, uncurated genotypes achieve high ASD coverage alongside curated lines. A high ASD score therefore reflects alignment between experimental assays and clinical nosology, not proof that ASD mouse models recapitulate human pathology more faithfully than other disorders’ models. The schizophrenia field predominantly favored endophenotypes over full-syndrome modeling [9,18], establishing models via neurophysiological or sensorimotor assays outside DSM-5-TR symptom criteria. Our mapping rules flag these endophenotype terms, and the pair-judgment module independently rejected every such pairing against criterion anchors (S1 Text, S20 Table). The NCD set is dominated by Parkinson and Huntington models recorded through motor assays. Here, 18 of the 47 credited NCD models qualify solely via criterion A’s perceptual-motor pathway, likely inflating apparent cognitive coverage. An alternative interpretation is considered: FACET’s discriminative power declines where clinical criteria lack direct rodent analogs, given that three SCZ criteria rely on human speech and expert confirmation rates are lowest for SCZ and NCD admissions. Evaluating SCZ models against dedicated endophenotype anchors would distinguish between these two explanations and remains an important next step.

### LLM judgment matches expert consistency, with the rejection rate stronger than the confirmation rate

The LLM admits some pairs the experts reject and refuses some they would admit, with objected admissions proving more frequent than objected refusals. Substituting alternative LLMs keeps the ASD support AUROC within our pre-specified equivalence margin while raising the SCZ score, confirming that SCZ performance depends on the LLM’s stringency. The higher value under the more permissive LLMs does not establish that the SCZ result is understated, because the experts confirmed only half of the SCZ admissions. The experts agree with nine in ten refusals and confirm two in three admissions, and they agree with the LLM about as often as with each other. A majority vote of the three LLMs raises confirmation to 0.71, above the 0.70 floor, but not near 0.85. The human-LLM gap lies in how closely a rodent phenotype must approximate a human symptom to qualify as a valid analog. Auditing admitted human-mouse phenotype pairs directly against primary behavioral data remains an open task. Our recruited expert raters would admit about one in three refused pairs stopped by the strict requirement, so strict eligibility costs recall (S1 Text). Furthermore, prompt variations alter one-fifth to one-third of the admitted pair set while shifting the support AUROC by at most 0.045. The support AUROC therefore cannot certify the admitted set, and the expert study is the decisive benchmark. Against the experts, the LLM’s verdicts add 5 to 7 points of agreement over text-similarity metrics alone, a modest margin that the present sample does not resolve. Crucially, FACET’s main technical advance is criterion-anchored pair admission rather than its scoring formula. Given identical admitted pairs, PhenoDigm and FACET scores differ by at most 0.062, whereas no whole-profile method identifies criterion-specific matches. Because experts confirm two-thirds of admitted pairs, the released catalogue should be treated as a first-pass resource that further expert curation can progressively refine.

### Limitations

FACET evaluates semantic alignment between recorded mouse phenotype terms and human diagnostic criteria, and nothing beyond this scope. It makes no mechanistic, etiological or predictive claim, and a low score does not refute a model. The DSM-5-TR criteria are also the only standard applied. A genotype may model features of a disorder that the criteria do not specify, such as endophenotypes or associated features (cognition in SCZ), thus FACET assigns such phenotypes no credit. FACET also counts the criteria a genotype’s phenotypes model and does not account for recorded phenotypes that run against the clinical presentation. So, a model credited with several criteria may still diverge from the disorder in other respects. Diagnostic Coverage and the support score therefore quantify face validity against formal diagnostic criteria alone, representing only one dimension of model validity. Furthermore, because FACET parses ontology annotations rather than underlying experimental conditions, secondary behavioral artifacts can inflate coverage. For example, reduced social investigation caused by a primary sensory or motor deficit is credited toward ASD criterion A1 exactly as annotated in the source publication. Among the 44 uncurated genotypes meeting five or six ASD criteria, 8 carry deafness or vestibular mutations. Quantifying such experimental confounds across all disorders would require pair-level evaluation of assay dependence beyond the scope of our current outputs.

Accordingly, the ASD count serves as a conservative lower bound, as sensory and motor confounds are not restricted to auditory or vestibular mutants.

Several design choices could have influenced these results. The fixed parameters and three of the eight mapping rules were first set on ASD, so the ASD result is not out of sample for them. SCZ, IDD and NCD received them unchanged and no setting was re-tuned per disorder. A higher judgment floor and a change of channel weights (Methods) both leave ASD and IDD unchanged and lower the SCZ or NCD support AUROC, the two disorders whose admissions the experts confirm least. The weights matter where the result already depends on which pairs the LLM sees. A rule adopted during development holds that a motor term cannot model NCD criterion B (interference with independence). It discards 15 admitted pairs, including terms that curated NCD models carry, so the NCD result is reported with and without the rule, and the NCD coverage and support AUROCs move by 0.014 or less. Similarly, while SCZ mapping rules flag endophenotype terms such as prepulse inhibition, they did not prevent curated SCZ annotations from reaching the judgment stage. All 234 behavioral annotations across the 42 curated SCZ models entered the pair-judgment stage (relevance tiers 1 to 3; Methods). The endophenotype pairs among them were independently refused against criterion anchors (S1 Text, S20 Table), obviating the need to rerun SCZ without those rules. Finally, LLM judgment is bounded by upstream candidate retrieval, which passes at most 20 terms by embedding similarity and 20 by lexical overlap per anchor. Since false negatives at the retrieval stage were not quantified, the admitted set may omit valid matches that experts would otherwise endorse.

The interim expert study represents eight of the 12 invited raters, at the design’s minimum. One of the eight is the first author, whose block is reported in a sensitivity analysis that excludes it. A second of the eight later became an author, having rated blind as an invited rater. The first author wrote the rating instrument (its two questions, answer scales, and instructions) without prior pilot review by external domain experts. Thus, inter-rater agreement reflects consistency under that specific operational framing. Finally, all measurements are sourced to a single database (MGI), fixed releases of the MP and HPO ontologies, and our project-specific paraphrasing of DSM-5-TR criteria.

## Conclusions and future work

FACET turns face validity, a quality long debated on a model-by-model basis, into criterion-level profiles for 32,535 MGI genotypes across five DSM-5-TR disorders. Our findings support quantitative model ranking where phenotypic evidence separates curated mouse lines from background genotypes at equal annotation depth (ASD and IDD). Elsewhere (SCZ and NCD), the outputs serve strictly as a structured catalogue of criterion-specific phenotype matches. Across all four primary contrasts, FACET identifies admitted phenotypes for most curated models and recovers curated lines above chance under both metrics. It also partitions unmodeled diagnostic criteria into those for which no rodent analog was admitted, one lacking an MP vocabulary term, and four awaiting empirical behavioral assay. We release three open resources with this paper: the per-criterion catalogue, the criterion-level HPO anchor sets, and the criterion-mapped taxonomy of the MP vocabulary.

We record every admitted pair against one criterion unit, so results can be regrouped under the Research Domain Criteria (RDoC) or a revised manual [32,33] without rerunning any judgment. Only new criteria require new anchors. Extending the pipeline with a dedicated endophenotype layer represents the immediate next step, as SCZ and NCD annotations largely reflect intentional neurophysiological and lesion-confirming assays rather than clinical symptom modeling. Evaluating models against endophenotype anchors will directly test whether lower DSM coverage stems from experimental design choices or cross-species boundaries.

Looking ahead, a prospective validation of label recovery, which scores genotypes whose phenotyping followed their proposal as disease models and was therefore not produced by the paper that made the claim, will depend on such genotypes being phenotyped and deposited. Future iterations of this method should integrate domain experts from the outset to co-design seed anchors, analog grading scales, and admission thresholds, alongside multi-provider LLM panels [34] that outperformed single-model judgment here. In the immediate term, the released catalogue enables investigators to bound translational claims explicitly by the criteria each mouse model recapitulates, and to target the actionable criteria where experimental evidence is still missing.

## Methods

### Data sources and the genotype set

The primary analysis run reads fixed snapshots of every source (S25 Table). File names, download dates, and checksums are listed in S2 Table, and later releases were used only to measure database drift. Every diagnostic criterion is given in this paper, the catalogue, the expert rating instrument and the code release as a paraphrased item written for the project (those of this paper are in S1 Table), and none of them reproduces DSM-5-TR text. The LLM steps of the primary analysis run read a paraphrase of each disorder’s criteria and supporting text, written for the project, and no DSM-5-TR text enters any prompt. The paraphrased criteria are deposited with the code release. The paraphrase of the supporting text is not redistributed, and the deposited anchor sets are the starting point from which the later steps can be repeated. The mouse vocabulary is the 14,750 non-obsolete MP terms reachable from the ontology root through is_a relations. Of these, the 11,485 annotated to at least one MGI genotype are candidates for judgment. The scored genotype set is the 32,535 of 76,060 MGI genotypes carrying four or more MP annotations. The threshold excludes genotypes whose score would report annotation sparsity in place of phenotype, and its effect is measured in Results by lowering it to one annotation. A genotype with no matching annotation is retained at a score of zero.

### Disease labels and primary disorders

Disease labels come from MGI’s curation of genotypes as disease models [20] through the Disease Ontology [39]. A genotype is a curated model (a positive) for a disorder when an MGI disease annotation names the disorder under the label rules of S3 Table. The positives in the set number 85 for ASD, 42 for SCZ, 121 for IDD, 71 for NCD, and 8 for MDD. Two genotypes that the disease-name rule matched on “Parkinson” are models of Wolff-Parkinson-White syndrome (a cardiac conduction disorder) and were removed from the NCD positives. They remain in the genotype set as unlabeled background. MGI curation records almost no explicit negatives, so uncurated genotypes are treated as unlabeled background in the positive-unlabeled sense [40], and the few explicit negative records are analyzed separately. A disorder has adequate sample size to separate curated models from the background with 30 or more positives, where the 95% AUROC interval has a half-width near 0.10 [41] (S32 Table). ASD, SCZ, IDD and NCD meet this and are the four primary disorders, and MDD is reported descriptively. Genes implicated by human genetic studies that used no mouse data (H genes, S5 Table) supply the attention control, which is the set of genotypes of H genes that have no curated model. NCD has no such list and uses its MGI disease-gene associations in their place.

### Language models and reproducibility

Each LLM step runs on a fixed LLM version with fixed settings (Table 4). The instructions and the material to be read are passed in separate roles. The answer is returned as a form whose fields and permitted values are fixed in advance, so the LLM cannot answer outside the enumerated options. Every output is cached under a hash (a fingerprint) of its full prompt, and the served LLM version is recorded where the interface returns it. A provider can serve an updated LLM under the same name, and seeds are honored only on a best-effort basis, so a fresh LLM run cannot be guaranteed to repeat the primary analysis run. The primary verdicts are therefore stored unchanged and released with the code, and every reported number is recomputed from them by deterministic steps. The prompts that ran are reproduced in the supplement. They carry the paraphrased criteria and no DSM-5-TR text. Deterministic steps run in Python 3 and use rapidfuzz for lexical matching [42] and NumPy for permutation draws [43]. The code release pins package versions and contains the configuration files, the frozen verdicts, and the scripts for every table and figure.

**Table 4.** The LLM steps of the pipeline and their validation. Every call uses temperature 0, seed 1 and a JSON schema (a machine-readable answer form) that enumerates its answers, except where stated. The last column gives each validation test and the date it was specified. The deterministic steps are covered by a reproduction gate (2026-10-19) under which the released code reproduces every score from the frozen verdicts.

| Step | Purpose | Input visible to the step | LLM | Output | Validation (date specified) |
|---|---|---|---|---|---|
| Criterion decomposition | Break each criterion into its observable features | The disorder’s DSM-5-TR chapter | gpt-5.4 (served gpt-5.4-2026-03-05) | Phenotype phrases with polarity and centrality | Not validated separately |
| Anchor resolution | Attach one HPO term to each phrase | The phrase and a 12-term HPO shortlist | gpt-5.4, answers restricted to the shortlist or NONE | HPO anchors | Not validated separately |
| Anchor attribution | Assign each anchor to the criterion unit it serves | The anchor and the criterion units | gpt-5.4 (served gpt-5.4-2026-03-05), provider default sampling, out-of-vocabulary answers abort the run | Anchor to unit map | Re-attribution by claude-opus-4-8 and gemini-2.5-pro (2026-10-19) |
| Mouse domain labeling | Place each MP term in a symptom domain | Five MP fields per term, 15 terms per call | gpt-5.4-mini | Domain and eligibility flags | Hand-curated anchor sets of 13 to 17 terms per disorder, run before release (2026-06 to 2026-07) and rerun 2026-11-08 (S21 and S22 Tables) |
| Mouse criterion mapping | Assign each MP term to the criteria it could serve | MP identifier, label and definition (parent and ancestor fields empty) and the criteria text | gpt-5.4, prompt v4_2026-07-27 | Criterion assignments and relevance tiers | A 62-term pre-declared pilot for ASD (2026-07-27), reclassification of a stratified sample by both alternate LLMs (2026-10-20) and the 15-term rerun of 2026-11-08 (S22 Table) |
| Pair judgment | Decide whether a mouse term models a feature of the anchor’s criterion | Anchor HPO label with its criterion phrase, domain label and up to five candidates with their scores | gpt-5.4 (alias, served snapshot not logged by the step, inferred gpt-5.4-2026-03-05) | Enumerated verdict fields | Admission negative control (2026-09), LLM swap (2026-10-19, amended 2026-10-27) and expert rating study (2026-09-13) |
| Text embeddings | Semantic retrieval of HPO and MP terms | Term text | text-embedding-3-large | Vectors | Not validated separately |

Analyses described as pre-specified were specified in dated internal documents before their outputs existed. The documents are deposited with the code release, and every deviation is reported with its result. The design fixes every scoring parameter and mapping rule on ASD and then applies them unchanged to SCZ, IDD, and NCD. No parameter was compared with another disorder’s labels before it was fixed, and none was re-tuned afterward, so the three disorders are out of sample for the parameters. MDD’s mapping ran on an earlier prompt and carries none of the rules (S26 Table). The judgment floor, the channel weights, the aggregation rule, and the polarity regime were swept on the primary analysis run (S30 and S31 Tables, S1 Text). The bundle rule and the clustering threshold were not swept. SCZ is the cleanest held-out test. IDD and NCD carry three changes made after their labels were compared, each reported with its unchanged arm: the IDD label rule, the removal of the two NCD labels, and the rebuild of the NCD criterion B anchors. The dated record of what was fixed when is given in S1 Text. Call counts and the usage of the validation runs are given in S1 Text.

### Anchoring

Anchoring resolves each DSM-5-TR criterion into a set of HPO terms (its anchors, each representing one observable feature of the criterion), and these sets are the criterion-level substrate against which mouse terms are assessed. Three LLM calls build the anchor sets (Table 4). The first reads the disorder’s criteria and supporting text in paraphrase and returns the observable content of each criterion as short phenotype phrases with polarity and centrality labels. The second resolves each phrase to at most one HPO term from a shortlist of 12. The shortlist is built by a lexical retriever over the label and synonyms and a semantic retriever over the label, synonyms and definition of every HPO term (S1 Text). A resolution is kept when its polarity is present or either, its centrality is core or supporting, and its relation to the phrase is exact, broader, or narrower. The third call attributes each anchor to the criterion unit it serves or marks it unattributable. No confidence floor is applied at any LLM step. A criterion unit is the scoreable content of one criterion. It is each numbered item where the manual counts items (ASD 7 units, SCZ 5, MDD 9) and one unit per criterion where the manual lists examples (IDD 2, NCD 2). Criteria that state onset, duration, exclusions, or functional consequence without phenotypic content (ASD C to E, SCZ B to F, IDD C and NCD C and D) are not scored. Anchors from supporting text or attributed to such criteria are judged but never credited. Each unit of the four primary disorders declares in the configuration the direction of change it requires, applied in code and shown to no LLM. For SCZ A1 to A3, the mouse criterion mapping prompt carried a note that no organism-level analog exists, while the pair judgment received no such note. ASD A2 was measured with no note at any step. No hand-curated gate was applied to this step.

### Mouse criterion mapping

Mouse criterion mapping assigns MP terms to the diagnostic criteria they could represent. Two LLM calls assign MP terms to the criteria they could serve (Table 4). The first places each of the 11,485 annotated MP terms in one symptom domain from a fixed per-disorder list (S1 Table) with eligibility flags. The second assigns each term in the disorder’s mapping scope (1,655 to 2,393 terms per disorder, one call per term) to the criterion units it could serve, from the MP identifier, label and definition, and the criterion text. The parent and ancestor fields of that prompt were empty in every primary analysis run. No HPO term, gene or genotype is in the prompt. The mapping prompt carries named per-disorder rules declared in the configuration (S26 Table). Eight rules exist, and each disorder opts into one to three of them, which the archived prompt of each run records. ASD carries grooming routing (patterned motor sequences to B1, not B2), involuntary motor signs (tremor, ataxia, seizures, and clasping refused for every criterion), and construct refusals (memory assays and sensory-capacity losses refused for the social and sensory criteria). IDD carries human schooling constructs (reading, arithmetic and IQ marked unassessable). NCD carries decline versus deficit (a deficit admitted with the missing longitudinal qualifier disclosed) and the language floor (vocalization terms flagged for review). SCZ carries the motor boundary from DSM (A4 admits posturing, negativism and agitation and refuses seizures, startle and bare activity) and mechanism is not face validity (prepulse inhibition, gating, latent inhibition and drug responses flagged as endophenotypes). MDD’s mapping predates the rules and carries none. Four rules flag terms for review without deciding them, so that a project decision stays testable against real candidates. Before each disorder’s mapping was released, a hand-curated anchor set with expected labels was run in a pilot mode that printed pass or fail per term. SCZ, IDD, NCD and MDD carried 13 to 17 MP terms with expected domains, and ASD a 62-term pre-declared set with expected criterion units, which passed its acceptance rule in full. Read against the primary taxonomies, the domain sets recover the expected domain for 61 of 62 terms and refuse all 22 negative controls. They admit 32 of the 40 terms expected admissible, and the eight misses are design decisions taken after the sets were written (S1 Text, S21 Table). A rerun on 2026-11-08 with 15 terms per disorder reproduced the primary verdict on 74 of 75 terms (S22 Table). These gates check whether mapping admits a hand-curated list and reproduces its verdicts, not retrieval recall, which was not measured. A relevance tier is then set in code. Tier 1 holds terms admitted to a unit with the required direction. Tiers 2 to 4 hold unadmitted terms that are behavioral (tier 2), anatomical or physiological (tier 3) or developmental and other (tier 4), and tier 5 holds terms outside the disorder’s mapping scope, which are never judged. Tiers 1 to 4 enter retrieval as candidates. The direction check applies after the LLM answers: an assignment whose mouse term states the direction opposite to the one the unit requires is vetoed, and a term with no stated direction passes. Of the 378 admitted pairs on scored criteria in the four primary disorders, 185 carry a mouse term with no stated direction and 45 a term that is not directional (an abnormality with no pole). The veto could therefore act on the remaining 148. The resulting assignment of every mouse term to domains, tiers and units is the mouse taxonomy.

### Retrieval and the combined retrieval score

For each anchor, retrieval keeps the 20 mouse terms with the highest cosine similarity of text-embedding-3-large vectors and the 20 with the highest rapidfuzz lexical score. The human side of each comparison is the anchor’s HPO label joined to its criterion phrase, and the mouse side is the MP label joined to its definition. Any annotated descendant of a retrieved term in tiers 1 to 4 is added as an injected candidate. Each candidate pair receives a combined retrieval score between 0 and 1 from four channels: the embedding cosine, the lexical score and the Resnik [44] and Jaccard [45] ontology similarities read from the Monarch table. Each channel is min-max normalized over the candidate table of the run, and the channels are combined with weights 0.45, 0.25, 0.20 and 0.10, renormalized over the channels present (S27 Table). When both ontology channels are absent, the lexical weight is halved before renormalization. The two ontology channels are absent for 97 to 99% of candidate pairs because the Monarch table covers few behavioral term pairs, so on nearly every pair the score is 78% embedding similarity and 22% lexical score. Its channels combine by a fixed rule, while its embedding channel is the output of a learned language model. The weights, the lexical factor, the bundle rule and the clustering threshold were set by hand on the ASD development run and were not tuned afterwards. A channel that cannot be computed is treated as missing.

### Pair judgment

The LLM is asked how well each candidate mouse term, as defined in MP, serves as an organism-level analog of the human target. It sees the anchor’s HPO label joined to the criterion phrase that decomposition returned and the anchor’s domain label. It also sees up to five candidates in descending score order (each with its MP label, definition, domain, score and channel values), assembled from fixed format strings. It sees no criterion text, gene, genotype, MGI disease label, curated list, or PhenoDigm score. The developer prompt is shared by every disorder and names the MDD symptom domains (S1 Text), while the user prompt carries the disorder’s own anchors and domain labels. The LLM returns only enumerated fields. They are a reranking action (promote, keep, demote, abstain), whether the mouse term is an organism-level observable, the scope of the observation, an analog grade (direct analog, close support, weak support, mechanistic only, mismatch), and strict and broad eligibility. A free-text reason is logged and read by no downstream step. A pair is admitted when it is strictly eligible, graded direct or close, and is not vetoed by the direction check. The final pair score is the combined retrieval score plus the adjustment of S27 Table, clipped to 0 to 1. Three routes carry a pair to the LLM. The five highest candidates of an anchor are judged together when the top two scores lie within 0.08 or the top score lies between 0.35 and 0.70. Every other pair scoring at least 0.57 and every injected descendant is then judged in groups of up to five per anchor with the same prompt and schema. The floor of 0.57 maximized balanced accuracy (0.736) against the LLM’s own broad eligibility verdicts on 180 ASD pairs and is applied to every disorder without per-disorder tuning. Raising the floor removes only pairs judged by the floor route, since bundle pairs and injected descendants are judged whatever their score. On the primary analysis run, a floor of 0.60 removes 6 to 14 admitted pairs per disorder and leaves the support AUROC equivalent to the primary run in every disorder. A floor of 0.65 removes 17 to 26 and leaves ASD and IDD equivalent, while SCZ falls by 0.078 and NCD by 0.045 (S30 Table). Lowering the floor needs new judgments and was not measured. The channel weights were swept the same way, and because they decide which pairs reach the LLM, the pairs a weighting newly routes were judged for the two weightings that moved a support AUROC. Every weighting leaves ASD and IDD equivalent to the primary run. Removing the lexical channel lowers SCZ by 0.059, and equal weights lower NCD by 0.136, because the broad cognitive and motor terms that credit most NCD models no longer reach the LLM (S31 Table). The lexical factor alone changes no support AUROC. Strict eligibility marks a high-quality organism-level analog suitable for strict support, and broad eligibility marks weaker but meaningful support. Only strict eligibility admits a pair. A disorder-specific variant of the developer prompt was run with the same LLM as a pre-specified sensitivity test, and its verdicts are released alongside the primary run. It moves the support AUROC by 0.045 or less in every disorder (inconclusive against the 0.05 margin for ASD, SCZ and IDD, equivalent for NCD) while replacing 19 to 32% of the admitted pairs (S1 Text). The shared prompt is kept as the primary run because every validation sample (the expert pairs, the alternate-LLM re-judgments and the admission control) was drawn from it, and the catalogue and the frozen verdicts are those of this run. For NCD, a construct rule adopted during development (that a motor term cannot model criterion B) sets the 15 admitted pairs it reaches (11 motor and reflex terms) to not eligible. The filtered configuration is the primary analysis run and the unfiltered one is reported in Results as a sensitivity test. Prompt construction follows the provider’s published guidance [46,47].

### Aggregation and the two measures

MP is a hierarchy in which the descendants of a term are its more specific forms and its ancestors are broader classes. A unit is credited to the matched term and to every descendant of it, never to an ancestor. A genotype is credited with a unit when its own annotations contain a credited term. An injected match is kept only when an admitted retrieved match under the same anchor is its ancestor, and the broadest behavioral term (abnormal behavior, MP:0004924) is excluded from crediting. The criterion profile of a genotype is the set of units credited to it, with the admitted pairs behind each. Diagnostic Coverage is the number of scored units in the profile, where units are weighted equally and a single admitted annotation suffices. Direct Diagnostic Coverage counts only units credited through an admitted leaf term (one with no descendants) that the genotype carries, and indirect credit is reported separately. The diagnostic rule of each disorder (S1 Table) is evaluated on the profile after scoring as a binary indicator and enters no score. For the support score, an anchor is assessable for a genotype when at least one of the genotype’s terms was judged against it. Each admitted match enters with its final pair score (the combined retrieval score after the LLM’s adjustment, Pair judgment). Within an anchor, the admitted matches are clustered (same term, shared ancestor, or token Jaccard of term text at or above 0.65); the match with the highest final pair score in each cluster is kept, and the kept scores combine by noisy-OR (one minus the product of one minus each score). Anchor values average without weights to the domain and domain means to the genotype, and an assessable anchor without an admitted phenotype contributes zero. The score is referred to a permutation null that holds the total annotation count fixed. For each count k of distinct MP terms carried by a genotype (capped at 200), 10,000 sets of k terms are drawn uniformly without replacement from the annotated MP terms. Each set is scored identically (NumPy generator, seed 1). The p value is one plus the number of null scores at or above the observed, divided by 10,001 [48]. The null draws terms uniformly and does not preserve the co-occurrence of real annotations, so a p value is a size-matched rarity index used for ranking and no per-genotype p is read as a test. A null that draws behavioral and other terms separately in the genotype’s own proportion is reported as a sensitivity analysis (S1 Text), a frequency-weighted null was not run, and all inference rests on the bootstrap intervals of the AUROCs. The ranking statistic is the negative base-10 logarithm of p capped at 4.0. Genotypes tied at the cap were re-tested with 1,000,000 permutations under a pre-specified rule (S1 Text). Because the re-test leaves most of each capped block tied, the block is reported as a tie. The catalogue orders it by a standardized support score (z against the permutation null) as a labeled secondary order.

### Evaluation framework

Every analysis evaluates whether a measure ranks the curated models of a disorder above the other genotypes and the substrate each ranking draws from. All analyses are deterministic given the frozen verdicts and are listed in the order of Results in S6 Table. We report two AUROC families, named by the measure that ranks them: the support AUROC ranks genotypes by the support score, and the coverage AUROC by Diagnostic Coverage.

The support AUROC is evaluated among genotypes carrying at least one candidate term for the disorder and repeated on the full background with the rest set to zero, and coverage analyses use the full genotype set. Clustering for every interval is by the mutated gene read from the allele symbols (S1 Text for transgene rulings), so that alleles of one gene are not counted as independent. A genotype is behaviorally annotated when it carries at least one annotation at or below MP:0004924. The phenotype classes of Figure 3 are MP branches, each term taking the first class in the order of S16 Table whose root is among its ancestors. The four behavioral classes kept separate (motor, cognition, emotion and affect, and social joined with vocalization) are the four largest children of abnormal behavior among curated models’ annotations (35, 19, 17 and 12%). Other behavior pools the remaining 15 children; two classes are the nervous-system branches, one joins mortality and growth, and the last pools every other branch. Behavioral depth is the number of behavioral annotations binned as 0, 1 to 2, 3 to 5, 6 to 10 and 11 or more. The 16 primary claims listed in S4 Table are corrected using the Holm method as a safeguard since the family was fixed after the results. The family holds the behavioral PhenoDigm configuration, specified before the recomputed configuration was adopted as primary, while the NCD lead over PhenoDigm recomputed and the H-gene odds ratios lie outside it. Two baselines are scored on the evaluation set. The behavioral count is the number of a genotype’s annotations at or below MP:0004924. The retrieval score without LLM admission takes for each scored anchor the highest combined retrieval score among the genotype’s own terms that retrieval listed for it, combines anchors within a unit by noisy-OR and averages the units. A second version of it is restricted to the admitted pairs. Depth matching weights each background genotype so that the background matches the curated models’ distribution over the five behavioral-depth bins, and the weighted support AUROC is bootstrapped by gene cluster with the weights recomputed in each replicate. S1 Text defines the composition of the annotation record, the lower bound on genotypes assayed with normal results, and the HPO gap. It also reports the supplementary sensitivity analyses: annotation detail, normal-result counting, explicit negatives, allele design (S9 Table), cross-disorder overlap (S1 Fig), and the mirrored permutation null.

### Comparison with PhenoDigm

PhenoDigm scores a genotype against a disease by the semantic similarity of their whole phenotype profiles, read from the positions of terms in the aligned human and mouse ontologies [21]. Its values are distributed by the Monarch Initiative as precomputed similarities between HPO and MP term pairs [37]. PhenoDigm scores here are the symmetric best-match average of Smedley et al. [21] between a genotype’s MP terms and the disorder’s resolved anchors. The anchors are those held in the comparator score files (46, 41, 25, and 28 for ASD, SCZ, IDD, and NCD). They are computed on the Monarch per-pair score (S1 Text). The anchor set FACET resolves serves as the human profile, because PhenoDigm has no criterion-relevant term set and HPO attaches none to these diagnoses. FACET’s score is referred to a permutation null while PhenoDigm’s is a raw similarity, and PhenoDigm was not referred to the same null. The published table omits term pairs below a similarity cutoff, which then read as zero, so the recomputed configuration applies PhenoDigm to the full phenotype profile without that cutoff. The behavioral configuration restricts PhenoDigm to behavioral phenotypes, and the admitted-pair configuration applies the PhenoDigm scorer to FACET’s admitted pairs. These two and the published table as distributed are reported in S10 Table.

### Expert rating study

The sampling frame is the 14,205 of the 14,209 judged pairs on scored criteria whose mouse term carries a definition, of which 420 are admitted. From this, 168 pairs were drawn into two classes of 84: admitted pairs and refused pairs, stratified by proximity to admission (S1 Text). Quotas were proportional to the refused population, so most refused pairs lie in the stratum farthest from admission. The pairs form 12 blocks partitioned into three groups, so each of 12 raters rates 42 pairs. The design assigns every pair to three raters from three expertise groups (clinician-scientists, psychiatric researchers with rodent experience and mouse phenotyping specialists). The design called for a minimum of eight and up to 12 raters. Eight blocks had been returned at the time of this analysis (seven invited raters and the first author), so the study is interim and the achieved coverage is one to three ratings per pair. One invited rater (ARC) joined the author list after returning his block. He rated under the same blinding as every invited rater, and his block is counted among the seven invited returns. Each pair is shown singly with a paraphrase of the criterion, the anchor and the mouse definition. The other candidates of its bundle and its score are withheld, and the raters are blind to the verdict. Raters answer whether the mouse phenotype belongs to the criterion (yes, no, cannot tell) and how strong an analog it is (unrelated, weak, close, direct, unsure). A rating confirms admission when the answer is yes and the analog close or direct. The rating instrument was written by the first author and was not reviewed by the expert groups before use. It asked whether the mouse phenotype is evidence for the criterion. This paper says that a phenotype models a feature of the criterion. The estimands are the confirmation rate among admitted pairs and the rejection rate among refused pairs, conditional on the verdict. They are estimated over distinct pairs with Wilson intervals and a one-sided lower bound whose standard error adds between-rater to between-pair variance. The declared rule reads the judgment as expert-supported when both rates reach 0.85 with lower bounds of 0.70, as not expert-confirmed when either rate is below 0.70 and as measured otherwise. The 0.85 bar was set before any rating was returned and exceeds the agreement later observed between two raters on the same pair (0.77), so it is a demanding standard. “Cannot tell” is excluded in the primary analysis and counted as disagreement and as agreement in two sensitivity analyses. Agreement on the membership question alone, agreement between two raters on the same pair and Cohen kappa against the rater majority are reported descriptively. The first author rated one unreturned block after the first five invited returns had been analyzed. The instrument pairs were drawn by script from the sealed key, so the first author neither chose nor knew them, and those ratings enter the primary analysis with their removal reported as a sensitivity analysis.

### Validation of the LLM steps

Validation effort follows the number of decisions a step makes and how easily a person can audit them (Table 4). We selected alternate LLMs from two other providers as an LLM evaluator tends to favor outputs of its own model family [34]. In the re-attribution of anchors, each alternate LLM receives the primary prompt and schema, and agreement with the primary run on each anchor’s primary unit is reported as a proportion and a Cohen kappa [49]. In the reclassification of mouse terms, the sample is every term the primary run admits to a disorder plus 300 refused terms drawn at random (seed 20261020).

Agreement is reported per stratum [50] and as a stratum-weighted kappa. The admission negative control compares the PhenoDigm scorer applied to the admitted pairs with the same scorer applied to 100 random draws of refused pairs. Each draw is matched in number, in the filters passed, and in the annotation frequency of its mouse terms. The one-sided p is one plus the number of draws whose PhenoDigm-scorer AUROC reaches that of the admitted pairs, divided by 101. In the LLM swap, every pair with a primary verdict in the four primary disorders (20,875 pairs) was re-judged by each alternate LLM with the primary user prompt and schema. Only the verdict fields change, and aggregation and the null are rerun with the primary configurations. Before the swap, the same procedure run on the cached primary verdicts reproduced the pair table and the primary support AUROCs exactly. The pre-specified outcome is the paired difference in support AUROC with a gene-clustered 90% interval, read against a margin of 0.05 as equivalent, not equivalent or inconclusive. Pair-level agreement is reported as agreement and kappa. Rules of the form at least k of n LLMs (compared with the expert ratings) are post hoc. The panel of Results is the majority, field by field, of three independent runs (the primary LLM under the disorder-specific prompt, claude-opus-4-8 and gemini-2.5-pro), with the primary run breaking a three-way split. It is compared with the raters on the 140 rated pairs of the four primary disorders with a definite rating. Every instrument pair now carries an invited rating, so no pairs are held out and any rule chosen on them is exploratory.

### Label provenance and secondary analyses

The claim paper of a curated model is the paper whose disease claim MGI cites. Removal of the claim paper deletes claim-paper annotations from curated models and background genotypes alike (S1 Text). Relatives are uncurated, behaviorally annotated genotypes of a curated model’s gene that cite none of its claim papers. The top coverage tier is Diagnostic Coverage of three or more units for ASD and MDD and two for SCZ, IDD and NCD. The top-tier odds ratios of S14 Table adjust for the logarithm of one plus the MP term count and of one plus the behavioral term count. The behavioral-annotation odds ratios adjust for the logarithm of one plus the PubMed record count per gene. The IMPC-only test takes the IMPC-tagged genotypes of the evaluation set (allele symbol carrying IMPC, KOMP, EUCOMM or NCOM) with curated models removed. It ranks the genotypes whose gene is an H gene against the rest by the support score, Diagnostic Coverage, the behavioral count and PhenoDigm recomputed, with gene-clustered intervals. Candidate models outside curation are the uncurated genotypes whose support score is at least the score at rank 100, and tie blocks are kept whole. This is the maximum support score for ASD (261 genotypes) and IDD (337) and the top-ranked genotypes for SCZ (176) and NCD (100). Each candidate is compared with controls matched on behavioral depth and annotation source for whether PubTator3 tags any of its papers with the disorder. Database drift is assessed by rescoring the genotype set on later MGI and MP releases with the frozen verdicts. Rescored on MGI’s report of 2026-10-05 and MP release 2026-09-23, the coverage AUROC moves by 0.0006 or less and the support AUROC by 0.0013 or less in every disorder.

### Statistical analysis and software

All computation ran in Python 3 with NumPy [43], pandas [51], SciPy [52] and statsmodels [53]. Lexical matching used rapidfuzz and the figures matplotlib [54]. Package versions are pinned in the released environment file. Separation of classes is the AUROC as the Mann-Whitney statistic with ties contributing one half [55], with average precision beside each support AUROC. Every interval is a 95% percentile interval from a gene-clustered bootstrap of 2,000 replicates [56,57], and a paired difference between two scores carries the two-sided bootstrap p value of its replicates. The 16 primary claims are corrected by the Holm method [58]. Equivalence of two AUROCs is read from the 90% interval of their difference against a pre-specified margin of 0.05, the two one-sided tests procedure [59]. Rates of expert agreement carry Wilson intervals [50] and a one-sided 95% lower bound on a standard error that adds the between-rater variance. Chance-corrected agreement is Cohen kappa against the rater majority, Fleiss kappa [60] among LLM runs and quadratic weighted kappa on the analog grade. Per-genotype permutation p values use the estimator of Phipson and Smyth [48]. H-gene enrichment is a logistic regression with Wald intervals. Power follows Hanley and McNeil [41]. No test is run on individual genotypes, and analyses marked post hoc are uncorrected.

### Data and code availability

The per-criterion catalogue is at https://dsm-5-map-mouse.vercel.app. The pipeline code, the HPO anchor sets, the criterion-mapped MP taxonomy, the frozen primary verdicts, the configuration files, the prompts and the analysis log are deposited in a public repository at submission. Every number in the paper is reproducible from the frozen verdicts by deterministic steps. The pair-judgment step did not log the snapshot served under the LLM alias. Every logged call to the same alias between 2026-06-04 and 2026-11-08 was served gpt-5.4-2026-03-05 (about 23,000 calls from the other steps, including runs on the same days). All pair judgments ran within that window, so they were judged by that snapshot (Table 4). A fresh run reproduces the procedure but not the verdicts bit for bit, and the alternate-LLM re-judgments measure that dependence. The mapping step reproduced 74 of 75 gate terms on a fresh run (S22 Table). The deposit omits the free-text reasons the LLM logged, which can quote the manual from memory, and carries the configuration files and prompts with the criteria in paraphrase. SHA-256 checksums of the primary data files are listed in S2 Table, and the code release will carry its own checksums. Third-party inputs are used under their own licenses. MGI data and the Mammalian Phenotype Ontology are licensed under CC BY 4.0 by The Jackson Laboratory, and the Disease Ontology under CC0. The Human Phenotype Ontology is used under the HPO license: its files are deposited unmodified with their release date (2026-02-16), and the HPO Consortium is cited [23]. The anchor sets and the criterion-mapped taxonomy add mappings to HPO and MP terms and change no term, label or definition of either ontology. DSM-5-TR diagnostic constructs were used as the clinical reference framework. FACET is not affiliated with or endorsed by the American Psychiatric Association.

## Ethics statement

The expert rating study collected professional judgments on phenotype term pairs from colleagues who agreed to take part. It involved no patients, no intervention and no personal data beyond each rater’s name or initials, primary role, board certification in psychiatry and years since terminal degree in bands. On November 9, 2026 the SUNY Upstate Medical University Office of Research Compliance determined that the study is not human subjects research. It does not require Institutional Review Board submission.

## Competing interests

The authors have declared that no competing interests exist.

## Author contributions

Conceptualization: AL, CL. Methodology, software, formal analysis, data curation, visualization and writing of the original draft: AL. Investigation (a preliminary gene-level comparison of mouse phenotype and human genetic gene lists that preceded the pipeline): DN. Supervision and resources: CL. Investigation (expert rating, as an invited rater before joining): ARC. Review and editing: ARC, JH, CL.

## Acknowledgments

We thank the expert raters who returned ratings or agreed to rate pairs. From SUNY Upstate Medical University: Ma-Li Wong, Nevena Radonjic, Frank Middleton, Andrea Viczian and Yanli Zhang-James. From Western Sydney University: Tim Karl. From the University of California San Diego: Abraham Palmer. From the University of Tennessee Health Science Center: Qian Ge, Jianyang Du and Michael McDonald. We thank the Mouse Genome Informatics, Human Phenotype Ontology, Monarch Initiative and PubTator teams for the resources the pipeline reads.

## Supporting information

**S1 Text. Supplementary Methods and Supplementary Results**.

The Methods give the anchor shortlist retrievers, the eight mapping rules in full, gene clustering and the host-locus transgene rulings, the re-test rule for the score cap and the expert sampling and analysis. They also give the dated record of pre-specification. They also give claim-paper removal, the composition of the annotation record, the shared developer prompt and the disorder-specific sensitivity run, and the mirrored permutation null. The PhenoDigm configurations and the prompts of the primary analysis run follow. The Results give the sensitivity of the support AUROC, the support and retrieval scores at equal depth, the PhenoDigm configurations and the alternate-LLM admissions. They also give the upstream LLM steps, the expert ratings in detail, human genetics and the describing literature, the reproducibility of the verdicts, the IMPC-only test, the depth-stratified null with PhenoDigm referred to the same nulls, and further analyses.

**S1 Table. Criterion units, symptom domains and diagnostic rules per disorder.**

**S2 Table. Data files of the primary analysis run with checksums.**

**S3 Table. Label rules and curated models.**

**S4 Table. The 16 primary claims under Holm correction.**

**S5 Table. Genes implicated by human genetics without mouse data (H genes).**

**S6 Table. Analyses in the order of Results with their record files.**

**S7 Table. Annotation composition of curated models and of the genotype set.**

**S8 Table. Sensitivity of the support AUROC, with the retrieval baseline and depth matching.**

**S9 Table. Allele design.**

**S10 Table. FACET against PhenoDigm in four configurations.**

**S11 Table. The 138 attributable HPO anchors.**

**S12 Table. Stability of the upstream LLM steps.**

**S13 Table. Expert agreement, including the developer sensitivity analysis.**

**S14 Table. Phenotyping attention and the describing literature.**

**S15 Table. The highest-credited genotypes per disorder by Diagnostic Coverage.**

**S16 Table. Phenotype classes, their MP root terms and the admitted terms per class.**

**S17 Table. The IMPC-only test.**

**S18 Table. The depth-stratified null and PhenoDigm referred to the same nulls.**

**S19 Table. The admitted terms and the curated models’ annotations behind the SCZ A5 and MDD A1 and A2 credits.**

**S20 Table. The SCZ candidate pairs built on endophenotype and drug-response terms, by term family, anchor and analog grade.**

**S21 Table. The hand-curated anchor sets that gated mouse domain labeling, with their outcome in the primary taxonomies.**

**S22 Table. The anchor-gate rerun of 2026-11-08, 15 terms per disorder, against the expectations and the primary run.**

**S23 Table. Validation of the pair judgment by alternate LLMs and the admission control.**

**S24 Table. Curated labels, claim papers and genes, with the IMPC-only test.**

**S25 Table. Data sources of the primary analysis run.**

**S26 Table. The eight named mapping rules.**

**S27 Table. Fixed parameters of anchoring, retrieval, routing and judgment.**

**S28 Table. Criteria with no admitted phenotype in any genotype or in no curated model.**

**S29 Table. LLM admission against retrieval alone on the rated pairs, scored against the rater majority.**

**S30 Table. The judgment floor raised on the primary analysis run: admitted pairs and criterion matches lost, and support AUROC.**

**S31 Table. The channel weights changed on the primary analysis run: admissions kept, pairs newly routed to the LLM and support AUROC.**

**S32 Table. Precision and power of an AUROC by the number of curated models.**

**S33 Table. The pairs the alternate LLMs admitted on SCZ A1 to A3.**

**S1 Fig. Cross-disorder overlap of uncurated genotypes.**

**S2 Fig. Agreement with the curated labels under both measures, coverage AUROC and support AUROC per disorder.**

## Notes

### Competing Interest Statement

The authors have declared no competing interest.

## References

1. Garner JP. The significance of meaning: why do over 90% of behavioral neuroscience results fail to translate to humans, and what can we do to fix it? ILAR J. 2014;55(3):438–456. doi:10.1093/ilar/ilu047

2. Stanford SC. Some reasons why preclinical studies of psychiatric disorders fail to translate: what can be rescued from the misunderstanding and misuse of animal ‘models’? Altern Lab Anim. 2020;48(3):106–115. doi:10.1177/0261192920939876

3. American Psychiatric Association. Diagnostic and Statistical Manual of Mental Disorders, fifth edition, text revision. Washington, DC: American Psychiatric Association Publishing; 2022. doi:10.1176/appi.books.9780890425787

4. Willner P. The validity of animal models of depression. Psychopharmacology. 1984;83(1):1–16. doi:10.1007/BF00427414

5. Belzung C, Lemoine M. Criteria of validity for animal models of psychiatric disorders: focus on anxiety disorders and depression. Biol Mood Anxiety Disord. 2011;1:9. doi:10.1186/2045-5380-1-9

6. Nestler EJ, Hyman SE. Animal models of neuropsychiatric disorders. Nat Neurosci. 2010;13(10):1161–1169. doi:10.1038/nn.2647

7. Chadman KK, Yang M, Crawley JN. Criteria for validating mouse models of psychiatric diseases. Am J Med Genet B Neuropsychiatr Genet. 2009;150B(1):1–11. doi:10.1002/ajmg.b.30777

8. Gottesman II, Gould TD. The endophenotype concept in psychiatry: etymology and strategic intentions. Am J Psychiatry. 2003;160(4):636–645. doi:10.1176/appi.ajp.160.4.636

9. van den Berg H. Evaluating the validity of animal models of mental disorder: from modeling syndromes to modeling endophenotypes. Hist Philos Life Sci. 2022;44(4):59. doi:10.1007/s40656-022-00537-4

10. Percelay S, Lahogue C, Billard JM, Freret T, Boulouard M, Bouet V. The 3-hit animal models of schizophrenia: improving strategy to decipher and treat the disease? Neurosci Biobehav Rev. 2024;157:105526. doi:10.1016/j.neubiorev.2023.105526

11. Silverman JL, Thurm A, Ethridge SB, et al. Reconsidering animal models used to study autism spectrum disorder: current state and optimizing future. Genes Brain Behav. 2022;21(5):e12803. doi:10.1111/gbb.12803

12. Ornoy A, Echefu B, Becker M. Animal models of autistic-like behavior in rodents: a scoping review and call for a comprehensive scoring system. Int J Mol Sci. 2024;25(19):10469. doi:10.3390/ijms251910469

13. Gencturk S, Unal G. Rodent tests of depression and anxiety: construct validity and translational relevance. Cogn Affect Behav Neurosci. 2024;24(2):191–224. doi:10.3758/s13415-024-01171-2

14. Czeh B, Fuchs E, Wiborg O, Simon M. Animal models of major depression and their clinical implications. Prog Neuropsychopharmacol Biol Psychiatry. 2016;64:293–310. doi:10.1016/j.pnpbp.2015.04.004

15. von Mucke-Heim IA, Urbina-Trevino L, Bordes J, Ries C, Schmidt MV, Deussing JM. Introducing a depression-like syndrome for translational neuropsychiatry: a plea for taxonomical validity and improved comparability between humans and mice. Mol Psychiatry. 2023;28(1):329–340. doi:10.1038/s41380-022-01762-w

16. Silverman JL, Yang M, Lord C, Crawley JN. Behavioural phenotyping assays for mouse models of autism. Nat Rev Neurosci. 2010;11(7):490–502. doi:10.1038/nrn2851

17. Ey E, Leblond CS, Bourgeron T. Behavioral profiles of mouse models for autism spectrum disorders. Autism Res. 2011;4(1):5–16. doi:10.1002/aur.175

18. Feifel D, Shilling PD. Promise and pitfalls of animal models of schizophrenia. Curr Psychiatry Rep. 2010;12(4):327–334. doi:10.1007/s11920-010-0122-x

19. Roubertoux PL, Tordjman S, Caubit X, et al. Construct validity and cross validity of a test battery modeling autism spectrum disorder (ASD) in mice. Behav Genet. 2020;50(1):26–40. doi:10.1007/s10519-019-09970-x

20. Baldarelli RM, Smith CL, Ringwald M, Richardson JE, Bult CJ, Mouse Genome Informatics Group. Mouse Genome Informatics: an integrated knowledgebase system for the laboratory mouse. Genetics. 2024;227(1):iyae031. doi:10.1093/genetics/iyae031

21. Smedley D, Oellrich A, Kohler S, et al. PhenoDigm: analyzing curated annotations to associate animal models with human diseases. Database (Oxford). 2013;2013:bat025. doi:10.1093/database/bat025

22. Robinson PN, Kohler S, Oellrich A, et al. Improved exome prioritization of disease genes through cross-species phenotype comparison. Genome Res. 2014;24(2):340–348. doi:10.1101/gr.160325.113

23. Gargano MA, Matentzoglu N, Coleman B, et al. The Human Phenotype Ontology in 2024: phenotypes around the world. Nucleic Acids Res. 2024;52(D1):D1333–D1346. doi:10.1093/nar/gkad1005

24. Brown TB, Mann B, Ryder N, et al. Language models are few-shot learners. Adv Neural Inf Process Syst. 2020;33:1877–1901. arXiv:2005.14165

25. Ouyang L, Wu J, Jiang X, et al. Training language models to follow instructions with human feedback. Adv Neural Inf Process Syst. 2022;35:27730–27744. arXiv:2203.02155

26. Gilardi F, Alizadeh M, Kubli M. ChatGPT outperforms crowd workers for text-annotation tasks. Proc Natl Acad Sci USA. 2023;120(30):e2305016120. doi:10.1073/pnas.2305016120

27. Caufield JH, Hegde H, Emonet V, et al. Structured prompt interrogation and recursive extraction of semantics (SPIRES): a method for populating knowledge bases using zero-shot learning. Bioinformatics. 2024;40(3):btae104. doi:10.1093/bioinformatics/btae104

28. Toro S, Anagnostopoulos AV, Bello SM, et al. Dynamic retrieval augmented generation of ontologies using artificial intelligence (DRAGON-AI). J Biomed Semantics. 2024;15(1):19. doi:10.1186/s13326-024-00320-3

29. Abrahams BS, Arking DE, Campbell DB, et al. SFARI Gene 2.0: a community-driven knowledgebase for the autism spectrum disorders (ASDs). Mol Autism. 2013;4:36. doi:10.1186/2040-2392-4-36

30. Dolensek N, Gehrlach DA, Klein AS, Gogolla N. Facial expressions of emotion states and their neuronal correlates in mice. Science. 2020;368(6486):89–94. doi:10.1126/science.aaz9468

31. Langford DJ, Bailey AL, Chanda ML, et al. Coding of facial expressions of pain in the laboratory mouse. Nat Methods. 2010;7(6):447–449. doi:10.1038/nmeth.1455

32. Ongur D, Abi-Dargham A, Clarke DE, et al. The future of DSM: a report from the Structure and Dimensions Subcommittee. Am J Psychiatry. 2026;183(5):301–309. doi:10.1176/appi.ajp.20250876

33. Borsboom D. A network theory of mental disorders. World Psychiatry. 2017;16(1):5–13. doi:10.1002/wps.20375

34. Verga P, Hofstatter S, Althammer S, et al. Replacing judges with juries: evaluating LLM generations with a panel of diverse models. arXiv:2404.18796. 2024. doi:10.48550/arXiv.2404.18796

35. Smith CL, Eppig JT. The Mammalian Phenotype Ontology as a unifying standard for experimental and high-throughput phenotyping data. Mamm Genome. 2012;23(9-10):653–668. doi:10.1007/s00335-012-9421-3

36. Groza T, Gomez FL, Mashhadi HH, et al. The International Mouse Phenotyping Consortium: comprehensive knockout phenotyping underpinning the study of human disease. Nucleic Acids Res. 2023;51(D1):D1038–D1045. doi:10.1093/nar/gkac972

37. Putman TE, Schaper K, Matentzoglu N, et al. The Monarch Initiative in 2024: an analytic platform integrating phenotypes, genes and diseases across species. Nucleic Acids Res. 2024;52(D1):D938–D949. doi:10.1093/nar/gkad1082

38. Wei CH, Allot A, Lai PT, et al. PubTator 3.0: an AI-powered literature resource for unlocking biomedical knowledge. Nucleic Acids Res. 2024;52(W1):W540–W546. doi:10.1093/nar/gkae235

39. Schriml LM, Munro JB, Schor M, et al. The Human Disease Ontology 2022 update. Nucleic Acids Res. 2022;50(D1):D1255–D1261. doi:10.1093/nar/gkab1063

40. Elkan C, Noto K. Learning classifiers from only positive and unlabeled data. In: Proceedings of the 14th ACM SIGKDD International Conference on Knowledge Discovery and Data Mining. 2008:213–220. doi:10.1145/1401890.1401920

41. Hanley JA, McNeil BJ. The meaning and use of the area under a receiver operating characteristic (ROC) curve. Radiology. 1982;143(1):29–36. doi:10.1148/radiology.143.1.7063747

42. Bachmann M. rapidfuzz/RapidFuzz: release 3.13.0. Zenodo. 2025. doi:10.5281/zenodo.15133267

43. Harris CR, Millman KJ, van der Walt SJ, et al. Array programming with NumPy. Nature. 2020;585(7825):357–362. doi:10.1038/s41586-020-2649-2

44. Resnik P. Using information content to evaluate semantic similarity in a taxonomy. In: Proceedings of the 14th International Joint Conference on Artificial Intelligence. 1995:448–453. arXiv:cmp-lg/9511007

45. Jaccard P. The distribution of the flora in the alpine zone. New Phytol. 1912;11(2):37–50. doi:10.1111/j.1469-8137.1912.tb05611.x

46. OpenAI. Prompt engineering. OpenAI API documentation. https://developers.openai.com/api/docs/guides/prompt-engineering. Accessed 2026-09-07.

47. OpenAI. Best practices for prompt engineering with the OpenAI API. OpenAI Help Center. https://help.openai.com/en/articles/6654000-best-practices-for-prompt-engineering-with-the-openai-api. Accessed 2026-09-07.

48. Phipson B, Smyth GK. Permutation p-values should never be zero: calculating exact p-values when permutations are randomly drawn. Stat Appl Genet Mol Biol. 2010;9(1):Article 39. doi:10.2202/1544-6115.1585

49. Cohen J. A coefficient of agreement for nominal scales. Educ Psychol Meas. 1960;20(1):37–46. doi:10.1177/001316446002000104

50. Wilson EB. Probable inference, the law of succession, and statistical inference. J Am Stat Assoc. 1927;22(158):209–212. doi:10.1080/01621459.1927.10502953

51. McKinney W. Data structures for statistical computing in Python. In: Proceedings of the 9th Python in Science Conference. 2010:56–61. doi:10.25080/Majora-92bf1922-00a

52. Virtanen P, Gommers R, Oliphant TE, et al. SciPy 1.0: fundamental algorithms for scientific computing in Python. Nat Methods. 2020;17(3):261–272. doi:10.1038/s41592-019-0686-2

53. Seabold S, Perktold J. Statsmodels: econometric and statistical modeling with Python. In: Proceedings of the 9th Python in Science Conference. 2010:92–96. doi:10.25080/Majora-92bf1922-011

54. Hunter JD. Matplotlib: a 2D graphics environment. Comput Sci Eng. 2007;9(3):90–95. doi:10.1109/MCSE.2007.55

55. Mann HB, Whitney DR. On a test of whether one of two random variables is stochastically larger than the other. Ann Math Stat. 1947;18(1):50–60. doi:10.1214/aoms/1177730491

56. Efron B, Tibshirani RJ. An Introduction to the Bootstrap. Boca Raton: Chapman and Hall/CRC; 1994. doi:10.1201/9780429246593

57. Field CA, Welsh AH. Bootstrapping clustered data. J R Stat Soc Series B. 2007;69(3):369–390. doi:10.1111/j.1467-9868.2007.00593.x

58. Holm S. A simple sequentially rejective multiple test procedure. Scand J Stat. 1979;6(2):65–70. [no DOI; JSTOR 4615733]

59. Schuirmann DJ. A comparison of the two one-sided tests procedure and the power approach for assessing the equivalence of average bioavailability. J Pharmacokinet Biopharm. 1987;15(6):657–680. doi:10.1007/BF01068419

60. Fleiss JL. Measuring nominal scale agreement among many raters. Psychol Bull. 1971;76(5):378–382. doi:10.1037/h0031619

