## Supporting Information (S1 Text, S1 to S33 Tables, S1 and S2 Figures) for "FACET: Criterion-level face-validity of 32,535 mouse genotypes across five DSM-5-TR psychiatric disorders"

### Supporting information for FACET: criterion-anchored measurement of mouse model face validity for psychiatric disorders at scale

An Le, Chunyu Liu

S1 Text holds the Supplementary Methods and the Supplementary Results. Supplementary Tables S1 to S14 and Supplementary Figure S1 follow. Every number is computed from the frozen verdicts of the primary analysis run under the primary labels (85, 42, 121, 71 and 8 curated models for ASD, SCZ, IDD, NCD and MDD). Where this supplement says “of record” or “the record”, it refers to the primary analysis run. Main-text tables and figures are cited by their numbers in the article.

DSM-5-TR diagnostic constructs were used as the clinical reference framework. FACET is not affiliated with or endorsed by the American Psychiatric Association.

#### S1 Text, part 1. Supplementary Methods

##### SM1. The anchor shortlist retrievers

Each phenotype phrase returned by criterion decomposition is resolved against a shortlist of at most 12 HPO terms, and the LLM may choose only from that shortlist or answer NONE. The shortlist is the union of two retrievers, merged on a fixed weighting and truncated to 12. The retrievers and the merge are deterministic code (pilot_asd_to_hpo_hybrid_rev.py, functions fuzzy_candidates, semantic_candidates and merge_candidate_lists), called by the disorder-agnostic anchoring script (stage1_curate_hpo.py) with its default sizes of 8 lexical, 12 semantic and 12 merged candidates. The archived retrieval tables of the five anchoring runs hold exactly 8 lexical, 12 semantic and 12 merged candidates for every phrase (files 03, 04 and 05 of each run folder).

##### The HPO record

Every HPO term of the release of record (2026-02-16) is read from the ontology file with its identifier, label, synonyms (all synonym entries, whatever their scope) and textual definition. Identifiers are normalized to the HP:nnnnnnn form, and nodes without an HPO identifier are dropped.

##### The lexical retriever

The lexical retriever compares the phrase with the label and with each synonym of every HPO term, and it does not read definitions. An alias table holds one row for each label and one row for each synonym. The phrase and each alias are lowercased, and content tokens are the alphabetic words of three or more letters outside a short stop list. The stop list holds abnormal, impaired, reduced, difficulty, deficit, deficits, current, history, used, ability, abilities, pattern and patterns.

When the phrase has two or more content tokens, an alias sharing none of them scores zero. Every other alias scores 0.6 times the rapidfuzz token-set ratio plus 0.4 times the rapidfuzz weighted ratio, on a scale of 0 to 100. Each HPO term keeps its best-scoring alias (the label before a synonym at equal score), and the 8 highest terms form the lexical list.

##### The semantic retriever

The semantic retriever compares the phrase with the label, synonyms and definition of every HPO term through text embeddings. Each term is represented by one text holding its label, its synonyms and its definition, each on a labeled line. These texts and the phrase are embedded with text-embedding-3-large and scaled to unit length, so the dot product is the cosine similarity. The 12 terms with the highest cosine form the semantic list. The embedding index is cached and rebuilt only when the SHA-256 digest of the indexed texts or the embedding model name changes.

##### The merge and the truncation

The two lists are joined on the HPO identifier, so the union holds between 12 and 20 terms. The cosine is rescaled to 0 to 1 as (cosine + 1) / 2, and the lexical score is divided by 100. A term missing from the semantic list enters with a rescaled semantic score of 0, and a term missing from the lexical list enters with a lexical score of 0. The combined score is 0.65 times the semantic score plus 0.35 times the lexical score, plus 0.05 when both retrievers returned the term. Terms are sorted by combined score, then by cosine and then by lexical score. The shortlist is the top 12. Up to 8 retrieved terms are therefore dropped at truncation.

##### What the LLM receives and what is kept

The resolution call receives the phrase with its source fragment, category, centrality, polarity, temporality and notes. It also receives each shortlisted term with its rank, identifier, label, synonyms, definition and retrieval scores. The response schema enumerates the shortlisted identifiers and NONE, so no other identifier can be returned, and a code check repeats that restriction. A resolution enters the anchor set when its centrality is core or supporting, its polarity is present or bidirectional and its relation to the phrase is exact, broader than the phrase or narrower than the phrase. A proxy relation is excluded. The prompts and schemas are reproduced in section SM12.

##### Release gates

No hand-curated gate was applied to anchor resolution. The hand-curated gates belong to the mouse domain labeling step (section SM1.3).

##### SM1.3 The anchor sets that gated mouse domain labeling

Each disorder’s domain labeling script carries a hand-curated anchor set (the --pilot mode of the rebuild scripts, which runs the set, prints pass or fail per term and exits without writing). Each entry names an MP term, the symptom domain it should receive and whether it should be admissible for strict face validity. The sets hold 13 (SCZ), 17 (IDD), 17 (NCD) and 15 (MDD) terms, of which 5, 10, 14 and 11 are expected admissible and the rest are negative controls (mechanistic, generic neurological, developmental and non-neurological terms). ASD, the development disorder, carried instead a 62-term pre-declared pilot at the criterion step (taxonomy_v2_pilot/pilot_terms_ASD.tsv, 2026-07-27): 14 terms expected to become admissible under the criterion-mapped design, 16 expected to stay admissible and 16 expected to stay refused, with an acceptance rule fixed before the run (recovery at least 60%, retention and specificity at least 85%). It scored 14 of 14, 16 of 16 and 16 of 16 (taxonomy_v2_pilot/PILOT_RESULT.md). The pilot outputs were not kept, so S21 Table reads each set against the primary taxonomy (2026-11-07, read-only): the domain the term received at the first call, and whether the second call admitted it to a criterion unit (IDD and NCD) or the strict flag of record admits it (SCZ and MDD). The expected domain is recovered for 61 of 62 terms (the exception is abnormal body weight, which MDD routed to non-neurological). All 22 negative controls are refused. Of the 40 terms expected admissible, 32 are admitted. The eight misses are the soft grooming anchor in IDD and NCD, IDD’s two developmental-onset terms (criterion C is not a scored unit) and NCD’s three neuropathology terms (amyloid, tau and neurofibrillary tangles, removed as a face-validity domain by the NCD construct rule), so each miss is a design decision taken after the sets were written rather than a classifier error.

##### SM1.4 The anchor-gate rerun of 2026-11-08

The gates were rerun with 15 terms per disorder (75 terms, S22 Table), expectations fixed in audit/anchor_gate_2026-11-07/build_gate_sets.py before the run. ASD took 15 terms of its 62-term pilot (12 expected admissible across A1, A3, B1, B2 and B4, with seizures, tremors and kidney morphology as controls). IDD took its set of record minus the two developmental-onset terms, and NCD its set of record minus the three neuropathology terms plus spatial working memory, since those expectations are undefined under the design of record. SCZ took its 13 of record plus abnormal emotion/affect behavior and seizures, and MDD its 15 of record. ASD, IDD and NCD ran through the criterion-mapped classifier with fresh output folders, and SCZ and MDD through their domain-labeling pilots with fresh caches, so no cached answer of record was replayed. The classifier prompt hashes matched the primary analysis runs for ASD and IDD. For NCD the hash differed (46712a03 against 99110981 of record), because the NCD criterion specification was edited after its primary analysis run, so the NCD rerun certifies the current specification.

The rerun reproduces the primary verdict on 74 of 75 terms: the one change is MDD’s abnormal body weight, routed to weight and appetite change on the rerun and to non-neurological in the record. Against the pre-set expectations, 67 of 75 pass (ASD 11, IDD 14, NCD 14, SCZ 13, MDD 15). The eight misses are the soft grooming anchor refused in IDD and NCD as in the record, ASD’s social withdrawal routed to A3 with A1 secondary, abnormal grooming routed to B1 rather than B2 and abnormal startle reflex refused, all as in the record, and three expectation errors in the gate file itself (MP:0001400 is hyperresponsive to touch, correctly routed to B4, and the two SCZ additions, where the record also holds the emotion term as non-neurological at tier 2 and seizures refused under a different domain label). No term expected to be refused was admitted in any disorder.

##### SM2. The eight named mapping rules

Each mapping rule is a block of prompt text held in one module of the code release (criterion_rules.py). A disorder opts into a rule by listing its name in the extra_rules field of its criterion specification, and the builder appends the listed blocks to that disorder’s mapping prompt. The archived provenance file of each primary taxonomy run (00_prompt_provenance.json) records the list, and the archived developer prompt (00_developer_prompt.txt) records the text that ran. Every rule a disorder opts into appears verbatim in its archived prompt, and no rule appears in the prompt of a disorder that does not list it (S2 Table.1). The SHA-256 digest of each archived prompt equals the prompt hash recorded by its run, and every mapping call of the four runs returned the served LLM version gpt-5.4-2026-03-05.

*S2 Table.1. Grounding rules per disorder in the primary taxonomy runs. Calls are one per MP term in the run’s scope (section SM12). MDD’s mapping ran in June 2026 on an earlier prompt and carries no named rule.*

| Disorder | Run folder (2026-07-28) | Prompt hash (first 16 characters) | Rules opted in | Grounding calls |
| --- | --- | --- | --- | --- |
| ASD | taxonomy_v2_full/run_ASD | 679539aea9a31e99 | grooming_routing, involuntary_motor_sign, asd_construct_refusals | 1,655 |
| SCZ | taxonomy_v2_full/prod_SCZ_2026-07-28 | c6853a6bd0a58fda | scz_motor_boundary_from_dsm, mechanism_is_not_face_validity | 1,657 |
| IDD | taxonomy_v2_full/run_ID | f495ff735b361000 | human_schooling_constructs | 2,393 |
| NCD | taxonomy_v2_full/prod_NCD_2026-07-28 | 2330f79d059f16b1 | decline_versus_deficit, language_floor | 1,710 |
| MDD | June 2026 taxonomy | not applicable | none | not applicable |

The texts below are reproduced as they ran, with British spellings, capitals and internal punctuation unchanged, because they are records of the prompt and not prose of this paper. The prompts carry the criteria in the project’s paraphrase, which is deposited with the code release, and no DSM-5-TR text. Short criterion labels that this paper also uses (insistence on sameness, for example) are kept, as are quotations of MP definitions. Two rules state a lock date of 2026-07-27 in their heading. A backup copy of the module saved on 2026-07-28 holds seven of the eight rules with identical text and lacks the SCZ motor-boundary rule, which was therefore added on 2026-07-28 before the SCZ primary taxonomy run.

Four rules end in a review flag rather than a verdict for part of their scope. These are circling and head movements under the involuntary motor sign rule, reasoning and planning terms under the human schooling rule, vocalization terms under the language floor and maternal or reproductive terms under A5 in the SCZ motor-boundary rule. A flagged term is still assigned and is listed on an audit worksheet.

##### Grooming routing (ASD, configuration name grooming_routing)

GROOMING AND STEREOTYPY, A ROUTING RULE LOCKED 2026-07-27
Rodent grooming is a repetitive, syntactically organised motor sequence, and the field treats it as
stereotypy. Route grooming terms to B1, "movements that are stereotyped or repeated", which is what
grooming IS. Do NOT route grooming to B2. B2 is for insistence-on-sameness and inflexibility constructs
such as cognitive inflexibility, perseveration, and impaired reversal, where the item's language about
routines and rigid patterns applies directly. Nest building and other patterned home-cage sequences follow
the same rule: B1 if the term names the repetitive motor act, B2 only if it names inflexibility.

##### Involuntary motor sign (ASD, configuration name involuntary_motor_sign)

```
VOLUNTARY BEHAVIOUR VERSUS INVOLUNTARY MOTOR SIGN, A REFUSAL RULE LOCKED 2026-07-27
Every example DSM B1 gives (simple repeated motor acts, arranging toys in rows, turning objects over and over,
repeating the words of others, phrases with a private meaning) is an ORGANISED ACT THE INDIVIDUAL PERFORMS. B1 does not describe movement that
merely repeats. REPETITION ALONE IS NOT SUFFICIENT FOR B1.

An involuntary neurological motor sign is repetitive and is still not a stereotypy. It is produced by
pathology of the motor system rather than emitted as behaviour. REFUSE the following for B1 and for every
other item, and set modality=physiological:
  tremor, myoclonus, muscle spasm, muscle twitch, fasciculation, dystonia, chorea, athetosis, clonus,
  catalepsy and waxy flexibility, ataxia and other incoordination, paralysis, palsy and paresis,
  reflex-elicited postures such as limb grasping and hindlimb clasping, and seizures of every kind,
  including convulsive, nonconvulsive, absence, clonic, tonic, tonic-clonic, focal, febrile, audiogenic,
  psychomotor and induced seizures and every seizure threshold or susceptibility measure.

THIS IS A CONSTRUCT REFUSAL, NOT A VOCABULARY ONE, so the vocabulary realism rule above does NOT override
it. Do not admit any of these because the definition contains the words "repetitive", "cyclical",
"stereotyped" or "movement". MP defines tremors as "repetitive, cyclical movements of the body or a body
part; usually involuntary": the kinematics repeat, and the sign is still not a stereotypy. Read the WHOLE
definition, including any clause naming the phenotype as involuntary, uncontrolled, reflexive, elicited,
induced or a loss of coordination or power.

THE BOUNDARY, STATED POSITIVELY. ADMIT to B1 a term naming an act the animal performs: grooming,
barbering, scratching, nest building, digging, burrowing, marble burying, object manipulation, bar biting,
rearing, jumping, and stereotypic behaviour as such.

WHERE THE BOUNDARY IS GENUINELY AMBIGUOUS, FLAG IT RATHER THAN GUESSING. Some locomotor and head movement
terms have both a stereotypy reading and a vestibular or reflexive one, and MP's own definition often
names the second (for example, circling is defined as "repeated movement in a circle; often associated
with inner ear defects"). For circling, spinning, head bobbing, head shaking, abnormal head movements and
any similar term whose definition names an inner ear, vestibular, reflex or neurological association,
assign your best item AND SET needs_manual_review=true, saying in reason which reading you took and what
the competing reading is. These terms go to a human audit worksheet. Do not silently resolve them.
```

##### ASD construct refusals (ASD, configuration name asd_construct_refusals)

```
ADDITIONAL CONSTRUCT REFUSALS SPECIFIC TO THIS DISORDER
  - The term measures a COGNITIVE FACULTY no item names. Object RECOGNITION MEMORY is a memory assay; it
    is not "smelling or touching objects to excess". Refuse it even though the word "object" appears. The same
    goes for spatial learning and working memory: autism's criteria name none of them.
  - The term is a loss of sensory CAPACITY where the item describes a RESPONSE and MP does offer a
    response-framed alternative. Deafness, blindness and anosmia are capacity losses with behavioral
    alternatives available; refuse them.
  - The term is maternal, agonistic, dominance or sexual behavior and the item describes peer social
    interaction. Note that the same term may legitimately serve a DIFFERENT item; judge against the item
    you assign, not against autism in general.
```

##### Human schooling constructs (IDD, configuration name human_schooling_constructs)

```
HUMAN SCHOOLING AND PSYCHOMETRIC CONSTRUCTS, A REFUSAL RULE FOR THIS DISORDER
Criterion A names learning in school and is confirmed by intelligence tests that are standardized and
given individually. Criterion B names everyday activities in more than one setting, at home, at school,
at work and in the community. Reading, writing, arithmetic, IQ, school and workplace performance are HUMAN
INSTITUTIONAL constructs. No mouse vocabulary instantiates them, and none ever will.

When an item's construct is of this kind, do NOT reach for a loose surrogate. Set item_id to that item,
admissible=false, and construct_inadmissible=true, so the pipeline can report the construct as
UNASSESSABLE rather than as a measured zero. A measured zero says the models were tested and failed; an
unassessable says the question cannot be put to a mouse. Those are different claims and the difference is
the point of this project.

TWO SPECIFIC TRAPS, both observed in this project's own prior ID spine and both to be avoided here:
  - Dyscalculia and dysgraphia are not mouse constructs. They are the schooling surrogate, not the
    criterion, and admitting them manufactures coverage.
  - LANGUAGE IS NOT HEARING. A prior ID target for language impairment pulled a pool made entirely of
    auditory threshold and hearing loss terms. Auditory CAPACITY is not communicative function. Refuse
    hearing terms for any communication construct, and say so in reason.

Where a component IS instantiable, admit normally. Learning and problem solving have direct mouse assays;
reasoning, planning and abstract thinking are the contested middle and should be admitted only when the
term names the operation rather than a task that merely requires it, with needs_manual_review=true.
```

##### Decline versus deficit (NCD, configuration name decline_versus_deficit)

DECLINE VERSUS DEFICIT, THE VOCABULARY REALISM CASE FOR THIS DISORDER
Criterion A asks for a DROP FROM THE PERSON'S EARLIER LEVEL OF PERFORMANCE. That is a within-individual longitudinal
comparison. MGI genotype-phenotype annotation is a BETWEEN-GENOTYPE comparison against wild type, and MP
has no longitudinal frame at all: it cannot say "worse than this animal used to be".

DO NOT REFUSE A TERM FOR LACKING THE LONGITUDINAL FRAME. If the term names a deficit in the right
cognitive domain, admit it and set qualifier_absent=true. Refusing every term because none is longitudinal
would zero the disorder for a reason that is about the vocabulary rather than about the models.

This substitution is a known and disclosed concession of the project, not a discovery, and it is recorded
so the reader can discount it. Criterion A's numbered sub-items 1 and 2 are EVIDENCE REQUIREMENTS
(informant or clinician concern; standardized neuropsychological testing). They are not symptom domains
and nothing may be admitted to them.

##### Language floor (NCD, configuration name language_floor)

LANGUAGE, WHICH IS FLAGGED RATHER THAN FORCED
A project decision holds that the language domain is a floor for this disorder, meaning no mouse analog.
That decision is NOT encoded here as a refusal, because a rule that assumes its own answer cannot be
tested. Judge language terms on their merits. If you find yourself admitting a vocalization or
communication term to the language domain, do so AND set needs_manual_review=true, naming in reason what
you think the term instantiates. The human audit worksheet then adjudicates the floor against real
candidates rather than against an assumption.

Note the distinction the previous rule also draws: auditory capacity (hearing thresholds, deafness) is
not language. Refuse those for language regardless.

##### SCZ motor boundary from DSM (SCZ, configuration name scz_motor_boundary_from_dsm)

```
WHAT COUNTS AS A4, DECIDED FROM THE DEFINITION OF A4 GIVEN IN THIS UNIT AND NOTHING ELSE
The governing text is the paraphrase of DSM-5-TR's definition given in this unit's definition addendum:
motor behavior that is grossly disorganized or abnormal can take many forms, from childlike silliness to
agitation that cannot be predicted; any kind of goal-directed behavior can be affected; and catatonic
behavior is a marked fall in how much the person responds to the surroundings, running from resisting
instructions (negativism), through holding a rigid, inappropriate or odd posture, to no verbal or motor
response at all (mutism and stupor).

Read that text literally. It names BEHAVIOR, and specifically GOAL-DIRECTED behavior, at one pole, and a
MARKED FALL IN RESPONSIVENESS TO THE SURROUNDINGS at the other. Three consequences follow from that
definition, and no other disorder's rules bear on this one.

ADMIT, because the definition names them. Holding a rigid, inappropriate or odd posture: catalepsy, waxy
flexibility, abnormal or bizarre posturing, and immobility or stupor-like unresponsiveness. Resistance to
instructions and negativism. Agitation and disorganization of goal-directed behavior where the term names
the disorganization rather than the activity level.

REFUSE, because the definition excludes them, and set modality=physiological.
  - SEIZURES of every kind, including absence, convulsive, clonic, tonic, tonic-clonic, focal, febrile,
    audiogenic and psychomotor seizures, and every seizure threshold or susceptibility measure. A seizure
    is a paroxysmal ictal event. It is neither behavior nor a fall in responsiveness to the surroundings,
    and the A4 definition describes no such phenomenon.
  - EXAGGERATED REFLEX RESPONSES to environmental stimulation, including hyperekplexia and every startle
    measure. Catatonia is defined here as a marked FALL in responsiveness to the surroundings. An
    exaggerated startle is an INCREASE in reactivity, which is the opposite pole of the construct DSM
    names, not an instance of it.
  - ELEMENTARY MOTOR SIGNS produced by motor system pathology rather than emitted as behavior: tremor,
    myoclonus, muscle twitch, muscle spasm, fasciculation, dystonia, chorea, athetosis, clonus, paralysis,
    palsy and paresis. None is a form of goal-directed behavior and none is a fall in responsiveness to
    the surroundings.
  - BARE ACTIVITY LEVEL. The definition names agitation that cannot be predicted and disorganized goal-directed behavior.
    Plain hyperactivity, increased or decreased locomotor activity, and stimulus-induced hyperactivity name
    an activity LEVEL and not a disorganization. Refuse them unless the term itself names agitation,
    unpredictability, disorganization, or a loss of goal direction.

DO NOT ADMIT A TERM TO A4 SIMPLY BECAUSE THE ITEM IS BIDIRECTIONAL. Bidirectional means DSM names both a
decreased and an increased pole. It does not mean any motor abnormality qualifies.

A5 AVOLITION, THE SAME DISCIPLINE
The definition describes avolition as a drop in purposeful activity that the person starts and sustains
from their own motivation, and reduced emotional expression as less emotion in face, eye contact, speech
melody and gesture. Terms
naming MATERNAL, REPRODUCTIVE or NURTURING behavior, for example pup retrieval, nursing and nest building
for pups, are self-started purposeful activities on a plain reading, but they are also species-specific
reproductive behaviors whose deficit is routinely produced by circuitry unrelated to motivation. DSM does
not settle this. Assign your best item AND SET needs_manual_review=true, saying in reason that the
competing reading is a reproductive rather than a motivational deficit. Do not silently resolve it.
```

##### Mechanism is not face validity (SCZ, configuration name mechanism_is_not_face_validity)

THE ENDOPHENOTYPE IS NOT FACE VALIDITY, A HARD SEPARATION FOR THIS DISORDER
Prepulse inhibition, sensorimotor gating, latent inhibition, and psychotomimetic hyper-response to
amphetamine, MK-801 or PCP are the aberrant-salience ENDOPHENOTYPE. They are the substrate a symptom is
theorised to arise from, not the symptom. They must NEVER be admissible for any Criterion A item, however
strongly the literature associates them with this disorder.

This separation is load-bearing rather than fastidious. A term-attribution probe on this project's own
data found that prepulse inhibition plus sensorimotor gating alone carry 57 percent of the competing
method's best matches for this disorder, and endophenotype plus circuit carriers about 80 percent. Keeping
them off the face-validity axis is what makes the comparison between the two methods interpretable.

##### SM3. Gene clustering and the host-locus transgene rulings

Every interval in this paper resamples genes in place of genotypes, so that alleles of one gene are not counted as independent. Each genotype therefore receives a cluster key naming the gene or genes it mutates. The key is read from the allele symbols of the genotype (field 2 of the MGI genotype phenotype report of record, held pipe-delimited in the genotype summary table). The rules below are those of the analysis code of record (audit/geneclust_2026-10-08/common.py, function allele_gene) and of the operational definitions of 2026-10-08 (section 5).

##### Reading one allele symbol

The rules are applied to each allele symbol in the order given, and the first rule that matches decides.

1. A wild-type allele (a symbol ending in <+>) and the placeholders “+” and “?” carry no gene.
2. Seven named host-locus transgene alleles take the gene fixed by the rulings of S3 Table.2.
3. A transgene inserted at a named host locus (a symbol of the form Marker<Tg(construct)code>) carries no gene when its construct names a tool. The tool list holds recombinases (cre, Flp, Dre), transactivators (tTA, rtTA), reporters (lacZ, GFP, EGFP, YFP, EYFP, RFP, tdTomato, mCherry, Venus, luciferase, GCaMP), Cas9, diphtheria toxin A and the ERT fusion. Any other construct carries its payload gene, read as the last element of the construct after splitting on hyphens, slashes and commas, with any variant suffix after an asterisk removed.
4. A transgene at no named locus (a symbol of the form Tg(construct)code) follows the same rule.
5. A Rosa26 knock-in (a symbol of the form Gt(ROSA)26Sor) carries no gene when the construct names a tool. Otherwise it carries the payload gene of its parenthesized construct, or a Rosa26-specific key when no parenthesized construct is present.
6. Every other allele (a symbol of the form Marker) carries its marker symbol.

##### From alleles to a cluster key

Gene symbols are compared in upper case, so a human transgene payload and the mouse gene of the same name share one cluster. The key of a genotype is the sorted set of genes carried by its alleles, joined into one string. A genotype mutating two genes therefore forms a cluster of its own, separate from the clusters of either gene alone. A genotype whose alleles carry no gene (for example a genotype holding only a driver and a reporter) is its own singleton cluster. The bootstrap draws clusters with replacement from the whole universe of the analysis (2,000 replicates, seed 20261008, percentile intervals). A replicate holding fewer than two genotypes of either class is discarded and counted.

##### The host-locus transgene class

A host-locus transgene names the disrupted insertion site as its marker, so the marker need not name the biology of the genotype. The class was enumerated by the rule that an allele symbol has the form Marker<Tg(payload)LabCode>, with the payload read by a parenthesis-balanced scan so that nested constructs parse. Of the 32,535 genotypes of the scored set, 584 (1.80%) carry at least one such allele, in 71 distinct pairs of host marker and payload. The same rule applied to the whole MGI genotype phenotype report flags 1,091 genotypes.

*S3 Table.1. Payload classes of the 71 host-locus pairs, assigned by hand (audit_transgene_leakage/classify.py).*

| Payload class | Pairs | Genotypes | What the payload is |
| --- | --- | --- | --- |
| Driver | 14 | 482 | Recombinase or transactivator (cre, icre, creERT2, FLP, rtTA or tTS, Sleeping Beauty transposase) |
| Reporter | 14 | 17 | lacZ, EGFP or a Tyr coat-color marker |
| Tool | 1 | 8 | Itgax-HBEGF/EGFP diphtheria toxin receptor for cell ablation |
| Biological | 39 | 74 | A gene or viral protein whose expression is the purpose of the construct |
| Unresolved | 3 | 3 | Named insertional genotype with no payload gene in the symbol |

Driver payloads make up 82.5% of the class, and for them the host marker is almost always a bystander. For the largest driver group (Alb-cre at Speer6-ps1, 138 genotypes) the host is a pseudogene carrying a 4 bp lesion. The flag is a triage instrument that asks for the allele to be read, and it was never applied as a filter. Seven host-locus alleles were read against the MGI allele records and the primary literature on 2026-08-01. The cluster key fixes the gene of seven alleles by hand (S3 Table.2). Five of them are among the seven read, and the SNCA and LRRK2 alleles follow the canonical rulings of the same date.

*S3 Table.2. Host-locus rulings applied by the cluster key. “Payload” means the transgene’s payload gene names the biology, and “host” means the disrupted host locus does.*

| Allele symbol | Host marker | Payload | Ruling | Key | Basis |
| --- | --- | --- | --- | --- | --- |
| Atp2b1<Tg(Thy1-CHMP2B*)1Rene> | Atp2b1 | mutant human CHMP2B | Payload | CHMP2B | Integration in Atp2b1 intron 1, MGI models frontotemporal dementia (DOID:9255), Vernay 2016 |
| Zbtb20<Tg(PDGFB-APPSwInd)20Lms> | Zbtb20 | human APP (Swedish and Indiana) | Payload | APP | Genotype J20, Zbtb20 heterozygotes tested directly and the host excluded (Gulbranson 2021) |
| Trib2<Tg(Thy1-Trh/APP*)2.1Ingm> | Trib2 | Trh and amyloid beta fusion | Payload | APP | Genotype TBA2.1, no integration effect found (Alexandru 2011). The payload is not full-length APP |
| 2310039L15Rik<Tg(Prnp-SNCA*A53T)23Mkle> | 2310039L15Rik | human SNCA A53T | Payload | SNCA | Canonical ruling of 2026-08-01 |
| Khdrbs2<Tg(LRRK2*R1441G)135Cjli> | Khdrbs2 | human LRRK2 R1441G | Payload | LRRK2 | Canonical ruling of 2026-08-01 |
| Nbea<Tg(GH1)240BNec> | Nbea | human GH1 | Host | NBEA | Mixed in the literature. Neuromuscular block and perinatal lethality are attributed to Nbea (Su 2004), and dwarfism, adiposity and bone phenotypes to ectopic GH (Nuytens 2014) |
| Maoa<Tg(H2-K1-Ifnb1)8Seif> | Maoa | Ifnb1 | Host | MAOA | A 17 kb deletion spans Maoa exons 2 and 3, MGI types the allele null, and MGI annotates both genotypes to MAOA deficiency (Brunner syndrome, OMIM:300615) citing Cases 1995 |

Two further alleles read in that audit need no entry in the ruling list, because the general rules already assign them. Alb-cre at Speer6-ps1 is a driver and carries no gene. The MAPT P301L responder at Fgf14 (rTg4510) carries its payload, MAPT, although a host deletion is a published contributor to its phenotype (Gamache 2019).

The Maoa ruling shows why the class is triaged by reading the allele and not filtered by rule: three of the seven alleles read are not resolved by the structural rule alone.

##### SM4. The re-test rule for the score cap

The support score has a cap because the permutation p value cannot fall below 1/10,001 at 10,000 permutations. In the evaluation set 323 ASD genotypes (62 of them curated) and 380 IDD genotypes (43 curated) share that smallest p value and the capped score of 4. SCZ and NCD do not reach the cap (support score at rank 100 of 3.222 and 2.796). The AUROC (computed with midranks) and average precision (thresholds grouped by distinct score) treat a tied block as one rank and are unaffected. A rank-k statement and the order of the released catalogue are not defined inside the block, so the order inside the cap was pre-specified on 2026-10-19 before any within-cap order was computed (audit/ties_2026-10-19/PREREG_TIES_2026-10-19.md).

##### The pre-specified rule

The primary tie-break is deterministic and uses the permutation null of record. Inside the cap, genotypes are ordered by the standardized excess of the observed support over the null, z = (observed score minus null mean) / null standard deviation. The null mean and standard deviation are those of the genotype’s annotation count in the per-count null primary table, in strict mode. The secondary key is the margin of the observed score over the null maximum. Outside the cap the record order is kept, so the tie-free score is the pair (negative log10 p, z) in lexicographic order.

The check reruns the permutation step with 1,000,000 permutations for ASD and IDD, with the seed and inputs of record. Its outcome is the Spearman correlation, inside the cap, between the order by z and the order by the 1,000,000-permutation p value. The decision rule was declared in advance. If that correlation is 0.8 or more, z orders the catalogue and the rerun is reported as its check. Otherwise the rerun order is used and z is reported as the deterministic alternative.

##### Gates and results

Both gates passed with 0 mismatches for ASD and IDD. The negative log10 p of the stored permutation files equals the support score of the primary table for every genotype of the evaluation set. The observed scores and counts of every capped genotype in the rerun equal the record. The results are given in S4 Table.1, computed under the labels of 2026-10-08, which equal the primary labels for ASD and IDD.

*S4 Table.1. Ordering inside the support-score cap.*

| Quantity | ASD | IDD |
| --- | --- | --- |
| Genotypes at the cap with 10,000 permutations (curated) | 323 (62) | 380 (43) |
| Still at the cap with 1,000,000 permutations (curated) | 314 (61) | 326 (43) |
| Distinct rerun values inside the original cap | 6 | 12 |
| Spearman correlation, z order against the rerun order | 0.232 | 0.417 |
| Order selected by the pre-specified rule | Rerun order | Rerun order |
| Genotypes left tied after ordering by z (identical observed score and count) | 45 | 79 |
| Spearman correlation, z against the margin over the null maximum | 0.563 | 0.596 |
| Curated models among the top 50, 100 and 200 by (score, z) | 10, 17, 38 | 3, 5, 18 |
| Precision at 50 (Wilson 95% interval) | 0.20 (0.11 to 0.33) | 0.06 (0.02 to 0.16) |
| Share of curated models in the whole capped block | 0.192 | 0.113 |

A hundredfold increase in permutations leaves 97% of the ASD block and 86% of the IDD block tied again, because their observed scores exceed every null draw. The order the rule selects is therefore itself almost one tie, a case the rule did not anticipate. The rerun moved one curated ASD model and no curated IDD model below the cap.

##### Reporting adopted

The capped block is reported as a tie in every rank statement. The catalogue lists it as a tie block, ordered by z as a labeled secondary order, and the rerun is reported as showing that the ties are real and not an artifact of resolution. This departs from the order the pre-specified rule selects and is reported here as a deviation. Ordering by z does not concentrate curated models: precision at 50 equals the block-wide rate for ASD and falls below it for IDD.

##### SM5. Expert sampling, instrument and analysis

The expert study measures how often a human expert makes the same call as the pipeline on a pair the LLM judged. Its design, sampling code and decision rule were fixed on 2026-09-13, before any rater saw the instrument (RATER_DESIGN_RATIONALE_2026-09-13 and _rating_pilot_build_v9.py, build v9_k3_168pairs_n12_judged). Only aggregates are reported here, and no verdict on a single pair is given.

##### Sampling frame

A pair is rateable when the LLM returned a verdict on it, its anchor belongs to a scored criterion unit and its mouse term carries a definition a rater can read. Of the 14,209 judged pairs on anchors of scored criteria, 14,205 meet the last condition. The four without a definition are SCZ pairs. Of the 14,205, 420 are admitted (prevalence 0.0296) and 13,785 are refused. Pairs on anchors of unscored text enter no score, so a rating of them would check nothing the paper reports, and they are outside the frame. So are retrieved pairs that the LLM never judged.

##### Strata by proximity to admission

Every rateable pair falls in one stratum, assigned in order from the LLM’s verdict fields and the mapping of the mouse term.

- S1: strictly eligible (the admitted class, 420 pairs).
- S2: broadly but not strictly eligible.
- S3: neither, but the mapping step assigned the mouse term to a criterion (relevance tier 1).
- S4: neither, with an analog grade and a mouse term outside tier 1.
- S5: no analog grade (no pair in the frame).

*S5 Table.1. The sampling frame by disorder and stratum (_rating_pilot_2026_v9/population.json). Drawn counts are per class.*

| Disorder | Rateable | S1 (admitted) | S2 | S3 | S4 | Drawn admitted | Drawn refused |
| --- | --- | --- | --- | --- | --- | --- | --- |
| ASD | 4,925 | 125 | 761 | 504 | 3,535 | 24 | 24 |
| SCZ | 3,209 | 50 | 392 | 320 | 2,447 | 18 | 18 |
| IDD | 3,201 | 108 | 677 | 311 | 2,105 | 18 | 18 |
| NCD | 2,607 | 95 | 411 | 259 | 1,842 | 16 | 16 |
| MDD | 263 | 42 | 104 | 49 | 68 | 8 | 8 |
| Total | 14,205 | 420 | 2,345 | 1,443 | 9,997 | 84 | 84 |

The two classes were drawn evenly, because a draw in proportion to the frame would hold two or three admitted pairs. The disorder allocation (24 ASD, 18 SCZ, 18 IDD, 16 NCD and 8 MDD per class) gives every disorder at least eight admitted pairs. The refused class was drawn under one quota across S2 to S5, set in proportion to the frame’s refused pairs by largest remainder (seed 20260913). The refused sample therefore follows the frame and consists mostly of S4 pairs, with the near-miss strata S2 and S3 in their frame proportions. Among refused pairs with a definite rating from the eight raters, 13 are S2, 8 are S3 and 54 are S4.

##### Blocks, partitions and raters

The 168 pairs form 12 blocks of 14 (7 admitted and 7 refused, with disorders dealt in turn). The 12 blocks are partitioned three ways into four triples of blocks. Rater codes A to D take one triple each from the first partition, E to H from the second and I to L from the third. Every block appears once in each partition, so every pair reaches three raters (one per partition), and two raters from different partitions share at most one block. Each rater rates 42 pairs (21 admitted and 21 refused) in two parts of 22 and 20 pairs, ordered so that a rater who stops after part one still returns a balanced sample. Three attention anchors sit in part one: two obvious matches and one obvious non-match. A rater who misses an anchor is discarded and the run repeated, and anchors enter no statistic. A session is 45 screens and about 30 minutes.

##### The instrument

Each screen shows a paraphrase of the DSM-5-TR criterion, the HPO anchor label and the MP term label with its definition. The instructions state that criterion wording is paraphrased from the DSM-5-TR and not quoted. The screen never shows the LLM’s verdict, any score, the other candidates of the pair’s bundle or the genotype. Two questions are asked. The first author wrote the questions, the answer scales and the instructions, and no expert group reviewed them before use.

1. Does this mouse phenotype belong to the criterion? (Is it evidence about what the criterion describes, directly or indirectly? This is the instrument’s own wording.) Answers: yes, no, cannot tell.
2. How closely does the mouse term map onto the human term? Answers: unrelated, weak, close, direct, unsure.

A rating confirms admission when the first answer is yes and the second is close or direct, and any other definite rating rejects. The first question alone is also compared with whether the mapping step assigned the mouse term to the criterion (the criterion-match comparison).

##### Estimands and the declared rule

The estimands are the confirmation rate among admitted pairs and the rejection rate among refused pairs, each conditional on the pipeline’s verdict, so neither depends on the rarity of admission. Each pair contributes the share of its raters who agree with the pipeline (the mean rate), and the share of pairs with a majority in agreement is reported beside it. Rates are estimated over distinct pairs with two-sided Wilson 95% intervals. The one-sided 95% lower bound is the point rate minus 1.645 standard errors. That standard error is the empirical one over pair means, with a between-rater component added (the rater-adjusted bound).

The declared rule reads the pair judgment as expert-supported when both point rates are at or above 0.85 and both lower bounds at or above 0.70. It reads it as not expert-confirmed when either point rate is below 0.70, and as measured otherwise. “Cannot tell” is excluded in the primary analysis and counted as rejection of the pair in a pre-specified sensitivity analysis. Counting it as acceptance is a further sensitivity analysis. Agreement between two experts on the same pair, Cohen kappa against the rater majority and the agreement coefficient AC1 are descriptive and carry no threshold.

The study was sized on the 0.70 line. A half-width of 0.10 at a true rate of 0.70 needs 81 pairs per class. The 84 drawn give a half-width of 0.098 at 0.70 and 0.076 at 0.85 under the single-rating formula.

##### Results at eight raters (aggregates)

The study is interim: eight of the 12 blocks had been returned at the time of this analysis. Eight raters returned clean runs with no anchor missed: seven of the 12 invited raters (B, C, D, E, G, H and L) and the first author, who rated block A after the first five invited returns had been analyzed. One of the seven invited raters later joined the author list of the paper. That block was rated blind, under the same conditions as every invited block, before its rater joined. The 336 ratings cover all 168 pairs. Of these ratings, 68 (20%) are “cannot tell”, and 152 pairs carry a definite composite rating. The instrument pairs were drawn by script from the sealed key, at random within each stratum, so the first author neither chose nor knew them. The ratings of block A enter the primary analysis and their removal is a sensitivity analysis.

*S5 Table.2. Agreement with the pipeline, eight raters, primary analysis (“cannot tell” excluded).*

| Comparison | Class | Distinct pairs | Ratings | Rate | Wilson 95% | Rater-adjusted lower bound |
| --- | --- | --- | --- | --- | --- | --- |
| Composite | Confirmation (admitted) | 77 | 138 | 0.662 | 0.551 to 0.758 | 0.517 |
| Composite | Rejection (refused) | 75 | 130 | 0.884 | 0.793 to 0.939 | 0.824 |
| Criterion match | Confirmation | 74 | 132 | 0.788 | 0.682 to 0.866 | 0.711 |
| Criterion match | Rejection | 78 | 136 | 0.733 | 0.625 to 0.818 | 0.657 |

The rule returns not expert-confirmed, because confirmation (0.662) is below 0.70. The sized half-width on the primary comparison is 0.106 against the target of 0.10, and the widest rater-adjusted half-width is 0.173, carried by between-rater spread (tau 0.211 on admissions). Counting “cannot tell” as rejection of the pair gives confirmation 0.528 (lower bound 0.362) and rejection 0.923 (lower bound 0.883) over pairs, or 91 of 168 and 154 of 168 over ratings. Counting it as acceptance gives 121 of 168 and 116 of 168 over ratings. The LLM agrees with the rater majority on 0.793 of the 140 rated pairs of the four primary disorders (kappa 0.635). Two raters rating the same pair agree 0.792 of the time (107 pairs, pair bootstrap 95% interval 0.713 to 0.866). By the same estimator the LLM agrees with a single rater on 0.772 of ratings (268 ratings on 152 pairs, 0.712 to 0.833). On the 107 pairs rated twice or more, the LLM agrees with a single rater on 0.776 of ratings, and the difference from the agreement between raters is -0.016 (95% interval -0.093 to 0.053). Two raters agree on 0.697 of the pairs the LLM admitted (56 pairs) and on 0.898 of the pairs it refused (51 pairs). Source: audit/rater_single_agreement_2026-11-13/. Weighted back to the frame, AC1 is 0.886 and kappa 0.253, which are descriptive only.

Without block A (seven invited raters, 294 ratings), confirmation is 0.654 (77 pairs, 118 ratings, Wilson 0.542 to 0.750, lower bound 0.492) and rejection 0.880 (75 pairs, 111 ratings, 0.787 to 0.936, lower bound 0.815), with the same verdict. The seven-rater and eight-rater values are reported side by side in S13 Table.

##### SM6. Claim-paper removal, relatives and the attention control

Claim-paper removal asks how much of the annotation FACET reads for a curated model comes from a paper other than the one that claimed it as a model. It was pre-specified on 2026-10-13 (audit/open_items_2026-10-13/PREREG_OPEN_ITEMS_2026-10-13.md, item 2). Its reading rule was fixed with it. A drop measures how far the annotation of curated models comes from the claim paper and is not by itself a test of FACET.

##### Definitions

- Claim references. For a curated model of disorder D, the references (MGI J numbers) that support the disease rows of its MGI genotype page whose Disease Ontology term belongs to D, mapped to PubMed identifiers. They were read from 16,021 fetched genotype pages, and 2 J numbers could not be mapped.
- Annotation references. For each MP annotation, the PubMed identifiers on its rows of the snapshot report of record.
- Variant B (primary). An annotation is removed from a curated model only when every reference supporting it is a claim reference. An annotation backed by any other reference stays.
- Variant A (strict). An annotation is removed from a curated model when any of its references is a claim reference.
- Symmetric control. Every background genotype loses the annotations supported only by its own reference with the most annotations, so that background genotypes lose their main describing paper as curated models lose theirs.

After removal, Diagnostic Coverage is rebuilt from the frozen verdicts and the support score is recomputed through aggregation and the permutation null. Both AUROC families are then evaluated with gene-clustered intervals. The paper reports variant B with the symmetric control.

##### Results under the primary labels

For most curated models every annotation reference is a claim reference: ASD 78 of 85, SCZ 34 of 42, IDD 105 of 121 and NCD 52 of 71 (MDD 7 of 8). Of the 85 curated ASD models, 74 cite a single paper in all their annotations. After removal under variant B, 4 of 85 ASD, 6 of 42 SCZ, 8 of 121 IDD and 6 of 71 NCD models keep any credited unit (MDD 1 of 8).

*S6 Table.1. Agreement with the curated labels after claim-paper removal (variant B with the symmetric control). Values are AUROCs with 95% gene-clustered intervals.*

| Family | ASD | SCZ | IDD | NCD |
| --- | --- | --- | --- | --- |
| Convergent, after removal | 0.504 (0.486 to 0.526) | 0.542 (0.494 to 0.603) | 0.520 (0.494 to 0.546) | 0.521 (0.495 to 0.548) |
| Discrimination, after removal | 0.487 (0.468 to 0.506) | 0.486 (0.484 to 0.487) | 0.509 (0.486 to 0.535) | 0.505 (0.483 to 0.532) |

Without the symmetric control, variant B gives coverage AUROCs of 0.473 for ASD, 0.490 for SCZ and 0.492 for IDD, and variant A gives 0.473, 0.479 and 0.487. Variant A leaves 4, 5 and 7 curated ASD, SCZ and IDD models with any credited unit. The SCZ support AUROC interval lies below 0.5 because no curated SCZ model keeps a nonzero support score after removal. Curated models with nonzero support fall from 81 to 3 for ASD, from 19 to 0 for SCZ and from 68 to 6 for IDD. The symmetric control leaves background genotypes with nonzero support at 753 of 1,921 for ASD, 355 of 1,039 for SCZ and 454 of 2,097 for IDD. Values without the symmetric control and the support counts for NCD were computed under the labels of 2026-10-08 (73 NCD positives) and are not restated here.

##### Relatives and the attention control

Relatives test whether annotation from papers that made no disease claim carries the disorder’s criteria, at the level of the gene. They were pre-specified on 2026-10-15 (Amendment 7 of the same document).

- A curated gene of disorder D is the cluster key (section SM3) of any curated model of D.
- A relative is a behaviorally annotated genotype of the scored set that carries a curated gene’s key and is not a curated model of D. Its annotation references include no claim reference for D of any curated model of that gene.
- The background is the behaviorally annotated genotypes that are not curated for D and whose key is no curated gene of D.
- The attention control is the behaviorally annotated, uncurated genotypes of genes on the disorder’s human-genetics list (H genes) that have no curated model. A transfer that persists against it cannot be explained by a gene being suspected for the disorder.

Relatives are compared by the support primary score inside the evaluation set as a depth-stratified AUROC. The strata are behavioral annotation counts of 1 to 2, 3 to 5, 6 to 10 and 11 or more, weighted by the relatives in each stratum, with gene-clustered percentile intervals (2,000 replicates, seed 20261008). Against background, relatives score 0.567 (0.491 to 0.644) for ASD, 0.591 (0.516 to 0.647) for SCZ, 0.614 (0.554 to 0.678) for IDD and 0.582 (0.525 to 0.633) for NCD. Against the attention control they score 0.498 (0.428 to 0.564), 0.573 (0.457 to 0.640), 0.548 (0.497 to 0.613) and 0.557 (0.484 to 0.630).

##### SM7. Composition of the annotation record and study attention

##### Composition of the annotation record

Behavior is about 7% of the annotated terms of the 32,535 scored genotypes. Non-neural physiology and morphology make up 64.0%, mortality and growth 13.4% and nervous-system morphology 8.1%. Within behavior, motor terms make up 4.2% of all annotated terms, cognitive terms 0.8, emotional terms 0.5 and social or vocal terms 0.4, with other behavioral terms the remainder. Of the 32,535 genotypes, 73.8% carry no annotation at or below abnormal behavior (MP:0004924). A genotype is behaviorally annotated when it carries at least one such annotation, and this is the only behavior restriction of record.

*S7 Table.1. Behavioral annotation by class. Percent of curated models with any annotation in the named branch, against 26.2% of all 32,535 genotypes behaviorally annotated.*

| Disorder | Curated models | Behaviorally annotated, percent | Branch named in the text, percent with any annotation in it |
| --- | --- | --- | --- |
| ASD | 85 | 97.6 | Social: 90.6 |
| SCZ | 42 | 100 |  |
| IDD | 121 | 83.5 | Cognitive: 49.6 (60 of 121) |
| NCD | 71 | 87.3 | Cognitive: 31.0 (22 of 71) |
| MDD | 8 | 100 |  |

Genotypes phenotyped only by IMPC are behaviorally annotated about twice as often as genotypes described in the literature (46.8 against 23.2%). Their behavioral terms are mostly motor (7.8% of their terms) and ear or eye terms (9.5%), with 0.3% social terms. No curated model of any disorder is a genotype phenotyped only by IMPC.

##### The record carries abnormal results only

The MGI genotype phenotype report read by the pipeline has seven fields (allelic composition, allele symbols, genetic background, MP identifier, PubMed identifier, marker and genotype) and no assay, qualifier or note. MGI curates normal results and assay notes on its genotype pages, and the bulk report omits them. For example, the genotype page of Shank3 homozygotes (MGI:5056404) flags normal prepulse inhibition, startle and novel object recognition, none of which appears among its 19 rows in the report. The only normal-result signal in the report is the generic term no abnormal phenotype detected (MP:0002169), carried by 4,001 genotypes. A sex-specific normal flag in a second MGI report marks 109 scored genotypes, 36 of them among the 24,014 genotypes without behavioral annotation (0.15%). Absence of a behavioral annotation therefore means that no abnormal behavioral result was recorded, never that behavior was not tested.

##### The IMPC bound on genotypes assayed with normal results

IMPC statistical results record every analyzed parameter of an IMPC genotype, normal or not. They give a lower bound on the genotypes without behavioral annotation that were assayed and found normal. The behavioral IMPC procedures were listed before any outcome was computed (66 of 275 procedure entries, 16 distinct procedure names), under a protocol pre-specified on 2026-10-09.

Of the scored genotypes, 3,688 completed at least one successful behavioral IMPC procedure (median 62 behavioral parameters per genotype). Of these, 48.7% (Wilson 95% interval 47.1 to 50.3) carry no abnormal behavioral MGI annotation. Therefore 1,795 of the 24,014 genotypes without behavioral annotation (7.5%) went through a standard battery and were found normal. For 103 of the 1,795, IMPC records a significant behavioral call that never became an MGI behavioral annotation.

MGI’s own normal flags were estimated from a random sample of 2,000 genotypes without behavioral annotation, parsed from their genotype pages. In that sample 1.25% (0.85 to 1.84) carry a recorded normal behavioral result, implying about 300 (204 to 442) of the 24,014. With these added, about 2,100 (8.7%) are assayed normals, although the two sets may overlap. The scored set requires four abnormal annotations, so IMPC genotypes with mostly normal results are underrepresented in it and the bound is conservative.

##### Annotation detail

Annotation detail measures how specific a genotype’s behavioral terms are, beyond how many it carries. It was pre-specified on 2026-10-09 (audit/detail_sensitivity_2026-10-09/PREREG_DETAIL_SENSITIVITY_2026-10-09.md) over each genotype’s distinct annotations at or below MP:0004924.

- D1 (primary): the mean propagated information content, where IC(t) is minus log2 of the share of the 76,060 MGI genotypes annotated to t or a descendant.
- D2: the number of terms at 11.0 bits or more. The cut was calibrated on SCZ catatonia terms only, and its generality is unproven.
- D3: the maximum propagated information content.
- D4: the mean is_a depth below MP:0004924 (shortest path).

Detail tertiles use D1 cut points of 6.968 and 8.460 bits, computed once over the 8,521 behaviorally annotated scored genotypes and shared by every disorder. Matching on count and detail uses 12 cells (4 behavioral-count strata by 3 D1 tertiles).

Among behaviorally annotated genotypes, curated ASD, SCZ and MDD models carry more detailed behavioral terms beyond their term count. The odds ratios for curated status per standard deviation of D1 are adjusted for the logarithm of the behavioral annotation count. They are 1.76 (1.35 to 2.29) for ASD, 1.85 (1.35 to 2.54) for SCZ, 3.60 (1.81 to 7.17) for MDD and 0.91 (0.71 to 1.17) for IDD. Genotypes phenotyped only by IMPC average 6.98 bits against 7.96 for literature genotypes. Weighting genotypes by annotation detail changes the support AUROC by 0.014 or less in every disorder, and every interval of the difference includes 0. The NCD values of this analysis were computed under the labels of 2026-10-08 (73 NCD positives) and are not restated here.

##### Study attention

Study attention measures how much a gene has been studied, independently of its phenotypes. It was pre-specified on 2026-10-09 (audit/mgi_additional_2026-10-09/PREREG_MGI_ADDITIONAL_2026-10-09.md).

- Gene attention is the number of distinct PubMed identifiers per marker in the MGI marker reference report.
- Genotype attention is the maximum over the genes of its cluster key, entered as the logarithm of one plus the count.
- The secondary covariate is the IMSR strain count per gene, built the same way.
- Genotypes with no mapped gene are excluded from attention models.
- Attention tertile cuts over behaviorally annotated scored genotypes with a mapped gene are 3.526 and 5.147 on the log scale.

The H-gene lists come from human genetic studies that used no mouse data: 125 ASD genes, 49 SCZ genes, 934 IDD genes and 186 MDD genes (S5 Table). NCD has no such list and uses its MGI disease-gene associations. Uncurated genotypes of H genes are behaviorally annotated more often than other uncurated genotypes for ASD (35.4 against 25.8%) and MDD (35.1 against 26.0), and not for SCZ (28.0 against 26.1) or IDD (25.4 against 26.1). Adjustment for attention does not remove the excess. The odds ratios for behavioral annotation move from 1.60 to 1.83 for ASD and from 1.55 to 1.68 for MDD (unadjusted to adjusted), and from 0.98 to 1.13 for IDD and 1.11 to 1.08 for SCZ.

##### SM8. The shared developer prompt and the disorder-specific sensitivity run

##### The primary prompt

The pair primary judgment ran on one developer prompt for all five disorders (reproduced in section SM12). That prompt names major depressive disorder as the pipeline’s disorder and lists the nine MDD criterion domains for context. The disorder-specific content of each run is carried by the user prompt, which gives the anchor label, the anchor’s criterion phrase, a domain label and one row per candidate. Each disorder’s configuration also holds a developer-prompt addendum (105, 1,424, 1,840, 2,212 and 961 characters for ASD, SCZ, IDD, NCD and MDD) that the primary analysis run did not append. The disorder-specific variant appends it to the base developer prompt and was run as a pre-specified sensitivity test.

##### How the two prompts differ

The shared prompt differs from the disorder-specific variant in three ways.

- It names major depressive disorder and lists the nine MDD criterion domains, where the variant names the disorder of the run and appends its addendum.
- It adds a rule for the three MDD criteria that name two directions. Under that rule an opposite-direction mouse phenotype counts as support for the sister target on the same axis and not as a mismatch.
- It omits a sentence of the base prompt stating that disorder-specific criteria are appended.

##### What is common to both

The user prompt, the response schema and its enumerations, candidate retrieval, the routing of pairs to the LLM, the score adjustment and every deterministic step are identical under both prompts. The user prompt carries no criterion text, so the developer prompt is the only text naming a disorder.

The one difference that could reach the direction logic was tested directly. Six verified pairs of increased and decreased MP sibling terms were checked for admission of both directions against one anchor in each pair primary table. Both directions were admitted once in all primary analysis runs: against the SCZ anchor aggressive behavior (HP:0000718), both increased aggression (MP:0005655) and decreased aggression (MP:0005656). No ASD, IDD, NCD or MDD table holds such a case. The SCZ pair is retained in the record and disclosed.

##### Why the shared prompt is the primary run

Every validation sample (the expert pairs, the alternate-LLM re-judgments and the admission control) was drawn from the primary verdicts, and the catalogue is built from them. The disorder-specific variant has not been shown to be more correct. The primary verdicts therefore remain the record, and the variant is reported as a pre-specified sensitivity test that bounds the effect of the prompt.

##### The same-LLM sensitivity rerun

Every pair with a primary verdict in the four primary disorders was re-judged with the same LLM under the disorder-specific prompts in a run completed on 2026-10-28 (pre-specified as amendment 3 of the LLM revalidation, outcome 1). Bundle pairs received the base developer prompt with the disorder’s addendum, and the remaining pairs received the base prompt. Every call was served by gpt-5.4-2026-03-05 (1,419, 1,394, 890 and 533 calls for ASD, SCZ, IDD and NCD). Aggregation and the permutation null were rerun with the primary configurations. The outcome is the paired difference in support AUROC, disorder-specific prompt minus record, with a gene-clustered 90% interval read against the margin of 0.05.

*S8 Table.1. Same-LLM rerun under the disorder-specific prompts. Admitted pairs are counted over all judged anchors, as in the column of all admitted pairs in Table 1 of the main text. Contrasts use the primary labels. Turnover is the share of the record’s admissions that the rerun refuses.*

| Quantity | ASD | SCZ | IDD | NCD |
| --- | --- | --- | --- | --- |
| Admitted pairs, record | 145 | 196 | 138 | 95 |
| Admitted pairs, disorder-specific prompt | 125 | 176 | 137 | 108 |
| Record admissions refused by the rerun | 47 | 51 | 36 | 18 |
| New admissions in the rerun | 27 | 31 | 35 | 31 |
| Turnover, percent of the record’s admissions | 32.4 | 26.0 | 26.1 | 18.9 |
| Support AUROC difference, disorder-specific minus record (90% interval) | -0.029 (-0.055 to +0.000) | -0.021 (-0.070 to +0.023) | +0.045 (+0.013 to +0.068) | -0.016 (-0.036 to +0.003) |
| Reading against the margin of 0.05 | Inconclusive | Inconclusive | Inconclusive | Equivalent |

No disorder is not equivalent, so the prompt does not move the record beyond the margin in any disorder. The point estimates favor the record for ASD and the disorder-specific prompt for IDD. The two prompts agree on strict eligibility for 98.1 to 98.9% of record pairs (kappa 0.72 to 0.77) and on the analog grade at kappa 0.56 to 0.70. One fifth to one third of the admitted set turns over while the AUROC moves by 0.045 or less, so the AUROC cannot certify the admitted set (S1 Text). With the disorder-specific prompt held fixed in both runs, replacing gpt-5.4 by claude-opus-4-8 moves the SCZ support AUROC by +0.203 (+0.144 to +0.265), so the SCZ dependence on the LLM is not an effect of the prompt.

##### SM9. Dated record of pre-specification

The judgment floor and the channel weights were set on the ASD development run (May 2026) before any other disorder was run. The primary mapping prompt (v4, 2026-07-27) and its eight named rules (S26 Table) were fixed on 2026-07-27 and 2026-07-28, after the first ASD, IDD and NCD development results and before the four primary taxonomy runs on 2026-07-28. MDD’s mapping ran in June 2026 on an earlier prompt and carries none of the rules. No scoring parameter was changed after a disorder’s scores had been compared with its labels. Three changes were made after such comparisons and are reported as such: the IDD label rule, the removal of the two NCD labels (2026-11-02) and the rebuild of the NCD criterion B anchors on 2026-07-31. The wording that a phenotype models a feature of the criterion was adopted after a rater’s comment on the instrument, which asked whether the mouse phenotype is evidence for the criterion.

##### SM10. The mirrored permutation null

The observed support score and the permutation null of record combine admitted matches differently. The observed score clusters near-duplicate matches within each anchor before combining them. Two matches fall in one cluster when they share the MP term, share an ancestor or have a token Jaccard of term text at or above 0.65. Only the best match per cluster is kept. The null of record combined every candidate-passing pair of a random set by noisy-OR without that clustering. The two score functions therefore differ, and the mirrored null tests whether the difference matters.

##### Procedure

The mirrored null applies the observed score’s clustering to each random set before combining, exactly as aggregation does for the genotype’s own terms. Everything else is held fixed. The random term sets are the identical draws of record (same seed and same sets for every annotation count), the observed scores are the record’s and the p value formula is unchanged. Every difference between the two nulls therefore comes from the score function alone. The comparison is the paired difference in support AUROC with a gene-clustered 90% interval, in the evaluation set and in the full background (permutation_null_mirrored.py, audit/nullmirror_2026-10-27/RESULT_nullmirror.md). It was run under the labels of 2026-10-08.

##### Result

Mirroring the clustering leaves every support AUROC unchanged to four decimals in both genotype sets, and every paired difference and its interval is zero to four decimals. The p value changes for 10, 2, 11 and 15 genotypes (ASD, SCZ, IDD and NCD) among the 18,439, 19,432, 18,716 and 17,472 genotypes the permutation step scored. All changes go in one direction (recorded as up for every changed genotype and down for none), and the largest change in the negative log10 p is 0.013 (IDD). The mean of the null at each annotation count shifts by at most 2.4 × 10⁻⁵. The number of genotypes at the smallest attainable p is identical under the two nulls (389, 57, 478 and 17 among the genotypes the permutation step scored). The record of 2026-10-28 reads the p primary values as conservative by these amounts, and no reported number moves.

*S9 Table.1. Support AUROC under the null of record and the mirrored null (labels of 2026-10-08).*

| Disorder | Evaluation set, record null | Evaluation set, mirrored null | Full background, record null | Full background, mirrored null |
| --- | --- | --- | --- | --- |
| ASD | 0.9557 | 0.9557 | 0.9689 | 0.9689 |
| SCZ | 0.6823 | 0.6823 | 0.7094 | 0.7094 |
| IDD | 0.7511 | 0.7511 | 0.7591 | 0.7591 |
| NCD | 0.6638 | 0.6638 | 0.7237 | 0.7237 |

The NCD values in the table use the 73 NCD positives of 2026-10-08. Under the primary labels (71 positives) the NCD primary value is 0.668 (Table 3 of the main text), and the comparison was not rerun under those labels. The two nulls assign the same p value to every NCD genotype but 15, and those 15 move by at most 0.0036 in negative log10 p.

##### SM11. PhenoDigm configurations and the published-table cutoff

##### The per-pair score

PhenoDigm scores a human term h against a mouse term m by the geometric mean of two ontology similarities read from the integrated phenotype ontology (Smedley et al. 2013):

s(h, m) = sqrt( simJ(h, m) × IC(MICA(h, m)) )

Here simJ is the Jaccard similarity of the is_a ancestor closures of the two terms, and IC(MICA) is the information content of their most informative common ancestor. The value is read from the phenodigm_score column of a Monarch semantic similarity table. A row of the published table, for example, has IC 1.4432 and Jaccard 0.16216, and their product’s square root, 0.48377, is the reported score. A table can hold several rows for one pair (one per common ancestor), and the comparator keeps the maximum.

##### The genotype score

For a genotype with MP term set M and a disorder with human term set H, the score is the symmetric best-match average:

BMA(H, M) = 0.5 × [ (1/|H|) Σ over h in H of max over m in M of s(h, m) + (1/|M|) Σ over m in M of max over h in H of s(h, m) ]

A human term with no scored pair to the genotype’s terms, and a genotype term with no scored pair to any human term, contributes 0. This is the average-score component of published PhenoDigm. The maximum-score component, the scaling of each score as a percentage of the hypothetical optimal model and the combined percentage score are omitted. The scaling does not change ranks within a disorder. The human term set is the anchor set FACET resolves for the disorder, because HPO attaches no symptom profile to these diagnoses. The comparison therefore isolates admission and aggregation from the choice of human vocabulary. PhenoDigm scores are raw similarities and were not referred to FACET’s permutation null.

The anchor sets of the comparator primary files hold 46 ASD, 41 SCZ, 25 IDD and 28 NCD terms (the n_hpo_targets field of every score file, from each disorder’s resolved anchor table). These are all resolved anchors, not only those attributable to a scored criterion unit.

##### The four configurations

*S10 Table.1. PhenoDigm configurations. All are compared with FACET on the support AUROC only, in the evaluation set, with the same genotypes and gene-clustered intervals.*

| Configuration | Similarity source | Genotype profile | Role |
| --- | --- | --- | --- |
| Published table | Monarch semantic similarity table version 0.4 (2024-08-14), which retains only pairs with Jaccard 0.4 or more, so every pair below the cutoff reads as 0 | All MP terms | Comparison as distributed |
| Recomputed | The same method recomputed on the 2026 Monarch integrated ontology (phenio-monarch.db, April 2026) with the configuration of the Monarch similarity profiles, without the cutoff | All MP terms | Primary comparison (Table 3 of the main text) |
| Behavioral | As recomputed | Only terms among the 548 is_a descendants of MP:0005386 (behavior/neurological phenotype) | Specified first and kept in the Holm family |
| Admitted pairs | As recomputed | Only FACET’s admitted pairs | Isolates the scorer on identical pairs |

Over a sample of 200,000 rows of the published table, the smallest observed Jaccard is 0.4019, so no row falls below the cutoff. An absent pair in that table means a pair below the publication cutoff and not a pair without similarity. A diagnostic configuration recomputed on the 2026 ontology with the cutoff kept separates release drift from the effect of the cutoff and is not reported as a comparison.

##### Results

*S10 Table.2. FACET minus PhenoDigm, support AUROC difference in the evaluation set (95% gene-clustered interval, two-sided bootstrap p where recorded), primary labels.*

| Configuration | ASD | SCZ | IDD | NCD |
| --- | --- | --- | --- | --- |
| Published table, PhenoDigm AUROC | 0.656 | 0.725 | 0.616 | 0.557 |
| Published table, difference | +0.300 (0.226 to 0.382) | -0.043 (-0.143 to 0.046, p 0.34) | +0.135 (0.068 to 0.240) | +0.111 (0.013 to 0.203, p 0.026) |
| Recomputed, difference | +0.184 (0.126 to 0.248, p below 0.001) | -0.090 (-0.191 to 0.001, p 0.054) | +0.045 (-0.028 to 0.158, p 0.28) | +0.075 (0.007 to 0.150, p 0.031) |
| Behavioral, difference | +0.269 (0.236 to 0.301), Holm p 0.002 | +0.043 (-0.053 to 0.135, p 0.36) | +0.065 (-0.012 to 0.171, p 0.10) | +0.045 (-0.028 to 0.126, p 0.22) |
| Admitted pairs, difference | +0.033 (0.025 to 0.041) | -0.062 (-0.144 to 0.016, p 0.10) | +0.019 (0.003 to 0.029, p 0.024) | 0.000 (-0.025 to 0.023, p 0.98) |

Removing the cutoff raises PhenoDigm in every disorder. The Holm family holds the behavioral configuration, and only its ASD difference survives correction. On FACET’s admitted pairs the two scorers differ by at most 0.062.

##### SM12. Prompts of the primary analysis run

This section reproduces, for each LLM step of Table 4 of the main text, the developer prompt, the user prompt and the response schema of the primary analysis run. The texts are records of what ran and are reproduced with their original wording, spelling, capitals and punctuation. They are not prose of this paper. The prompts carry the criteria in the project’s paraphrase, which is deposited with the code release, and no DSM-5-TR text. Short criterion labels that this paper also uses are kept. Placeholders of the form <…> mark values filled per call. In the mouse criterion mapping prompts of SCZ and NCD, the blocks that define a unit’s terms are also shown as a placeholder. Their text is in the criterion specifications of the code release and is available from the authors on request.

Two provenance levels apply. The mapping prompts are reproduced from the archived prompt files of the primary taxonomy runs, whose SHA-256 digests equal the prompt hashes recorded by those runs. No archived prompt file exists for criterion decomposition, anchor resolution, anchor attribution or the pair judgment. Those texts are reproduced from the code of record in the code release. The pair-judgment prompt that ran is in a file last modified on 2026-05-16, before every primary pair-judgment run.

##### 11.1 Criterion decomposition

The developer prompt of the configuration-driven anchoring script (stage1_curate_hpo.py) is reproduced below. The disorder’s name replaces {disorder_name}, and the disorder’s addendum (S11 Table.1) replaces {addendum}. The user prompt carries the disorder’s paraphrased core text (the criteria and supporting passages; the criteria are deposited with the code release and the supporting passages are not redistributed). Calls use temperature 0 and seed 1.

Developer prompt:

```
You are extracting machine-readable phenotype statements from DSM-style text
about {disorder_name}.

Goal:
Convert the input into a compact list of phenotype statements that can later
be matched to HPO. Each statement should be a granular feature, not a
diagnostic label.

Rules:
1. Extract phenotype FEATURES, not diagnoses or DSM labels.
2. Keep short normalized phrases (e.g., "anhedonia", "insomnia",
   "reduced eye contact", "sensory hyperreactivity").
3. Do NOT output disorder names, prevalence statements, risk factors,
   treatments, coding instructions, or comparisons to other disorders.
4. Do NOT force one input sentence into one output item. Split into
   granular features.
5. Keep negated/context-only material as polarity = "negated" or
   centrality = "context_only".
6. If a statement is about developmental timing, keep it as a phenotype-
   like item under whatever developmental_course-like category is in the
   schema.
7. Prefer descriptive, ontology-friendly phrasing over DSM prose.
8. Be conservative. If a phrase is not really a phenotype, exclude it.

{addendum}
```

User prompt:

Extract phenotype statements from the following {disorder_name} core text.

TEXT:
<the disorder's paraphrased core text>

The ASD variant of the original anchoring script (pilot_asd_to_hpo_hybrid_rev.py, function call_openai_extract) has the same eight rules with ASD-specific wording and examples. Its developer prompt is reproduced for completeness.

```
You are extracting machine-readable phenotype statements from DSM-style text.

Goal:
Convert the input into a compact list of phenotype statements that can later be matched to HPO.

Rules:
1. Extract phenotype FEATURES, not diagnoses.
2. Keep short normalized phrases like:
   <short example phrases, one per line>
3. Do NOT output disorder names, prevalence statements, risk factors, treatments,
   coding instructions, or comparisons to other disorders.
4. Do NOT force one input sentence into one output item. Split into granular features.
5. Keep negated/context-only material as polarity=negated or centrality=context_only.
6. If a statement is about developmental timing (e.g. early developmental onset,
   developmental regression), keep it as a phenotype-like item under developmental_course.
7. Prefer descriptive, ontology-friendly phrasing over DSM prose.
8. Be conservative. If a phrase is not really a phenotype, exclude it.
```

*S11 Table.1. Per-disorder category lists of criterion decomposition (configuration field stage1_hpo_curation). The addenda follow the table.*

| Disorder | Category enumeration |
| --- | --- |
| ASD | social_communication, restricted_repetitive_behavior, sensory, language, developmental_course, motor, other |
| SCZ | delusions, hallucinations, disorganized_speech, catatonic_disorganized_behavior, negative_symptoms, other |
| IDD | intellectual_functioning, adaptive_functioning, developmental_onset, other |
| NCD | cognitive_executive, motor_coordination, social_cognition_empathy, adaptive_functioning, other |
| MDD | depressed_mood, anhedonia, weight_appetite_change, sleep_disturbance, psychomotor_change, fatigue, worthlessness_guilt, concentration_indecisiveness, suicidality, other |

Per-disorder addenda, as they ran. The em dash character of the IDD addendum is shown as [U+2014].

ASD addendum:

Extract phenotype FEATURES, not diagnoses. Keep short normalized phrases. Capture sensory hyper-/hypo-reactivity per DSM-5-TR Criterion B.4 (added in 2013).

SCZ addendum:

DSM-5-TR Schizophrenia Criterion A is polythetic (2 or more of five distinct symptoms; at least one from A.1, A.2, A.3). Route each to its OWN domain: A.1 delusions -> delusions; A.2 hallucinations -> hallucinations; A.3 disorganized speech / formal thought disorder -> disorganized_speech; A.4 behavior that is grossly disorganized or catatonic -> catatonic_disorganized_behavior (bidirectional: immobility AND excessive purposeless activity, emit two items); A.5 negative symptoms -> negative_symptoms (emit diminished emotional expression and avolition as separate items). Do NOT map any symptom to a salience or PPI substrate: the aberrant-salience endophenotype is the mechanism axis, populated on the mouse side, NOT a face-valid analog of the phenomenology. Do NOT emit cognitive-executive or generic-social items as SCZ criteria (those are DSM associated features, not Criterion A) -> other. Extract phenotype FEATURES as short normalized phrases. Delusions, hallucinations, and disorganized speech have no direct mouse analog; that unmodellability is the result and is captured by those domains scoring near zero.

IDD addendum:

DSM-5-TR Intellectual Disability (Intellectual Developmental Disorder) requires ALL THREE criteria: A. Shortfalls in intellectual abilities (for example reasoning, solving problems, planning, abstract thought, judgment, school learning, learning from experience) -> domain intellectual_functioning. B. Deficits in adaptive functioning across conceptual, social, and/or practical domains (communication, social participation, independent living, self-care, daily-living skills) -> domain adaptive_functioning (a single umbrella domain; do NOT split conceptual/social/practical into separate domains, tag them all as adaptive_functioning). C. Shortfalls beginning during development -> domain developmental_onset (a temporal criterion; tag onset/milestone/developmental-period features here, temporality='developmental'). There are NO bidirectional domains for ID (all domains are deficit/loss-direction). Severity specifiers (mild/moderate/severe/profound) GRADE the same A/B deficits [U+2014] extract the described deficits into intellectual_functioning or adaptive_functioning, not as separate items. Extract phenotype FEATURES, not diagnoses or IQ cutoffs. Do NOT extract from Differential Diagnosis exclusions or from the ICD-11 terminology Note. Mouse models cannot directly have 'academic skills' [U+2014] operationalized downstream via learning/memory and adaptive-behavior paradigms [U+2014] but extraction here stays at the human-symptom level.

NCD addendum:

DSM-5-TR Major/Mild Neurocognitive Disorder (Chapter 16) is defined by ACQUIRED cognitive decline from a previous baseline in one or more of six domains. Map extracted human symptoms to these strict domains: (1) cognitive_executive - complex attention, executive function (planning, working memory, set-shifting, response inhibition, mental flexibility), and learning and memory (free recall, recognition, recent memory). Consolidates the attention/executive/memory poles into one domain. (2) motor_coordination - perceptual-motor function: coordination, gait, praxis, visuospatial navigation, and etiology-specific motor signs (parkinsonism in Lewy body / Parkinson disease, chorea and motor disorganization in Huntington disease, gait disorder in vascular / TBI). (3) social_cognition_empathy - recognition of others' emotions, theory of mind, empathy, social judgment; loss of empathy and social disinhibition are the behavioral-variant frontotemporal signature. (4) adaptive_functioning - Criterion B: interference with independence in everyday activities (instrumental activities of daily living). (5) neuropathology_progressive - the etiology-defining PROGRESSIVE neuropathology named by the subtype: amyloid and tau (Alzheimer), TDP-43 / tau (frontotemporal), alpha-synuclein (Lewy body / Parkinson), white matter / vascular injury, CAG-expansion degeneration (Huntington), spongiform change (prion). Tag the LANGUAGE domain (word-finding, naming, fluency, grammar) as 'other': language is a chapter-16 neurocognitive domain but has NO organism-level analog in the mouse, so it is not a strict NCD domain here. Extract phenotype FEATURES at the human-symptom level, not diagnoses. Do NOT extract from Differential Diagnosis or the delirium exclusion. The disorder-defining feature is PROGRESSIVE ACQUIRED decline; where the text marks progression or age-relatedness, set temporality accordingly. [2026-07-27 CRITERION-STRICT UPDATE: neuropathology_progressive is REMOVED as a face-validity domain; it is the etiology/pathology (mechanism axis), not a DSM Criterion A cognitive-decline criterion. Do NOT extract amyloid/tau/synuclein/pathology terms as scored NCD criteria -> other.]

MDD addendum:

Three MDD Criterion A symptoms are intrinsically BIDIRECTIONAL per DSM-5-TR: A.3 weight_appetite_change (gain OR loss), A.4 sleep_disturbance (insomnia OR hypersomnia), A.5 psychomotor_change (agitation OR retardation). When the DSM text gives both directions in one criterion, emit TWO separate items (one per direction) and set polarity = 'bidirectional' on each. Do NOT extract from 'Note:' blocks unless the Note directly qualifies a specific Criterion item (the clause of item A.1 on irritable mood in children and adolescents is part of that item). The A.1 clause on irritable mood in children and adolescents must use temporality='developmental'.

Response schema (ASD form, from the original script). The configuration-driven script builds the same schema with the disorder’s category list, and for every disorder except ASD the polarity list adds bidirectional.

```
{
 "name": "phenotype_extraction",
 "strict": true,
 "schema": {
  "type": "object",
  "additionalProperties": false,
  "properties": {
   "disorder_name": {
    "type": "string"
   },
   "items": {
    "type": "array",
    "items": {
     "type": "object",
     "additionalProperties": false,
     "properties": {
      "source_fragment": {
       "type": "string"
      },
      "normalized_phenotype": {
       "type": "string"
      },
      "category": {
       "type": "string",
       "enum": [
        "social_communication",
        "restricted_repetitive_behavior",
        "sensory",
        "language",
        "developmental_course",
        "motor",
        "other"
       ]
      },
      "centrality": {
       "type": "string",
       "enum": [
        "core",
        "supporting",
        "context_only"
       ]
      },
      "polarity": {
       "type": "string",
       "enum": [
        "present",
        "absent",
        "negated",
        "uncertain"
       ]
      },
      "temporality": {
       "type": "string",
       "enum": [
        "current",
        "history",
        "developmental",
        "unspecified"
       ]
      },
      "notes": {
       "type": "string"
      }
     },
     "required": [
      "source_fragment",
      "normalized_phenotype",
      "category",
      "centrality",
      "polarity",
      "temporality",
      "notes"
     ]
    }
   }
  },
  "required": [
   "disorder_name",
   "items"
  ]
 }
}
```

##### 11.2 Anchor resolution

The resolution call receives each phrase with its shortlist of at most 12 HPO terms (section SM1). Calls use temperature 0 and seed 1. No resolution-prompt addendum is configured for any disorder.

Developer prompt:

You adjudicate mappings from DSM-derived phenotype phrases to a bounded shortlist of HPO candidates.

You MUST choose from the provided shortlist only. Do not invent HPO IDs.
It is acceptable to abstain and return selected_hpo_id = NONE.

Decision guidance:
- Prefer semantically faithful phenotype matches over lexical similarity.
- If the candidate matches the source well, use mapping_relation = exact.
- If the candidate is broader than the source phrase, use broader_than_source.
- If the candidate is narrower/more specific than the source phrase, use narrower_than_source.
- If the candidate is only an approximate behavioral or conceptual proxy, use proxy_related.
- If no candidate is good enough, use no_match and selected_hpo_id = NONE.
- Use skip_not_phenotype only when the extracted item should not be mapped as a phenotype at all.
- Be conservative for psychiatry language and avoid forced mappings.

Confidence must be between 0 and 1.
Rationale should be concise and mention why the chosen candidate beats the top alternatives, or why abstention is warranted.

User prompt, a JSON object of this form (the source fragment is paraphrased criterion text):

```
{
  "phenotype_item": {
    "source_fragment": "<fragment of the paraphrased core text>",
    "normalized_phenotype": "<phrase returned by decomposition>",
    "category": "<category>",
    "centrality": "<core | supporting | context_only>",
    "polarity": "<polarity>",
    "temporality": "<temporality>",
    "notes": "<notes returned by decomposition>"
  },
  "candidate_hpo_terms": [
    {
      "candidate_rank": 1,
      "hpo_id": "HP:...",
      "label": "...",
      "synonyms": ["..."],
      "definition": "...",
      "retrieval": {"from_fuzzy": true, "from_semantic": true, "fuzzy_score": 0.0,
                    "semantic_score": 0.0, "combined_score": 0.0,
                    "matched_alias": "...", "matched_alias_type": "label | synonym"}
    }
  ]
}
```

Response schema. The enumeration of selected_hpo_id is rebuilt for every call from the shortlist, so no identifier outside it can be returned. A code check sets the outcome to unresolved if an identifier outside the shortlist is ever returned.

```
{
 "name": "hpo_resolution",
 "strict": true,
 "schema": {
  "type": "object",
  "additionalProperties": false,
  "properties": {
   "decision_status": {
    "type": "string",
    "enum": [
     "resolved",
     "unresolved",
     "skip_not_phenotype"
    ]
   },
   "selected_hpo_id": {
    "type": "string",
    "enum": [
     "<the shortlisted HPO identifiers>",
     "NONE"
    ]
   },
   "mapping_relation": {
    "type": "string",
    "enum": [
     "exact",
     "broader_than_source",
     "narrower_than_source",
     "proxy_related",
     "no_match"
    ]
   },
   "confidence": {
    "type": "number"
   },
   "manual_review": {
    "type": "boolean"
   },
   "rationale": {
    "type": "string"
   }
  },
  "required": [
   "decision_status",
   "selected_hpo_id",
   "mapping_relation",
   "confidence",
   "manual_review",
   "rationale"
  ]
 }
}
```

##### 11.3 Anchor attribution

The attribution call (decompose_targets_to_criterion_units.py) attributes each anchor to one criterion unit or marks it unattributable. It renders the same unit blocks as the mapping prompt, so those blocks follow 11.5. The call sets no temperature or seed (provider default sampling). Its schema leaves item_id as free text, and an answer outside the disorder’s units aborts the run. {spec.get(…)} is replaced by the disorder’s full name.

Developer prompt:

```
You attribute a HUMAN phenotype to the DSM-5-TR diagnostic unit it instantiates, for
{spec.get('disorder_long', spec['disorder'])}.

This is the mirror of the mouse-side task. There the question was which unit an MP term instantiates.
Here the phenotype is a human one, resolved from the DSM text itself into a Human Phenotype Ontology
term, so attribution should usually be possible and clean. The units below are paraphrased from
DSM-5-TR and are the same strings used to judge the mouse vocabulary.

[unit blocks, rendered as in the mapping prompt of 11.5]

HOW TO DECIDE
  - Choose the ONE unit the phenotype most directly instantiates and put it in `item_id`.
  - If it genuinely instantiates a second unit as well, name that in `second_item_id`, else "none".
  - A phenotype that belongs to this disorder but to NO scored unit, for example an associated
    feature, an onset statement, a severity specifier or an exclusion, is NOT attributable: set
    item_id to "none" and attributable to false. Do not force it into the nearest unit.
  - Judge against the unit's own text, not against the disorder in general. A phenotype
    about the disorder as a whole, with no unit that names it, is "none".
  - Set needs_manual_review when two units are genuinely close and the choice would change a count.

Valid `item_id` values: {", ".join(units)}, or "none".
Return JSON only.
```

User prompt:

HPO term: <anchor label>
HPO id: <anchor identifier>
Normalized phenotype as extracted from the DSM text: <phrase returned by decomposition>
Centrality recorded at stage 1: <centrality>
Rationale recorded at stage 1: <first 400 characters of the resolution rationale>

Response schema:

```
{
 "name": "target_unit_attribution",
 "schema": {
  "type": "object",
  "additionalProperties": false,
  "properties": {
   "item_id": {
    "type": "string"
   },
   "second_item_id": {
    "type": "string"
   },
   "attributable": {
    "type": "boolean"
   },
   "confidence": {
    "type": "number"
   },
   "needs_manual_review": {
    "type": "boolean"
   },
   "reason": {
    "type": "string"
   }
  },
  "required": [
   "item_id",
   "second_item_id",
   "attributable",
   "confidence",
   "needs_manual_review",
   "reason"
  ]
 },
 "strict": true
}
```

##### 11.4 Mouse domain labeling

Mouse domain labeling (gpt-5.4-mini, 15 MP terms per call) ran from per-disorder taxonomy scripts written before the criterion-mapped design (for example rebuild_behavioral_taxonomy.py and rebuild_scz_taxonomy_5domain.py in the code release). Its domain label is shown to the pair judgment as mouse_domain, and its tiers were replaced by the mapping step. Its prompts are not reproduced in this section, and the release files hold them.

##### 11.5 Mouse criterion mapping

The archived developer prompt of the ASD primary taxonomy run (taxonomy_v2_full/run_ASD/00_developer_prompt.txt, SHA-256 beginning 679539aea9a31e99) is reproduced in full below. Each named rule is replaced by a pointer to section SM2. Calls use temperature 0 and seed 1, one MP term per call.

You are classifying a single Mammalian Phenotype (MP) ontology term for a ASD mouse-model
face-validity pipeline. ASD is Autism Spectrum Disorder.

YOUR TASK
Decide which DSM-5-TR diagnostic item, if any, this mouse phenotype term could serve as a STRICT analog of,
meaning a defensible mouse instantiation of what that unit describes.

THIS DISORDER'S ENUMERATION IS COUNTED. DSM-5-TR requires ALL THREE Criterion A items and AT LEAST TWO of the four Criterion B items. Criterion A and Criterion B are each enumerated with a counting threshold, so each numbered item is a separately scored unit. So each numbered item is a separate unit, and a term serves the item it instantiates, not the criterion in general. There are 7 units.

THE UNITS, PARAPHRASED FROM DSM-5-TR. These are the ONLY definitions. Do not substitute your own summary of what
this disorder is, and do not apply criteria the text does not state. The parenthetical examples are part
of the text and are AUTHORITATIVE: a mouse term matching a named example matches the unit.

UNIT A1
 Criterion A stem: Lasting difficulty with social communication and social interaction that shows in more than one setting, now or in the past, through all three items below (the examples illustrate each item and do not exhaust it):
 Text (PARAPHRASED FROM DSM-5-TR): Trouble with the give and take of social and emotional exchange. Examples run from odd ways of approaching others and conversation that does not go back and forth; through sharing interests, feelings or emotional states less than others do; to not starting social contact or not answering it.

UNIT A2
 Criterion A stem: Lasting difficulty with social communication and social interaction that shows in more than one setting, now or in the past, through all three items below (the examples illustrate each item and do not exhaust it):
 Text (PARAPHRASED FROM DSM-5-TR): Trouble with the nonverbal behavior people use to communicate socially. Examples run from speech and nonverbal signals that fit poorly together; through unusual eye contact or body posture, or difficulty reading and making gestures; to no facial expression and no nonverbal communication at all.

UNIT A3
 Criterion A stem: Lasting difficulty with social communication and social interaction that shows in more than one setting, now or in the past, through all three items below (the examples illustrate each item and do not exhaust it):
 Text (PARAPHRASED FROM DSM-5-TR): Trouble building, keeping and making sense of relationships. Examples run from difficulty fitting one's behavior to different social situations; through difficulty joining others in pretend play or forming friendships; to showing no interest in peers.

UNIT B1
 Criterion B stem: Narrow, repeated patterns in behavior, interests or activities, now or in the past, through at least two items below (the examples illustrate each item and do not exhaust it):
 Text (PARAPHRASED FROM DSM-5-TR): Movements, handling of objects or speech that are stereotyped or repeated (for example simple repeated motor acts, arranging toys in rows or turning objects over and over, repeating the words of others, or phrases with a private meaning).

UNIT B2
 Criterion B stem: Narrow, repeated patterns in behavior, interests or activities, now or in the past, through at least two items below (the examples illustrate each item and do not exhaust it):
 Text (PARAPHRASED FROM DSM-5-TR): A need for things to stay the same, routines kept without flexibility, or ritual patterns of speech or nonverbal behavior (for example great upset at minor changes, trouble moving from one activity to the next, rigid ways of thinking, fixed greeting rituals, or needing the same route or the same food every day).

UNIT B3
 Criterion B stem: Narrow, repeated patterns in behavior, interests or activities, now or in the past, through at least two items below (the examples illustrate each item and do not exhaust it):
 Text (PARAPHRASED FROM DSM-5-TR): Interests that are very narrow and fixed and unusual in how strong or how focused they are (for example intense attachment to, or absorption in, unusual objects, or interests that are overly confined or perseverative).

UNIT B4
 Criterion B stem: Narrow, repeated patterns in behavior, interests or activities, now or in the past, through at least two items below (the examples illustrate each item and do not exhaust it):
 Text (PARAPHRASED FROM DSM-5-TR): Over- or under-response to sensory stimulation, or an unusual interest in the sensory side of the surroundings (for example seeming not to notice pain, heat or cold, strong aversion to particular sounds or textures, smelling or touching objects to excess, or fascination with watching lights or movement).

BLINDING, WHICH STILL APPLIES
Classify ONLY from the MP term label, definition, parent and ancestor text. Do NOT use or infer genotype,
gene identity, disease-gene status, disease association, literature prominence, or any ranking result.
This is a blinded ontology-term task. The blinding covers evidence about MODELS. It does NOT cover the
diagnostic criteria above, which you are required to use.

VOCABULARY REALISM, AND THIS IS THE MOST IMPORTANT INSTRUCTION
MP labels are terse and are written for mouse phenotyping, not for psychiatry. They almost never carry the
qualifiers the DSM text uses. MP has no way to say "rigid", "fixated", "excessive", "unusual in strength
or focus", "a behavioral response rather than a measured threshold", or "worse than this animal used to
be".

DO NOT REFUSE A TERM FOR LACKING A QUALIFIER THE VOCABULARY CANNOT EXPRESS. If the term names the right
construct in the right modality for the unit, admit it and set qualifier_absent=true. Recording the
uncertainty is correct; refusing every real term because none carries an impossible qualifier is not.

Worked application of the vocabulary realism rule:
 - The B4 text names seeming not to notice pain, heat or cold. MP expresses pain and temperature response only as nociception and thresholds (abnormal thermal nociception, abnormal pain threshold). Those ARE the mouse instantiation of that example. Admit them to B4 with qualifier_absent=true. Do NOT refuse them as "capacity rather than reactivity"; MP offers no reactivity-framed alternative.
 - The B4 text names smelling or touching objects to excess. Object-directed exploration terms (abnormal response to novel object, abnormal habituation to a novel object) ARE that instantiation. Admit.
 - The B1 text names the handling of objects. Object manipulation and use terms serve B1.
 - The B2 text names ritual patterns and the B3 text names narrow, fixed interests. Grooming, nest building and stereotypy terms may serve these regardless of whether the label says "rigid", since MP cannot say it.

WHERE TO STILL SAY NO. The rule above widens admission for VOCABULARY reasons only. It does not widen it
for CONSTRUCT reasons. Refuse when:
 - The term is anatomical, morphological, histological or molecular (set modality accordingly).
 - The term is a mechanistic or circuit-level measure (synaptic transmission, EEG, receptor currents,
 neuronal morphology). These support mechanism, not face validity.
 - The term is an ontology umbrella node so broad it names no specific phenotype ("abnormal behavior").
 - No unit fits. Then item_id is "none" and admissible is false. This is a normal and expected outcome
 for the large majority of MP terms.

[Named rule grooming_routing, reproduced in section SM2]

[Named rule involuntary_motor_sign, reproduced in section SM2]

[Named rule asd_construct_refusals, reproduced in section SM2]

COMPONENT
This disorder's units enumerate no named components, so set component to not_applicable.

DIRECTION
Report in term_direction the direction the MP LABEL ITSELF states for the underlying function:
 increase	the label states more of it ("increased grooming behavior", "excessive vocalization",
 "enhanced social recognition", "hyperresponsive to tactile stimuli")
 decrease	the label states less of it ("decreased vocalization", "impaired social recognition",
 "hyporesponsive to tactile stimuli", "reduced ...")
 unspecified	the label says only "abnormal ..." and does not state which way
 not_directional the term names no quantity that could go either way
Report what the LABEL SAYS. Do NOT judge whether the direction resembles the disorder; that comparison is
made downstream from the DSM text and is not your call. A term whose direction is opposite to the unit is
still correctly classified here, and will be handled later.

CONSTRUCT INADMISSIBILITY
Set construct_inadmissible=true only when the DSM unit names something NO mouse can instantiate under any
vocabulary, for example spoken idiosyncratic phrases, delusional belief content, or standardized
intelligence testing. This is a statement about the UNIT, not about this term, and it lets the pipeline
report the unit as unassessable rather than as a measured zero. Those are different claims: a measured
zero says the models were tested and failed, an unassessable says the question cannot be put to a mouse.

OUTPUT
Assign the single best unit to item_id. If a term genuinely serves two, put the second in second_item_id,
otherwise "none". Set needs_manual_review=true whenever the decision is close, and write one or two
sentences in reason naming which part of the unit text you relied on.

The prompts of the other three disorders were compared with the ASD prompt. From the heading BLINDING, WHICH STILL APPLIES to the end, they are identical to the ASD prompt except in three places. They have no worked examples of the vocabulary realism rule, they carry their own named rules (S2 Table.1) and they carry their own COMPONENT paragraph. Their opening sections and COMPONENT paragraphs are reproduced below. The SCZ notes on A1 to A3 are the project’s own wording and are kept.

SCZ (prod_SCZ_2026-07-28, SHA-256 beginning c6853a6bd0a58fda), opening section:

You are classifying a single Mammalian Phenotype (MP) ontology term for a SCZ mouse-model
face-validity pipeline. SCZ is Schizophrenia.

YOUR TASK
Decide which DSM-5-TR diagnostic unit, if any, this mouse phenotype term could serve as a STRICT analog of,
meaning a defensible mouse instantiation of what that unit describes.

THIS DISORDER'S UNITS ARE OF TWO KINDS, because DSM enumerates it two ways in the same criteria set. DSM-5-TR Criterion A requires TWO OR MORE of five numbered items, and at least one must be item 1, 2 or 3. That is a counting threshold with an additional gate, so each numbered item is a separately scored unit. Criterion B is a separate requirement of the diagnosis and carries no counting threshold: it lists major life areas as examples and asks for at least one, so it is ILLUSTRATIVE and is satisfied by marked dysfunction in any ONE named area. A diagnosis requires Criterion A AND Criterion B. There are 6 units in total. 5 are COUNTED items (A1, A2, A3, A4, A5): DSM counts them, so each is a separate unit and a term serves the item it instantiates rather than the criterion in general. 1 is a CRITERION-level unit (B): its own enumeration is ILLUSTRATIVE and is satisfied by any ONE named area, so do not hold a term to the whole criterion when it cleanly instantiates one named area of it.

THE UNITS, PARAPHRASED FROM DSM-5-TR. These are the ONLY definitions. Do not substitute your own summary of what
this disorder is, and do not apply criteria the text does not state. The parenthetical examples are part
of the text and are AUTHORITATIVE: a mouse term matching a named example matches the unit.

UNIT A1
 Criterion A stem: At least two of the five items below, each present for a substantial part of a one-month period (or for less time if treatment succeeded). One of the two must be delusions, hallucinations or disorganized speech:
 Text (PARAPHRASED FROM DSM-5-TR): Delusions.
 NOTE: A delusion is a belief with propositional content. No organism-level mouse analog exists under any vocabulary. construct_inadmissible.
 WHAT DSM ITSELF MEANS BY THE TERMS IN THIS UNIT. This paraphrases DSM-5-TR's definitional text from elsewhere in the same chapter, and it OVERRIDES the plain reading of a name. A term of art like `perceptual-motor` sounds broader than DSM's definition of it. Use the definition, not the name.
 <paraphrased definition of this unit's terms>

UNIT A2
 Criterion A stem: At least two of the five items below, each present for a substantial part of a one-month period (or for less time if treatment succeeded). One of the two must be delusions, hallucinations or disorganized speech:
 Text (PARAPHRASED FROM DSM-5-TR): Hallucinations.
 NOTE: A hallucination is a percept without a stimulus, reportable only by the experiencer. No mouse analog. construct_inadmissible.
 WHAT DSM ITSELF MEANS BY THE TERMS IN THIS UNIT. This paraphrases DSM-5-TR's definitional text from elsewhere in the same chapter, and it OVERRIDES the plain reading of a name. A term of art like `perceptual-motor` sounds broader than DSM's definition of it. Use the definition, not the name.
 <paraphrased definition of this unit's terms>

UNIT A3
 Criterion A stem: At least two of the five items below, each present for a substantial part of a one-month period (or for less time if treatment succeeded). One of the two must be delusions, hallucinations or disorganized speech:
 Text (PARAPHRASED FROM DSM-5-TR): Speech that is disorganized (for example often going off track, or incoherent).
 NOTE: Mice have no speech. Ultrasonic vocalization syntax is a remote non-criterion proxy and is NOT admitted as strict. construct_inadmissible.
 WHAT DSM ITSELF MEANS BY THE TERMS IN THIS UNIT. This paraphrases DSM-5-TR's definitional text from elsewhere in the same chapter, and it OVERRIDES the plain reading of a name. A term of art like `perceptual-motor` sounds broader than DSM's definition of it. Use the definition, not the name.
 <paraphrased definition of this unit's terms>

UNIT A4
 Criterion A stem: At least two of the five items below, each present for a substantial part of a one-month period (or for less time if treatment succeeded). One of the two must be delusions, hallucinations or disorganized speech:
 Text (PARAPHRASED FROM DSM-5-TR): Behavior that is grossly disorganized, or behavior that is catatonic.
 NOTE: Partially modellable and explicitly BIDIRECTIONAL: the item names grossly disorganized OR catatonic behavior, so both catalepsy, waxy flexibility, immobility and stupor at one pole, and hyperlocomotion and stereotypy at the other, are organism-level analogs.
 WHAT DSM ITSELF MEANS BY THE TERMS IN THIS UNIT. This paraphrases DSM-5-TR's definitional text from elsewhere in the same chapter, and it OVERRIDES the plain reading of a name. A term of art like `perceptual-motor` sounds broader than DSM's definition of it. Use the definition, not the name.
 <paraphrased definition of this unit's terms>

UNIT A5
 Criterion A stem: At least two of the five items below, each present for a substantial part of a one-month period (or for less time if treatment succeeded). One of the two must be delusions, hallucinations or disorganized speech:
 Text (PARAPHRASED FROM DSM-5-TR): Negative symptoms, that is, reduced expression of emotion or loss of drive (avolition).
 NOTE: Most modellable. The item's own parenthetical names diminished emotional expression and avolition. Absorbs asociality and social withdrawal.
 WHAT DSM ITSELF MEANS BY THE TERMS IN THIS UNIT. This paraphrases DSM-5-TR's definitional text from elsewhere in the same chapter, and it OVERRIDES the plain reading of a name. A term of art like `perceptual-motor` sounds broader than DSM's definition of it. Use the definition, not the name.
 <paraphrased definition of this unit's terms>

UNIT B
 Text (PARAPHRASED FROM DSM-5-TR): For a substantial part of the time since the illness began, functioning in at least one major area of life, for example work, relationships with other people or looking after oneself, is clearly below the level reached before onset (or, if the illness began in childhood or adolescence, the person never reaches the interpersonal, school or work functioning expected of them).
 NAMED COMPONENTS OF THIS CRITERION, 5, named in the same text: work; relationships with other people; looking after oneself; school functioning; occupational functioning
 These are EXAMPLES, not a list to be satisfied. The criterion is met by any ONE of them. Record which one in `component`.
 NOTE: WHAT THIS CRITERION ASKS FOR IS DETERIORATION, NOT A TRAIT DIFFERENCE. DSM asks whether functioning in a major life area is markedly below the level the individual previously achieved, or, for onset in childhood or adolescence, whether an expected level was never achieved. Both readings are about a CHANGE or a FAILURE TO ARRIVE, not about a stable characteristic. A mouse phenotype that is a constant lifelong difference from wild type is the weakest form of this criterion; one that emerges or worsens after development is the strongest. RECORD WHICH ONE YOU ARE LOOKING AT, because the project measures the size of this substitution rather than assuming it. Set qualifier_absent=false when the term itself names emergence, progression, deterioration, age-dependence or a loss of a previously present capacity. Set qualifier_absent=true when the term names only a stable between-genotype difference and is therefore standing in for a deterioration that MGI does not state. Do not default to either. THE BOUNDARY WITH A5, WHICH YOU MUST NOT COLLAPSE. A5 names the SYMPTOM, diminished emotional expression and avolition. B names the CONSEQUENCE at the level of major life areas: work, interpersonal relations, self-care, academic and occupational functioning. Many mouse terms can be read either way. Assign your best single unit in item_id, and when a term genuinely instantiates both, record the other in second_item_id rather than silently choosing one.

SCZ, COMPONENT paragraph:

COMPONENT
Only some units name components. Where a unit lists NAMED COMPONENTS above, set component to the one the term instantiates, choosing the most accurate rather than the most favourable, or none if the term serves that criterion but no named component. For every unit that lists no components, set component to not_applicable.

IDD (run_ID, SHA-256 beginning f495ff735b361000), opening section:

You are classifying a single Mammalian Phenotype (MP) ontology term for a ID mouse-model
face-validity pipeline. ID is Intellectual Developmental Disorder (Intellectual Disability).

YOUR TASK
Decide which DSM-5-TR diagnostic criterion, if any, this mouse phenotype term could serve as a STRICT analog of,
meaning a defensible mouse instantiation of what that unit describes.

THIS DISORDER'S ENUMERATION IS ILLUSTRATIVE, NOT COUNTED. DSM-5-TR requires ALL THREE criteria A, B and C to be met. There is NO counting threshold over a numbered list, so there are no numbered items to count. The lists inside A and B are given as examples and are ILLUSTRATIVE, not a menu to be satisfied in part. So the CRITERION is the unit, and the named components are examples of it. There are 2 units. SATISFYING ONE COMPONENT SATISFIES THE CRITERION: do not hold a term to the whole criterion when it cleanly instantiates one named component of it.

THE UNITS, PARAPHRASED FROM DSM-5-TR. These are the ONLY definitions. Do not substitute your own summary of what
this disorder is, and do not apply criteria the text does not state. The parenthetical examples are part
of the text and are AUTHORITATIVE: a mouse term matching a named example matches the unit.

UNIT A
 Text (PARAPHRASED FROM DSM-5-TR): Shortfalls in intellectual abilities, for example reasoning, solving problems, planning, thinking in the abstract, judgment, learning in school and learning from experience, shown both by clinical assessment and by intelligence tests that are standardized and given to the person individually.
 NAMED COMPONENTS OF THIS CRITERION, 7, named in the same text: reasoning; solving problems; planning; abstract thought; judgment; school learning; learning from experience
 These are EXAMPLES, not a list to be satisfied. The criterion is met by any ONE of them. Record which one in `component`.

UNIT B
 Text (PARAPHRASED FROM DSM-5-TR): Shortfalls in adaptive functioning, so that the person does not meet the developmental and sociocultural expectations for independence and social responsibility at their age. Without continuing support, these shortfalls restrict functioning in one or more everyday activities, for example communicating, taking part in social life and living independently, in more than one setting, for example at home, at school, at work and in the community.
 NAMED COMPONENTS OF THIS CRITERION, 3, named in the same text: communicating; taking part in social life; living independently
 These are EXAMPLES, not a list to be satisfied. The criterion is met by any ONE of them. Record which one in `component`.

IDD, COMPONENT paragraph:

COMPONENT
Set component to the named component the term instantiates. This is recorded for audit and is NEVER scored, so choose the most accurate one rather than the most favourable. Use none if the term serves the criterion but no named component.

NCD (prod_NCD_2026-07-28, SHA-256 beginning 2330f79d059f16b1), opening section:

You are classifying a single Mammalian Phenotype (MP) ontology term for a NCD mouse-model
face-validity pipeline. NCD is Major Neurocognitive Disorder.

YOUR TASK
Decide which DSM-5-TR diagnostic criterion, if any, this mouse phenotype term could serve as a STRICT analog of,
meaning a defensible mouse instantiation of what that unit describes.

THIS DISORDER'S ENUMERATION IS ILLUSTRATIVE, NOT COUNTED. DSM-5-TR requires criteria A through D. Criterion A is met by a decline in AT LEAST ONE cognitive domain, a threshold of one, which sets no counting gate over the domain list. Criterion A's numbered sub-items 1 and 2 are EVIDENCE REQUIREMENTS (informant or clinician concern; documented neuropsychological impairment), not symptom domains, and are not scoreable in a mouse under any vocabulary. So the CRITERION is the unit, and the named components are examples of it. There are 2 units. SATISFYING ONE COMPONENT SATISFIES THE CRITERION: do not hold a term to the whole criterion when it cleanly instantiates one named component of it.

THE UNITS, PARAPHRASED FROM DSM-5-TR. These are the ONLY definitions. Do not substitute your own summary of what
this disorder is, and do not apply criteria the text does not state. The parenthetical examples are part
of the text and are AUTHORITATIVE: a mouse term matching a named example matches the unit.

UNIT A
 Criterion A stem: Evidence of a substantial drop in cognitive performance compared with the person's earlier level, in at least one cognitive domain (learning and memory, executive function, complex attention, language, social cognition, or perceptual-motor function), resting on both of the following:
 Text (PARAPHRASED FROM DSM-5-TR): Evidence of a substantial drop in cognitive performance compared with the person's earlier level, in at least one cognitive domain (learning and memory, executive function, complex attention, language, social cognition, or perceptual-motor function), resting on both of the following: 1. The person, someone who knows the person well, or the clinician is concerned that cognitive function has clearly declined; and 2. Cognitive performance is substantially impaired, ideally as shown by standardized neuropsychological tests or, without such tests, by some other clinical assessment that yields numbers.
 NAMED COMPONENTS OF THIS CRITERION, 6, named in the same text: learning and memory; executive function; complex attention; language; social cognition; perceptual-motor
 These are EXAMPLES, not a list to be satisfied. The criterion is met by any ONE of them. Record which one in `component`.
 WHAT DSM ITSELF MEANS BY THE TERMS IN THIS UNIT. This paraphrases DSM-5-TR's definitional text from elsewhere in the same chapter, and it OVERRIDES the plain reading of a name. A term of art like `perceptual-motor` sounds broader than DSM's definition of it. Use the definition, not the name.
 <paraphrased definition of this unit's terms>

UNIT B
 Text (PARAPHRASED FROM DSM-5-TR): These deficits get in the way of independence in everyday life (at the least, the person needs help with demanding instrumental activities of daily living, for example handling bills or keeping track of medicines).
 NAMED COMPONENTS OF THIS CRITERION, 1, named in the same text: instrumental activities of daily living
 These are EXAMPLES, not a list to be satisfied. The criterion is met by any ONE of them. Record which one in `component`.
 NOTE: THIS CRITERION IS THE MAJOR VERSUS MILD DISCRIMINATOR, NOT A SEVERITY THRESHOLD. Major and Mild neurocognitive disorder share Criterion A apart from a substantial versus a modest decline: both require acquired cognitive decline in at least one of the six domains. What separates them is THIS criterion. For Major, the deficits GET IN THE WAY of independence in everyday life, so that at the least the person needs help with demanding instrumental activities of daily living, for example handling bills or keeping track of medicines. For Mild, those same activities are KEPT, though they may take more effort or compensating strategies. This spec is fenced to Major per register item D9, so a term admitted here is part of what makes the diagnosis Major rather than Mild. Judge it on interference with independent activity, not on the severity of the cognitive deficit itself, which belongs to Criterion A.

NCD, COMPONENT paragraph:

COMPONENT
Set component to the named component the term instantiates. This is recorded for audit and is NEVER scored, so choose the most accurate one rather than the most favourable. Use none if the term serves the criterion but no named component.

User prompt (classify_mp_terms_criterion_grounded.py, function build_user_prompt). In every primary taxonomy run the parent and ancestor fields of the input table were empty, so both fields read NA in every call.

Classify this blinded MP term.

mp_id: <MP identifier>
mp_label: <MP label>
mp_definition: <MP definition, or NA>
parent_label: <parent label, or NA>
ancestor_label: <ancestor label, or NA>

Response schema (criterion_prompt.py, function build_schema, joint mode). The unit and component enumerations are filled from the disorder’s criterion specification.

```
{
 "name": "mp_criterion_classification",
 "strict": true,
 "schema": {
  "type": "object",
  "additionalProperties": false,
  "required": [
   "item_id",
   "admissible",
   "second_item_id",
   "component",
   "modality",
   "term_direction",
   "qualifier_absent",
   "construct_inadmissible",
   "confidence",
   "needs_manual_review",
   "reason"
  ],
  "properties": {
   "item_id": {
    "type": "string",
    "enum": [
     "<the disorder's unit identifiers>",
     "none"
    ]
   },
   "second_item_id": {
    "type": "string",
    "enum": [
     "<the disorder's unit identifiers>",
     "none"
    ]
   },
   "component": {
    "type": "string",
    "enum": [
     "<the named components of the disorder's units>",
     "none",
     "not_applicable"
    ],
    "description": "Which NAMED component of that criterion the term instantiates. Reported, never scored. not_applicable when none are enumerated."
   },
   "admissible": {
    "type": "boolean"
   },
   "modality": {
    "type": "string",
    "enum": [
     "behavioral",
     "anatomical",
     "physiological",
     "developmental",
     "other"
    ]
   },
   "term_direction": {
    "type": "string",
    "enum": [
     "increase",
     "decrease",
     "unspecified",
     "not_directional"
    ]
   },
   "qualifier_absent": {
    "type": "boolean"
   },
   "construct_inadmissible": {
    "type": "boolean"
   },
   "confidence": {
    "type": "number"
   },
   "needs_manual_review": {
    "type": "boolean"
   },
   "reason": {
    "type": "string"
   }
  }
 }
}
```

##### 11.6 Pair judgment

The pair judgment ran in two passes with one prompt. Bundle pairs (the five highest candidates of an anchor, when the top two scores lie within 0.08 or the top score lies in 0.35 to 0.70) were judged together. Every other pair scoring at least 0.57, and every injected descendant, was then judged in chunks of up to five pairs per anchor with the same prompt and schema (rerank_score_floor_fill.py, chunk size 5). Calls use temperature 0 and seed 1. The primary analysis run used one shared developer prompt for every disorder (section SM8). It is reproduced first and the disorder-specific base prompt second.

Developer prompt that ran for every disorder (rerank_ambiguous_pairs_api_v3_mdd.py, function mdd_call_llm_bundle). The em dash character is shown as [U+2014].

```
You are reranking ambiguous cross-species phenotype candidate pairs for a
Major Depressive Disorder (MDD) face-validity pipeline aligned to DSM-5-TR.

Important constraints:
- You are not the primary scorer.
- Transparent non-LLM signals have already identified a small ambiguous bundle.
- Your job is only to promote, keep, demote, or abstain on those candidates.
- Ignore any hidden prior about disease genes or depression labels; you are not given them.
- Do not use any hand-built tier or relevance hierarchy; judge only behavioral/sensory/developmental analog quality.

The 9 MDD Criterion A domains for context:
  depressed_mood, anhedonia, weight_appetite_change, sleep_disturbance,
  psychomotor_change, fatigue, worthlessness_guilt,
  concentration_indecisiveness, suicidality.

Bidirectional-domain rule (CRITICAL [U+2014] MDD-specific):
- Three MDD Criterion A symptoms are INTRINSICALLY BIDIRECTIONAL per
  DSM-5-TR: A.3 weight_appetite_change (gain OR loss), A.4 sleep_disturbance
  (insomnia OR hypersomnia), A.5 psychomotor_change (agitation OR
  retardation). For HPO targets in these three domains, each target encodes
  ONE direction of the axis (e.g., HP:0000713 Agitation is the agitation
  direction of A.5; HP:0031843 Slow thought process is the retardation
  direction).
- When judging analog quality:
    * Same-direction mouse phenotype -> direct_analog or close_support
      (full credit).
    * Opposite-direction mouse phenotype that maps to the SISTER directional
      target on the same axis -> NOT a mismatch. The opposite-direction
      phenotype is the sister target's evidence, not this target's. Use
      weak_support if the phenotype loosely supports either direction; use
      mismatch only if the mouse phenotype is genuinely incompatible with
      the MDD axis (e.g., a metabolic phenotype unrelated to the
      psychomotor / sleep / weight axis).

Definitions:
- comparable_observable=true only if the mouse term is an organism-level or
  developmentally interpretable observable that could plausibly support the
  human target.
- evidence_scope=cellular_or_circuit_only if the term is primarily
  mechanistic physiology or cellular/circuit description rather than a
  behavioral/sensory/developmental organism-level phenotype.
- strict_eligible=true only for high-quality organism-level analogs suitable
  for strict MDD-domain support.
- broad_eligible=true for weaker but still meaningful support.
- abstain if none of the candidates are suitable organism-level analogs.

Do not invent new candidates. Evaluate only the listed bundle.
```

Disorder-specific base developer prompt (rerank_ambiguous_pairs_api_v3.py, function call_llm_bundle), which the same-LLM sensitivity run used with each disorder’s addendum for bundle pairs and alone for the other pairs (section SM8). The addendum sentence in its strict_eligible definition does not apply to the primary analysis run.

You are reranking ambiguous cross-species phenotype candidate pairs.

Important constraints:
- You are not the primary scorer.
- Transparent non-LLM signals have already identified a small ambiguous bundle.
- Your job is only to promote, keep, demote, or abstain on those candidates.
- Ignore any hidden prior about disease genes or disorder labels; you are not given them.
- Do not use any hand-built tier or relevance hierarchy; judge only behavioral/sensory/developmental analog quality.

Definitions:
- comparable_observable=true only if the mouse term is an organism-level or developmentally interpretable observable that could plausibly support the human target.
- evidence_scope=cellular_or_circuit_only if the term is primarily mechanistic physiology or cellular/circuit description rather than a behavioral/sensory/developmental organism-level phenotype.
- strict_eligible=true only for high-quality organism-level analogs suitable for strict domain support; the disorder-specific construct-validity criteria are appended by the wrapper as a developer_prompt_addendum.
- broad_eligible=true for weaker but still meaningful support.
- abstain if none of the candidates are suitable organism-level analogs.

Do not invent new candidates. Evaluate only the listed bundle.

User prompt (identical in both functions). The human target text is the anchor’s HPO label joined to its criterion phrase, and no criterion text is included.

Target ID: <anchor key>
Human target label: <anchor HPO label>
Human target text: <anchor HPO label>; <criterion phrase returned by decomposition>
Domain: <human domain label>

Ambiguous candidate bundle:
[1] mp_id=<id> | mp_term=<label> | mp_definition=<definition> | mouse_domain=<domain> | transparent_score=<0.000> | embed=<value> | lexical=<value> | resnik=<value> | jaccard=<value> | available_channels=<n> | structural_missing=<True or False>
[2] ...

Return evaluations for the listed candidates only. Use candidate_index values 1..<n>.

Response schema (RERANK_SCHEMA, identical in both functions):

```
{
 "name": "pair_rerank_bundle",
 "strict": true,
 "schema": {
  "type": "object",
  "additionalProperties": false,
  "properties": {
   "target_id": {
    "type": "string"
   },
   "selected_candidates": {
    "type": "array",
    "items": {
     "type": "object",
     "additionalProperties": false,
     "properties": {
      "candidate_index": {
       "type": "integer"
      },
      "rerank_action": {
       "type": "string",
       "enum": [
        "promote",
        "keep",
        "demote",
        "abstain"
       ]
      },
      "comparable_observable": {
       "type": "boolean"
      },
      "evidence_scope": {
       "type": "string",
       "enum": [
        "organism_behavior",
        "organism_sensory",
        "developmental_behavior",
        "cellular_or_circuit_only",
        "unclear"
       ]
      },
      "behavioral_analog_type": {
       "type": "string",
       "enum": [
        "direct_analog",
        "close_support",
        "weak_support",
        "mechanistic_only",
        "mismatch"
       ]
      },
      "strict_eligible": {
       "type": "boolean"
      },
      "broad_eligible": {
       "type": "boolean"
      },
      "reason": {
       "type": "string"
      }
     },
     "required": [
      "candidate_index",
      "rerank_action",
      "comparable_observable",
      "evidence_scope",
      "behavioral_analog_type",
      "strict_eligible",
      "broad_eligible",
      "reason"
     ]
    }
   }
  },
  "required": [
   "target_id",
   "selected_candidates"
  ]
 }
}
```

The score adjustment applied in code to the rerank action is +0.10 for promote, 0 for keep, -0.10 for demote and -0.25 for abstain, and the final pair score is clipped to 0 to 1. A pair is admitted when strict_eligible is true, behavioral_analog_type is direct_analog or close_support and the direction check of the unit does not veto it.

#### S1 Text, part 2. Supplementary Results

##### Sensitivity of the support AUROC

The separation does not depend on any one gene or on the minimum of four annotations, and it survives the restriction to behaviorally annotated genotypes. Intervals that resample genes in place of genotypes are 9 to 26% wider than genotype-level intervals, and none reaches chance. Keeping one random allele per gene moves the support AUROC by 0.018 or less for ASD, SCZ and NCD and raises IDD to 0.791 (0.763 to 0.813), because Mecp2 supplies 30 of the 111 IDD positives. Of those 30 Mecp2 genotypes, 16 score zero because they carry no behavioral annotation, so removing Mecp2 raises IDD support AUROC from 0.751 to 0.777 (0.717 to 0.835). Including every genotype with at least one annotation enlarges the universe to 17,472 to 19,432 genotypes and moves the support AUROC by 0.021 or less (ASD 0.950, SCZ 0.661, IDD 0.742, NCD 0.670). The IDD background also holds the 157 genotypes of syndromes with obligate intellectual disability that carry no IDD label. Counting them as positives lowers the IDD support AUROC to 0.692, or 0.727 with only their behaviorally annotated genotypes. Most of them carry no behavioral annotation and score low, so the primary IDD value overstates agreement with all genotypes of these syndromes by about 0.06. A genotype that no random annotation set matches as well reaches the maximum support score. For ASD, 323 genotypes tie at this maximum, 62 of them curated models (19.2%, 27 times prevalence), and for IDD 380 tie, 43 of them curated (11.3%, 13 times prevalence). SCZ and NCD genotypes never reach it. A re-test with 1,000,000 permutations leaves 314 of the 323 ASD and 326 of the 380 IDD genotypes tied, so the maximum is reported as a tie block. The catalogue orders the block by a standardized support score as a labeled secondary order, whose Spearman correlation with the re-test order is 0.23 for ASD and 0.42 for IDD. The order within the block is therefore not a ranking. The values are collected in S8 Table.

##### Contrasts between the support score and the retrieval score at equal depth

The retrieval score falls to 0.768, 0.540, 0.603 and 0.654 under the same weights. At equal depth the support score exceeds the retrieval score for ASD (+0.128, 0.086 to 0.165, p below 0.001) and IDD (+0.064, 0.009 to 0.135). The SCZ difference includes zero (+0.056, -0.041 to 0.146), as does the NCD difference, whose point estimate is negative (-0.072, -0.150 to 0.013). The two scores differ in the pairs they read, in how they combine them and in the permutation null, so these differences cannot be attributed to the LLM’s admission alone. Computed on the admitted pairs alone at equal depth, the retrieval score gives 0.893, 0.397, 0.650 and 0.579. For SCZ the admitted terms alone therefore rank curated models below equally deep background genotypes, and the SCZ separation comes from the comparison with random annotation sets of the same size. Without depth matching, the retrieval score computed on the admitted pairs alone gives 0.960, 0.590, 0.747 and 0.672 for ASD, SCZ, IDD and NCD (S8 Table).

##### PhenoDigm configurations

No automated method evaluates a genotype against individual diagnostic criteria, so the comparison with PhenoDigm (a widely used cross-species phenotype similarity method, Smedley et al. 2013) concerns the support AUROC only, in four configurations defined in Methods. The primary configuration recomputes PhenoDigm over the full phenotype profile without the published table’s similarity cutoff, which raises its score for every disorder.

FACET leads PhenoDigm recomputed for ASD by 0.184 (0.126 to 0.248) and does not differ significantly for IDD (S10 Table). FACET also leads for NCD (+0.075, 0.007 to 0.150, p 0.031) and trails for SCZ (-0.090, -0.191 to 0.001, p 0.054), both before correction. Neither comparison is in the corrected family, which was fixed on the behavioral configuration below before the recomputed configuration became primary. Against the published table FACET leads for ASD and IDD, leads for NCD before correction and does not differ for SCZ. A third configuration restricts PhenoDigm to behavioral phenotypes (the branch in which every admitted term lies), so that both methods read the same part of the record. It is the configuration in the Holm family (Methods), where FACET’s ASD lead survives correction while the other three differences are not significant (S4 Table). The two scorers converge once they read the same pairs. Applied to FACET’s own admitted pairs, the PhenoDigm scorer gains over its behavioral configuration in every disorder and the two then differ by at most 0.062. On the same admitted pairs the retrieval score gives 0.590 for SCZ, below both PhenoDigm (0.744) and FACET (0.682, Table 3 of the main text). The difference between the methods therefore lies in the pairs each reads and in the referral to the null. The ASD support AUROC stays between 0.94 and 0.96 whether 145 or 564 pairs are admitted (S23 Table), so agreement between the two scorers on ASD says little about which pairs should be admitted.

*S10 Table. FACET against PhenoDigm on discrimination, in the genotypes carrying a candidate term. Published uses the Monarch table as distributed. Recomputed applies PhenoDigm to the full phenotype profile without the published cutoff. Behavioral restricts PhenoDigm to behavioral phenotypes. Admitted pairs applies the PhenoDigm scorer to the pairs FACET admitted. Differences are FACET minus PhenoDigm with 95% gene-clustered intervals and the two-sided bootstrap p.*

| Disorder | FACET | PhenoDigm published | PhenoDigm recomputed | PhenoDigm behavioral | PhenoDigm on FACET’s admitted pairs | Difference, published | Difference, recomputed | Difference, behavioral | Difference, admitted pairs |
| --- | --- | --- | --- | --- | --- | --- | --- | --- | --- |
| ASD | 0.956 | 0.656 | 0.772 | 0.687 | 0.923 | +0.300 (0.226 to 0.382, p below 0.001) | +0.184 (0.126 to 0.248, p below 0.001) | +0.269 (0.236 to 0.301, p below 0.001, Holm p 0.002) | +0.033 (0.025 to 0.041, p below 0.001) |
| SCZ | 0.682 | 0.725 | 0.772 | 0.640 | 0.744 | -0.043 (-0.143 to 0.046, p 0.34) | -0.090 (-0.191 to 0.001, p 0.054) | +0.043 (-0.053 to 0.135, p 0.36, Holm p 0.51) | -0.062 (-0.144 to 0.016, p 0.10) |
| IDD | 0.751 | 0.616 | 0.706 | 0.686 | 0.732 | +0.135 (0.068 to 0.240, p below 0.001) | +0.045 (-0.028 to 0.158, p 0.28) | +0.065 (-0.012 to 0.171, p 0.10, Holm p 0.29) | +0.019 (0.003 to 0.029, p 0.024) |
| NCD | 0.668 | 0.557 | 0.593 | 0.623 | 0.668 | +0.111 (0.013 to 0.203, p 0.026) | +0.075 (0.007 to 0.150, p 0.031) | +0.045 (-0.028 to 0.126, p 0.22, Holm p 0.51) | 0.000 (-0.025 to 0.023, p 0.98) |

##### Alternate LLMs: composition of the extra admissions and the same-LLM prompt run

Re-judging every pair with either of two alternate LLMs leaves the ASD result within the pre-specified margin and raises the SCZ result. Every primary pair was re-judged by claude-opus-4-8 and gemini-2.5-pro and scored through the same pipeline (S23 Table). Both leave the ASD support AUROC within the pre-specified equivalence margin of 0.05 (equivalent when the 90% interval of the difference lies inside -0.05 to 0.05, not equivalent when it lies wholly outside and inconclusive otherwise). Both raise SCZ (by 0.182 and 0.159), so the SCZ result depends on which LLM judges. IDD is equivalent under claude-opus-4-8 and rises under gemini-2.5-pro by 0.090 (0.023 to 0.138), an interval that crosses the margin and is therefore inconclusive, and NCD is inconclusive under both. With 67 curated models the margin of 0.05 is narrower than the attainable NCD interval, so the two inconclusive NCD verdicts reflect power, while the SCZ intervals lie above the margin under both alternate LLMs. The three LLMs differ mainly in how readily they admit a pair. The alternate LLMs keep 87 to 99% of the pairs the primary LLM admitted and admit 2.2 to 4.2 times as many within the same candidate set. Of the 255 extra SCZ pairs that claude-opus-4-8 admits, 109 lie on A4, 21 on A5, 5 on A1 to A3 (S33 Table) and 120 on anchors of the unscored criterion B and of supporting text. Only 11 carry the endophenotype or drug-response terms the mapping rules name. The gemini-2.5-pro extras (502) divide the same way (162 on A4, 54 on A5, 41 on A1 to A3, 245 on unscored anchors and 33 on endophenotype or drug-response terms). The SCZ rise therefore comes from a broader reading of disorganized behavior and negative symptoms, while the terms the rules exclude account for few of the extra admissions. Because admitted pairs are a small fraction of the judged pairs, agreement on admission is high (90 to 97%) while kappa (agreement beyond chance, 0 at chance and 1 when perfect) is 0.34 to 0.54 (S23 Table). The two alternate LLMs agree with each other (kappa 0.53 to 0.66) more than with the primary LLM, which is the strictest of the three. The primary analysis run used one shared developer prompt for every disorder (Methods). Rerun with each disorder’s own prompt, the same LLM admits 125, 176, 137 and 108 pairs against 145, 196, 138 and 95 in the primary analysis run, so its strictness does not come from the prompt. It moves the support AUROC by -0.029 (90% interval -0.055 to 0.000) for ASD, -0.021 (-0.070 to 0.023) for SCZ, +0.045 (0.013 to 0.068) for IDD and -0.016 (-0.036 to 0.003) for NCD. Between the two prompts, 19 to 32% of the admitted set turns over.

##### Stability of the upstream LLM steps

The upstream LLM steps are more stable than the pair judgment. Re-attributing the anchors with the alternate LLMs agrees with the primary analysis run on all but at most two anchors for ASD, IDD and NCD and disagrees on 10 and 11 of the 41 SCZ anchors. Both alternate LLMs place five SCZ anchors elsewhere than the primary analysis run, among them diminished social interaction (under A5 for both alternate LLMs and under the unscored criterion B in the primary analysis run). The attribution of record is retained, and no downstream analysis was rerun with the alternate attributions. On a pre-specified sample of the mouse taxonomy the two alternate LLMs bracket the primary analysis run, one stricter and one more permissive. SCZ mapping is the least stable, since one alternate LLM keeps only half of the MP terms the primary analysis run assigns to an SCZ criterion (S23 Table). The re-attribution and reclassification values are in S12 Table.

##### Expert ratings in detail

Human raters agree with most of the pipeline’s refusals and with two in three of its admissions (Figure 5, S13 Table). Interim ratings from eight raters (seven of the 12 invited raters and the first author, section SM5) cover all 168 pairs (336 ratings). They agree with the pipeline on 88% of the sampled refused pairs, most of which lie in the stratum farthest from admission, and confirm 66% of admitted pairs. The pre-specified rule requires both rates to reach 0.85 with one-sided lower bounds of at least 0.70 for expert-supported, and reads either rate below 0.70 as not expert-confirmed. The rejection rate of refused pairs passes the bar (0.884, lower bound 0.824), while the confirmation rate of admitted pairs (0.662) falls below the floor, so the pair judgment is not expert-confirmed at this sample size. “Cannot tell” is excluded from these rates. Counting “cannot tell” as a rejection of the pair gives confirmation 0.54 and rejection 0.92 over ratings, and counting it as an acceptance gives 0.72 and 0.69. The LLM agrees with the rater majority on 79% of pairs (kappa 0.64). Two raters rating the same pair agree with each other 79% of the time (107 pairs, pair bootstrap 95% interval 0.71 to 0.87), and the LLM agrees with a single rater on 77% of ratings (section SM5). Much of the gap lies in how strong an analog must be to count. On the four-step analog scale (unrelated, weak, close, direct), raters grade the pairs the LLM called direct 0.83 steps lower on average and those it called close 0.31 steps lower. On the first rating question alone (whether the phenotype belongs to the criterion), raters agree with the mouse criterion mapping step on 79% of the terms it assigned to the criterion and 73% of the rest. Among ratings of admitted pairs, 13% reject the criterion match, 15% accept it but grade the analog weak, unrelated or unsure and 18% answer “cannot tell”. Confirmation of admitted pairs by disorder (with “cannot tell” excluded) is 77% for IDD, 75 for MDD, 67 for ASD, 62 for NCD and 52 for SCZ, and the raters agree with every SCZ refusal. The expert ratings and the admission control therefore single out the same two disorders. Some refused pairs had been found broadly eligible (bearing on the criterion) by the LLM or had been mapped to the criterion, and strict eligibility (a high-quality organism-level analog) then stopped them. Of the 36 definite ratings of such pairs, 13 (36%) admit the pair, against 1 of 94 ratings of pairs that met neither. Raters differ in leniency, and the between-rater spread (tau 0.21 on admissions) is the largest component of the rater-adjusted interval. No choice among the LLMs closes the gap. Four LLM runs were available for a vote: the primary analysis run, the same LLM under the disorder-specific prompt and the two alternate LLMs. Every rule that admits a pair when at least k of the four admit it agrees with the rater majority at 0.76 to 0.81, against 0.79 for the primary analysis run. The rules’ own admissions are confirmed at 0.65 to 0.71. Pairs refused by all four LLMs are rejected by the raters at 0.96. Removing the first author’s ratings gives confirmation 0.654 and rejection 0.880, with the same verdict.

##### Human genetics, phenotyping attention and the describing literature

For ASD and IDD, uncurated genotypes of genes implicated by human genetics (which the pipeline never reads) reach the top coverage tier more often than other uncurated genotypes, although these genes were also phenotyped more often. Background genotypes of genes implicated by human genetics without mouse data (H genes) reach the top ASD coverage tier (Methods) at 2.47 (1.64 to 3.73) times the adjusted odds of other background genotypes. The IDD odds ratio is 1.74 (1.32 to 2.30), and SCZ, NCD and MDD show no such effect (1.51, 1.08 and 0.83, every interval including 1). After adjustment for publication count, H-gene genotypes are also behaviorally annotated more often than other background genotypes for ASD (adjusted odds ratio 1.83, 1.57 to 2.13) and NCD (2.43, 2.20 to 2.68). The same holds for MDD (35.1 against 26.0%, 1.68, 1.39 to 2.04). The IDD excess is small (1.13, 1.06 to 1.21) and SCZ shows none (1.08, 0.69 to 1.67, S14 Table). Part of the higher coverage of H-gene genotypes is therefore phenotyping attention. The papers describing the top background candidates discuss the disorder more often than those of matched controls (a literature contrast that shares the dependence on the describing paper noted above). PubTator3 (a text-mining service that tags the diseases a paper discusses) tags a paper describing the genotype with the disorder for 40.2% of the ASD candidates. The rate is 5.7% among controls matched on behavioral depth and annotation source (risk ratio 7.0, 95% interval 4.2 to 11.7). The IDD rates are 34.4 against 11.6% (risk ratio 3.0, 2.1 to 4.1). SCZ has the literature contrast without the coverage effect (14.2 against 2.8%). NCD, which has no human-genetics list and uses its MGI disease-gene associations in its place, is the negative case on both counts. Its top background candidates by support score are cerebellar and channel mutants (Grid2, Reln, Scn8a) whose papers rarely discuss dementia (1.0 against 14.0%), while uncurated genotypes of dementia genes (Snca, Htt, App, Mapt) rank lower.

##### Genotypes meeting the strictest ASD requirement

The four genotypes meeting the strictest ASD requirement include Syngap1 and Tcf4 (neither curated for ASD). Their human orthologs cause intellectual disability (SYNGAP1, Hamdan et al. 2009), with autism reported in later cases (Hamdan et al. 2011), and Pitt-Hopkins syndrome (TCF4, Zweier et al. 2007 and Amiel et al. 2007). This illustrates what the catalogue surfaces without testing it, since the base rate of such genes among lower-coverage genotypes was not computed.

##### The IMPC-only test

The International Mouse Phenotyping Consortium phenotypes knockout genotypes by one fixed battery, independent of any disease claim, and 5,126 genotypes of the set carry an IMPC, KOMP, EUCOMM or NCOM allele. Within the IMPC-tagged genotypes of each evaluation set, with curated models removed, the genotypes whose gene is an H gene (S5 Table) were ranked against the other IMPC genotypes (S17 Table). The support score, Diagnostic Coverage, the behavioral annotation count and PhenoDigm recomputed all sit at chance in every disorder, with every gene-clustered interval including 0.5 except the IDD support AUROC (0.521, 0.498 to 0.546), and depth matching leaves the support AUROC at 0.49 to 0.52. Positives and background are behaviorally annotated at similar rates (82 to 89% against 82 to 86%), and 64 to 72% of the positives carry one or two behavioral terms, which reflects the IMPC adult battery rather than the genes. Script and result: audit/impc_test_2026-11-07/.

##### The depth-stratified null and PhenoDigm referred to the same nulls

Both analyses were added after review (post hoc) and run from the frozen verdicts. The stratified null draws, for a genotype with k distinct terms of which b lie at or below abnormal behavior, b terms from the 547 behavioral terms and k minus b from the other annotated terms, 10,000 draws per (k, b) stratum (542 to 548 strata per disorder), with the score function, cap and p primary value (permutation_null_depthstrat.py). The null itself moves: for SCZ, 1,218 of the 1,248 scored genotypes with a nonzero support score change their minus log10 p (mean change -0.39), and the null mean at k 10 rises from 0.003 to 0.043 when all ten terms are behavioral. The support AUROC does not move (S18 Table), because 84 to 92% of the background has no admitted term and scores zero under either null. The indicator of carrying any admitted term gives support AUROC 0.896 (ASD), 0.684 (SCZ), 0.721 (IDD) and 0.680 (NCD) by itself. Among the genotypes that carry one (81, 19, 68 and 39 curated models against 1,921, 1,039, 2,097 and 2,557 background genotypes), the support score ranks curated models at 0.890, 0.442, 0.788 and 0.404 under the null of record and at 0.896, 0.441, 0.796 and 0.381 under the stratified null. Six SCZ genotypes reach the cap under the stratified null and none did before: they are credited through terms outside the behavior branch and carry no behavioral term.

PhenoDigm recomputed (arm 3) was referred to the same two null constructions by its own symmetric best-match average over random sets (audit/phenodigm_null_2026-11-07/phenodigm_null.py, self-checked against the arm 3 primary scores). Size-matched, PhenoDigm moves by at most 0.020 (ASD, +0.020, p 0.001) and the record comparison is unchanged, except that the SCZ deficit of FACET reaches p 0.033 before correction (-0.099, -0.198 to -0.013). Behavior-matched, PhenoDigm falls by 0.115 (ASD), 0.092 (SCZ), 0.082 (IDD) and 0.052 (NCD), all p below 0.001, while FACET is unchanged, so FACET leads by +0.298, +0.127 and +0.127 for ASD, IDD and NCD and ties SCZ (+0.002, -0.118 to +0.108). PhenoDigm’s score is continuous and every genotype has one, so behavioral depth reorders the whole universe. FACET’s score is gated by admission, so depth can reorder only the admitted minority. Scripts and results: audit/nulldepth_2026-11-07/ and audit/phenodigm_null_2026-11-07/.

##### The SCZ mapping rules and the endophenotype terms

Whether the SCZ result depends on the mapping rules was checked on the record (2026-11-07, read-only). The 42 curated SCZ models carry 234 behavioral annotations (term occurrences at or below MP:0004924), of which 66 sit in relevance tier 1, 143 in tier 2 and 25 in tier 3, and none in tiers 4 or 5. Every behavioral term a curated SCZ model carries was therefore a candidate for judgment, and the rules removed none from candidacy. The terms the rules name as endophenotype or drug-response readouts reached the pair judgment. In the SCZ candidate primary table, 115 pairs carry prepulse inhibition, sensorimotor gating, latent inhibition, startle, amphetamine-response or methamphetamine-response terms (18 MP terms). Of these, 59 are against anchors of criteria A1 to A5, and 56 against anchors outside the five criteria (associated features such as sensory behavioral abnormality, aggressive behavior and abnormal affect), which credit no criterion. On the criterion anchors, 58 of the 59 pairs reached the judgment and none was strictly eligible (26 weak support, 22 mismatch, 9 mechanistic only, 1 close support below the strict requirement), under the shared primary developer prompt, which carries no SCZ rule (section SM8). On the anchors outside the criteria, five startle and gating pairs were strictly eligible (against sensory behavioral abnormality and reduced attention regulation), and they credit no criterion because those anchors are not scored units. S20 Table gives the pairs on criterion anchors by term family, anchor and grade. The SCZ result is therefore the judgment’s verdict on those terms.

##### SCZ A5 without the broad emotion term

SCZ A5 (negative symptoms) is credited through abnormal emotion/affect behavior (MP:0002572) and bradykinesia (MP:0005156). With the broad term removed and bradykinesia kept, A5 is credited to no curated model (from 31 of 42) and to 72 background genotypes (from 1,834), and the six curated models credited with A5 alone lose their only credit, so 36 of 42 models with any credited unit becomes 30. The A5 coverage of record is read as an upper bound.

##### Reproducibility of the verdicts

The reproducibility of the verdicts is set by the judgment floor, the instruction prompt and the served LLM snapshot. The judgment floor of 0.57 and the channel weights were set on ASD against the LLM’s own verdicts and applied unchanged to every disorder. The floor was swept on the primary analysis run (S30 Table, section SM14). The floor only selects which pairs outside a bundle reach the LLM by the floor route, and bundle pairs and injected descendants are judged whatever their score, so a higher floor un-judges the floor-route pairs below it and nothing else. At 0.60 the support AUROC is equivalent to the record in every disorder, and at 0.65 it is equivalent for ASD and IDD and falls for SCZ and NCD. A higher floor bounds the cost of the floor only. The gain of a lower floor was not measured, since it needs judgments the record never made (40 to 80 pairs per disorder at 0.55 and 165 to 333 at 0.50). Under a change of instruction prompt within one LLM, 19 to 32% of the admitted set turns over while the support AUROC moves by 0.045 or less (Results). The AUROC therefore cannot certify the admitted set, and the expert study is the decisive test of it. The pair-judgment script logged only the parsed verdicts and not the snapshot the provider served under the alias gpt-5.4. The snapshot is inferred from the record (2026-11-08, read-only). The pair judgments ran on the alias with seed 1 between 2026-06-04 (MDD) and 2026-09-10 (the IDD top-up), the four primary disorders on 2026-07-28 and the NCD B rescue on 2026-07-31. Every call to the same alias whose served model was logged returned gpt-5.4-2026-03-05: the criterion-mapped taxonomy runs and pilots of 2026-07-27 and 2026-07-28 (about 11,600 calls), the MDD domain labeling of 2026-06-04 and the gate rerun of 2026-11-08, about 23,000 calls in all with no exception. The alias therefore resolved to one snapshot from June to November, and the pair judgments fall inside that window. This is an inference from the other steps and not a log of the pair judgment itself, so the frozen verdicts remain the record. A future iteration of this work should build the LLM steps with experts involved from the start. The seed terms, the analog grades and the admission standard would then be set by the readers whose agreement is the decisive test. The released record permits that change without rerunning any judgment.

##### Further supplementary results

Values in this section come from the record of the run and its pre-specified or logged analyses. A value computed under the labels of 2026-10-08, which held 73 NCD positives before the two Wolff-Parkinson-White genotypes were removed, is marked as such. ASD, SCZ, IDD and MDD labels are identical in the two label sets.

##### Counting genotypes assayed with normal results leaves both measures in place

Treating genotypes assayed with normal results as assessed changes no AUROC by more than 0.015 under the pre-specified rule and raises most values under the post hoc rule. The pre-specified analysis restricted both classes to genotypes that are behaviorally annotated or carry an MGI normal behavioral flag, which adds 76 genotypes to the behaviorally annotated set. The full-background support AUROCs are then ASD 0.961, SCZ 0.662, IDD 0.747 and NCD 0.632, and the MDD coverage AUROC is 0.982. The post hoc analysis also counts the 1,795 genotypes that completed an IMPC behavioral procedure with no abnormal call (section SM7). The support AUROC then moves from 0.9608 to 0.9657 for ASD (0.943 to 0.983), from 0.6617 to 0.6734 for SCZ, from 0.7605 to 0.7565 for IDD and from 0.6348 to 0.6605 for NCD (0.581 to 0.727). Coverage AUROCs move from 0.944 to 0.952 (ASD), 0.732 to 0.767 (SCZ), 0.735 to 0.741 (IDD), 0.598 to 0.641 (NCD) and 0.982 to 0.985 (MDD). Intervals are gene-clustered with 1,000 replicates, and the NCD values use the labels of 2026-10-08. Adding assayed normals adds low-scoring background genotypes, so no separation depended on treating unassayed genotypes as negatives.

##### MGI records almost no explicit negatives, and those it records mark mechanism and not phenotype

MGI’s records stating that a genotype is not a model of a disease were matched to the scored set by allele pair and genetic background together. They number 1 for ASD (Tg(Camk2a-Bdnf)), 0 for SCZ and IDD, 12 for NCD and 1 for MDD (Pclo). The 12 NCD records are null alleles of genes whose disease acts through gain of function or a toxic protein (Htt 6, Prnp 2, Hap1 2, Apba2 1 and Drd2 1). Against these 12, curated NCD models score a full-background support AUROC of 0.655 (0.446 to 0.760) and a coverage AUROC of 0.572 (0.271 to 0.729), with 73 curated models under the labels of 2026-10-08. One record is discordant. Csmd1 homozygotes are a curated SCZ model on a mixed 129S5 and C57BL/6NTac background in the scored set, and a record states that they are not a model on a mixed B6 and 129S5 background outside it. A labeled-negative benchmark cannot be built from MGI, so uncurated genotypes are treated as unlabeled background.

##### Genotypes credited for one disorder are credited for others more often than behavioral depth predicts

High support scores for two disorders co-occur above depth-matched expectation, and the shared vocabulary explains most of it. The overlap ratio counts the uncurated genotypes in both disorders’ top 2% of support scores. It divides that count by the number expected at fixed behavioral depth (strata 0, 1 to 2, 3 to 5, 6 to 10 and 11 or more behavioral annotations). It is 4.58 for IDD and NCD, 2.55 for ASD and SCZ and 2.36 for ASD and IDD. Curated models also reach other disorders’ top coverage tiers above depth-matched expectation. Curated ASD models reach the IDD tier 3.6 times as often, curated SCZ models reach the ASD tier 4.0 times and the IDD tier 3.3 times, and curated IDD models reach the ASD tier 2.4 times. Curated NCD models almost never reach another disorder’s tier.

ASD and IDD share 13 admitted MP terms (all social terms plus cognitive inflexibility). IDD and NCD share 35 (nearly all learning and memory terms plus ataxia), and ASD and SCZ share 7 (stereotypy, circling, weaving and aggression). Removing the shared terms and rescoring, as pre-specified, brings the ratios to 1.05 for IDD and NCD (chance), 1.83 for ASD and IDD and 2.15 for ASD and SCZ. The ASD and SCZ correlation within depth strata strengthens from 0.248 to 0.357 after removal and is carried by distinct terms (social terms for ASD, locomotor and affect terms for SCZ). The ASD and IDD correlation among uncurated genotypes falls from 0.316 to -0.007, so it rests entirely on the 13 shared social terms. IDD’s adaptive-function anchors are social, so this is overlap between the criteria themselves. Removing shared terms lowers the support AUROC where they are the disorder’s own core content. ASD loses 0.093 (0.055 to 0.142) without the social terms shared with IDD, and IDD loses 0.101 (0.063 to 0.162) without the cognitive terms shared with NCD. ASD and SCZ each lose 0.007. SCZ and MDD overlap about twice expectation in the top tier after removal, while their correlation is null (0.027, -0.016 to 0.071). Correlations are Spearman coefficients pooled across depth strata by Fisher z. These analyses use the labels of 2026-10-08.

##### A change of prompt within one LLM turns over a fifth to a third of the admitted set while barely moving the AUROC

Rerun under the disorder-specific prompts (section SM8), the primary LLM refuses 47 of its 145 ASD admissions, 51 of 196 SCZ, 36 of 138 IDD and 18 of 95 NCD. It admits 27, 31, 35 and 31 pairs it had refused. The turnover is therefore 19 to 32% of the admitted set per disorder, while the support AUROC moves by 0.045 or less. The two prompts agree on strict eligibility for 98.1 to 98.9% of the primary pairs (kappa 0.72 to 0.77) and on the analog grade at kappa 0.56 to 0.70. Admissions are rare, so a small share of disagreements on all pairs is a large share of the admitted set. Under the same disorder-specific prompt, claude-opus-4-8 agrees with gpt-5.4 on strict eligibility for 94.8 to 96.0% of pairs (kappa 0.42 to 0.59). It admits 2.1 to 3.0 times as many pairs (370 against 125, 425 against 176, 363 against 137 and 224 against 108).

##### No rule combining several LLM runs raises expert confirmation of admissions to the bar

Rules of the form “admit when at least k of n LLM runs admit” were compared with the eight raters after the fact (post hoc). The comparison covers the 140 rated pairs of the four primary disorders with a definite rating. The four runs are the record, gpt-5.4 under the disorder-specific prompts, claude-opus-4-8 and gemini-2.5-pro. On the composite verdict every such rule agrees with the rater majority at 0.76 to 0.81, against 0.793 for the record alone (kappa 0.635). Each rule’s own admissions are confirmed at 0.65 to 0.71, and no single run or rule exceeds 0.72, against the 0.85 the declared rule requires. Pairs refused by all four runs are rejected by the raters at 0.96. On the membership question alone, requiring all four runs to agree raises agreement from 0.65 to 0.81 and confirmation from 0.59 to 0.75. This locates the permissive field in the record’s comparable-observable flag and not in strict eligibility. On the analog grade the record has the highest quadratic weighted kappa with the raters among single runs (0.54) and a mean step difference near zero (-0.02). Raters grade the pairs the LLM called direct analogs 0.83 steps lower on average and its close analogs 0.31 steps lower. Every instrument pair now carries a rating, so no pairs are held out and any rule chosen here is exploratory.

A majority vote gives a further comparison (post hoc). It combines, field by field, the three runs under the disorder-specific prompts (gpt-5.4, claude-opus-4-8 and gemini-2.5-pro), with the record breaking a three-way split. The three runs are unanimous on strict eligibility for 90 to 92% of judged pairs (Fleiss kappa ASD 0.43, SCZ 0.46, IDD 0.51, NCD 0.59). The majority vote admits 284, 356, 323 and 183 pairs for ASD, SCZ, IDD and NCD. Its support AUROC is 0.952 for ASD, 0.765 for SCZ and 0.777 for IDD, against the record a difference of -0.004 (equivalent), +0.083 (+0.014 to +0.149, inconclusive) and +0.026 (equivalent). The NCD comparison was computed under the labels of 2026-10-08 (-0.015, equivalent). Against the eight raters on the 140 rated pairs of the four primary disorders with a definite rating, the majority vote admits 62 pairs where the record admits 71. It agrees with the rater majority on 0.814 of pairs (0.754 to 0.871, kappa 0.672) against 0.793 (kappa 0.635) for the record, confirms 0.710 of its admissions against 0.662 and rejects 0.878 of its refusals against 0.906. Its confirmation rate clears the 0.70 floor of the declared rule and stays below the 0.85 bar, so the majority vote reads as measured. Against the rater majority on the same 140 pairs, precision (confirmation of admitted) and recall (share of the 53.5 rater-admitted pairs admitted, a tied pair counting half) are 0.662 and 0.879 for the record, 0.718 and 0.657 for the primary LLM under the disorder-specific prompt, 0.679 and 0.850 for claude-opus-4-8, 0.679 and 0.838 for gemini-2.5-pro and 0.710 and 0.822 for the majority vote. Recall is measured on an instrument drawn from the record’s verdicts, so pairs that only another run admits are represented only through the refused strata (PANEL_RULES_VS_RATERS_PR_ABCDEGHL_2026-11-17.md). The comparison is post hoc, and the record remains the primary result because every validation sample was drawn from it.

##### Raters would admit about one in three near-miss refusals and almost no clear refusal

Agreement with refusals depends on how close the refused pair came to admission. The counts are ratings of refused pairs, with “cannot tell” excluded. Raters agree with 93 of 94 ratings in S4 (no criterion assignment and no eligibility) and 10 of 14 in S3 (criterion assignment without eligibility). They agree with 13 of 22 in S2 (broad eligibility only). The raters would therefore admit 13 of the 36 rated near-miss refusals (36%) against 1 of 94 clear refusals. This is the recall cost of strict eligibility. By disorder, ratings of admitted pairs confirm 23 of 30 for IDD, 29 of 43 for ASD, 9 of 12 for MDD, 16 of 26 for NCD and 14 of 27 for SCZ. Raters agree with every rated SCZ refusal (29 of 29).

##### Call counts and usage of the primary analysis run

The pair primary judgment made 4,294 LLM calls: 1,419 for ASD, 1,394 for SCZ, 890 for IDD, 533 for NCD and 58 for MDD. The counts are the response records of the bundle and remaining-pair files of each run (ASD 39 and 1,380, SCZ 36 and 1,358, IDD 20, 778 and a top-up of 92, NCD 21 and 512, MDD 32 and 26). Mouse criterion mapping made 7,415 calls, one per MP term in scope (ASD 1,655, SCZ 1,657, IDD 2,393 and NCD 1,710). Token counts for the runs of May to September 2026 were not retained. Usage of the validation runs comes from the providers’ exports. OpenAI recorded 4,812 requests in October 2026 (9.8 million input and 2.0 million output tokens). Anthropic recorded 21.8 million input and 5.2 million output tokens over June, July and October 2026. Google billed 72 USD from August to November 2026.

##### Database drift

The scores are stable against six months of database change. Rescored on MGI’s report of 2026-10-05 and MP release 2026-09-23 with the frozen verdicts, the coverage AUROC moves by 0.0006 or less and the support AUROC by 0.0013 or less in every disorder.

#### Supplementary Tables

##### S1 Table. Criterion units, symptom domains and diagnostic rules per disorder

Glosses are paraphrases written for this table. No DSM-5-TR text is reproduced. Direction is the direction of change each unit declares in its configuration, applied in code after the LLM answers and shown to no LLM. Rule names are those of criterion_rules.py that each disorder opts into (S26 Table).

###### Part A. Scored criterion units

| Disorder | Unit | Paraphrased gloss | Declared direction |
| --- | --- | --- | --- |
| ASD | A1 | Reduced reciprocity in social and emotional exchange | deficit |
| ASD | A2 | Reduced nonverbal communication such as eye contact and gesture (no MP vocabulary, measured zero) | deficit |
| ASD | A3 | Difficulty forming, keeping or understanding relationships | deficit |
| ASD | B1 | Repetitive or stereotyped movement, object use or speech | increase |
| ASD | B2 | Insistence on sameness and rigid routines | increase |
| ASD | B3 | Narrow interests held with unusual intensity | either |
| ASD | B4 | Over- or under-reactivity to sensory input, or unusual sensory interest | either |
| SCZ | A1 | Delusions (no organism-level analog, flagged in the mapping prompt) | either |
| SCZ | A2 | Hallucinations (no organism-level analog, flagged in the mapping prompt) | either |
| SCZ | A3 | Disorganized speech (no organism-level analog, flagged in the mapping prompt) | either |
| SCZ | A4 | Disorganized or catatonic behavior, both poles admissible | either |
| SCZ | A5 | Negative symptoms, reduced emotional expression and motivation | deficit |
| IDD | A | Reduced reasoning and learning ability (one example domain suffices) | deficit |
| IDD | B | Reduced ability to manage everyday activities (one example domain suffices) | deficit |
| NCD | A | Acquired decline in at least one of six cognitive areas (attention, executive control, learning and memory, language, perceptual-motor function and social cognition) | deficit |
| NCD | B | Loss of independence in everyday activities | deficit |
| MDD | A1 | Depressed or irritable mood | not declared in a criterion spec |
| MDD | A2 | Loss of interest or pleasure | not declared in a criterion spec |
| MDD | A3 | Change in weight or appetite | bidirectional (config) |
| MDD | A4 | Sleeping too little or too much | bidirectional (config) |
| MDD | A5 | Psychomotor agitation or slowing | bidirectional (config) |
| MDD | A6 | Fatigue or loss of energy | not declared in a criterion spec |
| MDD | A7 | Low self-worth or undue guilt | not declared in a criterion spec |
| MDD | A8 | Difficulty thinking, concentrating or deciding | not declared in a criterion spec |
| MDD | A9 | Repeated thoughts of dying or of suicide | not declared in a criterion spec |

Criteria not scored (from excluded_criteria of each spec): ASD C (onset), D (impairment), E (exclusion). SCZ B (decline in functioning, judged as a provisional unit but outside the primary denominator of 5), C (duration), D, E, F (exclusions and qualifiers). IDD C (developmental onset). NCD C and D (exclusions). MDD has no criterion spec file (its mapping predates the criterion specs), and its nine A items are the scored units through the carried-over MDD configuration.

Unit grain (from scored_grain and breadth_denominator): ASD item, 7 units. SCZ item, 5 units (6 with provisional B in an opt-in arm). IDD criterion, 2 units. NCD criterion, 2 units. MDD item, 9 units.

Grounding rules opted into (extra_rules): ASD grooming_routing, involuntary_motor_sign, asd_construct_refusals. SCZ scz_motor_boundary_from_dsm, mechanism_is_not_face_validity. IDD human_schooling_constructs. NCD decline_versus_deficit, language_floor. MDD none.

###### Part B. Symptom domains of the mouse domain-labeling list

Symptom domains are domains.strict of each disorder configuration. Terms labeled is the number of the 11,485 annotated MP terms whose primary_domain in the primary taxonomy carries that label. Other labels are the non-symptom labels the same step used (support, mechanism and out-of-scope classes).

| Disorder | Symptom domain (terms labeled) | Other labels used (terms labeled) |
| --- | --- | --- |
| ASD | social_interaction (12), communication (3), repetitive_restricted (10), sensory_reactivity (11) | cognitive_behavioral_support (212), developmental_support (137), motor_support (111), generic_neurological_abnormality (14), mechanistic_neurobiology (1,052), non_neurological_or_other (9,923) |
| SCZ | delusions (0), hallucinations (0), disorganized_speech (1), catatonic_disorganized_behavior (67), negative_symptoms (35) | cognitive_associated (62), mechanism_substrate (82), mechanistic_neurobiology (1,385), non_neurological_or_other (9,853) |
| IDD | intellectual_functioning (61), adaptive_functioning (100), developmental_onset (13) | generic_neurological_abnormality (625), mechanistic_neurobiology (917), non_neurological_or_other (9,769) |
| NCD | cognitive_executive (90), motor_coordination (215), social_cognition_empathy (31), adaptive_functioning (33) | neuropathology_progressive (61), mechanistic_neurobiology (804), non_neurological_or_other (10,251) |
| MDD | depressed_mood (8), anhedonia (27), weight_appetite_change (28), sleep_disturbance (27), psychomotor_change (120), fatigue (25), worthlessness_guilt (8), concentration_indecisiveness (59), suicidality (3) | generic_neurological_abnormality (1,243), mechanistic_neurobiology (1,424), non_neurological_or_other (8,513) |

Each configuration also names a mechanism axis reported apart from face validity (ASD sensorimotor gating, SCZ aberrant salience substrate, NCD progressive neuropathology).

###### Part C. Diagnostic (symptom-count) rule as evaluated

The rule is evaluated on each genotype’s criterion profile after scoring, as a binary indicator, and enters no score. Counts were recomputed here from the catalogue of record and the primary labels.

| Disorder | Rule as evaluated on the profile | Genotypes meeting the rule (curated), all credit | Same, direct credit only | Highest units credited |
| --- | --- | --- | --- | --- |
| ASD | All of A1, A2, A3 and at least two of B1 to B4 | 0 (0) | 0 (0) | 6 of 7 |
| SCZ | At least two of A1 to A5, at least one of them A1, A2 or A3 | 0 (0) | 0 (0) | 2 of 5 |
| IDD | A and B | 333 (26) | 41 (8) | 2 of 2 |
| NCD | A and B | 66 (0) | 8 (0) | 2 of 2 |
| MDD | At least five of A1 to A9, at least one of them A1 or A2 | 0 (0) | 0 (0) | 4 of 9 |

The ASD, SCZ, IDD and NCD rules are those stated in satisfaction_rule of each criterion spec. MDD has no criterion spec, and its rule was evaluated here as five of nine with A1 or A2, the DSM-5-TR count rule. Under any reading the MDD count is zero, because no genotype is credited with more than four units. Only the symptom-count part of each rule is scored. Onset, duration, exclusion and impairment parts are not.

Sources: criterion_specs/{ASD,SCZ,ID,NCD}.json, {asd,scz,intellectualdisability,neurocognitive,mdd}.disorder.json (identical by SHA-256 to FACET_release/configs/), mp_term_taxonomy_criterion_v1_{ASD,SCZ,ID,NCD,MDD}/03_mp_term_taxonomy.tsv, mappability_catalog/genotype_face_validity_seco_criterion_strictonly_prev25_span1_k4_a2floor_ncdB_constructfiltered_idd20260910.tsv (ASD, SCZ, IDD, NCD), mappability_catalog/genotype_face_validity_seco_criterion_strictonly_prev25_span1_k4_a2floor_rescue_MDD_idd20260910.tsv (MDD), audit/geneclust_2026-10-08/labels_corrected_2026-11-02.tsv. Script scripts/s1_rule_check.py.

##### S2 Table. Data files of the primary analysis run with checksums

SHA-256 and size were computed on the device on 2026-11-02 with sha256sum and stat. Dates are file modification times (UTC) recorded on the device, except where a release is declared inside the file. MGI report files carry no release identifier, so for them the modification time is the only available date and is a lower bound on the download date (FACET_REPRODUCE/ONTOLOGY_VERSIONS.md, section 3). All paths are relative to the project root.

###### Part A. Files read by the primary analysis run

| Source | Local file | Version or date | Size (bytes) | SHA-256 |
| --- | --- | --- | --- | --- |
| MGI genotype phenotype report | MGI_PhenoGenoMP.rpt | file date 2026-03-23 | 53,498,021 | 213b36489df36434eedbe764ca082d386a51519601b60f85cdd54981068dbd08 |
| MP term labels and definitions (MGI report) | VOC_MammalianPhenotype.rpt | file date 2026-03-23 | 2,908,361 | 80af8bfdba3e00ac8a128f8a488665f0559fd4c437825cd18332f7b64aedbd83 |
| Mammalian Phenotype Ontology | MPheno_OBO.ontology | declared data-version: releases/2026-05-19/mp.obo (file date 2026-06-01) | 7,600,246 | 49b7ce934ba3885bb546a33a0641ed3ff63519a8fbfa4b5c0794501270b5a805 |
| Human Phenotype Ontology | hp.json | declared release 2026-02-16 (file date 2026-03-23) | 22,063,007 | cb3b756ac11d1308372136bf4f7f8db2ad1c51f64a795e09400e5ffb2a273243 |
| HPO disease annotations (HPO gap analysis) | phenotype.hpoa | declared release 2026-02-16 (file date 2026-05-08) | 35,261,380 | 5d7aedee5f6218e9952fceba5f883df2cc4a0c1938e55d8d2616c15c517964fb |
| Monarch semantic similarity table | phenio-monarch-hp-mp.0.4.semsimian.tsv | version 0.4 (from filename), file date 2024-11-19 | 250,525,100 | aefac634cc3622e1b78981a166ea30bc7ad809eca9b5cefb4ede680be868a13a |
| Monarch similarity table, IC companion (read by compute_b2_baselines.py and run_rescuepool_v2_cells.py) | phenio-monarch-hp-mp.0.4.semsimian.ic.tsv | version 0.4 (from filename), file date 2024-09-16 | 250,500,570 | 3bd29db9eabb81bac153bfe55e7eb9cb4f2010a4f4429604d0be03d2d179182c |
| MGI genotype disease report (labels) | MGI_Geno_DiseaseDO.rpt | file date 2026-04-06 | 8,473,426 | 6ea98fa3aed471aabb410ebcb3418e2bd310ef8251bcb12b50c880f45450fd58 |
| MGI Disease Ontology report (disease names) | MGI_DO.rpt | file date 2026-04-06 | 1,574,869 | fc5a4495e014689c9a005702cda96fbfbc8408e54782b32cd708afd4038f75f9 |
| MGI disease gene model report (disease-gene associations, NCD attention control) | MGI_DiseaseGeneModel.rpt | file date 2026-04-06 | 14,415,178 | 34d3606c38bd8c4911beb61effaf10e4a0a68bb43f516a0dc98c4ac4e56dfa02 |
| MGI marker list (genotype set build) | MRK_List2.rpt.gz | file date 2026-03-29 | 12,488,771 | 48de63a0058bf894a418172771d054b6ae91e53280c2c27d507aeac5787d9454 |
| Disease Ontology (label rules, closure under DOID:1059) | audit/doid.obo | file date 2026-07-27 | 7,162,791 | 079fbbfc6d39f5d6c87b7ad1d2db2e058916584aefdcde0a42156860edae2bbc |
| SFARI Gene animal-model export (genotype set build metadata) | SFARI-Gene_animal-genes_03-22-2026.csv | 2026-03-22 (from filename) | 19,061 | 74bb630bbb6cd08d771093bf313005669fc0719f33d0ded7d568930151e30e1e |

###### Part B. Files read by the secondary analyses (downloaded October 2026)

| Source | Local file | Version or date | Size (bytes) | SHA-256 |
| --- | --- | --- | --- | --- |
| MGI explicit negative disease records | MGI additional/MGI_Geno_NotDiseaseDO.rpt | file date 2026-10-09 | 192,165 | 201c3ce8f3b61d7d9db584d3b73fe4b3e84607f9bf03340aa44f0aac81bf47b4 |
| MGI marker references (publication counts per gene) | MGI additional/MRK_Reference.rpt | file date 2026-10-09 | 31,395,409 | 27a1c9be2560334c0bebf5b051468e467d630051bd95ce4d980094cc702c1b23 |
| MGI IMSR strain report (KOMP report) | MGI additional/MGI_IMSRKomp.rpt | file date 2026-10-09 | 1,297,831 | b37fc8f85de36ee4b6d2fb8035e64781223e0a4ea93759364e91e3b6f2d2e364 |
| MGI phenotypic allele report (allele type and attributes, allele design analysis) | MGI_PhenotypicAllele.rpt.txt | file date 2026-07-31 | 32,189,848 | b48e1110009e2679c7c6d8326491c5f6b2971e6f78ce3fe2787d5353d8ff1516 |
| MGI disease mouse model report | MGI additional/MGI_DiseaseMouseModel.rpt | file date 2026-10-09 | 14,660,624 | 4d752422afae84efc5cbfeb0e734a2946590b0af1336846d97c1c0636cc246d2 |
| IMPC statistical results (latest release folder) | _overnight/impc/statistical-results-ALL.csv.gz | file date 2026-10-09, release number not recorded in the files | 564,423,904 | 476ba65db8300a6ca8e2b3ac9a0d0a9d10094da3fd8ec40c5507d127b504ad93 |
| PubTator3 disease annotations | PubTator3/disease2pubtator3.gz | server date 2026-10-06 (download record of 2026-10-12) | 2,180,457,155 | 64caab45ce5784ea3343b350cd5ac3c72048643f330bce86d5d49f20b08a07b5 |
| PubTator3 gene annotations | PubTator3/gene2pubtator3.gz | server date 2026-10-06 | 757,625,998 | 94a6c301c50f34e6dce37a64e75b09899efced11e4af7d887a8bf530f59543e9 |
| PubTator3 species annotations | PubTator3/species2pubtator3.gz | server date 2026-10-06 | 490,625,738 | efb43fab02e0a4d6fed286d55fdccecda5a0b8fc07f1fa8f24ae6211c8355863 |

###### Part C. Later releases used only for the drift analyses

| Source | Local file | Version or date | Size (bytes) | SHA-256 |
| --- | --- | --- | --- | --- |
| MGI genotype phenotype report | MGI additional/MGI_PhenoGenoMP_2026-10-09.rpt | file date 2026-10-05 | 54,106,348 | 5c7265538a18910bbcde29295bd649626052495379874e5cf8e62892aafec935 |
| Mammalian Phenotype Ontology | MGI additional/MPheno_OBO_2026-10-09.ontology | declared data-version: releases/2026-09-23/mp.obo | 7,703,610 | 93bacc7a9731aec917377cf8d8f95e22b12e61d7665d180ce2d2a17d195728d3 |
| MGI genotype disease report | MGI additional/MGI_Geno_DiseaseDO_2026-10-13.rpt | file date 2026-10-12 | 8,569,474 | d8b6982756db0cf6c664e2673545ec4c66086de9d01608e508dcc76e05da3a2e |
| MGI Disease Ontology report | MGI additional/MGI_DO_2026-10-13.rpt | file date 2026-10-12 | 1,576,974 | 9440ba5bee417dc766063eccbb0b69f20de7c13f4849a72185a4062401897a2e |
| MP labels and definitions (MGI report) | MGI additional/VOC_MammalianPhenotype_2026-10-13.rpt | file date 2026-10-12 | 2,958,052 | abe051251d11b55a7116cfb3d002e33981c2e7bec7c4f379de0837c30cf090f2 |

###### Notes

1. Files of record were identified from broad_benchmark_setup/99_run_config.json (genotype set build: MGI_PhenoGenoMP.rpt, VOC_MammalianPhenotype.rpt, MRK_List2.rpt.gz, MGI_Geno_DiseaseDO.rpt, MGI_DO.rpt, SFARI export). They were also identified from FACET_REPRODUCE/ONTOLOGY_VERSIONS.md (MP, HPO, HPOA and Monarch versions) and from the label scripts audit/geneclust_2026-10-08/relabel.py and label_audit.py (MGI_Geno_DiseaseDO.rpt, MGI_DO.rpt, audit/doid.obo, phenotype.hpoa). Further sources were construct_disorder_tiers.py (MGI_DiseaseGeneModel.rpt), audit/mgi_additional_2026-10-09/mgi_add.py (negative records, marker references, IMSR), audit/overnight_audit_2026-10-09/extract_impc.py (IMPC), audit/open_items_2026-10-13/item1_pubtator_extract.py (PubTator3) and the drift scripts audit/consolidation_2026-10-12/c4_drift.py and audit/overnight_audit_2026-10-09/drift_quant.py.
2. The MGI primary files are not one snapshot. The genotype phenotype report and the MP labels date to 2026-03-23, the disease reports to 2026-04-06 and the marker list to 2026-03-29.
3. The IMPC release number, the phenio knowledge-graph release behind the Monarch table and download dates as opposed to file dates were not recorded and are not given.
4. Code-release versions and checksums named in the Data availability statement do not exist yet (deposit at submission) and are not listed.
5. Not listed: the DSM-5-TR itself, which no step of the run reads (the criteria enter in the project’s paraphrase, deposited with the code release), and the PhenoDigm recomputation inputs other than the Monarch table (the run uses the same table). The equivalent-flavour Monarch tables (phenio-equivalent-hp-mp.0.4.semsimian*.tsv) are on disk but no script in the project root or FACET_release names them, so they are omitted. Whether the IC companion table enters any number in the manuscript was not traced.

##### S3 Table. Label rules and curated models

A genotype is a curated model (positive) for a disorder when its MGI genotype disease annotation (MGI_Geno_DiseaseDO.rpt, disease names joined into disease_terms) satisfies the rule below. Rules were applied to all 76,060 MGI genotypes, and the positives counted here are those inside the 32,535-genotype set (four or more MP annotations).

###### Part A. Rules

| Disorder | Rule as run (name rule on the MGI disease names, case-insensitive) | Corrections applied to reach the primary labels |
| --- | --- | --- |
| ASD | autism spectrum disorder\|autistic disorder\|autism\|autistic features (genotype set build, broad_benchmark_setup/99_run_config.json), genotypes also positive for another psychiatric or neurodevelopmental disorder held apart as “mixed” | Mixed genotypes (annotated to ASD and to another disorder) counted as positives (ruling of 2026-10-08) |
| SCZ | schizophren\|schizoaffective\|schizophreniform\|brief psychotic\|delusional disorder\|psychotic disorder\|schizotypal\|psychosis\|hallucinatory (disorders_rescue_config.json, tier 0), mixed held apart | Mixed genotypes counted as positives |
| IDD | intellectual disability\|mental retardation\|fragile x\|rett syndrome\|cognitive disability\|cognitive impairment\|down syndrome\|williams syndrome\|prader[- ]willi\|angelman (tier 0), mixed held apart | Mixed genotypes counted, disease name containing “intellectual developmental disorder” added, any Disease Ontology term at or below intellectual disability (DOID:1059) in the is_a closure of audit/doid.obo added, Pitt-Hopkins syndrome (DOID:0060488) added by override (2026-10-08) |
| NCD | alzheimer\|frontotemporal dementia\|lewy body\|vascular dementia\|major neurocognitive\|mild neurocognitive\|delirium\|prion disease\|creutzfeldt\|huntington\|parkinson\|traumatic brain injury\|\btbi\b\|cognitive decline\|dementia (tier 0), mixed held apart | Mixed genotypes counted, two Prkag2 genotypes (MGI:8187632, MGI:8187633) annotated to Wolff-Parkinson-White syndrome (DOID:384), matched on “parkinson”, removed and kept as unlabeled background (ruling of 2026-11-02) |
| MDD | depression\|depressive\|melancholic\|dysthymic\|disruptive mood dysregulation\|premenstrual dysphoric\|seasonal affective\|postpartum depress\|atypical depress\|major depressive (tier 0), mixed held apart | Mixed genotypes counted as positives |

The tier 0 positive of construct_disorder_tiers.py is a match of the disorder’s disease_terms_regex on the genotype’s disease names. A tier 0 positive that is also positive for another disorder at the same tier was labeled <disorder>_mixed by build_per_tier_benchmark_tables.py and was outside the positives as run. The doids lists of disorders_rescue_config.json feed the gene-level tiers (MGI disease-gene associations), not the genotype labels.

###### Part B. Resulting positives (primary labels)

Counts from audit/geneclust_2026-10-08/labels_corrected_2026-11-02.tsv (388 rows). “In set” means inside the 32,535-genotype set (in_k4_pool = 1).

| Disorder | Positives as run, in set (outside) | Added: mixed, in set (outside) | Added: name “intellectual developmental disorder” | Added: DO closure under DOID:1059 | Added: Pitt-Hopkins override | Removed: Wolff-Parkinson-White | Positives of record in set | Positives of record, all MGI genotypes |
| --- | --- | --- | --- | --- | --- | --- | --- | --- |
| ASD | 75 (11) | 10 (1) |  |  |  |  | 85 | 97 |
| SCZ | 39 (12) | 3 (0) |  |  |  |  | 42 | 54 |
| IDD | 80 (11) | 7 (0) | 12 | 18 | 4 |  | 121 | 132 |
| NCD | 72 (23) | 1 (0) |  |  |  | 2 | 71 | 94 |
| MDD | 7 (1) | 1 (0) |  |  |  |  | 8 | 9 |
| Total |  |  |  |  |  |  | 327 labels on 317 genotypes | 386 labels |

Positives in the set: 85, 42, 121, 71 and 8 (327 labels). NCD positives as run in the set (72) include the two Wolff-Parkinson-White genotypes, which the file records as label_as_run 1 and label_corrected 0.

Sources: audit/geneclust_2026-10-08/labels_corrected_2026-11-02.tsv, audit/geneclust_2026-10-08/relabel.py (rulings of 2026-10-08, the code that generated the 2026-10-08 file of which the 2026-11-02 file is the WPW-corrected successor), audit/geneclust_2026-10-08/NCD_WPW_REMOVED_2026-11-02.md, disorders_rescue_config.json (disease_terms_regex), broad_benchmark_setup/99_run_config.json (autism_regex), construct_disorder_tiers.py and build_per_tier_benchmark_tables.py (tier 0 and mixed logic). The labels count and the 317 distinct genotypes were recomputed by scripts/s7_composition.py.

##### S4 Table. The 16 primary claims under Holm correction

###### Part A. The Holm family of 16 primary claims

The family was specified after the results were known (Amendment 5 of 2026-10-13) and is reported as a descriptive safeguard. Interval-derived p values are two-sided normal approximations with standard error equal to the width of the 95% gene-clustered percentile interval divided by 3.92. Bootstrap p values of zero are entered as 1/2001. “Below 1e-15” marks p values that underflow to zero in double precision (stored as 0 in the source file).

Values are computed on the primary labels (NCD 71 positives, 67 in the evaluation set).

| Claim | Kind of p | Estimate | p | Holm p | BH q |
| --- | --- | --- | --- | --- | --- |
| Support AUROC above 0.5, ASD | interval-derived | 0.9557 | below 1e-15 | below 1e-15 | below 1e-15 |
| Support AUROC above 0.5, SCZ | interval-derived | 0.6823 | 6.21e-06 | 3.11e-05 | 8.29e-06 |
| Support AUROC above 0.5, IDD | interval-derived | 0.7511 | below 1e-15 | below 1e-15 | below 1e-15 |
| Support AUROC above 0.5, NCD | interval-derived | 0.6679 | 3.86e-06 | 2.32e-05 | 5.62e-06 |
| Coverage AUROC above 0.5, ASD | interval-derived | 0.9641 | below 1e-15 | below 1e-15 | below 1e-15 |
| Coverage AUROC above 0.5, SCZ | interval-derived | 0.8772 | below 1e-15 | below 1e-15 | below 1e-15 |
| Coverage AUROC above 0.5, IDD | interval-derived | 0.7705 | below 1e-15 | below 1e-15 | below 1e-15 |
| Coverage AUROC above 0.5, NCD | interval-derived | 0.7614 | 1.62e-14 | 1.46e-13 | 3.25e-14 |
| FACET minus PhenoDigm behavioral, ASD | bootstrap | +0.2691 | 0.0005 | 0.002 | 0.00062 |
| FACET minus PhenoDigm behavioral, SCZ | bootstrap | +0.0425 | 0.363 | 0.448 | 0.363 |
| FACET minus PhenoDigm behavioral, IDD | bootstrap | +0.0654 | 0.097 | 0.291 | 0.111 |
| FACET minus PhenoDigm behavioral, NCD | bootstrap | +0.0453 | 0.224 | 0.448 | 0.239 |
| Average precision, FACET minus PhenoDigm recomputed, ASD | interval-derived | +0.1536 | 4.28e-08 | 3.0e-07 | 6.85e-08 |
| Papers of top candidates discussing the disorder, candidates minus matched controls, ASD (PubTator3) | two-proportion z | +0.3448 | below 1e-15 | below 1e-15 | below 1e-15 |
| Same, IDD | two-proportion z | +0.2285 | 1.81e-12 | 1.45e-11 | 3.22e-12 |
| Coverage AUROC change under claim-paper removal, ASD | interval-derived, unpaired SE (conservative) | -0.4599 | below 1e-15 | below 1e-15 | below 1e-15 |

Claims surviving Holm at 0.05: 13 of 16 (all except the SCZ, IDD and NCD PhenoDigm behavioral differences).

Sources: audit/open_items_2026-10-13/item5c_abstract_family.json and item5c_abstract_family.py, audit/geneclust_2026-10-08/ncd_wpw_removed_2026-11-02.json and ncd_wpw_removed_phenodigm_ladder_2026-11-02.json (NCD rows). Script scripts/s4_holm_recompute.py.

##### S5 Table. Genes implicated by human genetics without mouse data (H genes)

Genes implicated by human genetic studies that used no mouse data. They define the attention control (genotypes of H genes with no curated model) and the H-gene rows of S14 Table. The lists are those read by audit/geneclust_2026-10-08/bias.py (variable G), whose stored output bias.json records the list sizes 125 (ASD), 49 (SCZ), 934 (IDD), 186 (MDD) and 252 (NCD). Gene counts below are distinct upper-case symbols recomputed from the files. Genes are human symbols, matched to mouse genotypes through the gene key of the allele symbols.

| Disorder | Source list | Selection in the file | Local file | Genes | Union used | File SHA-256 |
| --- | --- | --- | --- | --- | --- | --- |
| ASD | Satterstrom et al. 2020 (Cell), Autism Sequencing Consortium, S2 Table, sheet “102_ASD” | TADA FDR at or below 0.1 (102 genes) | dnv_catalogs/asc_satterstrom2020_asd_genes.tsv (downloaded 2026-05-28) | 102 | 125 (49 genes in both lists) | f8317f4f90e212e8ccb6e1afaddc1339754f83d68ce0b7cfa158b0f070b1c57a |
| ASD | Fu et al. 2022 (Nature Genetics), Supplementary Table 11 | ASD72 flag true | dnv_catalogs/fu2022_asd_genes.tsv (downloaded 2026-05-28) | 72 | (as above) | 7201de50626537cec86d2c317f987c48fdc8bb8b94d822c4af2981432a7c62e0 |
| SCZ | SCHEMA, Singh et al. 2022 (Nature) | Genes with meta-analysis q below 0.05 (10 of them exome-wide significant at p below 2.14e-6) | dnv_catalogs/schema_scz_gene_list.tsv (downloaded 2026-05-26) | 32 | 49 (no overlap) | 88b587033261d960b1b04fc51dbb1206826aea367cbe6719fa94fc20164d59a7 |
| SCZ | denovo-db published de novo variants (McRae compilation), schizophrenia trios of Fromer et al. 2014 (Nature) | Genes with two or more de novo variants (loss of function or missense), per dnv_catalogs/README.md and verified on the counts in the file | dnv_catalogs/denovo_db_scz_genes.tsv (downloaded 2026-05-26) | 17 | (as above) | 0385365d3138046eebcbaecdc067a50bdd52a30fc8764cea073143f7f64d263d |
| IDD | DDG2P, PanelApp version 7.1 | Confidence “Green (confirmed)” | dnv_catalogs/ddg2p_id_confirmed_dominant.tsv (downloaded 2026-05-28) | 934 | 934 | 8971677cc2a31e7f3a8de80e8dad27071a97e3475b8c48c0b5f5ab8ace837118 |
| MDD | PGC MDD 2025 GWAS (no 23andMe, European ancestry), FUMA job 738723, MAGMA gene test | Bonferroni-significant MAGMA genes, MHC region excluded, assigned to a FUMA genomic risk locus by proximity (the “REAL” set, built 2026-05-28) | rescue_line_output/mdd_t1g_REAL_human_genes.tsv | 186 | 186 | b690c9d97d387638ee83c4c7d7daaf28a02616907f8a64ac63616fbea3782e59 |
| NCD | No human-genetics list. MGI disease-gene associations used in its place | Human genes that MGI_DO.rpt associates with the 54 NCD Disease Ontology terms of disorders_rescue_config.json | MGI_DO.rpt (file date 2026-04-06) | 252 | 252 | see S2 Table |

Notes

1. The IDD file name says “dominant”, but its rows include X-linked and biallelic allelic requirements. The selection recorded in the file is the confirmed-confidence tier only.
2. The MDD build summary (rescue_line_output/mdd_t1g_build_summary.json) records 474 Bonferroni-significant MAGMA genes, 419 after MHC exclusion and 186 after the proximity assignment, with the Bonferroni threshold 2.51e-06 over 19,941 genes.
3. Because NCD’s list comes from MGI’s own disease-gene curation, it is not independent of mouse data in the way the other four lists are. Methods states this.
4. The counts of background genotypes carrying an H gene are not tabulated here.

Sources: the files listed, audit/geneclust_2026-10-08/bias.py and bias.json, rescue_line_output/mdd_t1g_build_summary.json, rescue_line_output/RESCUE_SOURCE_AND_JUSTIFICATION_HANDOFF.md, disorders_rescue_config.json. Counts and checksums computed on the device by an inline script (scripts/s5_hgenes.py).

##### S6 Table. Analyses in the order of Results

Every analysis is deterministic given the frozen verdicts of the primary analysis run and the primary labels (audit/geneclust_2026-10-08/labels_corrected_2026-11-02.tsv). Paths are relative to the project root.

| Results subsection | Analysis | Record files |
| --- | --- | --- |
| FACET credits each genotype | Diagnostic Coverage and direct coverage per genotype | mappability_catalog/genotype_face_validity_seco_criterion_strictonly_prev25_span1_k4_a2floor_ncdB_constructfiltered_idd20260910.tsv, built by FACET_release/mappability_catalog/rebuild_fv_seco_criterion.py (depth rule leaf) |
| FACET credits each genotype | Anchor sets and attribution (Table 1, S11 Table) | target_units_{ASD,SCZ,ID,NCD,MDD}/03_target_criterion_units.tsv |
| Behavioral phenotypes are a small and uneven part of the record | Composition of the annotation record and the IMPC bound (S7 Table) | broad_benchmark_setup/01_all_genotype_summary.csv, MPheno_OBO.ontology |
| Curated models recapitulate part of their definition | Unit landscape (Figure 2) | audit/geneclust_2026-10-08/unit_landscape_2026-11-02.json |
| Curated models recapitulate part of their definition | Coverage AUROC and the behaviorally annotated restriction (Table 2) | audit/geneclust_2026-10-08/ncd_wpw_removed_2026-11-02.py and .json, audit/geneclust_2026-10-08/NCD_WPW_REMOVED_2026-11-02.md |
| Curated models recapitulate part of their definition | Route of NCD criterion A credit | catalogue file above with the anchor-to-domain map of target_units_NCD/03_target_criterion_units.tsv |
| Six criteria are credited to no genotype | Criteria without admitted phenotypes and the NCD construct rule (S28 Table) | catalogue file above and mappability_catalog/..._ncdB_asmeasured.tsv (unfiltered configuration) |
| The support score separates curated models from the background | Support AUROC, intervals and average precision (Table 3, S2 Fig) | audit/geneclust_2026-10-08/geneclust_main.json and ncd_wpw_removed_2026-11-02.json (NCD under the primary labels) |
| The support score separates curated models from the background | Robustness rows (S8 Table) | audit/geneclust_2026-10-08/geneclust_supp.json, ncd_wpw_removed_2026-11-02.json and trivial_baselines_ncd_wpw_removed_2026-11-02.py (inclusion threshold) |
| Much of the separation outside ASD reflects phenotyping | Baselines, LLM ablation and depth matching (Table 3) | audit/geneclust_2026-10-08/trivial_baselines_ncd_wpw_removed_2026-11-02.py, llm_ablation_ncd_wpw_removed_2026-11-02.py, depth_matched_intervals_2026-11-02.py and depth_matched_intervals_2026-11-02b.py with their .json outputs |
| FACET exceeds PhenoDigm for ASD | PhenoDigm configurations (Table 3, S10 Table) | audit/geneclust_2026-10-08/ncd_wpw_removed_phenodigm_ladder_2026-11-02.json |
| Alternate LLMs and experts | LLM swap, agreement and admission control (S23 Table) | audit/llm_revalidation_2026-10-19/ (rejudge_record_pairs.py, evalA3.py, RESULTS_AMENDMENT3_AND_NULLMIRROR_2026-10-28.md) and audit/geneclust_2026-10-08/ncd_wpw_removed_judgeswap_2026-11-02.json |
| Alternate LLMs and experts | Upstream re-attribution and reclassification (S12 Table) | audit/llm_revalidation_2026-10-19/ (reassign_units_independent.py, reclassify_mp_criterion_independent.py, PARTB_RESULTS.md, PARTC_RESULTS.md) and _reassign/ |
| Alternate LLMs and experts | Expert agreement (Figure 5, S13 Table) | _rating_analysis_v9.py, audit/llm_revalidation_2026-10-19/PANEL_RULES_VS_RATERS_ABCDEGHL_2026-11-17.md, audit/rater_breakdown_2026-10-20/RATER_BREAKDOWN_ABCDEGHL_2026-11-17.md, audit/rater_single_agreement_2026-11-13/ and FACET_SEND_PACKAGE_2026-09-10/rater_returns/FACET_rater_report_A_B_C_D_E_G_H_L_2026-11-17.md |
| Label agreement traces to the describing paper | Claim-paper removal and relatives (S24 Table) | audit/geneclust_2026-10-08/ncd_wpw_removed_claimpaper_2026-11-02.py and .json, ncd_wpw_removed_relatives_2026-11-02.py and .json |
| Label agreement traces to the describing paper | H genes, attention and the literature (S5 and S14 Tables) | audit/geneclust_2026-10-08/bias.py and bias.json |
| Supporting analyses | Allele design (S9 Table) | audit/geneclust_2026-10-08/landscape.py and landscape.json, recomputed under the primary labels |
| Supporting analyses | Cross-disorder overlap (S1 Fig) | audit/geneclust_2026-10-08/cross.py, recomputed under the primary labels with the ASD top tier of Methods |
| Supporting analyses | Multiplicity (S4 Table) | audit/open_items_2026-10-13/item5c_abstract_family.json and .py, recomputed for the NCD rows under the primary labels |
| Methods | Database drift | rescoring of the frozen verdicts on the MGI report of 2026-10-05 and MP release 2026-09-23 (later-release files of S2 Table) |
| Supporting analyses | Annotation detail and normal-result sensitivity | audit/detail_sensitivity_2026-10-09/ (detail.py, tier_sweep.py) |
| Supporting analyses | Mirrored permutation null | audit/nullmirror_2026-10-27/ |

##### S7 Table. Annotation composition of curated models and of the genotype set

Curated models are the positives of record within the 32,535 genotypes of the genotype set (primary labels, 2026-11-02). Each MP annotation is assigned to the first matching class, in this order. The classes are motor behavior (MP:0002066), sensory behavior and reflexes (MP:0002067), seizures (MP:0002064), social and vocal (MP:0002557, MP:0001529), cognition (MP:0014114), emotion and affect (MP:0002572), other behavior (MP:0004924) and then non-behavioral classes, read through the is_a closure of MP 2026-05-19. Behaviorally annotated means at least one annotation at or below abnormal behavior (MP:0004924). The class-based definition of the source script (any annotation in the seven behavioral classes, seizures included) gives the same counts in every row.

###### Part A. Genotypes carrying each kind of annotation

| Set | Genotypes | Behaviorally annotated, n (percent) | Any social or vocal term, n (percent) | Any cognitive term, n (percent) | Any motor behavior term, n (percent) | Any emotion or affect term, n (percent) | Median MP terms | Median behavioral terms |
| --- | --- | --- | --- | --- | --- | --- | --- | --- |
| ASD curated | 85 | 83 (97.6) | 77 (90.6) | 49 (57.6) | 66 (77.6) | 47 (55.3) | 11 | 6 |
| SCZ curated | 42 | 42 (100) | 18 (42.9) | 31 (73.8) | 32 (76.2) | 23 (54.8) | 9 | 5.5 |
| IDD curated | 121 | 101 (83.5) | 36 (29.8) | 60 (49.6) | 85 (70.2) | 49 (40.5) | 10 | 4 |
| NCD curated | 71 | 62 (87.3) | 2 (2.8) | 22 (31.0) | 54 (76.1) | 10 (14.1) | 10 | 3 |
| MDD curated | 8 | 8 (100) | 5 (62.5) | 4 (50.0) | 2 (25.0) | 8 (100) | 9.5 | 5.5 |
| All genotypes in the set | 32,535 | 8,521 (26.2) | 836 (2.6) | 1,592 (4.9) | 6,383 (19.6) | 1,301 (4.0) | 8 | 0 |

###### Part B. Share of all annotations in each class (percent of the set’s annotations)

| Set | Annotations | Motor behavior | Social and vocal | Cognition | Emotion and affect | Sensory behavior and reflexes | Seizures | Other behavior | All behavioral classes | Nervous system morphology | Nervous system physiology | Ear and eye | Mortality and growth | Other physiology and morphology |
| --- | --- | --- | --- | --- | --- | --- | --- | --- | --- | --- | --- | --- | --- | --- |
| ASD curated | 1,016 | 14.6 | 16.7 | 9.6 | 7.0 | 0.4 | 1.4 | 6.1 | 55.7 | 17.6 | 14.3 | 0.6 | 5.2 | 6.6 |
| SCZ curated | 440 | 16.8 | 6.1 | 15.5 | 8.2 | 0.5 | 0.0 | 3.9 | 50.9 | 19.6 | 14.6 | 0.5 | 4.3 | 10.2 |
| IDD curated | 1,447 | 15.8 | 4.2 | 8.9 | 4.3 | 0.5 | 1.7 | 2.8 | 38.0 | 21.8 | 6.8 | 1.0 | 14.4 | 18.0 |
| NCD curated | 868 | 21.1 | 0.2 | 3.8 | 1.5 | 0.5 | 0.9 | 2.5 | 30.5 | 27.8 | 4.5 | 2.0 | 8.0 | 27.3 |
| MDD curated | 86 | 2.3 | 16.3 | 12.8 | 22.1 | 0.0 | 0.0 | 7.0 | 60.5 | 17.4 | 17.4 | 0.0 | 0.0 | 4.7 |
| All genotypes in the set | 321,517 | 4.2 | 0.4 | 0.8 | 0.5 | 0.1 | 0.3 | 0.9 | 7.3 | 8.1 | 1.9 | 5.3 | 13.4 | 64.0 |

The “all behavioral classes” column is the sum of the seven behavioral class shares as computed from the rounded four-decimal shares, so it can differ from the sum of the one-decimal values shown by 0.1.

The primary labels place 327 disorder labels on 317 distinct genotypes of the set.

Sources: broad_benchmark_setup/01_all_genotype_summary.csv (mp_ids per genotype), MPheno_OBO.ontology (MP 2026-05-19), mappability_catalog/genotype_face_validity_seco_criterion_strictonly_prev25_span1_k4_a2floor_ncdB_constructfiltered_idd20260910.tsv (the 32,535 genotypes), audit/geneclust_2026-10-08/labels_corrected_2026-11-02.tsv. Method replicated from audit/geneclust_2026-10-08/landscape.py (function prof). Script scripts/s7_composition.py.

##### S8 Table. Sensitivity of the support AUROC

Support AUROCs with 95% gene-clustered intervals, except the one-allele row, which gives the median and the 2.5 and 97.5 percentiles over 1,000 draws. The primary values are in main Table 3.

| Comparison | ASD (85 curated) | SCZ (42) | IDD (111) | NCD (67) |
| --- | --- | --- | --- | --- |
| Curated models against the full background (32,535) | 0.969 (0.936 to 0.993) | 0.709 (0.633 to 0.787) | 0.759 (0.715 to 0.833) | 0.731 (0.661 to 0.803) |
| One random allele per gene (median of 1,000 draws) | 0.969 (0.958 to 0.970) | 0.676 (0.661 to 0.704) | 0.791 (0.763 to 0.813) | 0.686 (0.650 to 0.726) |
| Every genotype with at least one annotation (universe 17,472 to 19,432) | 0.950 | 0.661 | 0.742 | 0.670 |
| IDD with Mecp2 removed |  |  | 0.777 (0.717 to 0.835) |  |
| IDD with the 157 unlabeled genotypes of obligate intellectual disability syndromes counted as positives (behaviorally annotated only) |  |  | 0.692 (0.727) |  |
| Retrieval score on the admitted pairs only, without depth matching | 0.960 | 0.590 | 0.747 | 0.672 |

##### S9 Table. Allele design

Each genotype of the 32,535-genotype set is assigned one design class from its alleles (MGI allele type and attributes in MGI_PhenotypicAllele.rpt.txt), by the rule of audit/geneclust_2026-10-08/landscape.py. The classes are tested in order: recombinase driver with a conditional-ready allele (conditional), driver only, humanized sequence, transgene by random insertion, hypomorph, null or knockout, chemically induced, spontaneous, gene trap, other targeted or endonuclease-mediated allele, other. Units credited is Diagnostic Coverage (all credit). Curated models follow the primary labels (2026-11-02).

###### Part A. Curated models by design: n, share with any credited unit, mean units credited

| Design class | Genotypes in set | ASD | SCZ | IDD | NCD | MDD |
| --- | --- | --- | --- | --- | --- | --- |
| Null or knockout | 18,707 | 56, 0.964, 3.68 | 29, 0.793, 1.41 | 63, 0.619, 0.87 | 29, 0.517, 0.52 | 3, 1.0, 2.67 |
| Conditional (Cre plus floxed allele) | 4,313 | 0 | 0 | 0 | 0 | 0 |
| Humanized sequence | 1,848 | 10, 1.0, 3.30 | 2, 1.0, 1.50 | 31, 0.677, 0.84 | 26, 0.692, 0.69 | 3, 1.0, 2.00 |
| Targeted, not null (knock-in, point, other) | 1,682 | 11, 1.0, 3.18 | 1, 1.0, 2.00 | 20, 0.450, 0.60 | 11, 0.909, 0.91 | 0 |
| Chemically induced (ENU) | 1,690 | 1, 1.0, 5.00 | 4, 1.0, 1.25 | 0 | 0 | 1, 1.0, 2.00 |
| Spontaneous | 1,589 | 1, 1.0, 3.00 | 2, 1.0, 1.50 | 0 | 4, 0.75, 0.75 | 0 |
| Transgene (random insertion) | 1,434 | 1, 1.0, 3.00 | 1, 1.0, 1.00 | 0 | 1, 1.0, 1.00 | 0 |
| Hypomorph | 768 | 5, 0.6, 2.20 | 3, 1.0, 2.00 | 6, 0.833, 1.17 | 0 | 0 |
| Other or unresolved | 383 | 0 | 0 | 0 | 0 | 1, 1.0, 2.00 |
| Driver or reporter only | 79 | 0 | 0 | 1, 0.0, 0.00 | 0 | 0 |
| Gene trap | 42 | 0 | 0 | 0 | 0 | 0 |
| Total curated | 32,535 | 85 | 42 | 121 | 71 | 8 |

###### Part B. All genotypes of each design: share with any credited unit (crude, and directly standardized to the set’s distribution of behavioral annotation counts 0, 1 to 2, 3 to 5, 6 or more)

| Design class | ASD | SCZ | IDD | NCD | MDD |
| --- | --- | --- | --- | --- | --- |
| Null or knockout | 0.109 (0.110) | 0.171 (0.170) | 0.086 (0.087) | 0.138 (0.138) | 0.108 (0.107) |
| Conditional (Cre plus floxed allele) | 0.050 (0.079) | 0.113 (0.170) | 0.056 (0.087) | 0.099 (0.149) | 0.077 (0.109) |
| Humanized sequence | 0.172 (0.090) | 0.320 (0.190) | 0.154 (0.084) | 0.278 (0.161) | 0.142 (0.085) |
| Targeted, not null | 0.086 (0.086) | 0.161 (0.167) | 0.083 (0.082) | 0.128 (0.132) | 0.104 (0.112) |
| Chemically induced (ENU) | 0.112 (0.143) | 0.146 (0.185) | 0.051 (0.064) | 0.108 (0.137) | 0.084 (0.103) |
| Spontaneous | 0.162 (0.113) | 0.284 (0.198) | 0.121 (0.084) | 0.176 (0.120) | 0.141 (0.104) |
| Transgene (random insertion) | 0.082 (0.086) | 0.177 (0.187) | 0.079 (0.083) | 0.126 (0.132) | 0.114 (0.117) |
| Hypomorph | 0.089 (0.096) | 0.150 (0.175) | 0.083 (0.094) | 0.118 (0.135) | 0.095 (0.103) |
| Other or unresolved | 0.099 (0.136) | 0.123 (0.173) | 0.044 (0.065) | 0.073 (0.105) | 0.097 (0.127) |
| Driver or reporter only | 0.051 (0.061) | 0.076 (0.101) | 0.063 (0.072) | 0.127 (0.153) | 0.063 (0.098) |
| Gene trap | 0.000 (0.000) | 0.119 (0.144) | 0.048 (0.065) | 0.119 (0.158) | 0.048 (0.061) |

Standardized shares are label-independent and are read from landscape.json.

###### Part C. Conditional deletions against germline nulls of the same gene (mean units credited per gene, Wilcoxon signed-rank test over genes)

| Disorder | All genotypes: genes, conditional mean, null mean, genes conditional higher, genes null higher, p | Both behaviorally annotated: genes, conditional mean, null mean, conditional higher, null higher, p |
| --- | --- | --- |
| ASD | 1,322, 0.099, 0.151, 98, 205, 1.0e-06 | 203, 0.562, 0.822, 45, 81, 0.00026 |
| SCZ | 1,322, 0.143, 0.160, 203, 257, 0.091 | 203, 0.791, 0.755, 76, 66, 0.50 |
| IDD | 1,322, 0.072, 0.072, 122, 138, 0.91 | 203, 0.347, 0.400, 50, 56, 0.17 |
| NCD | 1,322, 0.108, 0.116, 178, 224, 0.25 | 203, 0.547, 0.575, 56, 59, 0.39 |
| MDD | 1,322, 0.092, 0.100, 168, 212, 0.093 | 203, 0.420, 0.432, 61, 62, 0.72 |

Part C does not depend on labels. No curated model of any disorder is a conditional deletion.

Sources: audit/geneclust_2026-10-08/landscape.py and landscape.json (sections design, conditional_vs_null_within_gene, conditional_vs_null_within_gene_behavior_tested), MGI_PhenotypicAllele.rpt.txt, broad_benchmark_setup/01_all_genotype_summary.csv, the catalogue primary files, audit/geneclust_2026-10-08/labels_corrected_2026-11-02.tsv, audit/geneclust_2026-10-08/FINDINGS_LOG_2026-10-08.md (findings F14, F17, F44). Script scripts/s9_allele_design.py.

##### S10 Table. FACET against PhenoDigm in four configurations

Discrimination in the genotypes carrying a candidate term. Published uses the Monarch table as distributed, which omits term pairs below a similarity cutoff. Recomputed (the main-text configuration, Table 3) applies PhenoDigm to the full phenotype profile without that cutoff. Behavioral restricts PhenoDigm to behavioral phenotypes and is the configuration in the Holm family. Admitted pairs applies the PhenoDigm scorer to the pairs FACET admitted. Differences are FACET minus PhenoDigm with 95% gene-clustered intervals and the two-sided bootstrap p.

| Disorder | FACET | PhenoDigm published | PhenoDigm recomputed | PhenoDigm behavioral | PhenoDigm on FACET’s admitted pairs | Difference, published | Difference, recomputed | Difference, behavioral | Difference, admitted pairs |
| --- | --- | --- | --- | --- | --- | --- | --- | --- | --- |
| ASD | 0.956 | 0.656 | 0.772 | 0.687 | 0.923 | +0.300 (0.226 to 0.382, p below 0.001) | +0.184 (0.126 to 0.248, p below 0.001) | +0.269 (0.236 to 0.301, p below 0.001, Holm p 0.002) | +0.033 (0.025 to 0.041, p below 0.001) |
| SCZ | 0.682 | 0.725 | 0.772 | 0.640 | 0.744 | -0.043 (-0.143 to 0.046, p 0.34) | -0.090 (-0.191 to 0.001, p 0.054) | +0.043 (-0.053 to 0.135, p 0.36, Holm p 0.45) | -0.062 (-0.144 to 0.016, p 0.10) |
| IDD | 0.751 | 0.616 | 0.706 | 0.686 | 0.732 | +0.135 (0.068 to 0.240, p below 0.001) | +0.045 (-0.028 to 0.158, p 0.28) | +0.065 (-0.012 to 0.171, p 0.10, Holm p 0.29) | +0.019 (0.003 to 0.029, p 0.024) |
| NCD | 0.668 | 0.557 | 0.593 | 0.623 | 0.668 | +0.111 (0.013 to 0.203, p 0.026) | +0.075 (0.007 to 0.150, p 0.031) | +0.045 (-0.028 to 0.126, p 0.22, Holm p 0.45) | 0.000 (-0.025 to 0.023, p 0.98) |

##### S11 Table. The 138 attributable HPO anchors

The HPO anchors that the anchor-attribution step of the primary analysis run assigned to a criterion unit (column attributable = True). Units are the scored criterion units of Methods. Category and centrality are the values returned by the decomposition step. Second unit is the secondary unit the attribution step named, if any (crediting uses the primary unit only). HPO labels are the HPO 2026-02-16 labels carried in the files. No DSM-5-TR text is reproduced, and the free-text reason column of the source files is not included.

Source: target_units_{ASD,SCZ,ID,NCD,MDD}/03_target_criterion_units.tsv.

Total 138 attributable anchors of 174. Five of the 15 SCZ attributable anchors are attributed to criterion B, which is not scored, so 133 lie on scored units.

###### ASD

Count: 37 attributable of 43 anchors (37 on scored units).

| HPO id | HPO label | Criterion unit | Second unit | Category | Centrality |
| --- | --- | --- | --- | --- | --- |
| HP:0012433 | Abnormal social behavior | ASD A1 |  | social communication | core |
| HP:0012760 | Reduced social responsiveness | ASD A1 |  | social communication | core |
| HP:4000084 | Reduced sharing of interests | ASD A1 |  | social communication | core |
| HP:4000092 | Poor conversational reciprocity | ASD A1 |  | social communication | core |
| HP:5200027 | Infrequent social initiations | ASD A1 |  | social communication | supporting |
| HP:5200028 | Reduced responsiveness to social cues | ASD A1 |  | social communication | supporting |
| HP:5200121 | Objectification of others | ASD A1 | A3 | social communication | supporting |
| HP:5200310 | Diminishment of social interactions | ASD A1 | A3 | social communication | supporting |
| HP:5200316 | Diminishment of interpersonal communication | ASD A1 |  | social communication | core |
| HP:0000758 | Abnormal nonverbal communicative behavior | ASD A2 |  | social communication | core |
| HP:0004673 | Decreased facial expression | ASD A2 |  | social communication | core |
| HP:0034435 | Abnormal eye contact | ASD A2 |  | social communication | core |
| HP:4000081 | Reduced production of gestures | ASD A2 |  | social communication | core |
| HP:5200030 | Diminished integration of verbal and non-verbal communicative behavior | ASD A2 |  | social communication | core |
| HP:0000728 | Reduced ability to form peer relationships | ASD A3 |  | social communication | core |
| HP:4000083 | Lack of interest in peers | ASD A3 |  | social communication | core |
| HP:5200016 | Abnormal peer relationships | ASD A3 |  | social communication | supporting |
| HP:5200035 | Reduced cooperative imaginative play | ASD A3 |  | social communication | core |
| HP:5200136 | Reduced social reciprocity | ASD A3 | A1 | social communication | core |
| HP:0000733 | Motor stereotypy | ASD B1 |  | restricted repetitive behavior | core |
| HP:0010529 | Echolalia | ASD B1 |  | language | core |
| HP:0025774 | Idiosyncratic use of words | ASD B1 |  | language | core |
| HP:5200108 | Nonfunctional or atypical use of objects in play | ASD B1 |  | restricted repetitive behavior | core |
| HP:0000732 | Inflexible adherence to routines | ASD B2 |  | restricted repetitive behavior | core |
| HP:0033360 | Impaired ability to shift attention | ASD B2 |  | restricted repetitive behavior | supporting |
| HP:5200052 | Resistance to trivial environmental changes | ASD B2 |  | restricted repetitive behavior | core |
| HP:5200129 | Abnormal rituals | ASD B2 |  | restricted repetitive behavior | core |
| HP:5200130 | Insistence on sameness | ASD B2 |  | restricted repetitive behavior | core |
| HP:4000070 | Fixated interests | ASD B3 |  | restricted repetitive behavior | core |
| HP:0007328 | Impaired pain sensation | ASD B4 |  | sensory | core |
| HP:0010829 | Impaired temperature sensation | ASD B4 |  | sensory | core |
| HP:4000079 | Sensory seeking | ASD B4 |  | sensory | core |
| HP:5200058 | Sensory hypersensitivity | ASD B4 |  | sensory | core |
| HP:5200059 | Sensory hyposensitivity unexplained by sensory deficit | ASD B4 |  | sensory | core |
| HP:5200064 | Olfactory sensory seeking | ASD B4 |  | sensory | core |
| HP:5200067 | Visual sensory seeking | ASD B4 |  | sensory | core |
| HP:5200132 | Atypical sensory examination of objects | ASD B4 |  | sensory | core |

###### SCZ

Count: 15 attributable of 41 anchors (10 on scored units, 5 on criterion B, which is not scored).

| HPO id | HPO label | Criterion unit | Second unit | Category | Centrality |
| --- | --- | --- | --- | --- | --- |
| HP:0000746 | Delusion | SCZ A1 |  | delusions | core |
| HP:0000738 | Hallucinations | SCZ A2 |  | hallucinations | core |
| HP:0025769 | Disordered formal thought process | SCZ A3 |  | disorganized speech | core |
| HP:0000708 | Atypical behavior | SCZ A4 |  | catatonic disorganized behavior | supporting |
| HP:0000718 | Aggressive behavior | SCZ A4 |  | other | supporting |
| HP:0002311 | Incoordination | SCZ A4 |  | other | supporting |
| HP:5200045 | Reduced impulse control | SCZ A4 |  | other | supporting |
| HP:0000745 | Abnormal diminished volition | SCZ A5 | B | negative symptoms | core |
| HP:0025766 | Abnormal affect | SCZ A5 |  | other | supporting |
| HP:5200328 | Diminishment of emotional expression | SCZ A5 |  | negative symptoms | core |
| HP:0007086 | Social and occupational deterioration | SCZ B (not scored) |  | other | supporting |
| HP:0031058 | Impairment of activities of daily living | SCZ B (not scored) |  | other | supporting |
| HP:0033695 | Occupational disability | SCZ B (not scored) |  | other | supporting |
| HP:5200024 | Abnormal relationship | SCZ B (not scored) | A5 | other | supporting |
| HP:5200310 | Diminishment of social interactions | SCZ B (not scored) | A5 | negative symptoms | supporting |

###### IDD

Count: 21 attributable of 25 anchors (21 on scored units).

| HPO id | HPO label | Criterion unit | Second unit | Category | Centrality |
| --- | --- | --- | --- | --- | --- |
| HP:0001328 | Specific learning disability | IDD A |  | intellectual functioning | core |
| HP:0002354 | Memory impairment | IDD A |  | adaptive functioning | supporting |
| HP:0002442 | Dyscalculia | IDD A |  | adaptive functioning | supporting |
| HP:0010522 | Dyslexia | IDD A |  | adaptive functioning | supporting |
| HP:0010526 | Dysgraphia | IDD A |  | adaptive functioning | supporting |
| HP:0033051 | Impaired executive functioning | IDD A |  | intellectual functioning | core |
| HP:0033055 | Impaired ability to plan | IDD A |  | intellectual functioning | core |
| HP:0033056 | Impaired ability to organize | IDD A |  | adaptive functioning | supporting |
| HP:0100543 | Cognitive impairment | IDD A |  | intellectual functioning | core |
| HP:5200401 | Abnormal judgment | IDD A |  | intellectual functioning | core |
| HP:0000728 | Reduced ability to form peer relationships | IDD B |  | adaptive functioning | supporting |
| HP:0002463 | Language impairment | IDD B |  | adaptive functioning | supporting |
| HP:0007086 | Social and occupational deterioration | IDD B |  | adaptive functioning | supporting |
| HP:0031058 | Impairment of activities of daily living | IDD B |  | adaptive functioning | core |
| HP:0034434 | Abnormal communication | IDD B |  | adaptive functioning | core |
| HP:5200021 | Reduced social insight | IDD B |  | adaptive functioning | supporting |
| HP:5200024 | Abnormal relationship | IDD B |  | adaptive functioning | supporting |
| HP:5200037 | Lack of expressed empathy | IDD B |  | adaptive functioning | supporting |
| HP:5200118 | Difficulty recognizing another’s emotions | IDD B |  | adaptive functioning | supporting |
| HP:5200310 | Diminishment of social interactions | IDD B |  | adaptive functioning | core |
| HP:5200316 | Diminishment of interpersonal communication | IDD B |  | adaptive functioning | supporting |

###### NCD

Count: 28 attributable of 28 anchors (28 on scored units). The criterion B anchors are those of the rebuild of 2026-07-31 (the file carries a backup suffixed .bak_pre_ncdB_2026-07-31, not used).

| HPO id | HPO label | Criterion unit | Second unit | Category | Centrality |
| --- | --- | --- | --- | --- | --- |
| HP:0001268 | Mental deterioration | NCD A |  | cognitive executive | core |
| HP:0002186 | Apraxia | NCD A |  | motor coordination | supporting |
| HP:0002354 | Memory impairment | NCD A |  | cognitive executive | core |
| HP:0002463 | Language impairment | NCD A |  | other | core |
| HP:0006977 | Deficit in grammar | NCD A |  | other | supporting |
| HP:0010524 | Disturbed sensory perception | NCD A |  | motor coordination | supporting |
| HP:0010794 | Impaired visuospatial constructive cognition | NCD A |  | motor coordination | supporting |
| HP:0010863 | Receptive language delay | NCD A |  | other | supporting |
| HP:0030784 | Anomic aphasia | NCD A |  | other | supporting |
| HP:0033051 | Impaired executive functioning | NCD A |  | cognitive executive | core |
| HP:0033055 | Impaired ability to plan | NCD A |  | cognitive executive | supporting |
| HP:0033071 | Impaired task monitoring | NCD A |  | cognitive executive | supporting |
| HP:0033692 | Declarative memory loss | NCD A |  | cognitive executive | supporting |
| HP:0100543 | Cognitive impairment | NCD A |  | cognitive executive | core |
| HP:5200021 | Reduced social insight | NCD A |  | social cognition empathy | core |
| HP:5200044 | Reduced attention regulation | NCD A |  | cognitive executive | supporting |
| HP:5200118 | Difficulty recognizing another’s emotions | NCD A |  | social cognition empathy | supporting |
| HP:0000020 | Urinary incontinence | NCD B |  | adaptive functioning | supporting |
| HP:0002607 | Bowel incontinence | NCD B |  | adaptive functioning | supporting |
| HP:0010992 | Stress urinary incontinence | NCD B |  | adaptive functioning | supporting |
| HP:0031058 | Impairment of activities of daily living | NCD B |  | adaptive functioning | core |
| HP:0031059 | Impaired ability to bathe oneself | NCD B |  | adaptive functioning | supporting |
| HP:0031060 | Impaired ability to dress oneself | NCD B |  | adaptive functioning | supporting |
| HP:0031061 | Impaired toileting ability | NCD B |  | adaptive functioning | supporting |
| HP:0031062 | Impaired transferring ability | NCD B |  | adaptive functioning | supporting |
| HP:0031063 | Impaired feeding ability | NCD B |  | adaptive functioning | supporting |
| HP:0031064 | Impaired continence | NCD B |  | adaptive functioning | supporting |
| HP:0040183 | Encopresis | NCD B |  | adaptive functioning | supporting |

###### MDD

Count: 37 attributable of 37 anchors (37 on scored units). MDD attribution was carried over from the June 2026 MDD run (the file records “carried_from_mdd_v1”, adapter of 2026-07-29), not produced by a new attribution call.

| HPO id | HPO label | Criterion unit | Second unit | Category | Centrality |
| --- | --- | --- | --- | --- | --- |
| HP:0000737 | Irritability | MDD A1 |  | depressed mood | core |
| HP:0033705 | Tearfulness | MDD A1 |  | depressed mood | supporting |
| HP:5200271 | Hopelessness | MDD A1 |  | depressed mood | supporting |
| HP:5200273 | Pathological sadness | MDD A1 |  | depressed mood | core |
| HP:0000741 | Apathy | MDD A2 |  | anhedonia | supporting |
| HP:0012154 | Anhedonia | MDD A2 |  | anhedonia | core |
| HP:0046504 | Decreased libido | MDD A2 |  | anhedonia | supporting |
| HP:4000068 | Abnormal interest | MDD A2 |  | anhedonia | supporting |
| HP:0001508 | Failure to thrive | MDD A3 |  | weight appetite change | supporting |
| HP:0001824 | Weight loss | MDD A3 |  | weight appetite change | core |
| HP:0002591 | Polyphagia | MDD A3 |  | weight appetite change | core |
| HP:0004324 | Increased body weight | MDD A3 |  | weight appetite change | core |
| HP:0004396 | Poor appetite | MDD A3 |  | weight appetite change | core |
| HP:0030221 | Sweet craving | MDD A3 |  | weight appetite change | supporting |
| HP:0025288 | Excessive sleep | MDD A4 |  | sleep disturbance | core |
| HP:0025298 | Extended nighttime sleep | MDD A4 |  | sleep disturbance | supporting |
| HP:0031354 | Sleep onset insomnia | MDD A4 |  | sleep disturbance | supporting |
| HP:0031355 | Maintenance insomnia | MDD A4 |  | sleep disturbance | supporting |
| HP:0031356 | Terminal insomnia | MDD A4 |  | sleep disturbance | supporting |
| HP:0100785 | Insomnia | MDD A4 |  | sleep disturbance | core |
| HP:0000711 | Restlessness | MDD A5 |  | psychomotor change | supporting |
| HP:0000713 | Agitation | MDD A5 |  | psychomotor change | core |
| HP:0000733 | Motor stereotypy | MDD A5 |  | psychomotor change | supporting |
| HP:0002300 | Mutism | MDD A5 |  | psychomotor change | supporting |
| HP:0002465 | Poor speech | MDD A5 |  | psychomotor change | supporting |
| HP:0012171 | Stereotypical hand wringing | MDD A5 |  | psychomotor change | supporting |
| HP:0031843 | Abnormally slow thought process | MDD A5 |  | psychomotor change | supporting |
| HP:0012378 | Fatigue | MDD A6 |  | fatigue | supporting |
| HP:0025406 | Asthenia | MDD A6 |  | fatigue | core |
| HP:0031058 | Impairment of activities of daily living | MDD A6 |  | fatigue | supporting |
| HP:0025791 | Delusion of guilt | MDD A7 |  | worthlessness guilt | supporting |
| HP:0031469 | Low self-esteem | MDD A7 |  | worthlessness guilt | core |
| HP:6000011 | Guilt | MDD A7 |  | worthlessness guilt | supporting |
| HP:0000736 | Short attention span | MDD A8 |  | concentration indecisiveness | supporting |
| HP:0031987 | Diminished ability to concentrate | MDD A8 |  | concentration indecisiveness | core |
| HP:0031589 | Suicidal ideation | MDD A9 |  | suicidality | core |
| HP:5200330 | Suicide behaviors | MDD A9 |  | suicidality | core |

Totals: 174 anchors, 138 attributable, 133 attributable to scored units.

##### S12 Table. Stability of the upstream LLM steps

Re-attribution reports agreement with the primary analysis run on the primary criterion unit. Reclassification reports the percent of the MP terms the primary analysis run assigns to a criterion that the alternate LLM also assigns and, in parentheses, the percent of unassigned terms it also leaves unassigned.

| Check | ASD | SCZ | IDD | NCD |
| --- | --- | --- | --- | --- |
| Anchor re-attribution, claude and gemini | 42 and 40 of 42 (of the 43 ASD primary anchors, one was absent from both alternate resolutions) | 31 and 30 of 41 | 24 and 24 of 25 | 27 and 28 of 28 |
| Taxonomy reclassification, assigned terms kept (unassigned kept out): claude, gemini | 84 (99), 93 (91) | 50 (99), 89 (93) | 70 (100), 100 (95) | 76 (99), 98 (92) |

##### S13 Table. Expert agreement

Rates are estimated over distinct pairs. The lower bound is one-sided 95% on the rater-adjusted standard error and is the quantity the pre-specified rule reads. The criterion-match rows read the membership answer against the LLM’s criterion mapping in place of its admission verdict, so their denominators differ from the composite rows. The primary analysis is interim and uses all eight raters returned so far (seven invited raters and the first author, who rated block A). The last two rows remove the first author’s ratings.

| Comparison | Pairs (ratings) | Rate | Wilson 95% interval | One-sided lower bound |
| --- | --- | --- | --- | --- |
| Composite confirmation of admitted pairs | 77 (138) | 0.662 | 0.551 to 0.758 | 0.517 |
| Composite rejection of refused pairs | 75 (130) | 0.884 | 0.793 to 0.939 | 0.824 |
| Criterion match alone, pairs the LLM mapped to the criterion | 74 (132) | 0.788 | 0.682 to 0.866 | 0.711 |
| Criterion match alone, pairs the LLM did not ground to the criterion | 78 (136) | 0.733 | 0.625 to 0.818 | 0.657 |
| Seven invited raters only: composite confirmation | 77 (118) | 0.654 | 0.542 to 0.750 | 0.492 |
| Seven invited raters only: composite rejection | 75 (111) | 0.880 | 0.787 to 0.936 | 0.815 |

##### S14 Table. Phenotyping attention and the describing literature

H genes are genes implicated by human genetics without mouse data (S5 Table). Odds ratios are adjusted for the logarithm of one plus the PubMed record count per gene, with Wald 95% intervals. Literature rates compare the top uncurated candidates with controls matched on behavioral depth and annotation source, by whether PubTator3 tags a describing paper with the disorder.

| Quantity | ASD | SCZ | IDD | NCD | MDD |
| --- | --- | --- | --- | --- | --- |
| H-gene genotypes behaviorally annotated against other background genotypes, percent (odds ratio adjusted for publication count) | 35.4 against 25.8 (1.83, 1.57 to 2.13) | 28.0 against 26.1 (1.08, 0.69 to 1.67) | 25.4 against 26.1 (1.13, 1.06 to 1.21) | 38.4 against 25.2 (2.43, 2.20 to 2.68) | 35.1 against 26.0 (1.68, 1.39 to 2.04) |
| Top uncurated candidates whose papers discuss the disorder, against matched controls, percent (risk ratio) | 40.2 against 5.7 (7.0, 4.2 to 11.7) | 14.2 against 2.8 | 34.4 against 11.6 (3.0, 2.1 to 4.1) | 1.0 against 14.0 |  |

#### Supplementary Figure

##### S1 Fig. Cross-disorder overlap

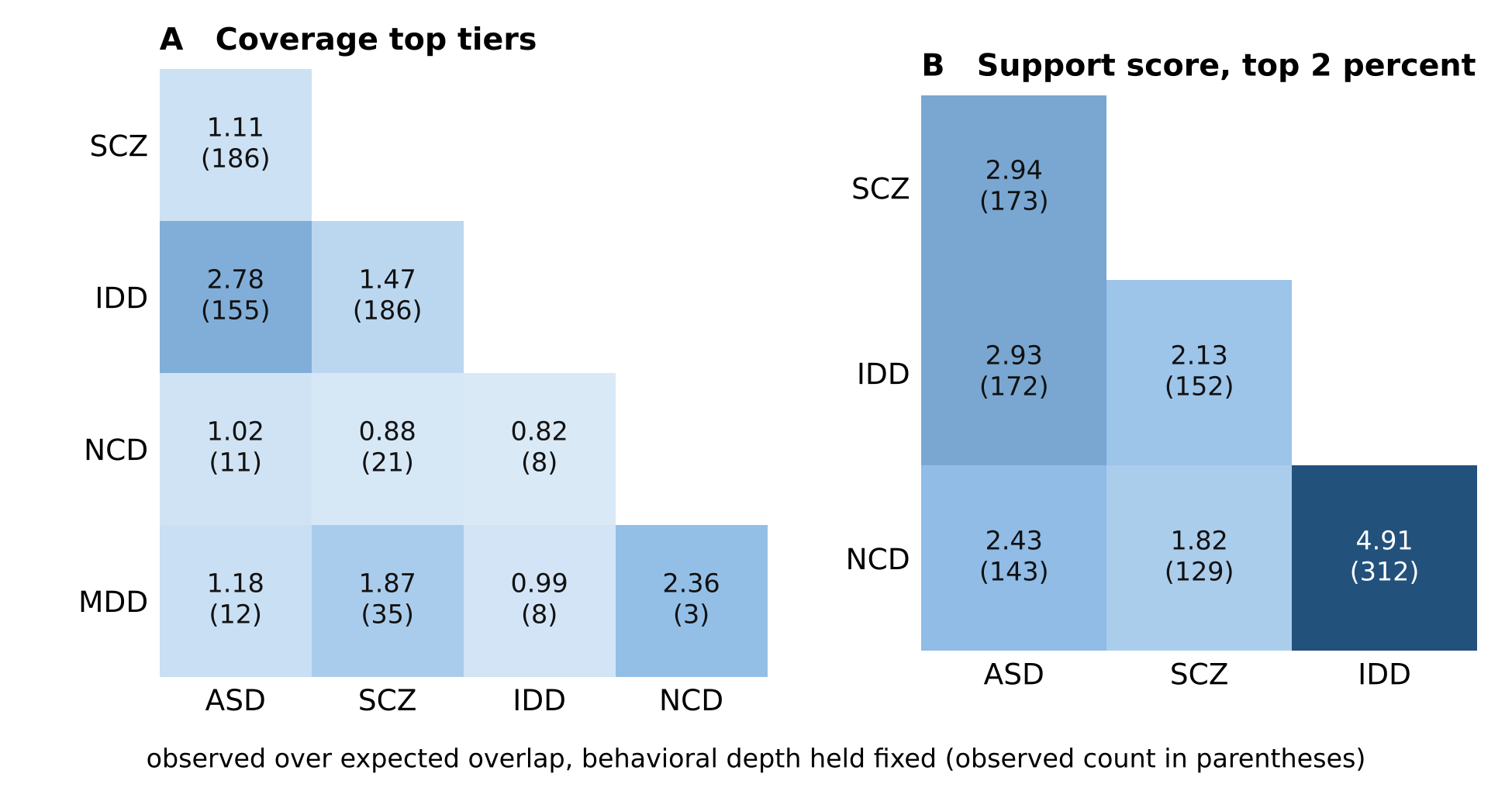

*S1 Fig. Cross-disorder overlap of uncurated genotypes. Each cell gives the observed overlap of two disorders’ top sets over the overlap expected with behavioral depth held fixed (five bins of the behavioral count), with the observed count in parentheses. Curated models of either disorder are excluded. (A) Coverage top tiers: Diagnostic Coverage of three or more units for ASD and MDD and two for SCZ, IDD and NCD. (B) Support score at or above the 98th percentile of the genotype set (primary disorders only). Labels of record.*

Against independence the top sets of every pair overlap 8 to 46 times more than chance, but most of that excess follows behavioral phenotyping. With behavioral depth held fixed, coverage overlaps run from 0.82 to 2.78 times expectation, the highest for ASD and IDD. Support overlaps run from 1.82 to 4.91, the highest for IDD and NCD. The values are computed by the logic of audit/geneclust_2026-10-08/cross.py under the primary labels, with the ASD top tier set to three units as in Methods.

| Pair | Coverage top sets (first, second) | Overlap | Expected, independent | Expected, depth fixed | Ratio, depth fixed | Support top sets (first, second) | Overlap | Expected, depth fixed | Ratio, depth fixed |
| --- | --- | --- | --- | --- | --- | --- | --- | --- | --- |
| ASD and SCZ | 429, 1,181 | 186 | 15.6 | 167.7 | 1.11 | 560, 578 | 173 | 58.9 | 2.94 |
| ASD and IDD | 417, 264 | 155 | 3.4 | 55.7 | 2.78 | 546, 548 | 172 | 58.7 | 2.93 |
| ASD and NCD | 437, 63 | 11 | 0.9 | 10.8 | 1.02 | 573, 652 | 143 | 58.8 | 2.43 |
| ASD and MDD | 439, 46 | 12 | 0.6 | 10.2 | 1.18 |  |  |  |  |
| SCZ and IDD | 1,191, 292 | 186 | 10.7 | 127.0 | 1.47 | 604, 592 | 152 | 71.4 | 2.13 |
| SCZ and NCD | 1,215, 66 | 21 | 2.5 | 23.8 | 0.88 | 638, 681 | 129 | 70.8 | 1.82 |
| SCZ and MDD | 1,228, 43 | 35 | 1.6 | 18.7 | 1.87 |  |  |  |  |
| IDD and NCD | 306, 66 | 8 | 0.6 | 9.8 | 0.82 | 600, 644 | 312 | 63.6 | 4.91 |
| IDD and MDD | 304, 46 | 8 | 0.4 | 8.1 | 0.99 |  |  |  |  |
| NCD and MDD | 65, 46 | 3 | 0.1 | 1.3 | 2.36 |  |  |  |  |

Curated models of one disorder in the coverage top tier of another (models not curated for the second disorder, expected rate from genotypes curated for neither within each behavioral-depth bin):

| Curated for | Top tier of | Models | Observed | Expected | Ratio |
| --- | --- | --- | --- | --- | --- |
| ASD | SCZ | 84 | 50 | 35.0 | 1.43 |
| ASD | IDD | 78 | 43 | 12.0 | 3.59 |
| ASD | NCD | 85 | 3 | 2.4 | 1.23 |
| ASD | MDD | 85 | 0 | 2.3 | 0.00 |
| SCZ | ASD | 41 | 12 | 6.1 | 1.97 |
| SCZ | IDD | 41 | 15 | 4.5 | 3.31 |
| SCZ | NCD | 42 | 0 | 0.8 | 0.00 |
| SCZ | MDD | 42 | 3 | 0.7 | 4.50 |
| IDD | ASD | 114 | 24 | 14.3 | 1.67 |
| IDD | SCZ | 120 | 40 | 34.2 | 1.17 |
| IDD | NCD | 121 | 0 | 2.4 | 0.00 |
| IDD | MDD | 121 | 0 | 1.9 | 0.00 |
| NCD | ASD | 71 | 4 | 6.6 | 0.61 |
| NCD | SCZ | 71 | 16 | 15.4 | 1.04 |
| NCD | IDD | 71 | 1 | 4.9 | 0.20 |
| NCD | MDD | 70 | 0 | 0.7 | 0.00 |
| MDD | ASD | 8 | 2 | 1.9 | 1.08 |
| MDD | SCZ | 8 | 3 | 3.3 | 0.91 |
| MDD | IDD | 8 | 3 | 1.5 | 1.96 |
| MDD | NCD | 7 | 1 | 0.2 | 4.44 |

#### Supplementary Results, passages moved from the main text (v18)

##### SR-M1. Behavioral phenotypes are a small and uneven part of the MGI record

About 7% of all annotated terms in the genotype set describe behavior (motor 4.2%, cognition 0.8, emotion 0.5, social or vocal 0.4 and other behavioral terms the remainder, S1 Text). Three in four genotypes (73.8%) carry no “abnormal behavior” annotation. The MGI phenotype report carries abnormal results only. At least 7.5% of genotypes without behavioral annotation were assayed and found normal: 1,795 of 24,014 completed a behavioral procedure of the International Mouse Phenotyping Consortium (IMPC) with no abnormal call. Absence of behavioral annotation therefore cannot be read as absence of testing. Curated models are behaviorally annotated at 83.5 to 100% by disorder against 26.2% of the genotype set (Table 1), and their terms reflect the assays each field favors. Nine in ten curated ASD models (90.6%) carry a social term. Only half of curated IDD models (49.6%) and under a third of curated NCD models (22 of 71, 31.0%) carry any cognitive term (S7 Table).

Each MGI disease label records that a cited paper described the genotype as a model of the disease. The curation places 327 disorder labels on the genotype set under the label rules of Methods and records almost no negatives (ASD 1, NCD 12, MDD 1, SCZ and IDD none). It leaves 157 genotypes from 57 syndromes with obligate intellectual disability without an IDD label. Every uncurated genotype is therefore unlabeled rather than negative.

##### SR-M2. The NCD construct rule

NCD B is credited through decreased grooming behavior and aphagia under the construct rule that a motor term does not model interference with independence. Without that rule, impaired coordination alone credits criterion B to 1,292 genotypes (25 of them curated NCD models, which then meet the symptom-count rule), and 1,103 genotypes in all would meet it. The NCD coverage AUROC is then 0.775 (0.701 to 0.839) against 0.761 with the rule, and the support AUROC is 0.674 (0.602 to 0.746) against 0.668.

##### SR-M3. The retrieval score without LLM admission

Curated models received more behavioral phenotyping than the background, and two baselines without criterion content measure how much of the separation that alone produces (S8 Table, part B). The first is the number of a genotype’s behavioral annotations. It separates curated models from the background at support AUROC 0.895 for ASD, 0.849 for SCZ, 0.787 for IDD and 0.714 for NCD, above the support score for the last three. The second is the retrieval score, which takes for each anchor the best similarity score among the candidate terms a genotype carries, with no LLM verdict and no permutation null (Methods). It reaches 0.938, 0.827, 0.784 and 0.781 in the same order, because it also grows with the number of annotations. The random sets behind the support score match a genotype’s total annotation count but not its number of behavioral annotations (its behavioral depth). Depth matching therefore reweights the background so that its distribution of behavioral depth equals that of the curated models. Because the weights are built from binned behavioral counts, the count itself falls to near chance (0.51 to 0.53), which shows that the matching worked. The support score falls from 0.956 to 0.895 (0.852 to 0.931) for ASD and from 0.682 to 0.596 (0.517 to 0.674) for SCZ. It falls from 0.751 to 0.667 (0.618 to 0.736) for IDD and from 0.668 to 0.582 (0.508 to 0.653) for NCD. The separation that survives equal phenotyping depth is therefore large for ASD, moderate for IDD and small for SCZ and NCD. The retrieval score falls to 0.768, 0.540, 0.603 and 0.654 under the same weights, and the support score exceeds it with an interval above zero for ASD and IDD only (S8 Table, part B). The two scores differ in the pairs they read, in how they combine them and in the permutation null, so the difference cannot be attributed to the LLM’s admission alone. For SCZ the retrieval score on the admitted pairs alone falls to 0.397 at equal depth, so the SCZ separation comes from the comparison with random annotation sets of the same size.

*S8 Table, part B. Baselines and depth matching. Support AUROCs in the evaluation set with 95% gene-clustered intervals. The behavioral count is the number of a genotype’s annotations at or below abnormal behavior (MP:0004924). The retrieval score without LLM admission takes for each anchor the best combined retrieval score among the genotype’s candidate terms, combines anchors within a unit by noisy-OR and averages the units, with no verdict and no permutation null. The permutation null matches each genotype to random annotation sets of its own total size without regard to its behavioral depth. Depth matching therefore reweights the background to the curated models’ distribution of behavioral annotation depth (0, 1 to 2, 3 to 5, 6 to 10, 11 or more). Because the bins are bins of the behavioral count, the depth-matched count row is a check that the matching worked.*

| Comparison | ASD (85 curated) | SCZ (42) | IDD (111) | NCD (67) |
| --- | --- | --- | --- | --- |
| Support score (Table 3 of the main text) | 0.956 (0.920 to 0.983) | 0.682 (0.603 to 0.761) | 0.751 (0.705 to 0.820) | 0.668 (0.595 to 0.737) |
| Support score, depth matched | 0.895 (0.852 to 0.931) | 0.596 (0.517 to 0.674) | 0.667 (0.618 to 0.736) | 0.582 (0.508 to 0.653) |
| Number of behavioral annotations, no criterion content | 0.895 | 0.849 | 0.787 | 0.714 |
| Number of behavioral annotations, depth matched | 0.524 (0.499 to 0.545) | 0.527 (0.493 to 0.565) | 0.511 (0.499 to 0.531) | 0.508 (0.484 to 0.533) |
| Retrieval score without LLM admission | 0.938 | 0.827 | 0.784 | 0.781 |
| Retrieval score without LLM admission, depth matched | 0.768 (0.718 to 0.819) | 0.540 (0.487 to 0.592) | 0.603 (0.563 to 0.652) | 0.654 (0.611 to 0.707) |
| Retrieval score on the admitted pairs only, depth matched | 0.893 (0.851 to 0.935) | 0.397 (0.324 to 0.478) | 0.650 (0.595 to 0.731) | 0.579 (0.503 to 0.663) |
| Support minus retrieval without admission, depth matched | +0.128 (0.086 to 0.165, p below 0.001) | +0.056 (-0.041 to 0.146, p 0.24) | +0.064 (0.009 to 0.135, p 0.024) | -0.072 (-0.150 to 0.013, p 0.08) |

##### S15 Table. The highest-credited genotypes per disorder

*S15 Table. The highest-credited genotypes per disorder and the number of criterion units each is credited with. Curated models are the highest-ranked curated genotypes by Diagnostic Coverage. Background genotypes are the highest-ranked genotypes carrying no label for the disorder.*

| Disorder | Curated models at the top (units credited) | Background genotypes at the top (units credited) |
| --- | --- | --- |
| ASD | Arid1b 6, then at 5 of 7: Shank3, Shank2, Arhgap32, Nexmif, Slc6a4, Cntnap2 and 13 others | CHMP2B transgene at Atp2b1 6 (curated for frontotemporal dementia), then at 5: Caskin1, Gnao1, Zeb1, Slc12a2, Ext1, Tshz3 and 37 others |
| SCZ | 25 models at 2 of 5 (A4 and A5), among them Disc1, Dtnbp1, Syngap1, Map6, Gria1, Cplx2 | 1,231 genotypes at 2 of 5, among them Lrit3, Npas4, Shank3, Sez6l |
| IDD | 26 models at 2 of 2, among them Nexmif, Arid1b, Dyrk1a, Tmco1, Capza2, Tcf4, Fmr1 | 307 genotypes at 2 of 2, among them Gabrb3, Adcy3, Nfix, Gria1, Syngap1 |
| NCD | 47 models at 1 of 2 (A), among them Snca, Grn, Htt, Chmp2b, Prnp, App, Psen1, Park7 | 66 genotypes at 2 of 2, among them Psip1, Dmrta2, Cckbr, Cebpa, Cplx2 |
| MDD | Adcy3 4 of 9 (A1 to A4), then at 2: Tph2, Crtc1, Gabrd, Gpr156, Car1, Disc1, Calhm2 | Slc18a2, Il6, Qrfp at 4 of 9, then Hivep2, Iglon5, St3gal4 at 3 |

##### S16 Table. Phenotype classes of Figure 3 and the admitted terms per class

Classes are read from the Mammalian Phenotype Ontology (release 2026-05-19) by ancestry, each term taking the first class in the order listed whose root is among its ancestors. Shares are of the behavioral annotations of the 327 curated models (primary labels). Admitted terms are distinct MP terms admitted on scored criteria in the match export of the primary analysis run. Script audit/composition_2026-11-06/composition.py.

| Class | MP root terms | Share of curated models’ behavioral annotations, percent | Admitted terms per disorder (ASD, SCZ, IDD, NCD, MDD) |
| --- | --- | --- | --- |
| social and vocal | MP:0002557 abnormal social/conspecific interaction behavior, MP:0001529 abnormal vocalization | 12.3 and 2.7 | 17, 4, 13, 3, 2 |
| cognition | MP:0014114 abnormal cognition | 18.7 | 3, 1, 36, 32, 0 |
| emotion and affect | MP:0002572 abnormal emotion/affect behavior | 16.6 | 2, 5, 1, 0, 3 |
| motor behavior | MP:0002066 abnormal motor capabilities/coordination/movement | 34.9 | 21, 30, 2, 7, 9 |
| other behavior | the other 15 children of MP:0004924 abnormal behavior (sensory capabilities and reflexes 5.4, grooming 3.4, seizures 2.5, the rest below 1.5 each) | 14.8 | 18, 3, 1, 6, 12 |
| nervous system morphology | MP:0003632 | not behavioral | 0, 0, 0, 1, 0 |
| nervous system physiology | MP:0003633 | not behavioral | 0, 0, 0, 1, 0 |
| mortality and growth | MP:0010768 mortality/aging, MP:0005378 growth/size/body region | not behavioral | 0, 0, 0, 0, 5 |
| other physiology and morphology | every other top-level MP branch | not behavioral | 0, 0, 0, 0, 0 |

##### S17 Table. The IMPC-only test

Universe: IMPC-tagged genotypes of the evaluation set with curated models removed. Positives: genotypes whose gene is an H gene. Intervals: 95% gene-clustered bootstrap, 2,000 replicates. Depth matched: background reweighted to the positives’ behavioral-depth bins, gene-clustered, 500 replicates.

| Disorder | IMPC genotypes | H-gene positives (genes) | Support AUROC | Coverage AUROC | Behavioral count AUROC | PhenoDigm recomputed | Support minus PhenoDigm (p) | Support AUROC, depth matched |
| --- | --- | --- | --- | --- | --- | --- | --- | --- |
| ASD | 2,460 | 36 (33) | 0.491 (0.428 to 0.563) | 0.593 (0.494 to 0.696) | 0.577 (0.481 to 0.672) | 0.499 (0.406 to 0.593) | -0.008 (-0.081 to +0.065, p 0.82) | 0.489 (0.430 to 0.555) |
| SCZ | 2,543 | 8 (7) | 0.514 (0.442 to 0.712) | 0.513 (0.326 to 0.740) | 0.517 (0.342 to 0.741) | 0.483 (0.321 to 0.667) | +0.031 (-0.147 to +0.207, p 0.68) | 0.517 (0.436 to 0.695) |
| IDD | 2,588 | 213 (190) | 0.521 (0.498 to 0.546) | 0.524 (0.492 to 0.556) | 0.565 (0.521 to 0.610) | 0.502 (0.456 to 0.546) | +0.020 (-0.027 to +0.070, p 0.41) | 0.517 (0.495 to 0.540) |
| NCD | 2,438 | 45 (44) | 0.484 (0.470 to 0.509) | 0.513 (0.440 to 0.591) | 0.475 (0.394 to 0.563) | 0.477 (0.398 to 0.559) | +0.007 (-0.074 to +0.087, p 0.87) | 0.486 (0.469 to 0.513) |

##### S18 Table. The depth-stratified null and PhenoDigm referred to the same nulls

Evaluation set, corrected labels, K4. Intervals are 95% gene-clustered bootstrap (2,000 replicates), and the paired difference of the two nulls is given with its 90% interval as in S23 Table. Depth matched: background reweighted to the curated models’ behavioral-depth bins.

| Disorder | n (curated) | Support AUROC, null of record | Support AUROC, stratified null (95%) | Stratified minus record (90%) | Any admitted term, indicator AUROC | Depth matched, record | Depth matched, stratified |
| --- | --- | --- | --- | --- | --- | --- | --- |
| ASD | 11,961 (85) | 0.956 | 0.957 (0.921 to 0.983) | +0.001 (+0.000 to +0.002) | 0.896 | 0.895 | 0.898 |
| SCZ | 12,486 (42) | 0.682 | 0.682 (0.604 to 0.760) | +0.000 (-0.002 to +0.002) | 0.684 | 0.596 | 0.593 |
| IDD | 12,382 (111) | 0.751 | 0.752 (0.706 to 0.821) | +0.001 (-0.000 to +0.002) | 0.721 | 0.667 | 0.672 |
| NCD | 11,603 (67) | 0.668 | 0.665 (0.593 to 0.735) | -0.003 (-0.010 to +0.005) | 0.680 | 0.582 | 0.581 |

| Disorder | PhenoDigm raw | PhenoDigm, size-matched null | PhenoDigm, behavior-matched null | FACET minus PhenoDigm, size matched (95%, p) | FACET minus PhenoDigm, behavior matched (95%, p) |
| --- | --- | --- | --- | --- | --- |
| ASD | 0.772 (0.707 to 0.831) | 0.792 (0.730 to 0.848) | 0.657 (0.574 to 0.734) | +0.163 (+0.112 to +0.222, p below 0.001) | +0.298 (+0.220 to +0.383, p below 0.001) |
| SCZ | 0.772 (0.703 to 0.840) | 0.781 (0.717 to 0.844) | 0.680 (0.596 to 0.763) | -0.099 (-0.198 to -0.013, p 0.033) | +0.002 (-0.118 to +0.108, p 0.98) |
| IDD | 0.706 (0.623 to 0.757) | 0.714 (0.626 to 0.767) | 0.624 (0.532 to 0.680) | +0.037 (-0.040 to +0.157, p 0.38) | +0.127 (+0.042 to +0.256, p 0.002) |
| NCD | 0.593 (0.543 to 0.637) | 0.594 (0.540 to 0.638) | 0.541 (0.488 to 0.586) | +0.074 (+0.007 to +0.147, p 0.028) | +0.127 (+0.057 to +0.202, p below 0.001) |

##### S19 Table. The admitted terms and the curated models’ annotations behind the SCZ A5 and MDD A1 and A2 credits

Admitted terms of record per unit, and the curated models’ own annotations at or below them (number of curated models carrying each, of 42 SCZ and 8 MDD).

| Unit | Admitted terms | Curated models’ annotations crediting the unit (models) |
| --- | --- | --- |
| SCZ A5 (negative symptoms) | abnormal emotion/affect behavior (MP:0002572), bradykinesia (MP:0005156) | increased thigmotaxis (7), increased exploration in new environment (6), decreased anxiety-related response (5), abnormal response to novel object (5), abnormal emotion/affect behavior (5), increased anxiety-related response (5), decreased exploration in new environment (4), impaired passive avoidance behavior (4), behavioral despair (3), increased coping response (2), anhedonia (2), decreased aggression towards mice (2), and one each of abnormal response to social novelty, decreased thigmotaxis, increased aggression towards mice, decreased freezing behavior, decreased aggression, abnormal habituation to a new environment and increased freezing behavior, none through bradykinesia |
| MDD A1 (depressed mood) | increased depression-related behavior (MP:0014414) | behavioral despair (6), anhedonia (2), decreased coping response (2), increased depression-related behavior (1) |
| MDD A2 (loss of interest or pleasure) | anhedonia (MP:0009710), abnormal depression-related behavior (MP:0003360), decreased female mating frequency (MP:0014602) | behavioral despair (6), abnormal depression-related behavior (3), anhedonia (2), decreased coping response (2), increased depression-related behavior (1) |

##### S20 Table. The SCZ candidate pairs built on endophenotype and drug-response terms, by term family, criterion anchor and analog grade

Pairs on anchors of criteria A1 to A5 (59 of the 115 pairs carrying these terms in the SCZ candidate primary table). Grades are the pair judgment’s analog grade. No pair is strictly eligible. The 56 pairs on anchors outside the five criteria are omitted, since those anchors credit no criterion.

| Term family (MP terms) | Criterion anchor | Pairs | Weak support | Mismatch | Mechanistic only | Close support | Not judged |
| --- | --- | --- | --- | --- | --- | --- | --- |
| Startle reflex (abnormal, absent, increased, decreased) | A1 delusions (Delusional perception, Abnormal judgment) | 8 | 4 | 4 | 0 | 0 | 0 |
| Startle reflex | A2 hallucinations (Hallucinations, Abnormal perceptual state) | 8 | 6 | 2 | 0 | 0 | 0 |
| Startle reflex | A3 disorganized speech (Disordered formal thought process) | 4 | 0 | 4 | 0 | 0 | 0 |
| Startle reflex | A4 disorganized behavior (Atypical behavior) | 4 | 1 | 3 | 0 | 0 | 0 |
| Startle reflex | A5 negative symptoms (Abnormal diminished volition) | 4 | 0 | 4 | 0 | 0 | 0 |
| Amphetamine and methamphetamine response (6 terms) | A1 delusions | 12 | 6 | 3 | 3 | 0 | 0 |
| Amphetamine and methamphetamine response | A4 disorganized behavior | 6 | 1 | 1 | 4 | 0 | 0 |
| Latent inhibition of conditioning | A1 delusions | 3 | 3 | 0 | 0 | 0 | 0 |
| Latent inhibition of conditioning | A2 hallucinations | 2 | 2 | 0 | 0 | 0 | 0 |
| Latent inhibition of conditioning | A3 disorganized speech | 1 | 1 | 0 | 0 | 0 | 0 |
| Latent inhibition of conditioning | A4 disorganized behavior | 1 | 0 | 0 | 1 | 0 | 0 |
| Prepulse inhibition and sensorimotor gating (6 terms) | A1 delusions | 6 | 2 | 1 | 1 | 1 | 1 |
| All |  | 59 | 26 | 22 | 9 | 1 | 1 |

Source: FACET_release/data/pairs/SCZ_judged_pairs.tsv, columns mp_label, human_domain, behavioral_analog_type, strict_eligible, read 2026-11-07.

##### S21 Table. The hand-curated anchor sets that gated mouse domain labeling, with their outcome in the primary taxonomies

Expected domain and admissibility as written in each disorder’s rebuild script. Outcome read from the primary taxonomy on 2026-11-07: the domain of the first call, and admission to a criterion unit at the second call (IDD, NCD) or the strict flag of record (SCZ, MDD). Domain names are the project’s symptom-domain labels (S1 Table).

| Disorder | MP term | Expected domain | Expected | Domain of record | Domain match | Outcome of record |
| --- | --- | --- | --- | --- | --- | --- |
| SCZ | MP:0002822 catalepsy | catatonic disorganized behavior | admissible | catatonic disorganized behavior | yes | admitted (catatonic_disorganized_behavior) |
| SCZ | MP:0001399 hyperactivity | catatonic disorganized behavior | admissible | catatonic disorganized behavior | yes | admitted (catatonic_disorganized_behavior) |
| SCZ | MP:0001361 social withdrawal | negative symptoms | admissible | negative symptoms | yes | admitted (negative_symptoms) |
| SCZ | MP:0009710 anhedonia | negative symptoms | admissible | negative symptoms | yes | admitted (negative_symptoms) |
| SCZ | MP:0021111 decreased social investigation | negative symptoms | admissible | negative symptoms | yes | admitted (negative_symptoms) |
| SCZ | MP:0003088 abnormal prepulse inhibition | mechanism substrate | control (refuse) | mechanism substrate | yes | refused |
| SCZ | MP:0002634 abnormal sensorimotor gating | mechanism substrate | control (refuse) | mechanism substrate | yes | refused |
| SCZ | MP:0002635 reduced sensorimotor gating | mechanism substrate | control (refuse) | mechanism substrate | yes | refused |
| SCZ | MP:0008428 abnormal spatial working memory | cognitive associated | control (refuse) | cognitive associated | yes | refused |
| SCZ | MP:0001463 abnormal spatial learning | cognitive associated | control (refuse) | cognitive associated | yes | refused |
| SCZ | MP:0001900 impaired synaptic plasticity | mechanistic neurobiology | control (refuse) | mechanistic neurobiology | yes | refused |
| SCZ | MP:0001905 abnormal dopamine level | mechanistic neurobiology | control (refuse) | mechanistic neurobiology | yes | refused |
| SCZ | MP:0002135 abnormal kidney morphology | non neurological or other | control (refuse) | non neurological or other | yes | refused |
| IDD | MP:0008428 abnormal spatial working memory | intellectual functioning | admissible | intellectual functioning | yes | admitted (A) |
| IDD | MP:0002062 abnormal associative learning | intellectual functioning | admissible | intellectual functioning | yes | admitted (A) |
| IDD | MP:0001458 abnormal object recognition memory | intellectual functioning | admissible | intellectual functioning | yes | admitted (A) |
| IDD | MP:0001463 abnormal spatial learning | intellectual functioning | admissible | intellectual functioning | yes | admitted (A) |
| IDD | MP:0001469 abnormal contextual conditioning behavior | intellectual functioning | admissible | intellectual functioning | yes | admitted (A) |
| IDD | MP:0001360 abnormal social investigation | adaptive functioning | admissible | adaptive functioning | yes | admitted (B) |
| IDD | MP:0001447 abnormal nest building behavior | adaptive functioning | admissible | adaptive functioning | yes | admitted (B) |
| IDD | MP:0001440 abnormal grooming behavior | adaptive functioning | admissible | adaptive functioning | yes | refused |
| IDD | MP:0021165 behavioral developmental delay | developmental onset | admissible | developmental onset | yes | refused |
| IDD | MP:0001290 delayed eyelid opening | developmental onset | admissible | developmental onset | yes | refused |
| IDD | MP:0000003 abnormal adipose tissue morphology | non neurological or other | control (refuse) | non neurological or other | yes | refused |
| IDD | MP:0002135 abnormal kidney morphology | non neurological or other | control (refuse) | non neurological or other | yes | refused |
| IDD | MP:0004859 abnormal synaptic plasticity | mechanistic neurobiology | control (refuse) | mechanistic neurobiology | yes | refused |
| IDD | MP:0001905 abnormal dopamine level | mechanistic neurobiology | control (refuse) | mechanistic neurobiology | yes | refused |
| IDD | MP:0003861 abnormal nervous system development | generic neurological abnormality | control (refuse) | generic neurological abnormality | yes | refused |
| IDD | MP:0000060 delayed bone ossification | non neurological or other | control (refuse) | non neurological or other | yes | refused |
| IDD | MP:0002174 abnormal gastrulation movements | non neurological or other | control (refuse) | non neurological or other | yes | refused |
| NCD | MP:0001463 abnormal spatial learning | cognitive executive | admissible | cognitive executive | yes | admitted (A) |
| NCD | MP:0001458 abnormal object recognition memory | cognitive executive | admissible | cognitive executive | yes | admitted (A) |
| NCD | MP:0002062 abnormal associative learning | cognitive executive | admissible | cognitive executive | yes | admitted (A) |
| NCD | MP:0001469 abnormal contextual conditioning behavior | cognitive executive | admissible | cognitive executive | yes | admitted (A) |
| NCD | MP:0001516 abnormal motor coordination/balance | motor coordination | admissible | motor coordination | yes | admitted (A) |
| NCD | MP:0001406 abnormal gait | motor coordination | admissible | motor coordination | yes | admitted (A) |
| NCD | MP:0001393 ataxia | motor coordination | admissible | motor coordination | yes | admitted (A) |
| NCD | MP:0021112 impaired social recognition | social cognition empathy | admissible | social cognition empathy | yes | admitted (A) |
| NCD | MP:0020396 abnormal social recognition | social cognition empathy | admissible | social cognition empathy | yes | admitted (A) |
| NCD | MP:0001447 abnormal nest building behavior | adaptive functioning | admissible | adaptive functioning | yes | admitted (B) |
| NCD | MP:0001440 abnormal grooming behavior | adaptive functioning | admissible | adaptive functioning | yes | refused |
| NCD | MP:0003329 amyloid beta deposits | neuropathology progressive | admissible | neuropathology progressive | yes | refused |
| NCD | MP:0004250 tau protein deposits | neuropathology progressive | admissible | neuropathology progressive | yes | refused |
| NCD | MP:0003214 neurofibrillary tangles | neuropathology progressive | admissible | neuropathology progressive | yes | refused |
| NCD | MP:0001900 impaired synaptic plasticity | mechanistic neurobiology | control (refuse) | mechanistic neurobiology | yes | refused |
| NCD | MP:0001905 abnormal dopamine level | mechanistic neurobiology | control (refuse) | mechanistic neurobiology | yes | refused |
| NCD | MP:0002135 abnormal kidney morphology | non neurological or other | control (refuse) | non neurological or other | yes | refused |
| MDD | MP:0002573 behavioral despair | depressed mood | admissible | depressed mood | yes | admitted (depressed_mood) |
| MDD | MP:0014414 increased depression-related behavior | depressed mood | admissible | depressed mood | yes | admitted (depressed_mood) |
| MDD | MP:0020379 abnormal sucrose solution preference | anhedonia | admissible | anhedonia | yes | admitted (anhedonia) |
| MDD | MP:0005449 abnormal food intake | weight appetite change | admissible | weight appetite change | yes | admitted (weight_appetite_change) |
| MDD | MP:0001259 abnormal body weight | weight appetite change | admissible | non neurological or other | no | refused |
| MDD | MP:0001501 abnormal sleep pattern | sleep disturbance | admissible | sleep disturbance | yes | admitted (sleep_disturbance) |
| MDD | MP:0003194 abnormal frequency of paradoxical sleep | sleep disturbance | admissible | sleep disturbance | yes | admitted (sleep_disturbance) |
| MDD | MP:0001399 hyperactivity | psychomotor change | admissible | psychomotor change | yes | admitted (psychomotor_change) |
| MDD | MP:0001402 decreased locomotor activity | psychomotor change | admissible | psychomotor change | yes | admitted (psychomotor_change) |
| MDD | MP:0010053 decreased grip strength | fatigue | admissible | fatigue | yes | admitted (fatigue) |
| MDD | MP:0014370 impaired spatial working memory | concentration indecisiveness | admissible | concentration indecisiveness | yes | admitted (concentration_indecisiveness) |
| MDD | MP:0000003 abnormal adipose tissue morphology | non neurological or other | control (refuse) | non neurological or other | yes | refused |
| MDD | MP:0004859 abnormal synaptic plasticity | mechanistic neurobiology | control (refuse) | mechanistic neurobiology | yes | refused |
| MDD | MP:0001905 abnormal dopamine level | mechanistic neurobiology | control (refuse) | mechanistic neurobiology | yes | refused |
| MDD | MP:0003861 abnormal nervous system development | generic neurological abnormality | control (refuse) | generic neurological abnormality | yes | refused |

Sources: rebuild_scz_taxonomy_5domain.py, rebuild_id_taxonomy.py, rebuild_neurocognitive_taxonomy.py, rebuild_mdd_taxonomy.py (ANCHOR_SET), read against mp_term_taxonomy_scz_5domain_v1, mp_term_taxonomy_criterion_v1_ID, mp_term_taxonomy_criterion_v1_NCD and mp_term_taxonomy_mdd_v1 (03_mp_term_taxonomy.tsv).

##### S22 Table. The anchor-gate rerun of 2026-11-08, 15 terms per disorder, against the expectations and the primary run

Expected: the criterion unit (ASD, IDD, NCD) or domain and admissibility (SCZ, MDD) fixed before the run. Record: the primary taxonomy. Rerun: the 2026-11-08 run. For SCZ and MDD, admitted means the strict face-validity flag is set.

| Disorder | MP term | Expected | Record | Rerun | Rerun = record | Rerun = expected |
| --- | --- | --- | --- | --- | --- | --- |
| ASD | MP:0001360 abnormal social investigation | A1 | A1 | A1 | yes | yes |
| ASD | MP:0001361 social withdrawal | A1 | A3 | A3 | yes | no |
| ASD | MP:0021111 decreased social investigation | A1 | A1 | A1 | yes | yes |
| ASD | MP:0001529 abnormal vocalization | A1 | A1 | A1 | yes | yes |
| ASD | MP:0020396 abnormal social recognition | A3 | A3 | A3 | yes | yes |
| ASD | MP:0021112 impaired social recognition | A3 | A3 | A3 | yes | yes |
| ASD | MP:0001400 hyperresponsive | B1 | B4 | B4 | yes | no |
| ASD | MP:0001441 increased grooming behavior | B1 | B1 | B1 | yes | yes |
| ASD | MP:0001440 abnormal grooming behavior | B2 | B1 | B1 | yes | no |
| ASD | MP:0002733 abnormal thermal nociception | B4 | B4 | B4 | yes | yes |
| ASD | MP:0003998 decreased thermal nociceptive threshold | B4 | B4 | B4 | yes | yes |
| ASD | MP:0001486 abnormal startle reflex | B4 | refused | refused | yes | no |
| ASD | MP:0002064 seizures | refuse | refused | refused | yes | yes |
| ASD | MP:0000745 tremors | refuse | refused | refused | yes | yes |
| ASD | MP:0002135 abnormal kidney morphology | refuse | refused | refused | yes | yes |
| IDD | MP:0008428 abnormal spatial working memory | A | A | A | yes | yes |
| IDD | MP:0002062 abnormal associative learning | A | A | A | yes | yes |
| IDD | MP:0001458 abnormal object recognition memory | A | A | A | yes | yes |
| IDD | MP:0001463 abnormal spatial learning | A | A | A | yes | yes |
| IDD | MP:0001469 abnormal contextual conditioning behavior | A | A | A | yes | yes |
| IDD | MP:0001360 abnormal social investigation | B | B | B | yes | yes |
| IDD | MP:0001447 abnormal nest building behavior | B | B | B | yes | yes |
| IDD | MP:0001440 abnormal grooming behavior | B | refused | refused | yes | no |
| IDD | MP:0000003 abnormal adipose tissue morphology | refuse | refused | refused | yes | yes |
| IDD | MP:0002135 abnormal kidney morphology | refuse | refused | refused | yes | yes |
| IDD | MP:0004859 abnormal synaptic plasticity | refuse | refused | refused | yes | yes |
| IDD | MP:0001905 abnormal dopamine level | refuse | refused | refused | yes | yes |
| IDD | MP:0003861 abnormal nervous system development | refuse | refused | refused | yes | yes |
| IDD | MP:0000060 delayed bone ossification | refuse | refused | refused | yes | yes |
| IDD | MP:0002174 abnormal gastrulation movements | refuse | refused | refused | yes | yes |
| NCD | MP:0001463 abnormal spatial learning | A | A | A | yes | yes |
| NCD | MP:0001458 abnormal object recognition memory | A | A | A | yes | yes |
| NCD | MP:0002062 abnormal associative learning | A | A | A | yes | yes |
| NCD | MP:0001469 abnormal contextual conditioning behavior | A | A | A | yes | yes |
| NCD | MP:0008428 abnormal spatial working memory | A | A | A | yes | yes |
| NCD | MP:0001516 abnormal motor coordination/balance | A | A | A | yes | yes |
| NCD | MP:0001406 abnormal gait | A | A | A | yes | yes |
| NCD | MP:0001393 ataxia | A | A | A | yes | yes |
| NCD | MP:0021112 impaired social recognition | A | A | A | yes | yes |
| NCD | MP:0020396 abnormal social recognition | A | A | A | yes | yes |
| NCD | MP:0001447 abnormal nest building behavior | B | B | B | yes | yes |
| NCD | MP:0001440 abnormal grooming behavior | B | refused | refused | yes | no |
| NCD | MP:0001900 impaired synaptic plasticity | refuse | refused | refused | yes | yes |
| NCD | MP:0001905 abnormal dopamine level | refuse | refused | refused | yes | yes |
| NCD | MP:0002135 abnormal kidney morphology | refuse | refused | refused | yes | yes |
| SCZ | MP:0002822 catalepsy | catatonic disorganized behavior, admissible | catatonic disorganized behavior, admitted | catatonic disorganized behavior, admitted | yes | yes |
| SCZ | MP:0001399 hyperactivity | catatonic disorganized behavior, admissible | catatonic disorganized behavior, admitted | catatonic disorganized behavior, admitted | yes | yes |
| SCZ | MP:0001361 social withdrawal | negative symptoms, admissible | negative symptoms, admitted | negative symptoms, admitted | yes | yes |
| SCZ | MP:0009710 anhedonia | negative symptoms, admissible | negative symptoms, admitted | negative symptoms, admitted | yes | yes |
| SCZ | MP:0021111 decreased social investigation | negative symptoms, admissible | negative symptoms, admitted | negative symptoms, admitted | yes | yes |
| SCZ | MP:0003088 abnormal prepulse inhibition | mechanism substrate, refuse | mechanism substrate, refused | mechanism substrate, refused | yes | yes |
| SCZ | MP:0002634 abnormal sensorimotor gating | mechanism substrate, refuse | mechanism substrate, refused | mechanism substrate, refused | yes | yes |
| SCZ | MP:0002635 reduced sensorimotor gating | mechanism substrate, refuse | mechanism substrate, refused | mechanism substrate, refused | yes | yes |
| SCZ | MP:0008428 abnormal spatial working memory | cognitive associated, refuse | cognitive associated, refused | cognitive associated, refused | yes | yes |
| SCZ | MP:0001463 abnormal spatial learning | cognitive associated, refuse | cognitive associated, refused | cognitive associated, refused | yes | yes |
| SCZ | MP:0001900 impaired synaptic plasticity | mechanistic neurobiology, refuse | mechanistic neurobiology, refused | mechanistic neurobiology, refused | yes | yes |
| SCZ | MP:0001905 abnormal dopamine level | mechanistic neurobiology, refuse | mechanistic neurobiology, refused | mechanistic neurobiology, refused | yes | yes |
| SCZ | MP:0002135 abnormal kidney morphology | non neurological or other, refuse | non neurological or other, refused | non neurological or other, refused | yes | yes |
| SCZ | MP:0002572 abnormal emotion/affect behavior | negative symptoms, admissible | non neurological or other, refused | non neurological or other, refused | yes | no |
| SCZ | MP:0002064 seizures | mechanistic neurobiology, refuse | non neurological or other, refused | non neurological or other, refused | yes | no |
| MDD | MP:0002573 behavioral despair | depressed mood, admissible | depressed mood, admitted | depressed mood, admitted | yes | yes |
| MDD | MP:0014414 increased depression-related behavior | depressed mood, admissible | depressed mood, admitted | depressed mood, admitted | yes | yes |
| MDD | MP:0020379 abnormal sucrose solution preference | anhedonia, admissible | anhedonia, admitted | anhedonia, admitted | yes | yes |
| MDD | MP:0005449 abnormal food intake | weight appetite change, admissible | weight appetite change, admitted | weight appetite change, admitted | yes | yes |
| MDD | MP:0001259 abnormal body weight | weight appetite change, admissible | non neurological or other, refused | weight appetite change, admitted | no | yes |
| MDD | MP:0001501 abnormal sleep pattern | sleep disturbance, admissible | sleep disturbance, admitted | sleep disturbance, admitted | yes | yes |
| MDD | MP:0003194 abnormal frequency of paradoxical sleep | sleep disturbance, admissible | sleep disturbance, admitted | sleep disturbance, admitted | yes | yes |
| MDD | MP:0001399 hyperactivity | psychomotor change, admissible | psychomotor change, admitted | psychomotor change, admitted | yes | yes |
| MDD | MP:0001402 decreased locomotor activity | psychomotor change, admissible | psychomotor change, admitted | psychomotor change, admitted | yes | yes |
| MDD | MP:0010053 decreased grip strength | fatigue, admissible | fatigue, admitted | fatigue, admitted | yes | yes |
| MDD | MP:0014370 impaired spatial working memory | concentration indecisiveness, admissible | concentration indecisiveness, admitted | concentration indecisiveness, admitted | yes | yes |
| MDD | MP:0000003 abnormal adipose tissue morphology | non neurological or other, refuse | non neurological or other, refused | non neurological or other, refused | yes | yes |
| MDD | MP:0004859 abnormal synaptic plasticity | mechanistic neurobiology, refuse | mechanistic neurobiology, refused | mechanistic neurobiology, refused | yes | yes |
| MDD | MP:0001905 abnormal dopamine level | mechanistic neurobiology, refuse | mechanistic neurobiology, refused | mechanistic neurobiology, refused | yes | yes |
| MDD | MP:0003861 abnormal nervous system development | generic neurological abnormality, refuse | generic neurological abnormality, refused | generic neurological abnormality, refused | yes | yes |

Source: audit/anchor_gate_2026-11-07/ (gate files, RESULT_gate.md, pilot_SCZ.log, pilot_MDD.log).

##### S23 Table. Validation of the pair judgment by alternate LLMs and the admission control

Contrasts are paired differences in support AUROC with 90% gene-clustered intervals against the pre-specified margin of 0.05. The admission control reports the one-sided Monte Carlo p against 100 matched draws of refused pairs.

| Check | ASD | SCZ | IDD | NCD |
| --- | --- | --- | --- | --- |
| claude-opus-4-8 minus primary analysis run | -0.006 (-0.010 to -0.002), equivalent | +0.182 (+0.124 to +0.242), not equivalent | +0.028 (+0.003 to +0.045), equivalent | +0.025 (-0.029 to +0.095), inconclusive |
| gemini-2.5-pro minus primary analysis run | -0.014 (-0.024 to -0.006), equivalent | +0.159 (+0.085 to +0.235), not equivalent | +0.090 (+0.023 to +0.138), inconclusive | +0.020 (-0.016 to +0.065), inconclusive |
| Support AUROC: primary analysis run, claude-opus-4-8, gemini-2.5-pro | 0.956, 0.950, 0.942 | 0.682, 0.864, 0.842 | 0.751, 0.779, 0.841 | 0.668, 0.693, 0.688 |
| Admitted pairs: primary analysis run, claude-opus-4-8, gemini-2.5-pro | 145, 370, 564 | 196, 425, 683 | 138, 363, 574 | 95, 224, 266 |
| Agreement on admission, % (kappa): claude, gemini against primary analysis run | 96.5 (0.51), 93.5 (0.34) | 95.9 (0.53), 92.5 (0.39) | 94.6 (0.50), 90.0 (0.35) | 94.7 (0.54), 92.6 (0.43) |
| Admission negative control, p | 0.010 | 0.109 | 0.020 | 0.554 |

##### S24 Table. Curated labels, claim papers and genes, with the IMPC-only test

Removal of the claim paper is symmetric between curated models and comparable background genotypes. Relatives are uncurated, behaviorally annotated genotypes of a curated model’s gene that cite none of its claim papers, and the attention control is the genotypes of H genes without a curated model. AUROC rows are support AUROC with 95% gene-clustered intervals unless the row names the coverage family, and the relatives rows are stratified by behavioral depth. The IMPC rows rank, among IMPC-tagged genotypes of the evaluation set with curated models removed, the genotypes of H genes (36, 8, 213 and 45) against the other IMPC genotypes (about 2,400 per disorder). The H-gene odds ratios and the describing-literature contrast are in S14 Table.

| Quantity | ASD | SCZ | IDD | NCD |
| --- | --- | --- | --- | --- |
| Curated models whose every annotation comes from the claim paper | 78 of 85 | 34 of 42 | 105 of 121 | 52 of 71 |
| Curated models with any credited unit after claim-paper removal | 4 of 85 | 6 of 42 | 8 of 121 | 6 of 71 |
| Coverage AUROC after claim-paper removal | 0.504 (0.486 to 0.526) | 0.542 (0.494 to 0.603) | 0.520 (0.494 to 0.546) | 0.521 (0.495 to 0.548) |
| Support AUROC after claim-paper removal | 0.487 (0.468 to 0.506) | 0.486 (0.484 to 0.487) | 0.509 (0.486 to 0.535) | 0.505 (0.483 to 0.532) |
| IMPC genotypes only, H-gene genotypes against the rest: support AUROC | 0.491 (0.428 to 0.563) | 0.514 (0.442 to 0.712) | 0.521 (0.498 to 0.546) | 0.484 (0.470 to 0.509) |
| IMPC genotypes only: PhenoDigm recomputed | 0.499 (0.406 to 0.593) | 0.483 (0.321 to 0.667) | 0.502 (0.456 to 0.546) | 0.477 (0.398 to 0.559) |

##### S25 Table. Data sources of the primary analysis run

| Source | Version or snapshot | Use in FACET |
| --- | --- | --- |
| DSM-5-TR [3] | 2022 text revision | Diagnostic criteria, paraphrased for the project. S1 Table gives the short form used in this paper. The LLM steps of the primary analysis run read the fuller paraphrase deposited with the code release and no DSM-5-TR text |
| Human Phenotype Ontology [23] | Release 2026-02-16 | Human vocabulary for anchors |
| Mammalian Phenotype Ontology [35] | Release 2026-05-19 | Mouse vocabulary (14,750 reachable terms, 11,485 annotated) |
| MGI genotype phenotype report [20] | 2026-03-23 | Genotype annotations. The report carries abnormal phenotypes only, so a normal result leaves no annotation |
| MGI disease model reports | 2026-04-06 | Disease labels, disease-gene associations and explicit negative records |
| MGI marker references and IMSR strain report (KOMP report) | October 2026 download, secondary analyses only | Publication and strain counts per gene, as measures of study attention |
| IMPC statistical results [36] | Snapshot of record | Behavioral procedures completed with normal results, to bound untested from normal |
| Monarch semantic similarity table [37] | Version 0.4 (file of 2024-11-19) | Ontology similarity between HPO and MP terms, two channels of the similarity score and the PhenoDigm comparator [21] |
| PubTator3 [38] | Accessed 2026-10 | Disease annotations of the papers describing candidate genotypes |
| Human genetic gene lists | S5 Table | Genes implicated by human genetics without mouse data |
| MGI report 2026-10-05, MP release 2026-09-23, disease reports 2026-10-12 | Later releases | Drift analyses only |

##### S26 Table. The eight named mapping rules

Each is a block of prompt text that a disorder’s configuration opts into. Locked means the date the text was fixed, before the primary taxonomy runs of 2026-07-28. Full text in S1 Text.

| Rule | Disorder | What it does | Locked |
| --- | --- | --- | --- |
| Grooming routing | ASD | Routes grooming, nest building and other patterned motor sequences to stereotyped movements (B1) and keeps them out of insistence on sameness (B2) | 2026-07-27 |
| Involuntary motor sign | ASD | Refuses tremor, dystonia, ataxia, clasping, seizures and other involuntary signs for every criterion, since repetition alone is not a stereotypy, and flags circling and head movements for review | 2026-07-27 |
| ASD construct refusals | ASD | Refuses memory and learning assays, losses of sensory capacity and maternal, agonistic or sexual behavior for the social and sensory criteria | 2026-07-27 |
| Human schooling constructs | IDD | Marks reading, writing, arithmetic, IQ and school or workplace performance as unassessable in a mouse so that they are not counted as measured zeros, and refuses hearing terms for language | 2026-07-27 |
| Decline versus deficit | NCD | Admits a deficit in the right cognitive domain although MP has no longitudinal frame, and records the missing qualifier as a disclosed concession | 2026-07-28 |
| Language floor | NCD | Admits vocalization and communication terms on their merits and flags them for review, so that the project’s language floor is tested against real candidates | 2026-07-28 |
| SCZ motor boundary from DSM | SCZ | Reads A4 from the DSM-5-TR definition of disorganized and catatonic behavior, admitting posturing, negativism and agitation and refusing seizures, startle, elementary motor signs and bare activity level, and flags reproductive behavior under A5 | 2026-07-28 |
| Mechanism is not face validity | SCZ | Flags prepulse inhibition, sensorimotor gating, latent inhibition and psychotomimetic drug responses at mapping as endophenotypes without a criterion counterpart. The pair judgment, which carries no SCZ rule, independently refused every such pair on a criterion anchor (S20 Table) | 2026-07-27 |

##### S27 Table. Fixed parameters of anchoring, retrieval, routing and judgment

Full definitions in S1 Text.

| Parameter | Value |
| --- | --- |
| Anchor shortlist | Top 8 lexical plus top 12 semantic matches, merged at 0.65 semantic and 0.35 lexical, truncated to 12 |
| Candidates per anchor | Top 20 by embedding and top 20 by lexical score, plus injected annotated descendants of tiers 1 to 4 |
| Channel weights (embedding, lexical, Resnik, Jaccard) | 0.45, 0.25, 0.20, 0.10, renormalized over present channels |
| Lexical factor when both ontology channels are absent | 0.5, so that the score is 0.78 embedding and 0.22 lexical on the 97 to 99% of pairs lacking ontology channels. Set on ASD, swept in S31 Table |
| Bundle | 5 candidates, when the top two scores lie within 0.08 or the top score lies in 0.35 to 0.70 |
| Judgment floor | 0.57 |
| Score adjustment by the LLM | +0.10 promote, 0 keep, -0.10 demote, -0.25 abstain |
| Admission | Strictly eligible, direct or close analog, direction not vetoed |

##### S28 Table. Criteria with no admitted phenotype in any genotype or in no curated model

The last column gives the number of curated models credited against the number of background genotypes credited. Candidates are the pairs retrieval produced for the criterion’s anchors, and judged are those routed to the LLM. One of the four admitted NCD B pairs is dropped at aggregation by the rule for added descendant terms (main text, Methods), so three pairs on two terms credit the criterion.

| Criterion | Candidates retrieved | Judged | Admitted | Curated vs background genotypes credited |
| --- | --- | --- | --- | --- |
| SCZ A1 to A3 (delusions, hallucinations, disorganized speech) | 1,233 | 1,137 | 0 | 0 vs 0 |
| MDD A7 (worthlessness or guilt) | 112 | 15 | 0 | 0 vs 0 |
| MDD A9 (thoughts of death) | 76 | 10 | 0 | 0 vs 0 |
| ASD A2 (nonverbal communication) | no mouse vocabulary | 0 | 0 | 0 vs 0 |
| MDD A5 (psychomotor change) | 260 | 51 | 14 | 0 vs 1,859 |
| MDD A6 (fatigue) | 109 | 24 | 4 | 0 vs 132 |
| MDD A8 (impaired concentration) | 76 | 13 | 1 | 0 vs 207 |
| NCD B (interference with independence) | 1,183 | 837 | 4 | 0 vs 251 |

##### S2 Fig. Agreement with the curated labels under both measures

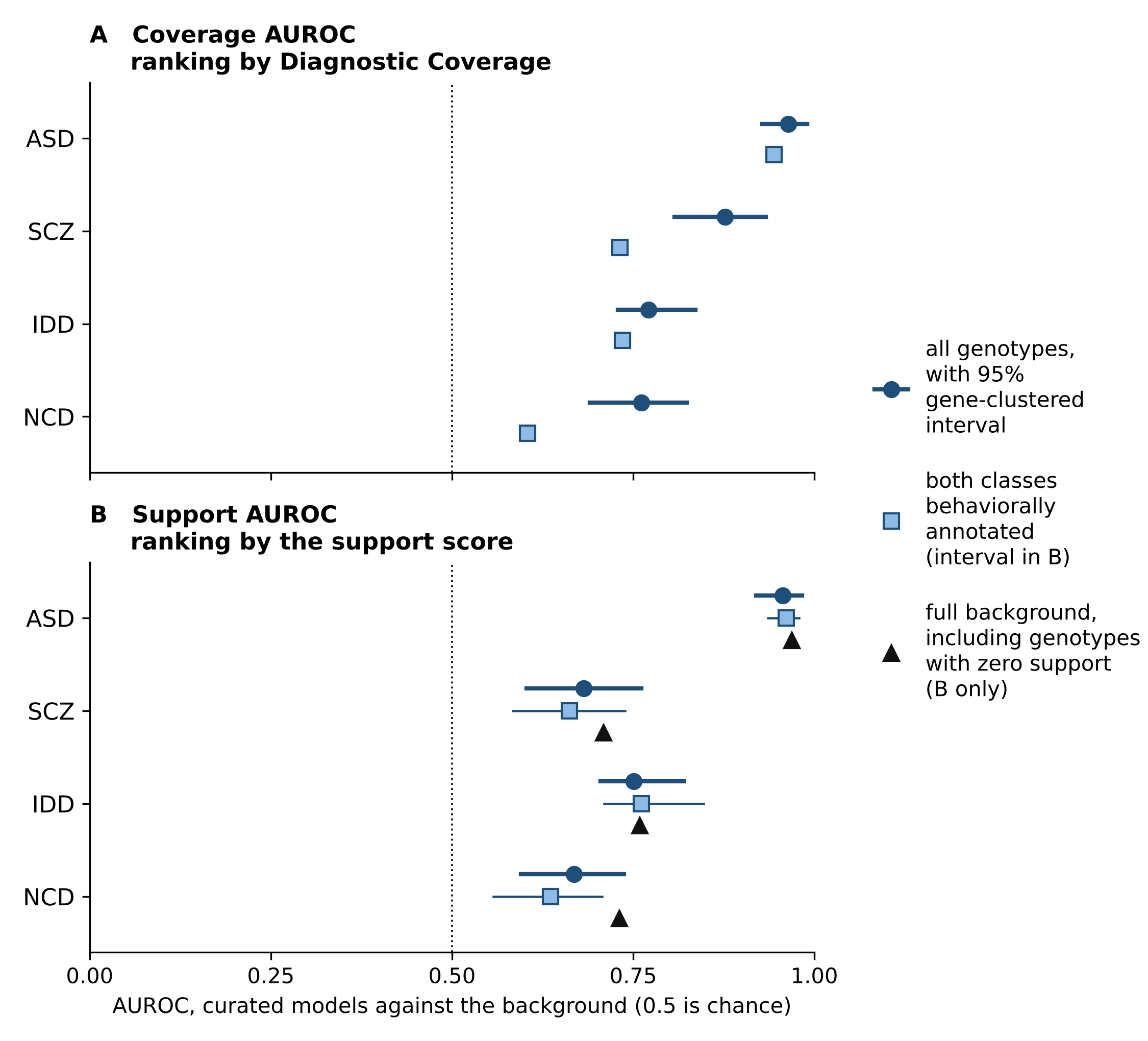

(A) Coverage AUROC ranks the genotype set by Diagnostic Coverage. (B) Support AUROC ranks the genotypes carrying a candidate term by the support score. Circles carry 95% gene-clustered intervals. Squares restrict both classes to behaviorally annotated genotypes, as point estimates in A and with intervals in B. Triangles use the full background, including genotypes with zero support (S8 Table).

##### Human-genetics genes and the IMPC-only test (moved from the main text)

Uncurated genotypes of genes implicated by human genetics without mouse data (H genes, which the pipeline never reads) reach the top coverage tier more often than other uncurated genotypes for ASD and IDD, partly through phenotyping attention. The papers behind the top uncurated ASD candidates discuss autism far more often than matched controls (S1 Text, S14 Table).

**Where phenotyping is uniform and tied to no claim, neither method finds the human-genetics genes.** The International Mouse Phenotyping Consortium (IMPC) phenotypes knockout genotypes by one fixed battery, and 5,126 genotypes of the set carry an IMPC allele, almost none of them curated. Within the IMPC genotypes of each evaluation set, with curated models removed, the genotypes of H genes rank against the rest at a support AUROC of 0.48 to 0.52. PhenoDigm recomputed and the behavioral annotation count are also at chance (S17 and S24 Tables, main text Table 3 for the primary values). The test is weak by construction: the positives are few (8 for SCZ), the H-gene prior is per gene and not per genotype, and the IMPC adult battery carries almost no social or cognitive assay. It shows only that in the one uniformly phenotyped set, no score finds a signal, not that none exists.

##### SM13. LLM admission against retrieval alone on the rated pairs

Whether the LLM verdict agrees with the experts better than text matching does was tested on the 140 rated pairs of the four primary disorders (2026-11-08, rerun at eight raters on 2026-11-17, no LLM call). The primary LLM admits 71 of them. Each retrieval rule admits the 71 pairs with the highest score, pooled across disorders or with each disorder’s own count, and every rule is scored against the rater majority of composite ratings with ties counted half. The retrieval score is the combined retrieval score of the primary analysis run, which combines the embedding, lexical and ontology channels with no LLM verdict, and embedding similarity is its embedding channel alone. Differences are paired, with a pair bootstrap of 4,000 replicates.

##### S29 Table. LLM admission against retrieval alone on the rated pairs, scored against the rater majority

| Rule (71 pairs admitted) | Agreement with raters | Kappa | Confirmation of admitted | Rejection of refused | Recall of rater-admitted | LLM minus rule, agreement (95% interval) |
| --- | --- | --- | --- | --- | --- | --- |
| LLM primary admission | 0.793 | 0.635 | 0.662 | 0.906 | 0.879 |  |
| Embedding similarity alone, pooled | 0.743 | 0.507 | 0.606 | 0.848 | 0.804 | +0.050 (-0.018 to +0.121) |
| Embedding similarity alone, within disorder | 0.746 | 0.521 | 0.613 | 0.855 | 0.813 | +0.046 (-0.021 to +0.114) |
| Retrieval score, pooled | 0.729 | 0.474 | 0.592 | 0.833 | 0.785 | +0.064 (-0.004 to +0.136) |
| Retrieval score, within disorder | 0.721 | 0.446 | 0.577 | 0.819 | 0.766 | +0.071 (+0.000 to +0.139) |

Without a cutoff, against the rater majority over the 123 pairs without a tie, the AUROC is 0.835 for the LLM verdict, 0.803 for the retrieval score, 0.811 for embedding similarity and 0.860 for the final pair score (retrieval plus the LLM’s adjustment). Within the pairs the LLM admitted, the retrieval score ranks rater-confirmed above rater-rejected pairs at 0.559, and within the pairs it refused at 0.656. The instrument was drawn by stratum from the record’s verdicts, and the retrieval score separates the record’s admitted from refused rated pairs at 0.859, so many sampled pairs are clear cases that both rules decide alike and the comparison is not neutral between them. Source: audit/retrieval_vs_llm_pairs_2026-11-08/ (script and results, aggregates only).

##### SM14. The judgment floor raised on the primary analysis run

The arms were built from the record’s pair table (the copy that reproduced the record through stages 3 and 4 on 2026-10-20) by un-judging every pair that reached the LLM by the floor route with a combined retrieval score below the new floor. Its verdict fields were emptied, its adjustment set to zero and its final score set to the combined retrieval score, as for every unjudged primary pair. Bundle pairs and injected descendants were left as judged. Each arm then ran through stages 3 and 4 with the commands of the alternate-LLM arms, and its support AUROC was compared with the record by a gene-clustered paired bootstrap (2,000 replicates, seed 20261008) against the pre-specified margin of 0.05, on the primary labels. A criterion match (criterion unit, admitted MP term) of the catalogue is lost when every admitted pair behind it is un-judged. The coverage AUROC was not recomputed. Source: audit/floor_sweep_record_2026-11-08/.

##### S30 Table. The judgment floor raised on the primary analysis run: admitted pairs and criterion matches lost, and support AUROC

Support AUROC in the evaluation set, with the difference from the record (floor 0.57) and its 90% gene-clustered interval. Equivalent: the interval lies inside -0.05 to 0.05. Inconclusive: it crosses a margin bound.

| Disorder | Admitted primary pairs | Lost at 0.60 | Lost at 0.65 | Criterion matches lost at 0.60, 0.65 | Support AUROC, record | Floor 0.60 | Floor 0.65 |
| --- | --- | --- | --- | --- | --- | --- | --- |
| ASD | 145 | 6 | 20 | 2, 8 | 0.956 | 0.956 (+0.000, -0.000 to +0.001), equivalent | 0.955 (-0.000, -0.001 to +0.000), equivalent |
| SCZ | 196 | 6 | 26 | 0, 6 | 0.682 | 0.682 (+0.000, +0.000 to +0.000), equivalent | 0.605 (-0.078, -0.125 to -0.032), inconclusive |
| IDD | 138 | 7 | 17 | 0, 1 | 0.751 | 0.751 (-0.000, -0.001 to +0.000), equivalent | 0.730 (-0.021, -0.048 to -0.005), equivalent |
| NCD | 95 | 14 | 17 | 4, 6 | 0.668 | 0.646 (-0.021, -0.045 to -0.002), equivalent | 0.623 (-0.045, -0.075 to -0.015), inconclusive |

On the full pool the differences are the same within 0.005 (SCZ -0.081 and NCD -0.042 at 0.65). The SCZ and NCD falls exclude zero, so a higher floor removes admissions that separate their curated models from the background, and the sweep cannot tell whether those admissions are analogs or label-correlated matches. Unjudged floor-route pairs a lower floor would add: 80, 57, 51 and 40 at 0.55 and 333, 257, 165 and 234 at 0.50 (ASD, SCZ, IDD, NCD).

##### SM15. The channel weights changed on the primary analysis run

The combined retrieval score is a weighted mean of the normalized embedding, lexical, Resnik and Jaccard channels, renormalized per pair over the channels present, with the lexical weight halved when both ontology channels are absent. The weights enter the record in four places. They set the combined retrieval score and the final pair score that the support score aggregates. They also decide the bundle route (the top five candidates of an anchor whose top two scores lie within 0.08 or whose top score lies in 0.35 to 0.70) and the fill route (a combined retrieval score of 0.57 or more outside a bundle). Injected descendants are judged whatever their score. Six arms were built from the record’s pair table (2026-11-09, no LLM call): equal weights without the lexical factor, the weights of record without the lexical factor and each channel left out. For each arm the score was recomputed from the stored normalized channels and both routes were re-derived. Every recorded verdict whose pair was still routed was kept. Every judged pair that lost its route was un-judged (verdict fields emptied, adjustment zero, final score equal to the combined retrieval score), and the pairs newly routed but never judged were counted. Replayed under the weights of record, the routing reproduced every bundle and every judged pair of the record in all four disorders. Each arm ran through stages 3 and 4 with the commands of the other arms. It was compared with the record by a gene-clustered paired bootstrap (2,000 replicates, seed 20261008) against the margin of 0.05, on the primary labels.

The arms hold the recorded verdicts fixed, so they bound the cost of a weighting and cannot show its gain. For the two cells that moved a support AUROC, the newly routed pairs were judged with the record’s bundle prompt and schema, the same LLM alias and seed, and the arm was rerun (the topped-up rows below). Of 219 pairs newly routed under equal weights for NCD, 4 were admitted, and of 68 newly routed for SCZ without the lexical channel, 11 were admitted. Source: audit/channel_weights_2026-11-08/.

##### S31 Table. The channel weights changed on the primary analysis run: admissions kept, pairs newly routed to the LLM and support AUROC

Evaluation set, primary labels. Admissions are strict admissions kept of the record’s count. The difference is arm minus record with its 90% gene-clustered interval. Equivalent: the interval lies inside -0.05 to 0.05. Not equivalent: it lies wholly outside. Inconclusive: it crosses a margin bound. Rows without a top-up hold the recorded verdicts fixed.

| Arm | ASD (145 admitted, 0.956) | SCZ (196, 0.682) | IDD (138, 0.751) | NCD (95, 0.668) |
| --- | --- | --- | --- | --- |
| Equal weights, no lexical factor | 121 kept, 198 newly routed, 0.955 (-0.001, -0.003 to +0.000), equivalent | 178, 193, 0.686 (+0.004, +0.003 to +0.005), equivalent | 128, 148, 0.751 (+0.000, -0.001 to +0.001), equivalent | 80, 219, 0.512 (-0.156, -0.205 to -0.101), not equivalent |
| Equal weights, newly routed pairs judged |  |  |  | 84 admitted, 0.532 (-0.136, -0.181 to -0.084), not equivalent |
| Weights of record, no lexical factor | 139, 80, 0.956 (-0.000, -0.001 to +0.000), equivalent | 185, 86, 0.685 (+0.003, +0.002 to +0.004), equivalent | 132, 59, 0.751 (-0.000, -0.001 to +0.001), equivalent | 84, 62, 0.651 (-0.017, -0.041 to +0.004), equivalent |
| Embedding left out | 109, 747, 0.954 (-0.002, -0.004 to -0.000), equivalent | 165, 409, 0.663 (-0.019, -0.052 to +0.006), inconclusive | 119, 257, 0.729 (-0.022, -0.049 to -0.003), equivalent | 79, 517, 0.513 (-0.155, -0.203 to -0.100), not equivalent |
| Lexical left out | 144, 88, 0.956 (-0.000, -0.000 to +0.000), equivalent | 190, 68, 0.615 (-0.067, -0.112 to -0.025), inconclusive | 135, 42, 0.751 (-0.000, -0.001 to +0.000), equivalent | 89, 39, 0.671 (+0.003, +0.001 to +0.005), equivalent |
| Lexical left out, newly routed pairs judged |  | 201 admitted, 0.623 (-0.059, -0.108 to -0.011), inconclusive |  |  |
| Resnik left out | 145, 3, 0.956, equivalent | 195, 6, 0.682, equivalent | 138, 1, 0.751, equivalent | 95, 4, 0.668, equivalent |
| Jaccard left out | 145, 8, 0.956, equivalent | 196, 1, 0.682, equivalent | 138, 2, 0.751, equivalent | 95, 2, 0.668, equivalent |

On the full pool every difference is within 0.015 of its primary value (NCD equal weights -0.170, topped up -0.149, SCZ lexical left out -0.070, topped up -0.059). The two ontology channels are absent for 97 to 99% of pairs, so leaving either out changes almost nothing. Under equal weights the NCD admissions lost are the broad terms that credit most curated NCD models (abnormal cognition, abnormal learning/memory/conditioning, impaired coordination and the apraxia route). They fall out of the bundle or below the floor when the lexical channel gains weight, and the newly routed pairs are mostly refused. Under lexical left out the six SCZ admissions lost are planning, declarative memory and social recognition terms, and the eleven new admissions do not restore the separation.

##### SM16. Precision and power of an AUROC by the number of curated models

A disorder is treated as primary at 30 or more curated models (Methods). The table gives the basis by the formula of Hanley and McNeil [41] for the coverage AUROC, whose positives are the curated models of record and whose negatives are the remaining genotypes of the 32,535. It reports the half-width of the 95% interval at an AUROC of 0.5 and of 0.7, and the smallest AUROC that a two-sided test at 0.05 detects against 0.5 with 80% power. The formula treats genotypes as independent. The intervals of the paper come from a gene-clustered bootstrap, which is wider where several curated models share a gene, so the table is a lower bound on the uncertainty. Source: audit/power_2026-11-18/ (2026-11-18, no LLM call).

##### S32 Table. Precision and power of an AUROC by the number of curated models

| Disorder (curated models) | Background genotypes | Half-width of the 95% interval at AUROC 0.5 | Half-width at AUROC 0.7 | Smallest AUROC detected with 80% power |
| --- | --- | --- | --- | --- |
| ASD (85) | 32,450 | 0.061 | 0.063 | 0.589 |
| SCZ (42) | 32,493 | 0.087 | 0.089 | 0.626 |
| IDD (121) | 32,414 | 0.052 | 0.052 | 0.574 |
| NCD (71) | 32,464 | 0.067 | 0.068 | 0.597 |
| MDD (8) | 32,527 | 0.200 | 0.204 | 0.782 |
| Threshold for a primary disorder (30) | 32,505 | 0.103 | 0.105 | 0.649 |

##### S33 Table. The pairs the alternate LLMs admitted on SCZ A1 to A3

The primary LLM admitted no pair on SCZ A1 to A3. The table lists every pair that an alternate LLM admitted there and the primary LLM refused: 5 for claude-opus-4-8 and 41 for gemini-2.5-pro, the 5 being among the 41. None is an analog of a delusion, a hallucination or disorganized speech. The mouse terms are endophenotype and cognitive terms (latent inhibition, startle, conditioning, learning, habituation), vocalization and stereotypy terms, eye-movement terms and brain-morphology terms. None of these pairs was expert-rated.

| LLM | Unit | HPO anchor | MP term | Analog grade |
| --- | --- | --- | --- | --- |
| claude-opus-4-8 | A1 | Delusion (HP:0000746) | abnormal latent inhibition of conditioning behavior (MP:0006299) | close |
| claude-opus-4-8 | A2 | Hallucinations (HP:0000738) | decreased startle reflex (MP:0001489) | close |
| claude-opus-4-8 | A3 | Disordered formal thought process (HP:0025769) | abnormal cognition (MP:0014114) | close |
| claude-opus-4-8 | A3 | Disordered formal thought process (HP:0025769) | abnormal latent inhibition of conditioning behavior (MP:0006299) | close |
| claude-opus-4-8 | A3 | Disordered formal thought process (HP:0025769) | decreased vocalization (MP:0020351) | close |
| gemini-2.5-pro | A1 | Delusion (HP:0000746) | abnormal associative learning (MP:0002062) | close |
| gemini-2.5-pro | A1 | Delusion (HP:0000746) | abnormal cognition (MP:0014114) | close |
| gemini-2.5-pro | A1 | Delusion (HP:0000746) | abnormal habituation to a novel odor (MP:0011622) | close |
| gemini-2.5-pro | A1 | Delusion (HP:0000746) | abnormal latent inhibition of conditioning behavior (MP:0006299) | direct |
| gemini-2.5-pro | A1 | Delusion (HP:0000746) | abnormal sensitization to xenobiotic (MP:0011624) | close |
| gemini-2.5-pro | A1 | Delusion (HP:0000746) | cognitive inflexibility (MP:0014115) | close |
| gemini-2.5-pro | A1 | Delusion (HP:0000746) | enhanced cued conditioning behavior (MP:0009455) | close |
| gemini-2.5-pro | A1 | Delusion (HP:0000746) | impaired conditioning behavior (MP:0012317) | close |
| gemini-2.5-pro | A1 | Delusion (HP:0000746) | impaired cued conditioning behavior (MP:0009456) | close |
| gemini-2.5-pro | A1 | Delusion (HP:0000746) | impaired learning (MP:0012315) | close |
| gemini-2.5-pro | A1 | Delusion (HP:0000746) | impaired spatial memory extinction (MP:0021088) | close |
| gemini-2.5-pro | A2 | Hallucinations (HP:0000738) | abnormal latent inhibition of conditioning behavior (MP:0006299) | close |
| gemini-2.5-pro | A2 | Hallucinations (HP:0000738) | abnormal startle reflex (MP:0001486) | close |
| gemini-2.5-pro | A2 | Hallucinations (HP:0000738) | absent startle reflex (MP:0006359) | close |
| gemini-2.5-pro | A2 | Hallucinations (HP:0000738) | decreased startle reflex (MP:0001489) | close |
| gemini-2.5-pro | A3 | Disordered formal thought process (HP:0025769) | abnormal Purkinje cell focal axonal swelling (MP:0014330) | close |
| gemini-2.5-pro | A3 | Disordered formal thought process (HP:0025769) | abnormal cognition (MP:0014114) | close |
| gemini-2.5-pro | A3 | Disordered formal thought process (HP:0025769) | abnormal eye blink conditioning behavior (MP:0002807) | close |
| gemini-2.5-pro | A3 | Disordered formal thought process (HP:0025769) | abnormal eye movement (MP:0001389) | close |
| gemini-2.5-pro | A3 | Disordered formal thought process (HP:0025769) | abnormal habituation (MP:0011617) | close |
| gemini-2.5-pro | A3 | Disordered formal thought process (HP:0025769) | abnormal habituation to a new environment (MP:0011620) | close |
| gemini-2.5-pro | A3 | Disordered formal thought process (HP:0025769) | abnormal inhibitory learning (MP:0020340) | close |
| gemini-2.5-pro | A3 | Disordered formal thought process (HP:0025769) | abnormal latent inhibition of conditioning behavior (MP:0006299) | close |
| gemini-2.5-pro | A3 | Disordered formal thought process (HP:0025769) | abnormal response to novelty (MP:0003107) | close |
| gemini-2.5-pro | A3 | Disordered formal thought process (HP:0025769) | abnormal short-term object recognition memory (MP:0002800) | close |
| gemini-2.5-pro | A3 | Disordered formal thought process (HP:0025769) | abnormal startle reflex (MP:0001486) | close |
| gemini-2.5-pro | A3 | Disordered formal thought process (HP:0025769) | abnormal temporal memory (MP:0001468) | close |
| gemini-2.5-pro | A3 | Disordered formal thought process (HP:0025769) | abnormal visual pursuit (MP:0006156) | close |
| gemini-2.5-pro | A3 | Disordered formal thought process (HP:0025769) | absent Cajal-Retzius cell (MP:0000344) | close |
| gemini-2.5-pro | A3 | Disordered formal thought process (HP:0025769) | absent cerebellum vermis (MP:0000865) | close |
| gemini-2.5-pro | A3 | Disordered formal thought process (HP:0025769) | absent corpus callosum (MP:0002196) | close |
| gemini-2.5-pro | A3 | Disordered formal thought process (HP:0025769) | absent vestibuloocular reflex (MP:0004845) | close |
| gemini-2.5-pro | A3 | Disordered formal thought process (HP:0025769) | cognitive inflexibility (MP:0014115) | close |
| gemini-2.5-pro | A3 | Disordered formal thought process (HP:0025769) | decreased startle reflex (MP:0001489) | close |
| gemini-2.5-pro | A3 | Disordered formal thought process (HP:0025769) | decreased vocalization (MP:0020351) | close |
| gemini-2.5-pro | A3 | Disordered formal thought process (HP:0025769) | ectopic Purkinje cell (MP:0000885) | close |
| gemini-2.5-pro | A3 | Disordered formal thought process (HP:0025769) | enlarged lateral ventricles (MP:0008535) | close |
| gemini-2.5-pro | A3 | Disordered formal thought process (HP:0025769) | excessive vocalization (MP:0013140) | close |
| gemini-2.5-pro | A3 | Disordered formal thought process (HP:0025769) | impaired discrimination learning (MP:0012296) | close |
| gemini-2.5-pro | A3 | Disordered formal thought process (HP:0025769) | increased stereotypic behavior (MP:0001409) | close |
| gemini-2.5-pro | A3 | Disordered formal thought process (HP:0025769) | stereotypic behavior (MP:0001408) | close |

#### Supplementary references

Alexandru A, Jagla W, Graubner S, et al. Selective hippocampal neurodegeneration in transgenic mice expressing small amounts of truncated Abeta is induced by pyroglutamate-Abeta formation. J Neurosci. 2011;31(36):12790-12801. doi:10.1523/JNEUROSCI.1794-11.2011

Amiel J, Rio M, de Pontual L, et al. Mutations in TCF4, encoding a class I basic helix-loop-helix transcription factor, are responsible for Pitt-Hopkins syndrome, a severe epileptic encephalopathy associated with autonomic dysfunction. Am J Hum Genet. 2007;80(5):988-993. doi:10.1086/515582

Cohen J. A coefficient of agreement for nominal scales. Educ Psychol Meas. 1960;20(1):37-46. doi:10.1177/001316446002000104

de Leeuw CA, Mooij JM, Heskes T, Posthuma D. MAGMA: generalized gene-set analysis of GWAS data. PLoS Comput Biol. 2015;11(4):e1004219. doi:10.1371/journal.pcbi.1004219

Fleiss JL. Measuring nominal scale agreement among many raters. Psychol Bull. 1971;76(5):378-382. doi:10.1037/h0031619

Fromer M, Pocklington AJ, Kavanagh DH, et al. De novo mutations in schizophrenia implicate synaptic networks. Nature. 2014;506(7487):179-184. doi:10.1038/nature12929

Fu JM, Satterstrom FK, Peng M, et al. Rare coding variation provides insight into the genetic architecture and phenotypic context of autism. Nat Genet. 2022;54(9):1320-1331. doi:10.1038/s41588-022-01104-0

Gamache J, Benzow K, Forster C, et al. Factors other than hTau overexpression that contribute to tauopathy-like phenotype in rTg4510 mice. Nat Commun. 2019;10:2479. doi:10.1038/s41467-019-10428-1

Gulbranson DR, Ho K, Yu GQ, et al. Phenotypic differences between the Alzheimer’s disease-related hAPP-J20 model and heterozygous Zbtb20 knock-out mice. eNeuro. 2021;8(3):ENEURO.0089-21.2021. doi:10.1523/ENEURO.0089-21.2021

Hamdan FF, Daoud H, Piton A, et al. De novo SYNGAP1 mutations in nonsyndromic intellectual disability and autism. Biol Psychiatry. 2011;69(9):898-901. doi:10.1016/j.biopsych.2010.11.015

Hamdan FF, Gauthier J, Spiegelman D, et al. Mutations in SYNGAP1 in autosomal nonsyndromic mental retardation. N Engl J Med. 2009;360(6):599-605. doi:10.1056/NEJMoa0805392

Holm S. A simple sequentially rejective multiple test procedure. Scand J Stat. 1979;6(2):65-70. [no DOI; JSTOR 4615733]

Jaccard P. The distribution of the flora in the alpine zone. New Phytol. 1912;11(2):37-50. doi:10.1111/j.1469-8137.1912.tb05611.x

Nuytens K, Tuand K, Fu Q, et al. The dwarf phenotype in GH240B mice, haploinsufficient for the autism candidate gene neurobeachin, is caused by ectopic expression of recombinant human growth hormone. PLoS One. 2014;9(10):e109598. doi:10.1371/journal.pone.0109598

Satterstrom FK, Kosmicki JA, Wang J, et al. Large-scale exome sequencing study implicates both developmental and functional changes in the neurobiology of autism. Cell. 2020;180(3):568-584.e23. doi:10.1016/j.cell.2019.12.036

Singh T, Poterba T, Curtis D, et al. Rare coding variants in ten genes confer substantial risk for schizophrenia. Nature. 2022;604(7906):509-516. doi:10.1038/s41586-022-04556-w

Smedley D, Oellrich A, Kohler S, et al. PhenoDigm: analyzing curated annotations to associate animal models with human diseases. Database (Oxford). 2013;2013:bat025. doi:10.1093/database/bat025

Su Y, Balice-Gordon RJ, Hess DM, et al. Neurobeachin is essential for neuromuscular synaptic transmission. J Neurosci. 2004;24(14):3627-3636. doi:10.1523/JNEUROSCI.4644-03.2004

Vernay A, Therreau L, Blot B, et al. A transgenic mouse expressing CHMP2B intron 5 mutant in neurons develops histological and behavioural features of amyotrophic lateral sclerosis and frontotemporal dementia. Hum Mol Genet. 2016;25(15):3341-3360. doi:10.1093/hmg/ddw182

Wilson EB. Probable inference, the law of succession, and statistical inference. J Am Stat Assoc. 1927;22(158):209-212. doi:10.1080/01621459.1927.10502953

Zweier C, Peippo MM, Hoyer J, et al. Haploinsufficiency of TCF4 causes syndromal mental retardation with intermittent hyperventilation (Pitt-Hopkins syndrome). Am J Hum Genet. 2007;80(5):994-1001. doi:10.1086/515583
